# The reliability and heritability of cortical folds and their genetic correlations across hemispheres

**DOI:** 10.1101/795591

**Authors:** Fabrizio Pizzagalli, Guillaume Auzias, Qifan Yang, Samuel R. Mathias, Joshua Faskowitz, Joshua Boyd, Armand Amini, Denis Rivière, Katie L. McMahon, Greig I. de Zubicaray, Nicholas G. Martin, Jean-François Mangin, David C. Glahn, John Blangero, Margaret J. Wright, Paul M. Thompson, Peter Kochunov, Neda Jahanshad

**Affiliations:** Imaging Genetics Center, Mark and Mary Stevens Neuroimaging and Informatics Institute, Keck School of Medicine of USC, Marina del Rey, CA, USA; Institut de Neurosciences de la Timone, UMR7289, Aix-Marseille Université & CNRS, Marseille, France; Laboratoire des Sciences de l’Information et des Systèmes, UMR7296, Aix-Marseille Université & CNRS, Marseille, France; Department of Psychiatry, Boston Children’s Hospital and Harvard Medical School, Boston, MA, USA & Yale University School of Medicine, CT, USA; Olin Neuropsychiatry Research Center, Institute of Living, Hartford Hospital, USA; UNATI Lab, Neurospin, CEA, Paris Saclay University, Gif sur Yvette, France; CATI, Multicenter Neuroimaging Platform, Paris, France; School of Clinical Sciences and Institute of Health and Biomedical Innovation, Queensland University of Technology, Brisbane, QLD, 4000, Australia; Faculty of Health and Institute of Health and Biomedical Innovation, Queensland University of Technology (QUT), Brisbane, QLD 4059, Australia; QIMR Berghofer Medical Research Institute, Brisbane, QLD, Australia; South Texas Diabetes and Obesity Institute, University of Texas Rio Grande Valley School of Medicine, Brownsville, TX, USA; Queensland Brain Institute, University of Queensland, Brisbane, QLD 4072, Australia; Maryland Psychiatric Research Center, Department of Psychiatry, Univ. of Maryland School of Medicine, Baltimore, MD, USA

**Keywords:** Cortical folding, Heritability, Brain Imaging, Sulci Morphometry, Genetic Asymmetry, Imaging Genetics, Reliability

## Abstract

The structure of the brain’s cortical folds varies considerably in human populations. Specific patterns of cortical variation arise with development and aging, and cortical traits are partially influenced by genetic factors. The degree to which genetic factors affect cortical folding patterning remains unknown, yet may be estimated with large-scale *in-vivo* brain MRI. Using multiple MRI datasets from around the world, we estimated the reliability and heritability of sulcal morphometric characteristics including length, depth, width, and surface area, for 61 sulci per hemisphere of the human brain. Reliability was assessed across four distinct test-retest datasets. We meta-analyzed the heritability across three independent family-based cohorts (N > 3,000), and one cohort of largely unrelated individuals (N~9,000) to examine the robustness of our findings. Reliability was high (interquartile range for ICC: 0.65−0.85) for sulcal metrics. Most sulcal measures were moderately to highly heritable (heritability estimates = 0.3−0.7). These genetic influences vary regionally, with the earlier forming sulci having higher heritability estimates. The central sulcus, the subcallosal and the collateral fissure were the most highly heritable regions. For some frontal and temporal sulci, left and right genetic influences did not completely overlap, suggesting some lateralization of genetic effects on the cortex.

## Introduction

Understanding the mechanisms underlying brain structural and functional variations is essential for advancing neuroscience. Genetic drivers of brain differences are important to identify as potential risk factors for heritable brain diseases, and targets for their treatment. Large-scale neuroimaging consortia, including the Enhancing Neuro Imaging Genetics through Meta Analysis (ENIGMA^1^) consortium, have identified common genetic variants that have small but significant associations with variations in brain morphology^2^. The same brain regions show, on average, consistent structural abnormalities in populations of individuals diagnosed with neuropsychiatric or neurodevelopmental disorders, including schizophrenia^3^, bipolar disorder^4^, major depressive disorder (MDD)^5^, obsessive-compulsive disorder (OCD)^6^, ADHD^7^, and autism spectrum disorder (ASD)^8^. Studies have even identified genetic correlations between the human brain structure and risk for disease^9–11^.

Enriched in neuronal cell bodies, the brain’s cortical gray matter is involved in almost all human cognitive functions and behavior, including sensory perception and motor control^12^. Macroscale anatomical features of the human cortex can be reliably extracted from structural MRI scans, and among the most common are regional thickness and surface area measures. These MRI-based features show robust alterations in several neurological, neurodevelopmental, and psychiatric disorders^5^, influenced by both environmental and genetic variation^13–15^.

Gyrification, or folding of the cortical surface, takes place during brain development, forming sulci (“fissures”) and gyri (“ridges”) in the cortical gray matter. Gyrification is among the fundamental features of human and non-human primate brain anatomy^16,17^ and occurs in an orchestrated pattern^17^ that changes during fetal life and into adolescence^18^. The mechanisms of brain folding are not fully understood^19,20^, but the process is largely preserved among humans and non-human primates. The pattern of brain gyri (or sulci) tend to delimit areas with specific functions and are generally consistent across subjects^21–24^. The complexity and inter-subject variability of brain gyrification are influenced by many factors, including genetically and environmentally influenced developmental, aging and pathological processes^25,26^.

Large-scale imaging studies have aimed to discover both common and rare genetic variants that contribute to brain variability as estimated using *in-vivo* brain scans, such as MRI^27^; genome-wide association studies (GWAS) find that, as with other complex traits, individual common variants typically explain less than 1% of the population variance, despite accounting for a large fraction of the variance in aggregate^2,10,28,29^. Therefore, successful efforts to discover common variants that affect cortical structure require tens of thousands of scans, as well as independent samples for replication and generalization. Large-scale biobanks have amassed tens of thousands of MRI scans^30^. Even so, replicating effects and ensuring generalizability of findings to other scanned populations, requires assurance that the brain measures being studied are reliably extracted across a variety of possible MRI scanning paradigms. Features must also be reliably extracted if effects are to be pooled across studies as in multisite consortia such as ENIGMA and CHARGE^1^.

*Sulcal-based morphometry* provides in-depth analyses of the brain’s cortical fissures, or folds, as seen on MRI. Measures of sulcal morphometry - including length, depth, width and surface area - among others - have been associated with developmental maturation in adolescents^31^, degenerative changes in the elderly^31,32^, and disorders such as schizophrenia^33,34^, bipolar disorder^35^ and autism spectrum disorder^36^; altered fissuration is also found in several genetic disorders, such as Williams syndrome^37,38^ where abnormal cortical patterns have known genetic influences. The depth of the central sulcus has been reported to be highly heritable, and the degree of this heritability varies along the profile of the central sulcus^39^. Effects on sulcal patterns are partially independent of those on cortical thickness or surface area^14,31^. Here we set out to understand the degree to which these additional, more in-depth, cortical features may be influenced by genetic factors, yet before studying the heritability of these features, a multisite effort is needed to assess the reliability of measures derived from the automatic extraction of these sulci from MRI.

Here, we perform an extensive reliability (N=110) and heritability (N=13,113) analysis. Reliability was estimated in data from four cohorts, totalling 110 participants (19-61 years of age, 47% females on average) who underwent a T1-weighted brain MRI scan twice within three months. A subset of data from the Human Connectome Project (HCP) and the Queensland Twin Imaging Study (QTIM) were used along with the Kennedy Krieger Institute - Multi-Modal MRI Reproducibility Resource (KKI) and the Open Access Series of Imaging Studies (OASIS) datasets for reproducibility analysis. We included datasets for which we would expect minimal or no structural changes between scans, so we limited the analysis to healthy individuals between the ages of 18 and 65, and ensured the inter-scan interval was less than 90 days. See **Table S2** in Supplementary Material for more details.

We analyzed heritability in four independent cohorts, three with a family based design and one using single-nucleotide polymorphism (SNP) based heritability estimates. The cohorts included two twin-based samples (QTIM and HCP), one cohort of extended pedigrees (the Genetics of Brain Structure and Function; GOBS), and another of over 9,000 largely unrelated individuals (the UK Biobank). Heritability estimates are population specific, but here, our aim was to understand the heritability pattern across populations and demonstrate that genetic effects are consistently observed. We pooled information from all family based cohorts to estimate the generalized heritability values using meta-and mega-analytic methods^40,41^.

We estimated reliability and heritability for measures of each sulcus on the left and right hemisphere separately. As there is limited evidence for genetic lateralization across most of the human brain^42–44^. we also evaluated the heritability estimates of the measures for each sulcus averaged across the two hemispheres. This may lead to more stable measurements and, if the bilateral measures are influenced by similar genetic factors, then more stable measures could lead to better powered genetic studies. We also assessed the degree of genetic correlation between the measures across hemispheres. Sulci with limited genetic correlations between hemispheres may reveal novel insight into the brain’s lateralization and identify key biomarkers for relating lateralized traits, such as language and handedness, to brain structure^45^.

## Results

### Measurement reliability and its relationship to heritability

**Table SA** in the **Supplementary Materials**, reports the sulcal nomenclature, including both the abbreviations and full name for each sulcus. Reliability estimates can be found in **Supplementary Tables S3-S5** for ICC and **Supplementary Tables S6-S9** for the bias evaluation; heritability estimates are reported in **Supplementary Tables S10-S21** for the univariate analysis and **Supplementary Tables S22-S25** for the bivariate analysis. We summarize the results below.

We evaluated the reliability of the sulcal measurements using two indices: (1) the intraclass correlation coefficient (ICC^46,47^, equation 5), in both left and right hemispheres and after averaging bilateral measures; (2) the bias^48^ (*b, equation 4*) computed as the difference between the sulcal shape measures for the two scans divided by their averaged value. This index will give an estimate of consistency across subjects for each cohort.

### Intraclass correlation (ICC)

Overall, the meta-analysis showed an ICC interquartile range of 0.62 to 0.86 with the length as least robust global metric tested across the full brain (**Table 1**, **Figure 1-B**). Averaging left and right hemispheres improved the reliability across the brain for individual sulci;a higher fraction of sulci reached an acceptable ICC > 0.75, for all the descriptors (**Figure 1-B**).

**Figure 1:**
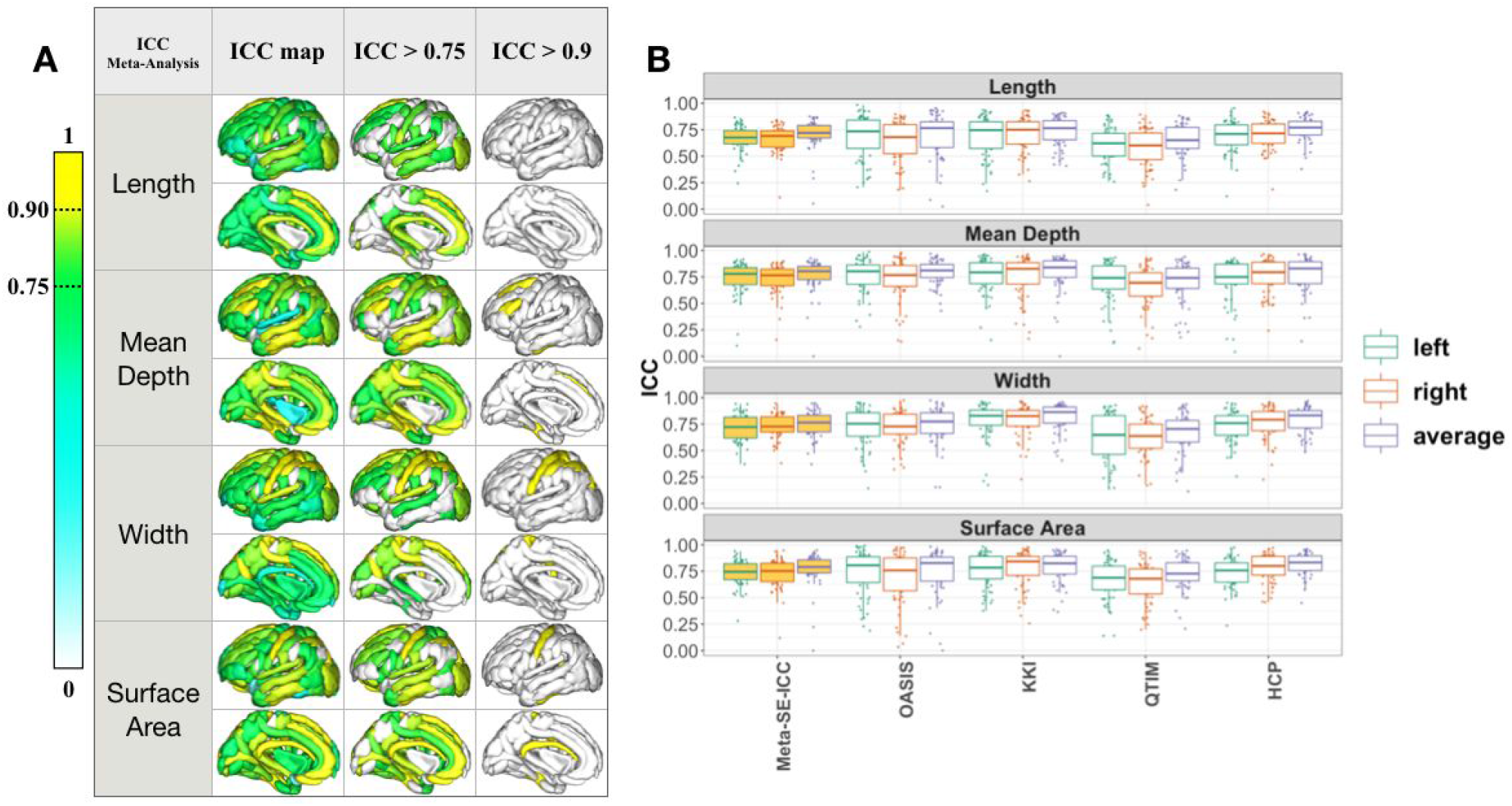
A) Sulcal-based meta-analysis of intraclass correlation (ICC) for bilaterally averaged sulcal measures. **Sulcal length** shows generally ‘good’ reproducibility, although no regions show ICC > 0.9^47^. **Mean depth** shows ‘excellent’ reproducibility (ICC > 0.9) for: the inferior frontal sulcus (S.F.inf.) and the superior frontal sulcus (S.F.sup.); **Sulcal width** shows ‘excellent’ reproducibility for: intraparietal sulcus (F.I.P.), superior postcentral intraparietal superior sulcus (F.I.P.Po.C.inf.), central sulcus (S.C.), superior postcentral sulcus (S.Po.C.sup.). **Surface area** shows ‘excellent’ reproducibility for: the central sulcus (S.C.), subcallosal sulcus (S.Call.), anterior occipito-temporal lateral sulcus (S.O.T.lat.ant.). B) The intra-class correlation (ICC) for **left, right** and **bilaterally averaged** sulcal length, mean depth, width, and surface area across the whole brain is plotted for four test-retest cohorts as well as the meta-analysis across them KKI shows the highest ICC across sulci.

**Table 1:** Meta-analysis of ICC estimated from 4 independent cohorts for sulcal length, mean depth, width, and surface. Left and right hemisphere and bilaterally averaged Mean ± SD are reported with ICC interquartile range [25% - 75%] across sulci.

| Meta-Analysis | Length | Mean Depth | Width | Surface Area |
| --- | --- | --- | --- | --- |
| Left | 0.66 ± 0.15<br>[0.61 - 0.76] | 0.72 ± 0.18<br>[0.68 - 0.83] | 0.71 ± 0.14<br>[0.62 - 0.82] | 0.73 ± 0.13<br>[0.66 - 0.82] |
| Right | 0.65 ± 0.16<br>[0.59 - 0.74] | 0.73 ± 0.14<br>[0.66 - 0.82] | 0.71 ± 0.14<br>[0.64 - 0.81] | 0.73 ± 0.13<br>[0.67 - 0.81] |
| Average | 0.70 ± 0.17<br>[0.66-0.79] | 0.77 ± 0.15<br>[0.73-0.86] | 0.76 ± 0.11<br>[0.69-0.83] | 0.76 ± 0.17<br>[0.72 - 0.86] |

The meta-analysis of ICC captures the variability in the reliability across cohorts for each sulcus (**Sup. Figure S1, Figure S2 and Figure S3**). Reliability measures depend to some extent on the cohort examined, or the scanning acquisition parameters. For example, for QTIM, which was collected at 4 tesla, the ICC can be classified as “good” (ICC > 0.75) for the left sulcal surface area of the *collateral sulcus* (*F.Coll*.), but “poor or moderate” (ICC<0.75) in OASIS for the same trait. Figure 1-A shows the meta-analysis of ICC across the 4 cohorts. We also show the patterns for “good” (ICC >0.75) and “excellent” (ICC>0.9) reliability.

For a detailed breakdown of the ICC for measures of sulci morphometry per cohort, please see **Sup. Figure S1** for the left hemisphere, **Sup. Figure S2**, for the right, and **Sup. Figure S3** for bilaterally averaged measures.

For the complete meta-analyzed ICC results, please see **Supp. Figures S4-S7**, for length, depth, width and surface area respectively, all of which are tabulated in **Supp. Tables S5**.

For each sulcus, we averaged the reliability estimates across all 4 sulcal descriptors to find the most reliable sulci overall. The *central sulcus (S.C.)* gave the most reliable sulcal measures, followed by the *median frontal sulcus* (S.F.median.), the *intraparietal sulcus* (F.I.P.), the *occipito-temporal lateral sulcus* (S.O.T.lat.ant.), the sylvian sulcus (S.C.sylvian), the *sub-parietal sulcus* (S.s.P.), the *occipital lobe*, and the *superior temporal sulcus* (S.T.s.) (**Supp. Figure S8**).

### Bias (*b*)

We explored test-retest consistency in terms of the ‘bias’ (*b*, equation 4), with Bland-Altman analyses. As in^48^, overall the bias values showed high test-retest consistency of sulcal shape measures (**Sup. Table S6**). Bias values greater than or equal to 0.1 are considered high, and were noted mainly for length estimates - e.g. for the length of the left and right *anterior/posterior sub-central ramus of the lateral fissure* (F.C.L.r.sc.ant./post.), and the length of the left and right insula (See **Supp. Table S7-S9** for bias estimates across the left, right and bilaterally averaged sulcal metrics). Paralleling the higher ICC in bilaterally averaged measures, lower ‘bias’ estimates were obtained with individual sulcal measures averaged across the left and right hemispheres (**Supp. Table S9**).

ICC and bias *b* of bilaterally averaged sulcal metrics were significantly negatively correlated for all metrics except for length, in particular *r_length_*=-0.11 [*p*=0.07], *r_mean-depth_*=−0.14 [p=0.02], r_*width*_=−0.25 [p=4.6×10^−5^], *r_surface-area_*=−0.25 [*p*=1.2×10^−5^]’ suggesting, as expected, that a lower bias between test and retest measurements relates to higher reproducibility as estimated by ICC.

### Heritability estimates for the brain’s cortical folding patterns

The profile of heritability (*h^2^*) estimates were calculated for bilaterally averaged sulcal measures of length, mean depth, width and surface area for the three family based cohorts. Heritabilities of all measures were estimated after adjusting for total intracranial volume (ICV), age, age^2^, sex and the interactions between sex and age and age^2^. Across descriptors and sulci, heritability estimates showed a similar pattern across the three family based cohorts, QTIM, HCP, GOBS (**Supp. Figure S9**, **Supp. Figure S10**, **Supp. Figure S11;** the GOBS cohort shows lower heritability, (h2 = 0.3±0.1), compared to QTIM (h2 = 0.4±0.1) and HCP (h2 = 0.4±0.1); GOBS is a cohort with an extended pedigree design and a wide age range (18-85 years of age), while both HCP and QTIM are twin based cohorts of young adults aged 25-35 years and 20-30 years, respectively.

The generalized heritability profile of cortical folding was obtained by meta-analyzing the estimates across these three independent family-design cohorts, and is highlighted in **Figure 2a**. Aggregate heritability estimates were also calculated in a mega-analytic manner, where 3,030 subjects from the family-based cohorts (QTIM, HCP and GOBS) were pooled together (after adjusting for covariates within cohort and normalizing across cohorts) before computing heritability as in previous works^40,41^. As expected, we found similarities between meta and mega-analysis derived heritability estimates as indicated by a significant Pearson’s correlation between these two approaches (r~0.84, p=10^−3^−10^−8^; **Supplementary Figure S13**). Individual heritability estimates, standard errors and p-values for bilaterally averaged sulcal length, mean depth, width and surface area are tabulated in **Supplementary Tables S10-S19** for each cohort, and in **Supplementary Table S20-S21** for the meta-and mega-analyses.

**Figure 2.**
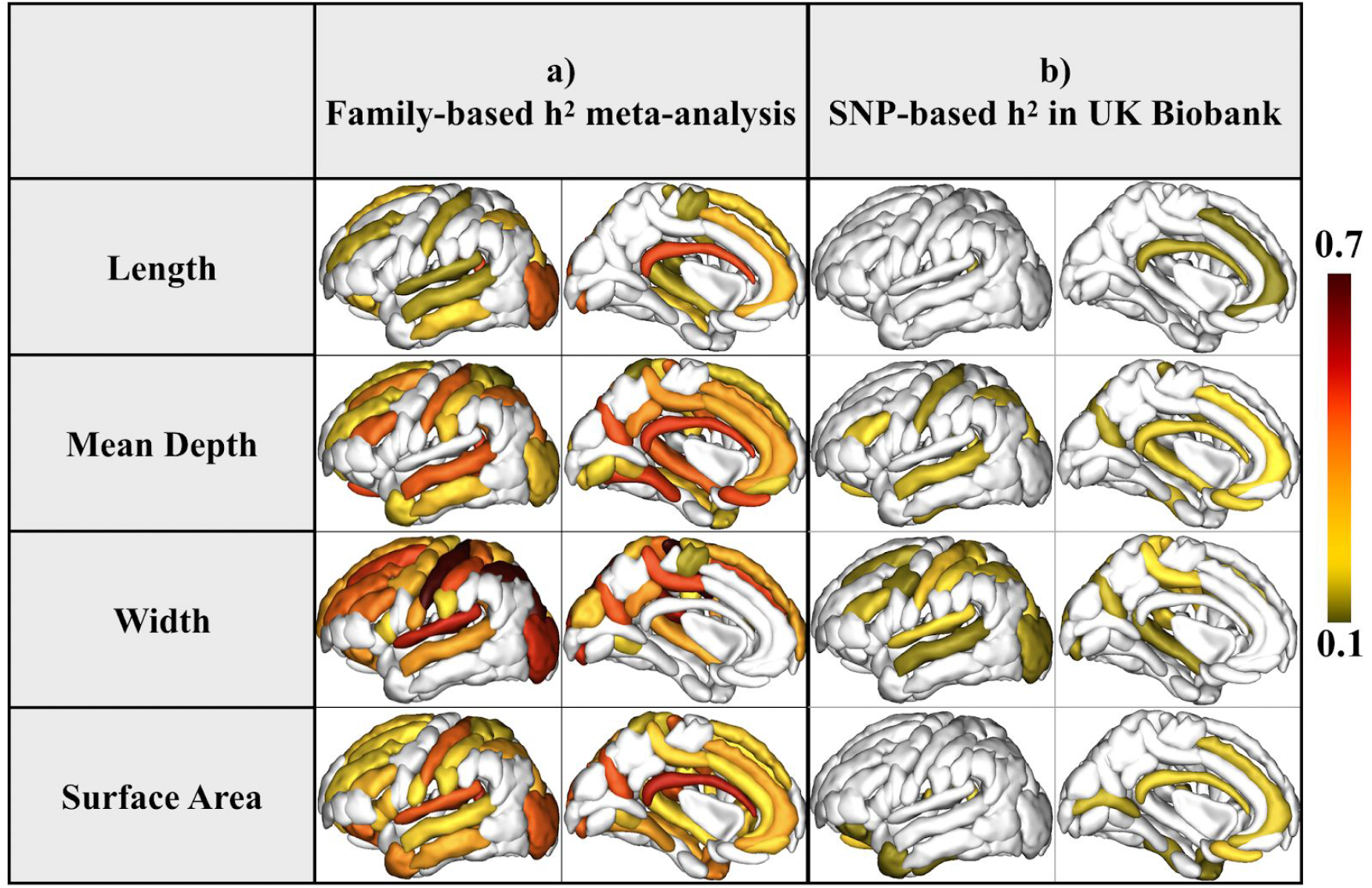
Heritability estimates (*h^2^*) for each bilaterally averaged sulcal descriptor are mapped. a) The results of the inverse-variance weighted meta-analysis of the heritability estimates across three family based cohorts QTIM, HCP, and GOBS highlight an overall heritability profile across 3,030 individuals. b) Heritability estimates (*h^2^*) calculated from sulcal features extracted from MRI scans of 10,083 unrelated individuals scanned as part of the UK Biobank were calculated using the genome-wide complex trait analysis (GCTA) package. Regional sulcal metrics found to be significantly heritable in the large population sample largely overlap with those found to be most highly heritable across the family based studies. We highlight only regions that had significant heritability estimates in sulci that had an ICC > 0.75 (See **Supp Table S3-S5** for sulcal-based values of ICC). Significant regions survived Bonferroni correction for multiple comparisons across all bilateral traits and regions (p < 0.05/(61*4)); darker red colors indicate higher heritability estimates. The left hemisphere was used for visualization purposes.

Using genome-wide complex trait analysis (GCTA), we estimated the heritability of sulcal variation in unrelated individuals of European ancestry from the UK Biobank and found many sulcal features for which the SNP-based heritability estimates were approximately 25% of the estimates derived from the family based studies, (*h*^2^=0.2 ± 0.1; **Figure 2b**. The heritability estimates for the UK Biobank are reported in **Supplementary TableS19**)

Overall, the pattern of heritability estimates were largely coherent between the family based and large-scale population studies. Among the sulcal descriptors analyzed, the *width* was the most heritable measurement, while the length was the least, showing significant heritability estimates for only sparse regions of the cortex. The heritability of sulcal length was more frequently significant when not adjusting for ICV, yet we find minimal differences in the overall *h^2^* estimates for sulcal depth and width before and after covarying for ICV (**Supp. Figure S14**); this suggests that there are minimal independent genetic influences on sulcal length beyond the known genetic influences on ICV, which was used as a covariate in the results presented.

Meta-analyzed heritability and reliability were significantly correlated for the family-based cohorts: r=0.36 (pval=1×10^−7^) for sulcal length, r=0.31 (pval=4.1×10^−6^) for mean depth, r=0.26 (pval=7×10^−5^) for sulcal width and r=0.25 (pval=1×l0^−4^) for surface area (**Supp. Figure S15**); for UK Biobank heritability and reliability were correlated for mean depth (r=0.43, pval=2×10^−3^) and sulcal width (r=0.38, pval=4×10^−3^) (**Supp. Figure S16**),Few regions with “poor” reliability (ICC < 0.75) show significant heritability estimated with the meta analysis on bilaterally averaged measures. Among others, the length of the *parieto-occipital fissure* (F.P.O.) [ICC=0.66, h2=0.18 (pval=1×10^−5^)], the mean depth of the *ascending ramus of the lateral fissure* (F.C.L.r.asc.) [ICC=0.74, h2=0.2 (pval=2.2×10^−6^)], the surface area of the *anterior inferior frontal sulcus* (S.F.inf.ant.) [ICC=0.65, h2=0.17 (pval=4.7×10^−6^)] and the width of the *calloso-marginal ramus of the lateral fissure* (F.C.M.ant.) [ICC=0.63, h2=0.34 (pval=1×10^−16^)] (**Supp. Table S5** and **Supp. Table S20**). For UK Biobank, the length of S.T.pol. [ICC=0.70, h2=0.14 (pval=6×10^−5^)], the width and the surface area for the *insula*, [ICC=0.65, h2=0.14 (pval=2.6×10^−5^)] and [ICC=0.65, h2=0.l6 (pval=3.8×10^−6^)] respectively (**Supp. Table S19)**.

Across the brain, the sulcal measurements were significantly heritable across cohorts, in particular for sulcal width. For sulcal width we found higher univariate heritability for the right hemisphere (h^2^=0.23±0.08) compared to the left hemisphere (h^2^=0.2±0.08) (paired t-test: 1.6×10^−6^, dof=50).

The heritability estimates for the global measures (i.e. the sum across sulci) of sulcal length, mean depth, width and surface area (covarying for ICV, age, and sex variables) are also reported in **Supplementary Figure S17**. QTIM, HCP and GOBS show similar trend across descriptor and hemispheres, except QTIM which had generally higher heritability for sulci on the right hemisphere compared to those on the left.

Sulci that were significantly heritable across descriptors included the intraparietal sulcus, the occipital lobe, the subcallosal sulcus, the internal frontal sulcus, the orbital sulcus, the anterior inferior temporal sulcus and the polar temporal sulcus, among others; in total 15 sulci were significant in the meta analysis, and 19 for the mega analysis. (for the corresponding values of *h^2^* see **Supplementary Table S20-S21)**.

Even though fewer sulci show significantly heritable estimates for their lengths, the heritability estimates for those sulci are not necessarily low and may be higher than that of width for a given sulcus. For example, for the subcallosal sulcus, its length shows higher *h^2^* than its width: *h^2^* = 0.5 and 0.3 respectively (**Supplementary Table S20**). This suggests a more morphometric specific genetic signature, where regionally different measurements may pick up a genetic effect.

33% (36% for mega-analysis) of the total number of bilaterally averaged sulci showed significant *h^2^* for sulcal length, 57% (59% for mega-analysis) for mean depth, 67% (65% for mega-analysis) for width and 62% (60% for mega-analysis) for the surface area. 6 sulci were significantly heritable for only one of the 4 descriptors (1 for mega-analysis). No sulcus show significant heritability for length only.

### Lateralized sulcal heritability estimates

#### Genetic correlation across the hemispheres

Averaging brain-imaging derived traits across the left and right hemispheres, as above, has been shown to reduce noise due to measurement error in large scale, multi-cohort efforts^2,3,28,49^. Improvements in the signal-to-noise ratio may be essential for discovering single common variants that explain less than one percent of the overall variability in a trait. However, by assessing left and right separately, we may be able to discover lateralized genetic effects if they exist. We confirmed here that the genetic correlations across the hemispheres of the brain for measures of the same sulcus were significant (*ρ_G_* ~ 0.92 ± 0.10) (**Supp Tables S22-S25**).

**Figure 3 (left)** shows the genetic correlation (ρ_G_) between left and right homologous regions for those sulci that were significantly heritable in both left and right hemispheres (see **Supplementary Table S22-S25**) after Bonferroni correction (p<0.05/[115×4]). The genetic correlation across left and right is generally higher when examining the width of the sulci. The width of the central sulcus, the inferior frontal sulcus, intermediate frontal sulcus, superior frontal sulcus, posterior lateral sulcus, the superior postcentral intraparietal superior sulcus and the intraparietal sulcus, and the surface of the occipital lobe show significant genetic correlations across all cohorts (see **Supplementary Table S22-S24**)

**Figure 3:**
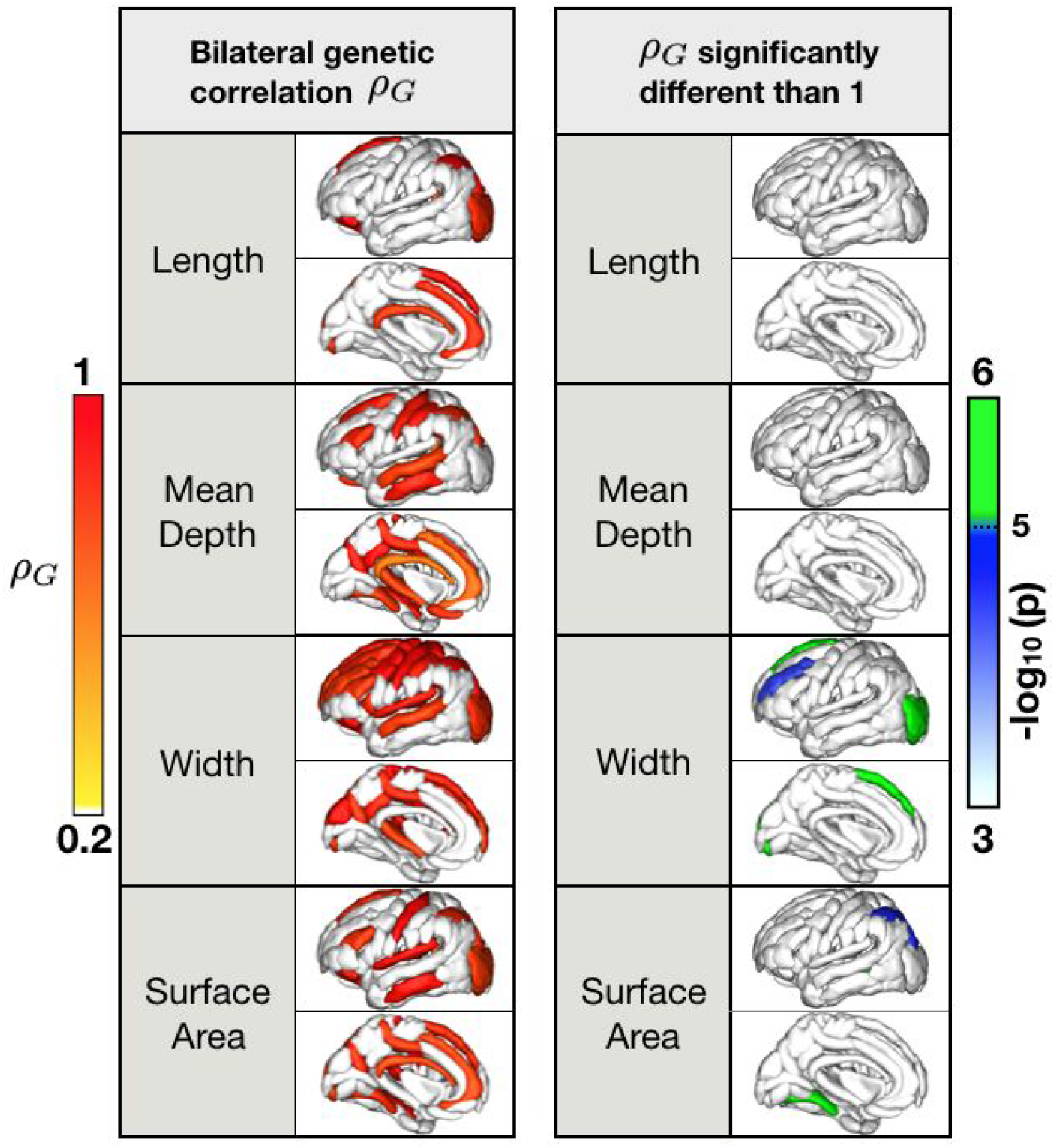
Left: The results of the meta-analysis across three family based cohorts testing the genetic correlation (*ρ_G_*) between measures from corresponding left and right hemispheres is shown. *ρ_G_* has been computed for those bilateral regions showing significant univariate heritability estimated with the meta-analysis as reported in Figure 2-a. Two sets of p-values are obtained when performing genetic correlations: a more traditional p-value comparing the correlation to the null, and another p-value comparing the genetic correlation to a perfect overlap, assessing the difference between the genetic correlation obtained and a correlation of 1. Only sulci showing a Bonferroni corrected significance comparing the correlation to the null for the tested regions and an overall ICC > 0.75 are highlighted. Bonferroni correction was conducted using the number of regions meeting the required ICC, so p < 0.05/[N], where N= 11 sulci for length + 31 for mean depth + 36 for width + 37 for surface area, for a total of 115 traits. Regional heritability estimates are tabulated in **Supplementary Table S10-S18**. Right: The −log_10_(p-value) relative to the difference between the correlation (*ρ*_G_) and 1 is mapped (**Supplementary Table S25**) for only those regions showing significant genetic correlation as on the left side (*ρ*_G_)

In contrast, phenotypic correlations (*ρ_P_*) between the left and right indices were on average less than *ρ_P_* < 0.5 in each cohort (**Supplementary Figure S18**). Sulcal width showed the highest (*ρ_P_* = 0.38 ± 0.15) meta-analyzed correlation between left and right homologs compared to the other sulcal descriptors (0.29 ± 0.07 for sulcal length, 0.30 ± 0.11 for mean depth and 0.33 ± 0.12 for surface area).

In **Figure 3 (right)**, we map regions with significant genetic correlations between left and right sulcal measures, but only those for which the genetic correlation is statistically different from 1; i.e., the 95% confidence interval surrounding the correlation estimate did not contain 1. These sulci represent brain regions that may have diverging genetic influences. In particular, the meta-analysis reveals that the occipital lobe, the intra-parietal lobe, the median frontal sulcus, the intermediate left frontal sulcus and the collateral sulcus may have lateralized genetic effects; sulcal length and mean depth did not show diverging genetic influences between the left and right hemispheres.

## Discussion

Our study has four main findings: 1) many of the sulci common across individuals can be reliably extracted, and some descriptors are more reliable than others; 2) cortical folding patterns are promising phenotypes for genetic analysis of cortical gyrification, yet some descriptors of sulci such as width may be more genetically influenced than for example, length; 3) these genetic influences vary regionally, with the earlier forming sulci have higher heritability estimates; 4) sulcal analyses may provide insight into the lateralization of genetic effects influencing brain structure.

Neuroimaging consortia such as ENIGMA^1^ and CHARGE (http://www.chargeconsortium.com/) require the evaluation of traits that can be reliably extracted regardless of the type of MRI scanner and scanning protocol used. Indeed, measurement error could lead to ceiling effect on heritability estimates, and genetic correlations. Highly heritable traits can be detected only if the traits are robustly measured^50^ and low reliability could lead to an underestimation of the true heritability^51^. While heritability is a population-specific estimate, one main goal of the imaging genetics field is to identify genetic variants that affect brain structure and function in populations around the world. Therefore, we aimed to ensure our measures were reliable across different datasets, as well as heritable across different populations.

Here we identified the most reliable sulcal regions in four independent cohorts with test-retest data. The ICC estimates the relation between within-subject variance and between-subjects variance, while the “bias” measure represents a subject-based index of consistency^48^. Our results show high consistency between test and retest (“bias” < 0.1^48^ on average) suggesting that the ICC could be affected by the homogeneity of the population under study: lower ICC values are expected when variability in global brain size or age range is limited. Combining several independent cohorts to assess the reliability of morphometric measures is beneficial. The meta-analysis of ICC measures estimated from these cohorts revealed that sulcal width was the most reliable metric among the descriptors analyzed.

Since cortical thickness has been reported as highly reliable^52^, the negative correlation between gray matter thickness and sulcal width shown in our study (**Figure S22**) supports the hypothesis that inter-subject variability, and by consequence sulcal labeling performance, affects the reliability of sulcal shape descriptors in addition to segmentation performance.

Prior studies described the association between reproducibility and heritability of shape measures for different brain structures in the QTIM cohort^53^. They found a correlation between reproducibility (ICC) and heritability, with a large percentage of traits showing “poor” reliability (ICC < 0.75)^53^. Here we show that most of the reliable sulcal shape descriptors are also under strong control. In four independent cohorts, we replicated prior findings^39,54,55^ and analyzing more than a hundred sulci across the brain, we demonstrated the metrics of the central sulcus shape as the most highly heritable traits^39,54,55^. Heritability of sulcal width was also higher for the right hemisphere than the left hemisphere, consistent with the hypothesis that there is less genetic control over the left hemisphere^56^.

Our results show variability of sulcal-based heritability patterns across sulcal descriptors in regions with “excellent” reliability (ICC > 0.9), confirming that not all the reliable traits are necessarily heritable^51^. These sulcal descriptors may serve as biomarkers for genetically mediated brain disorders and serve as phenotypes for large-scale GWAS, enhancing the discovery of specific genomic variants that influence brain structure and disease risk. In particular, our findings suggest that sulci appearing early in brain development^57,58^ that show high heritability such as the central sulcus, the Sylvian fissures, the parieto-occipital lobes and the superior temporal sulcus, may represent the optimal targets for a GWAS analysis. Indeed, we found a significant Pearson’s correlation between heritability averaged across sulcal descriptors and the appearance of sulci (in weeks)^59^ (**Figure S19**, r=−0.62, p=0.0025). We also found the frontal lobe to be significantly heritable, together with the temporal sulcus, even though this region develops later (secondary sulci)^58^.

We partially confirmed the results obtained in^60^. In particular, we found several medial frontal regions strongly heritable for sulcal surface area and width; our results confirm previous studies on the central sulcus^39^, the temporal lobe^56^ and the corpus callosum area^61^ and are also in line with studies showing high estimated heritability in prefrontal and temporal lobes for cortical thickness and surface area^62–68^, especially for sulcal mean depth and sulcal width.

Across three independent family based cohorts, QTIM - an Australian cohort of young adult twins and siblings - HCP, a North American cohort of twins and siblings, and GOBS - a Mexican American cohort of extended pedigrees, we found similar patterns of heritability for four descriptors of sulcal morphometry - length, depth, width, and surface area. Globally, we found sulcal heritability estimates of approximately 0.3−0.4, similar to estimates in other species, including *Papio* baboons^54^. Heritability estimates from GOBS were slightly lower than for QTIM or HCP, as expected for an extended pedigree design when compared to twin designs^69^. It has also been proposed that higher image quality, and therefore lower measurement error, could lead to higher heritability estimates^70^. GOBS and HCP volumes were acquired with a 3T scanner and HCP has higher spatial resolution compared to GOBS. QTIM was acquired with slightly lower spatial resolution but at higher magnetic field strength (4T). Further analyses will be needed to investigate how the SNR varies across QTIM, HCP and GOBS and how this affects heritability estimation^70^. SNP-based heritability estimated in the UK Biobank shows a similar pattern (**Figure S20**) across the brain, but with lower *h^2^* values compared to the family based cohorts. This might be due to the “missing heritability” effect arising in the SNP-based heritability estimation^71^.

Sulcal length is the descriptor with the least number of sulci showing significant heritability, and is likely more consistent throughout adult life. We found limited evidence for independent genetic influences on sulcal length once intracranial volume (ICV) was accounted for as a covariate.

Apart from work by the ENIGMA Laterality group^72^, most published ENIGMA studies^2,28,41,73^ performed analyses on pooled bilateral measures, averaging data from the left and right hemispheres; in fact, averaging the measures across hemispheres may provide more stable estimates, and higher signal-to-noise in measurements, yielding better power for genomic studies. To test whether this is acceptable when evaluating sulcal parameters, we performed a bivariate genetic analysis to estimate the genetic correlation between left and right sulcal measures. The sulcal measures showed not only significant heritability estimates when averaging the left and right hemispheres, but a strong genetic correlation was also observed between the contralateral measures for a subset of sulci.

A genetic correlation between right and left hemispheres, indicates pleiotropy, suggesting that genetic influences underlying the structure and variability in the measures tend to overlap. In family-based studies using bivariate variance components analysis to determine the genetic correlation components of variance, when a significant genetic correlation is identified, the confidence interval around the genetic correlation often includes one, suggesting the underlying genetic influences of the measures were not statistically distinguished from each other. Incomplete pleiotropy is suggested when genetic correlations are significant, but the confidence intervals do not include one. While in SNP-based genetic correlation models, incomplete pleiotropy may be suggested over complete pleiotropy in the presence of measurement error, in a bivariate polygenic model, measurement error falls into the environmental component of variance and the environmental correlation, and therefore does not influence the maximum-likelihood estimate of the genetic correlation; i.e, measurement error makes it more difficult to reject the null hypothesis that the genetic correlation is one. Features that exhibit unique genetic influences in one hemisphere may reveal insights into the biological causes of brain lateralization that may play an important role in neurodevelopmental or psychiatric disorders. Evidence of less genetic control in the left hemisphere has been found in^56^ and confirmed in^59^ where the authors found higher cortical gyrification complexity in the right hemisphere at an early development stage.

Here we found genetic asymmetries in the frontal lobe (width) that have been reported to be reduced in volumes in the right hemisphere when correlated with the duration of illness in schizophrenia^74^. This may relate to disorder-specific abnormalities seen in brain folding patterns rather than volume, as reported in a post-mortem study on schizophrenia^75^. Significant genetic differences were also detected in sulci of the occipital lobe, a region which has been associated with Parkinson’s disease^76,77^, posterior cortical atrophy (PCA), a disorder causing visual dysfunction, and logopenic aphasia^78^.

Some regions showing lateralization of genetic effects for sulcal descriptors, such as the collateral fissure (sulcal surface area), show the same effect for other measures extracted from the cortex such as the cortical surface area (**Figure S21**). However, for other regions, such as the occipital lobe, we find lateralization of genetic factors for sulcal measures (*width*) and not for regional cortical thickness or surface area measures. This suggests that sulcal descriptors could offer additional insights into brain lateralization, beyond more commonly collected metrics.

The genetic influences on brain structure are regionally dependent, and differ according to the measurement under study. For example, the genetic correlations between cortical thickness and surface area are weak and negative^14^. In non-human primates, brain cortical folding was also found to be influenced by genetic factors largely independent of those underlying brain size^16,79^. Measuring cortical folding through sulcal-based morphometry could therefore highlight brain metrics beyond thickness and surface area, and may complement these more traditional measures to reveal a deeper understanding of the genetic architecture of human brain structure. This may be particularly relevant for the sulcal width -the most heritable of the four tested metrics.

Given our results, conducting a GWAS of each hemisphere’s features for the most heritable features from the frontal, temporal and occipital regions, may provide insight into the biological mechanisms that drive the structure and function of each hemisphere, as well as hemispheric specialization. However, for most sulcal regions, the genetic architecture of the two hemispheres is largely correlated. The discovery and replication of specific genetic influences on brain structure requires very highly powered analyses, achievable through large-scale studies and collaboration. Harmonized imaging and genetic protocols, rigorous quality assurance, reproducibility assessments, along with statistical rigor are vital in these collaborative endeavors such as those by ENIGMA. We have made these customized protocols using BrainVISA software available at: http://enigma.ini.usc.edu/protocols/imaging-protocols/.

## Materials and Methods

### Participants and MRI Imaging

#### QTIM (Queensland Twin Imaging study)

1,008 right-handed participants^80^, 370 females and 638 males were studied, including 376 dizygotic (DZ) and 528 monozygotic (MZ) twins (one set of DZ triplets) and 104 siblings, with an average age of 22.7 ± 2.7 years [range: 18–30]. T1-weighted images were acquired on a 4 T Bruker Medspec scanner with an inversion recovery rapid gradient echo sequence. Acquisition parameters were: inversion/repetition/echo time (TI/TR/TE) = 700/1500/3.35 ms; flip angle = 8 degrees; with an acquisition matrix of 256 × 256; voxel size= 0.94 × 0.90 × 0.94 mm^3^.

#### HCP (Human Connectome Project)

816 participants^81^, 362 females and 454 males, average age, 29.1 ± 3.5 years [range: 22–36]. These included 412 siblings, 205 dizygotic (DZ) and 199 monozygotic (MZ) twins, including triplets. T1-weighted images were acquired using a 3T Siemens scanner. MRI parameters: (TI/TR/TE) = 1000/2400/2.14 ms; flip angle = 8 degrees; voxel size = 0.7 mm isotropic, acquisition matrix = 224 × 224. The subset of test-retest scans includes all right-handed subjects.

#### GOBS (Genetics of Brain Structure and Function project)

A total of 1,205 Mexican-American individuals from extended pedigrees (71 families, average size 14.9 [1–87] people) were included in the analysis. 64% of the participants were female and ranged in age from 18 to 97 (mean ± SD: 47.1 ± 14.2) years. Individuals in this cohort have actively participated in research for over 18 years and were randomly selected from the community with the constraints that they are of Mexican-American ancestry, part of a large family, and live within the San Antonio region. Imaging data were acquired at the UTHSCSA Research Imaging Center on a Siemens 3 T Trio scanner (Siemens, Erlangen, Germany). Isotropic (800 μm) 3D Turbo-flash T1-weighted images were acquired with the following parameters: TE/TR/TI = 3.04/2100/785 ms, flip angle = 13 degrees. Seven images were acquired consecutively using this protocol for each subject and the images were then co-registered and averaged to increase the signal-to-noise ratio and reduce motion artifacts^82^.

#### UK Biobank

Analyses were conducted on the 2017 imputed genotypes restricted to variants present in the Haplotype Reference Consortium^83,84^. UK Biobank bulk imaging data were made available under application #11559 in July 2017. We analyzed 10,083 participant (4807 females), mean age= 62.4 ± 7.3 years [range: 45-79]. Voxel matrix: 1.0×1.0×1.0 mm - acquisition matrix: 208×256×256. 3D MP-RAGE, TI/TR=880/2000 ms, sagittal orientation, in-plane acceleration factor=2. Raw MRI data were processed using the ENIGMA FreeSurfer and sulcal analysis protocols. Following processing, all images were visually inspected for FreeSurfer quality control of grey/white matter classifications. The central sulcus segmented and labeled by BrainVISA was also visually controlled for labeling quality for all subjects.

#### KKI (Kennedy Krieger Institute - Multi-Modal MRI Reproducibility Resource)

Twenty-one healthy volunteers with no history of neurological conditions (10 F, 22-61 years old) were recruited. All data were acquired using a 3 T MRI scanner (Achieva, Philips Healthcare, Best, The Netherlands) with body coil excitation and an eight-channel phased array SENSitivity Encoding (SENSE) head-coil for reception. All scans were completed during a 2-week interval. The resulting dataset consisted of 42 “1-h” sessions of 21 individuals. MP-RAGE T1-weighted scans were acquired with a 3D inversion recovery sequence: (TR/TE/TI = 6.7/3.1/842 ms) with a 1.0 × 1.0 × 1.2 mm^3^ resolution over a field of view of 240 × 204 × 256 mm acquired in the sagittal plane. The SENSE acceleration factor was 2 in the right-left direction. Multi-shot fast gradient echo (TFE factor = 240) was used with a 3-s shot interval and the turbo direction being in the slice direction (right–left). The flip angle was 8 degrees. No fat saturation was employed^85^, https://www.nitrc.org/projects/multimodal/.

#### OASIS

This test-retest reliability data set contains 20 right-handed subjects (19–34 years old) without dementia imaged on a subsequent visit within 90 days of their initial session. MPRAGE T1-weighted scans were acquired on a 1.5-T Vision scanner (Siemens, Erlangen, Germany): (TR/TE/TI = 9.7/4.0/20 ms) with an in-plane resolution of 1.0 × 1.0 × mm^2^ resolution over a FOV of 256 × 256 mm acquired in the sagittal plane. Thickness/gap= 1.25/0 mm; flip angle = 10 degrees. (https://www.oasis-brains.org/)^86^.

**Table S1:** Genetic Analysis: demographics for the 4 cohorts analyzed in this study

| Cohort | N(%F) | Ethnicity | Age in years<br>(mean +/- stdev [range]) | Relatedness |
| --- | --- | --- | --- | --- |
| QTIM | 1,008<br>(37%) | Caucasian | $22.7 \pm 2.7$ [18–30] | 376 DZ<br>528 MZ<br>104 Siblings |
| HCP | 816<br>(44%) | Mixed US population | $29.1 \pm 3.5$ [ 22–36] | 205 DZ<br>199 MZ and triplets<br>412 Siblings |
| GOBS | 1205<br>(64%) | Mexican American | $47.1 \pm 14.2$ [18–97] | 71 families |
| UKBB | 10,083<br>(47%) | Caucasian | $62.4 \pm 7.3$ [45–79] | Unrelated |

**Table S2:** Cohorts analyzed for the test-retest study. HCP and QTIM were used for the reproducibility analysis as they were representative of subjects examined in the genetic analysis. Among publicly available datasets we selected KKI and OASIS, as in^48^, based on age (18< age <65) and inter-scan interval (<90 days)

| Cohorts | Age Range (mean) | N°subject (%F) | Inter-scan Interval (days) | Field Strength [T] | Voxel size [mm]<sup>3</sup> |
| --- | --- | --- | --- | --- | --- |
| KKI | 22 – 61 (31.8) | 21 (48%) | 14 | 3 | [1 x 1 x 1.2] |
| HCP | 24 - 35 (30.1) | 35 (44%) | 90 | 3 | [0.7 x 0.7 x 0.7] |
| OASIS | 19 - 34 (23.3) | 20 (60%) | 90 | 1.5 | [1.0 x 1.0 x 1.25] |
| QTIM | 21 – 28 (23.2) | 34 (37%) | 90 | 4 | [0.94 x 0.98 x 0.98] |

### MRI image processing and sulcal extraction

Anatomical images (T1-weighted) were corrected for intensity inhomogeneities and segmented into gray and white matter tissues using FreeSurfer (http://surfer.nmr.mgh.harvard.edu/); segmentations and regional labels were quality controlled using ENIGMA protocols for outlier detection and visual inspection (http://enigma.ini.usc.edu/protocols/imaging-protocols/). BrainVISA (http://brainvisa.info) was run for sulcal extraction, identification, and sulcal-based morphometry. Morphologist 2015, an image processing pipeline included in BrainVISA, was used to quantify sulcal parameters. Briefly, the Morphologist 2015 segmentation pipeline computes left and right hemisphere masks, performs gray and white matter classification, reconstructs a gray/white surface and a spherical triangulation of the external cortical surface, independently for both hemispheres. However, to improve sulcal extraction and build on current protocols used by hundreds of collaborators within ENIGMA, quality controlled FreeSurfer outputs (*orig.mgz, ribbon.mgz, and talairach.auto*) were directly imported into the pipeline to avoid re-computing several steps, including intensity inhomogeneity correction and gray/white matter classification. Sulci were then automatically labeled according to a pre-defined anatomical nomenclature of 62 sulcal labels for the left hemisphere and 61 sulcal labels for the right hemisphere^87,88^.

### Sulci descriptors and quality control

Analyzing the shape of the cortex through sulcal-based morphometry allows us to quantify the geometry of a sulcus in terms of several distinct and complementary descriptors, consisting of length, mean depth, surface area and width (or fold opening) of all extracted and labeled sulci. Cortical thickness and surface area have been found highly heritable and largely genetically correlated^13,89^. Moreover, specific genetic loci associated with cortical surface area and thickness are involved in cortical development^90^. Cortical thickness, surface area, and folding tend to exhibit different age-related trajectories^91,92^. In particular, cortical thickness is determined by the horizontal layers in the cortical mantle, surface area reflects the number of radial columns perpendicular to the pial surface^92^ and sulcal shape relates to the microstructure of the neuronal sheets and to the local axonal connectivity within a cortical region, which may influence the degree of folding^16^.

The *length* of a sulcus is measured in millimeters as the geodesic length of the junction between a sulcus and the hull of the brain. The *mean depth* corresponds to the average of the depth across all the vertices along the bottom of a sulcus (the depth of a vertex located at the bottom of a sulcus is defined as the geodesic distance along the sulcus to the brain hull). The *surface area* is the total area of the sulcal surface. The enclosed CSF volume divided by the sulcal surface area gives the *width*, a gross approximation of the average width of the CSF in the fold^55^.

To further quality control the extracted sulcal measures and identify subjects whose sulci were not optimally identified, we consider as outliers those subjects showing, for a particular sulcus, abnormal values for at least one of the descriptors. That is, for a given sulcus, the z-score across subjects is computed for each descriptor. The set of subjects showing an absolute z-score greater than 2.5 for one or more descriptor were discarded from further analysis^93^. This led to discard ~ 3% of subjects for each sulcus. An outlier for one sulcus has been included in the analysis for the other sulci if the absolute z-score was less than 2.5.

### Univariate and bivariate quantitative genetic analyses

The relative influences of genetic and environmental factors on human traits can be estimated by modeling the known genetic relationship between individuals and relating it to observed covariance in measured traits; in twin studies, monozygotic (MZ) twin pairs - who typically share all their common genetic variants - are compared to dizygotic (DZ) twin pairs, who share, on average, 50%. The same principle can be used for extended pedigrees, in which a large number of individuals have varying degrees of relatedness. Here, we use both twins and extended pedigrees to estimate the heritability of these in-depth cortical sulcal measures. For a given cohort of participants, the narrow-sense heritability (*h^2^*) is defined as the proportion of the observed variance in a trait (*σ*^2^_*P*_) that can be attributed to additive genetic factors 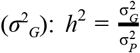.

As part of the ENIGMA consortium, we developed image-processing protocols based on BrainVISA software (http://brainvisa.info/web/index.html)^87,88,94^ to identify sulci and extract common descriptors (**Section 3.2**). Using SOLAR-ECLIPSE imaging genetics tools (http://www.nitrc.org/projects/se_linux)^95^, we investigated the heritability profile of sulci across the whole brain: 62 on the left and 61 on the right hemisphere.

Variance components methods, implemented in the Sequential Oligogenic Linkage Analysis Routines (SOLAR) software package^95^, were used for all genetic analyses. Heritability (*h^2^*) is the proportion of total phenotypic variance accounted for by additive genetic factors and is assessed by contrasting the observed phenotypic covariance matrix with the covariance matrix predicted by kinship. High heritability indicates that the covariance of a trait is greater among more closely related (genetically similar) individuals; here, for example, monozygotic twins as compared to dizygotic twins and siblings.

Prior to testing for the significance of heritability, sulcal descriptor values for each individual are adjusted for a series of covariates. We estimated the influence of specific variables (additive genetic variation, and covariates including intracranial volume, sex, age, age^2^, age × sex interaction, age^2^ × sex interaction) to calculate heritability and its significance (*p*-value) for accounting for a component of each trait’s variance within this population.

The significance threshold for heritability analysis of individual sulci was set to be p≤ (0.05/m), where *m* = 61 (number of bilateral sulci) times 4 (number of measurements). *m* = 123 when left and right sulcal heritability is estimated. This reduced the probability of Type 1 errors associated with multiple measurements.

Classical quantitative genetic models were used to partition the phenotypic correlation (ρ_P_) between the left and the corresponding right sulcal measures into the genetic (ρ_G_), and a unique environmental (ρ_E_) components, for each pair of traits. Just as with the univariate model, the bivariate phenotype of an individual is modeled as a linear function of kinship coefficients that express relatedness among all individuals within the cohort (MZ twins share all their additive genetic information and DZ twins and siblings share on average 50%). The significance of ρ_G_ and ρ_E_ were estimated from the likelihood ratio test when comparing the model to ones where the correlation components are constrained to be zero^95–97^. This estimates ρ_G_ and ρ_E_ and their standard errors. The significance of these coefficients is determined by a *z*-test of their difference from zero. If ρ_G_ differs significantly from zero then a significant proportion of the traits’ covariance is influenced by shared genetic factors.

In this case, we tested another model where the genetic correlation factor ρ_G_ is fixed to 1. Fixing ρ_G_ to 1 suggests that the additive genetic components comprising the two traits overlap completely, and there is no detectable unique genetic composition for the individual traits. Once again, the log-likelihood of this model is compared to one where the parameters are freely optimized. If ρ_G_ is not found to significantly differ from 1, then we cannot reject the hypothesis that both heritable traits are driven by the same set of genetic factors. If ρ_G_ is significantly different from 0 and significantly different from 1, then the traits share a significant portion of their variance, however each is also likely to be partially driven by a unique set of genetic factors.

Some considerations should be made regarding the measurement error of the traits analyzed here: ρ_G_ is the correlation between the latent genetic effects on the two traits irrespective of the proportion of phenotypic variance these latent effects explain (i.e., heritability). Measurement error, which is uncorrelated between individuals regardless of their relatedness, falls into the environmental component and environmental correlations. Measurement error therefore influences *h^2^*, ρ_E_, ρ_P_, but definitely never ρ_G_.

In practice, measurement error does make ρ_G_ harder to estimate, because low heritability means that the underlying genetic effects cannot be estimated with much precision. This causes the standard error of the ρ_G_ estimate to increase, but critically, doesn’t change its maximum-likelihood estimate systematically. So, measurement error makes it harder to reject the null hypothesis that ρ_G_=1.

Moreover, the bivariate polygenic model used here to estimate the left-right genetic correlation is a linear function of laterality (L-R). Indeed, the genetic variance of L-R is

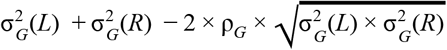

Where 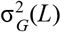 and 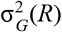 are the genetic variance for the left and right traits. The phenotypic variance is similarly defined so that the heritability of L-R can be obtained. But if L-R shows significant heritability, it could be because: 1) genetic overlap is incomplete and/or 2) L and R have unequal genetic variances. So, studying laterality is not recommended here because 1) and 2) are confounded.

### Meta-analysis of additive genetic variance

Meta-analysis calculate weighted mean heritability (h^2^) and standard error estimates based on measurements from individual cohorts^40,41^. We weight the heritability from each cohort by the heritability standard error, as extracted from the variance component model of SOLAR. The heritability weighted by standard error^40,41^ is:

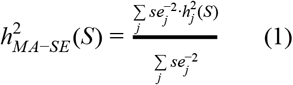

Where S=1 to *N_S_* indexes the sulci and *j* =1, 2,3 indexes the cohorts.

### Mega-analysis of additive genetic variance

While meta-analyses compute first the heritability independently for each cohort and then combine the results, mega-analyses combine first different cohorts and then run a single computation for heritability evaluation. We use a program (*polyclass*), developed for SOLAR^98^ for mega-analysis of heritability on sulci descriptors^41,99^. This function fits the model after combining the pedigrees of QTIM, HCP and GOBS into a single pedigree (for more details see^40,41^).

### Meta-analysis of genetic correlation

A meta-analysis of genetic correlation is calculated weighting the genetic correlation computed for each cohort by its sample size:

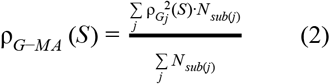

Where S=1 to *N_S_* indexes the sulci, *j* = 1..3 indexes the cohorts and *N_sub(j)_* is the sample size of cohort *j*. To combine p-values in a meta-analysis we used the Edgington’s method which represents a compromise between methods more sensitive to largest p-values (e.g. Pearson’s method) and methods more sensitive to smallest p-values (e.g. Fisher’s method)^100,101^:

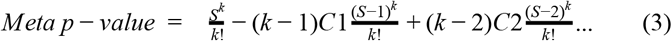

where *S* is the sum of o-values and *k* the number of tests (i.e. *k*=3 cohorts in our study). The corrective additional terms are used as long as the number subtracted from *S* in the numerator is less than *S*. All the p-values in the meta-analyses estimated were computed using this method.

### SNP-based heritability analysis

We used Genome-wide Complex Trait Analysis (GCTA)^102^ to calculate the heritability from the individual genotypes. Genotypes on the autosomal chromosomes were used to calculate the genetic relationship matrix (GRM) with GCTA^102^. Heritability, was calculated using a linear mixed model, with age, sex, ICV, and the first four genetic components from multidemential scaling analysis as fixed covariates; we also covaried for the presence of any diagnosed neurological or psychiatric disorder. In our analysis, we excluded participants with non-European ancestry, missing genotypes or phenotypes, and mismatched sex information.

### Reliability analysis

#### Sulcal measurement reliability

To evaluate the reliability of the sulcal shape descriptors we analyzed their variability, or reproducibility error, across the test–retest (TRT) sessions for each of the 4 TRT cohorts. For each MRI scan there are several sources of variability, including variability from hydration status, variability due to slightly different acquisitions in the two sessions (head position change in the scanner, motion artifacts, scanner instability, etc.), and finally variability due to the imaging processing methods themselves.

There could also be variability in the reliability estimates depending on the type of MRI system used (vendor, model, acquisition parameters), so it is important to address the issue of reliability across a variety of platforms. We used 2 indices of reliability: 1) the dimensionless measure of absolute percent bias of descriptor, *b*, (sulcal length, mean depth, width and surface area) of a sulcus with respect to its average; 2) the intraclass correlation coefficient (ICC). *b* is computed as follows:

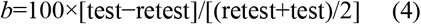

The estimation of the means is more robust than the estimation of the variance from the signed differences, in particular for smaller sets of subjects. The distributions of sulcal measurement differences plotted the mean across sessions were examined with a Bland–Altman analysis^103^. These plots show the spread of data, the bias (i.e. mean difference) and the limits of agreement (± 1.96 SD), and were used to confirm that the distributions were approximately symmetric around zero and to check for possible outliers. While the ICC estimates the relation between within-subject variance and between-subjects variance, *b* offers a subject-based index that might be used to find outliers. If scan and rescan are perfectly reliable, *b* should be equal to zero. The cases where *b* is greater than 0.1, as in^48^ are considered unreliable.

The intraclass correlation coefficient (ICC) was computed to quantify the reproducibility for sulcal-based measurements. ICC is defined as

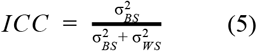

providing an adequate relation of within-subject (σ^2^_WS_) and between-subject (σ^2^_BS_) variability^46,104,105^.

The ICC estimates the proportion of total variance that is accounted for by the σ^2^_BS_. Values below 0.4 are typically classified as ‘poor’ reproducibility, between 0.4 and 0.75 as ‘fair to good’, and higher values as ‘excellent’ reproducibility^47^.

Equation 5 was used to estimate the ICC for each sulcal descriptor, independently for each cohort. The four cohorts were then combined into a meta-analysis (*ICC_MA-SE_*), similar to equation 1, in order to account for intra-site variability end to better estimate the sulcal reliability:

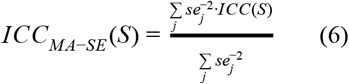

where *j* = 1, 2,3,4 indexes the cohorts. The standard error (SE) was computed like SE = ICC/ Z, where Z is obtained from a normal distribution knowing the p-value. *ICC_MA-SE_* was computed only if the cohort-based ICC computed with equation 5 was estimated for at least 3/4 cohorts.

## Data availability

**OASIS:** The OASIS data are distributed the greater scientific community under the Creative Commons Attribution 4.0 license. All data is available via www.oasis-brains.org^86^.

**KKI (Kennedy Krieger Institute - Multimodal MRI Reproducibility Resource):** Open access: https://www.nitrc.org/projects/multimodal/^85^.

**QTIM:** Data from the QTIM cohort used in this manuscript can be applied for by contacting Dr. Margaret Wright. Access to data by qualified investigators are subject to scientific and ethical review. Summary results from cohort QTIM are available as part of the supplementary data^49^.

**HCP:** Family status and other potentially sensitive information are part of the Restricted Data that is available only to qualified investigators after signing the Restricted Data Use Terms. Open access data (all imaging data and most of the behavioral data) is available to those who register and agree to the Open Access Data Use Terms.

Restricted data elements that could be potentially used to identify subjects include family structure (twin or non-twin status and number of siblings); birth order; age by year; handedness; ethnicity and race; body height, weight, and BMI; and a number of other categories. Each qualified investigator wanting to use restricted data must apply for access and agree to the Restricted Data Use Terms (https://humanconnectome.org/study/hcp-young-adult/data-use-terms)^81^.

**GOBS:** Data from the GOBS cohort used in this manuscript can be applied for by contacting Prof. David Glahn or Prof. John Blangero. Access to data by qualified investigators are subject to scientific and ethical review and must comply with the European Union General Data Protection Regulations (GDPR)/all relevant guidelines. The completion of a material transfer agreement (MTA) signed by an institutional official will be required. Summary results from cohort GOBS are available as part of the supplementary data.

**UKBB:** Access to data from the UK Biobank can be obtained by approved scientists through application with UK Biobank (www.ukbiobank.ac.uk/researchers)^84^.

## ACKNOWLEDGMENTS

This research was funded in part by the NIH Big Data to Knowledge (BD2K) program for supporting the ENIGMA Center of Excellence with grant U54EB020403. Additional support was provided by NIH grants R01AG059874, R01MH117601, R01MH121246, and P41EB015922.A research grant from Biogen Inc (Jahanshad/Thompson) helped support image data processing for the UK Biobank Resource, which was made available under application number 11559. Data collection for QTIM was supported by NIH grant R01HD050735, and Australian NHMRC grant 486682 (M Wright); financial support for GOBS was provided by the grants MH078143, MH078111, and MH083824 (PIs: DC Glahn & J Blangero); HCP data were provided [in part] by the Human Connectome Project, WU-Minn Consortium (PIs: D Van Essen & K Ugurbil; U54MH091657) funded by the 16 NIH Institutes and Centers that support the NIH Blueprint for Neuroscience Research; and by the McDonnell Center for Systems Neuroscience at Washington University; OASIS: Cross-Sectional was supported by P50AG05681, P01AG03991, P01AG026276, R01AG021910, P20MH071616, U24RR021382 (PIs: D Marcus, R Buckner, J Csernansky, J Morris); KKI was supported by NIH grants P41RR015241(PCM Zijl), R01NS056307 (J Prince), R21NS064534 (BA Landman/J Prince), R03EB012461 (BA Landman). BrainVISA’s Morphologist software development has received funding from the European Union’s Horizon 2020 Framework Programme for Research and Innovation under Grant Agreement No 720270 & 785907 (Human Brain Project SGA1 & SGA2), and by the FRM DIC20161236445.

We also thank Dr. Anderson M. Winkler for feedback on our original preprint, and Dr. Alessandra Griffa for assistance with visualization.

## Footnotes

1 http://www.chargeconsortium.com/

