## Supplementary Figures for "The reliability and heritability of cortical folds and their genetic correlations across hemispheres"

Pizzagalli F et al.

### Supplementary Material

**List of Figures:**

**ICC analysis**

Figure S1-S3: ICC plotted for each sulcus, highlighting the 4 test-retest cohorts analyzed in this work. The red line represents the meta-analysis of ICC.

Figure S4-S7: Sulcal regions ranked based on reliability (ICC)

Figure S8: Sulcal regions ranked based on the mean across sulcal descriptors of reliability (ICC)

**Heritability (h2) analysis**

Figure S9: Sulci showing univariate $h^2$ overlap between QTIM HCP and GOBS for the left sulci.

Figure S10: Sulci showing univariate $h^2$ overlap between QTIM HCP and GOBS for the right sulci.

Figure S11: Sulci showing univariate $h^2$ overlap between QTIM HCP and GOBS for the bilaterally averaged sulci.

Figure S12: Violin plots showing heritability estimates for QTIM, HCP, and GOBS cohorts and for mega and meta analyses.

Figure S13: Pearson's correlation coefficients estimated between Meta- and Mega-analysis.

Figure S14: Sulcal based heritability in GOBS comparing h2 estimation with and without ICV as covariate.

Figure S15: Pearson's correlation between Meta-$h^2$ and Meta-ICC analysis.

Figure S16: Pearson's correlation between UKBB-$h^2$ and Meta-ICC analysis.

Figure S17: Univariate heritability ($h^2$) with standard error of global sulcal shape descriptor, for left and right hemisphere.

Figure S18: Meta-analysis of Pearson's correlation between left and right sulcal length, mean depth, surface area, and width.

Figure S19: Pearson's correlation between heritability ($h^2$) and the appearance of sulci.

Figure S20: Pearson's correlation between heritability ($h^2$) estimated from UKBB sulcal shape descriptors and the meta analysis of heritability estimated from QTIM, HCP and GOBS.

Figure S21: ROI-based phenotypic and genetic correlation between left and right hemispheres computed for cortical thickness and surface.

Figure S22: Sulcal-based correlation between sulcal width and grey matter thickness computed with BrainVISA for each sulcal surrounding area.

**Figure S1: Left Hemisphere**: ICC across **left** sulci for the 4 test-retest cohorts analyzed here. The red line corresponds to the meta-analysis (see Supp. Table_S2). Missing values correspond to descriptors that were to be measured by BrianVISA for more than half of the subjects in a cohort, such as the length of the left *insula* and the left posterior sub-central ramus of the lateral fissure (*F.C.L.r.sc.post.)*. Measures for the anterior sub-central ramus of the lateral fissure (*F.C.L.r.sc.ant.*) and the diagonal ramus of the lateral fissure (*F.C.L.r.diag*) were missing for all the descriptors.

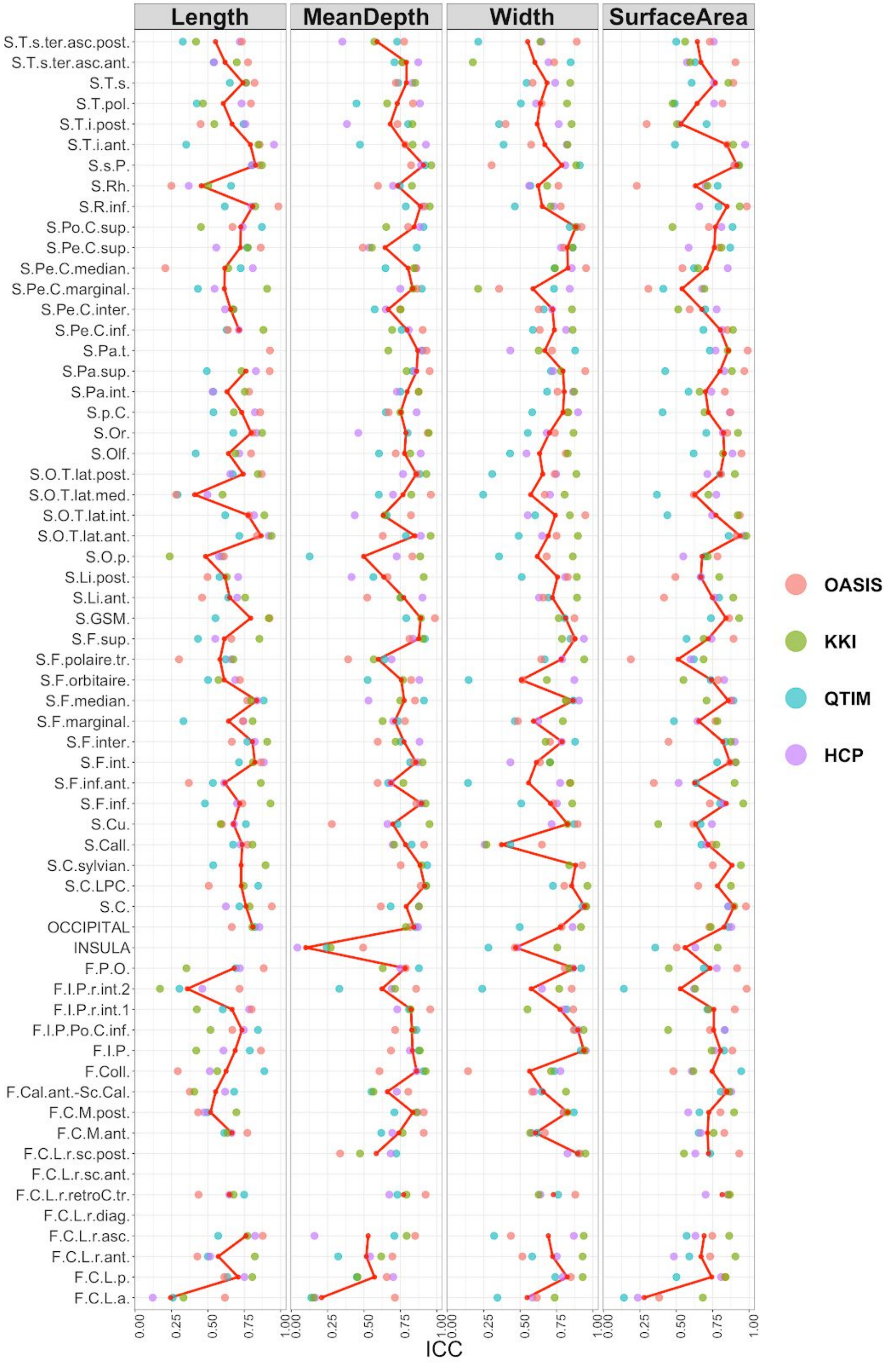

**Figure S2: Right Hemisphere**: ICC across right sulci for the 4 test-retest cohorts analyzed here. The red line corresponds to the meta-analysis (see Supp. Table_S3). Missing values correspond to descriptors that were to be measured by BrianVISA for more than half of the subjects in a cohort, such as the length of the left *insula* and the left posterior sub-central ramus of the lateral fissure (*F.C.L.r.sc.post.)*. Measures for the anterior sub-central ramus of the lateral fissure (*F.C.L.r.sc.ant.*), and the paracentral lobule central sulcus (SC.LPC.). For the diagonal ramus of lateral fissure (*F.C.L.r.diag*) measures were missing for 4/5 of the cohorts and thus discarded for meta-analysis and left/right average, for all the descriptors.

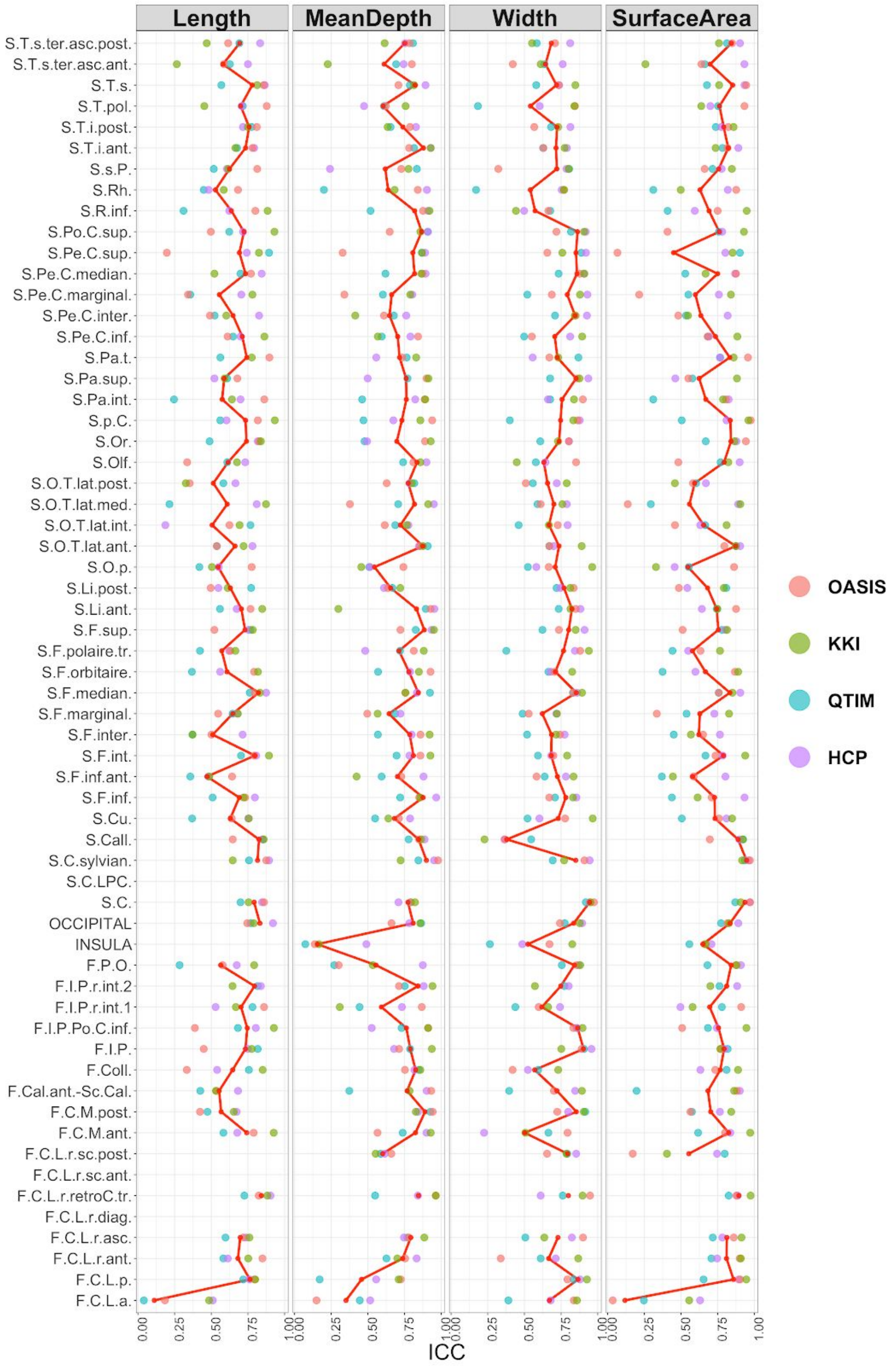

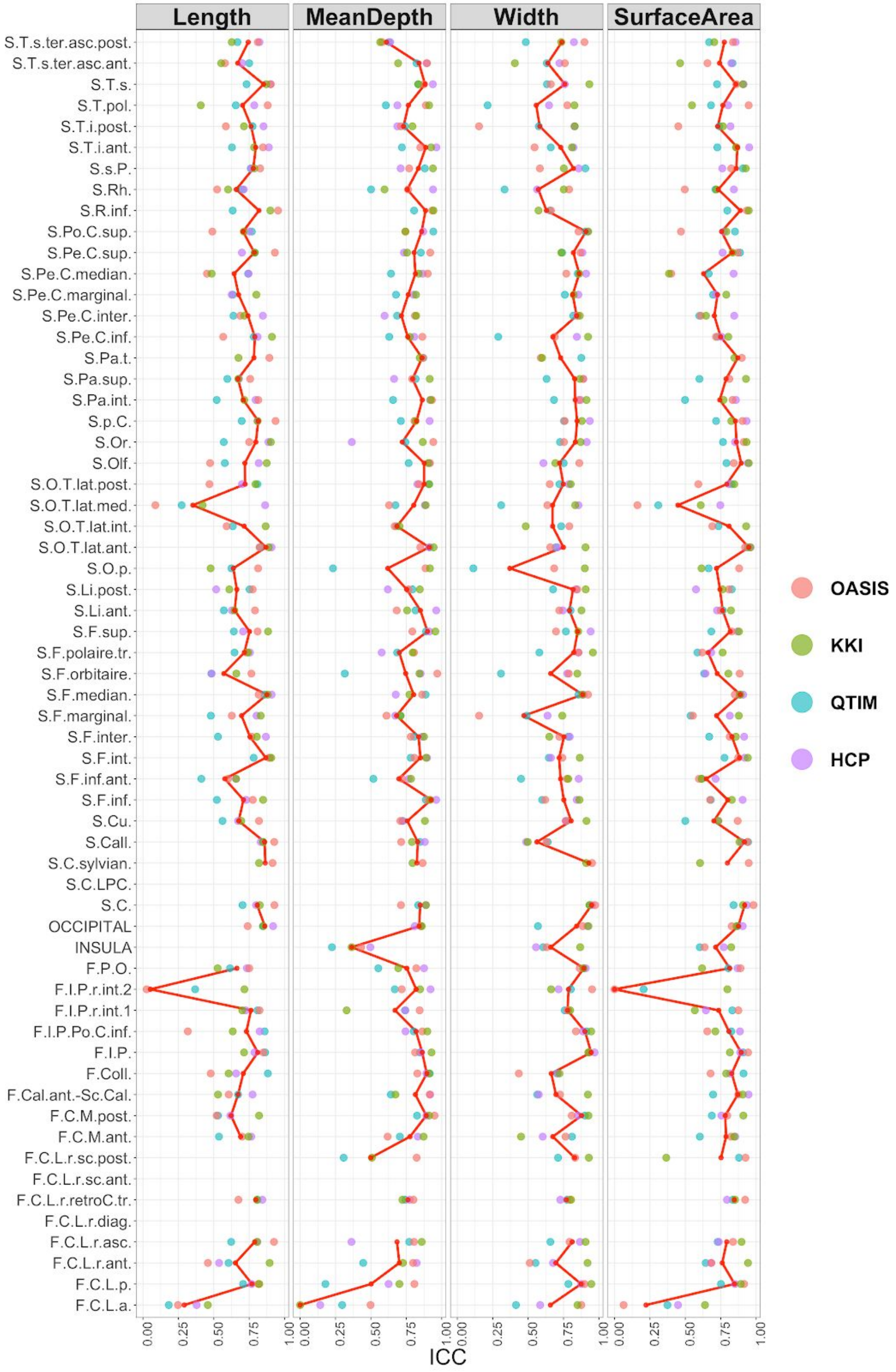


**Figure S3: Bilaterally averaged sulcal measure**: ICC across sulci for the 4 test-retest cohorts analyzed here. The red line corresponds to the meta-analysis (see Supp. Table_S4). Missing values correspond to descriptors that were to be measured by BrianVISA for more than half of the subjects in a cohort, such as the length of the right *insula,* the paracentral lobule central sulcus (SC.LPC.), the anterior sub-central ramus of the lateral fissure (*F.C.L.r.sc.ant.*) and the diagonal ramus of lateral fissure (*F.C.L.r.diag*) for all the descriptors.

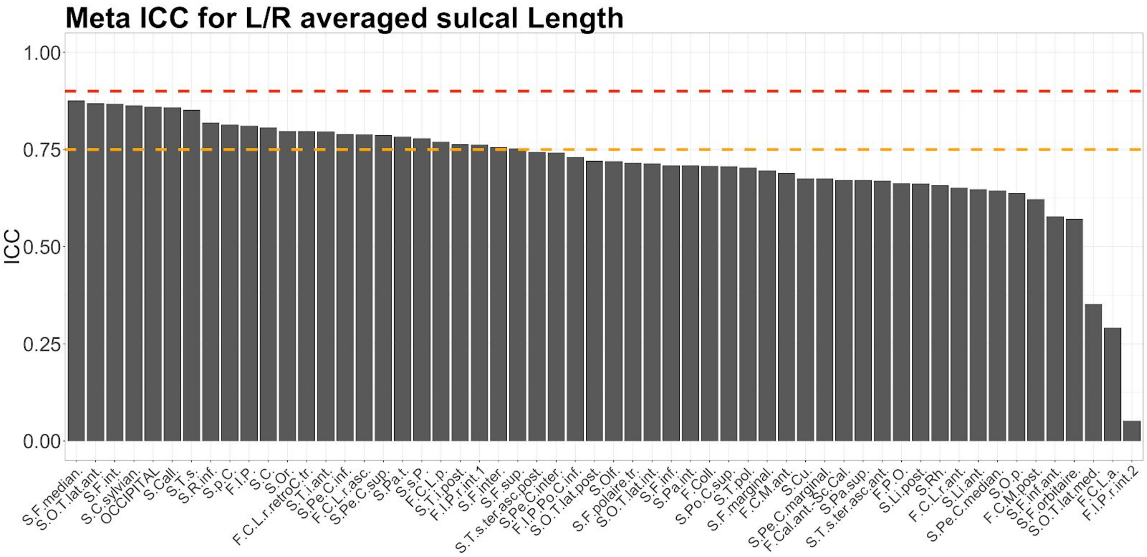


**Figure S4: Meta-Analysis:** Ranked ICC for Left/Right averaged sulcal **Length**. The dashed orange and red lines highlight the ‘good’ (ICC=0.75) and ‘excellent’ (ICC=0.9) reliability thresholds [1]. (see Figure 1 and Supp. Table_S4).

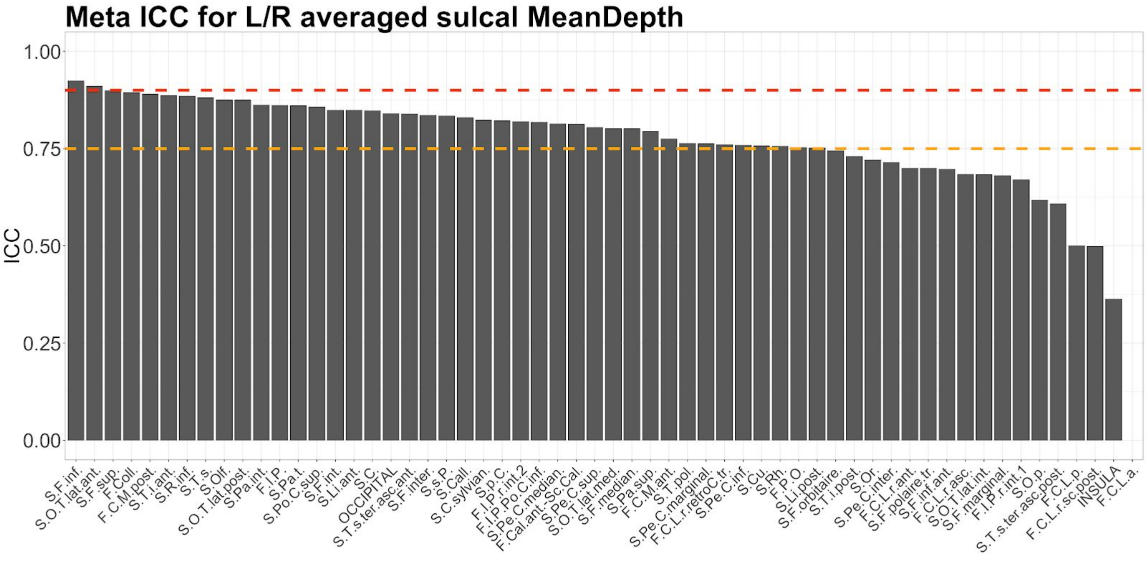


**Figure S5:** Meta-Analysis: Ranked ICC for Left/Right averaged sulcal **Mean Depth**. The dashed orange and red lines highlight the ‘good’ (ICC=0.75) and ‘excellent’ (ICC=0.9) reliability thresholds [1]. (see Supp. Table_S4).

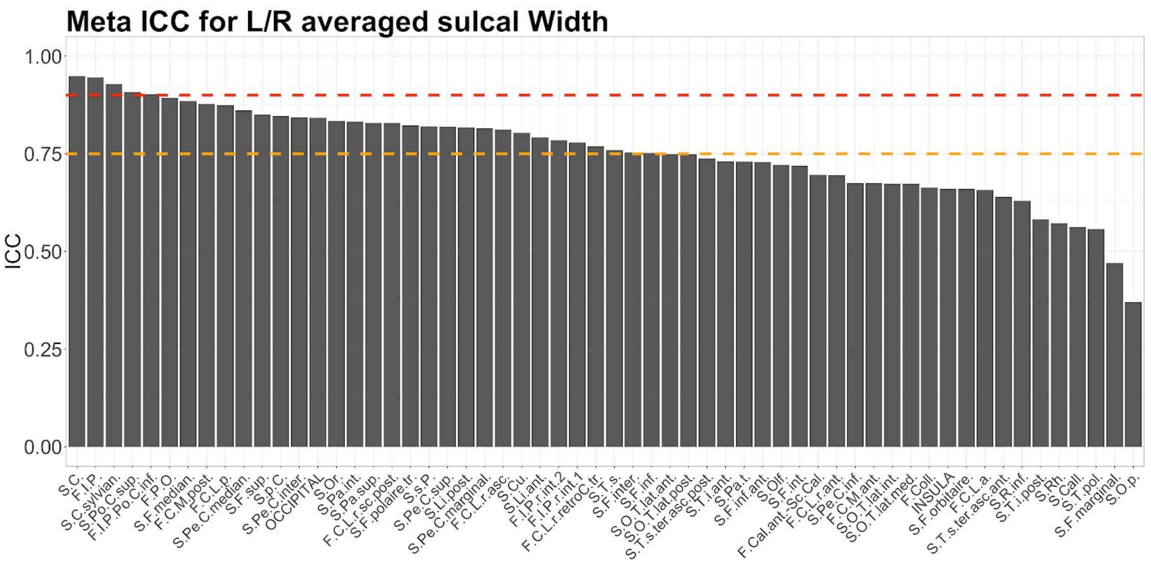


**Figure S6: Meta-Analysis:** Ranked ICC for Left/Right averaged sulcal **Width**. The dashed orange and red lines highlight the 'good' (ICC=0.75) and 'excellent' (ICC=0.9) reliability thresholds [1]. (see Supp. Table_S4).

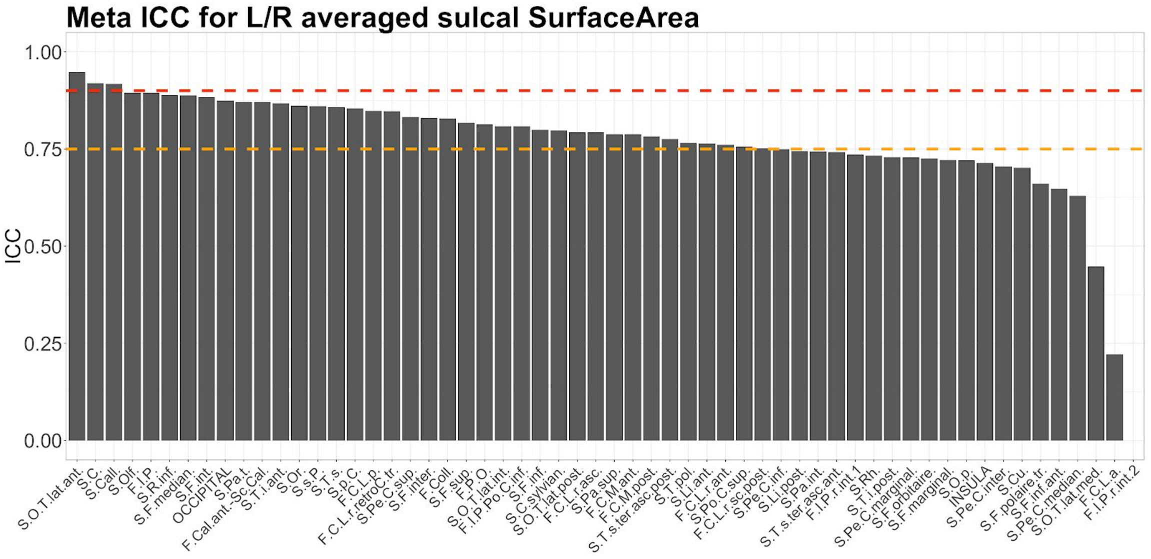


**Figure S7: Meta-Analysis:** Ranked ICC for Left/Right averaged sulcal **Surface Area**. The dashed orange and red lines highlight the 'good' (ICC=0.75) and 'excellent' (ICC=0.9) reliability thresholds [1]. (see Supp. Table_S4).

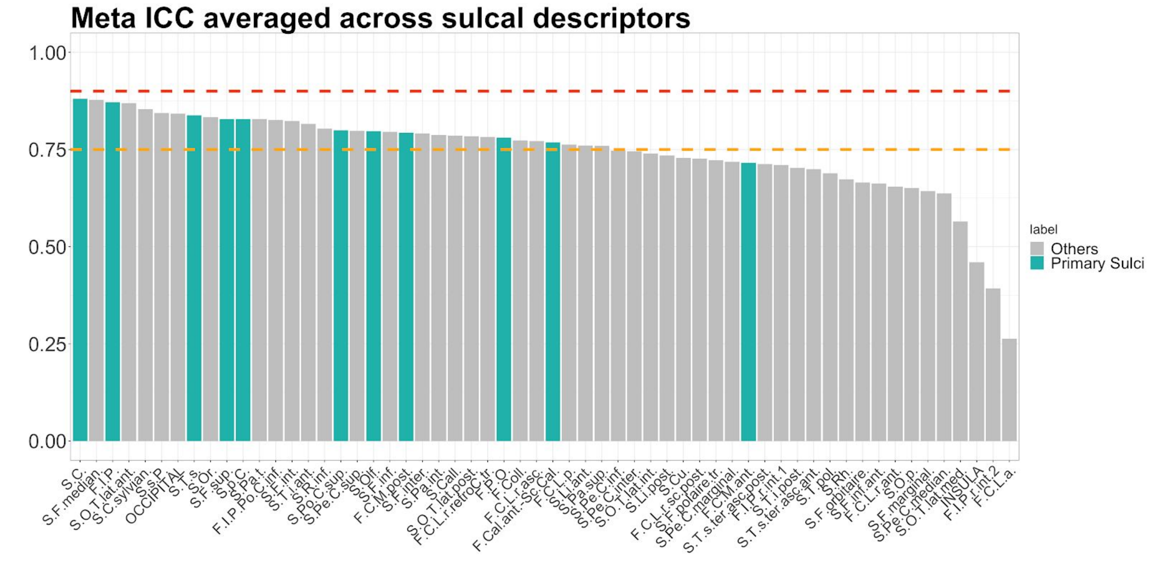


**Figure S8: Meta-Analysis:** Average across sulcal shape descriptors of meta-ICC computed for bilaterally averaged sulcal measures. Primary sulci are highlighted in green. The dashed orange and red lines highlight the ‘good’ (ICC=0.75) and ‘excellent’ (ICC=0.9) reliability thresholds [1]. On average the central sulcus (*S.C.*) seems the more reliable and the anterior lateral sulcus (*F.C.L.a.*) the less reliable.

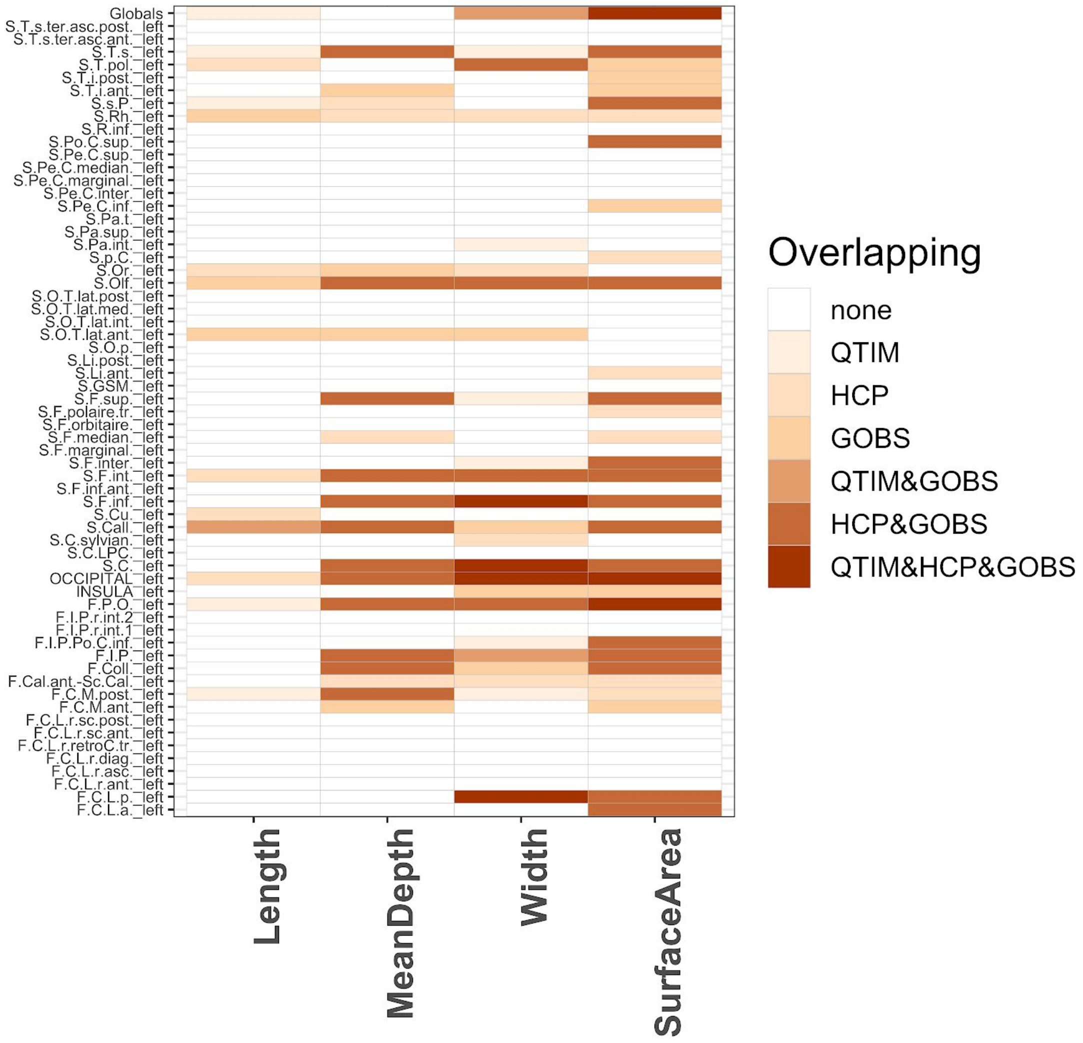


**Figure S9:** Sulci showing univariate $h^2$ overlap between QTIM HCP and GOBS **univariate h2** for the **left** hemisphere. Only the *Bonferroni* corrected results are reported. No overlap was found between QTIM and HCP only. Among others, regions like the *left central sulcus* (S.C._left) and the *left occipital area* (OCCIPITAL_left) show significant heritability for sulcal width for the three cohorts. **Figure S22**

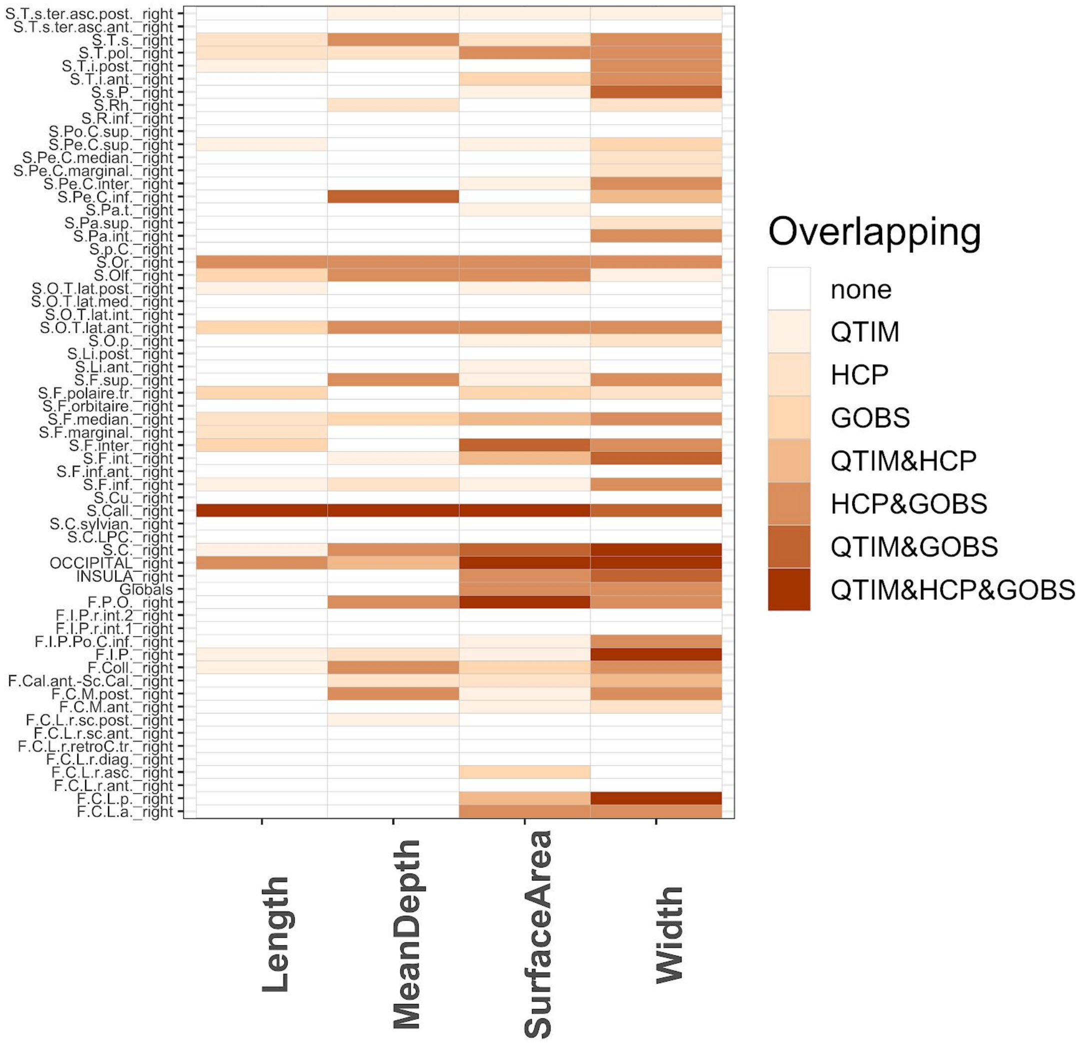


**Figure S10:** Sulci showing univariate $h^2$ overlap between QTIM HCP and GOBS **univariate h2** for the **right** hemisphere. Only the *Bonferroni* corrected results are reported. Among others, regions like the *right subcallosal sulcus* (S.Call._right) and the *right parieto-occipital fissure* (F.P.O._right) show significant heritability for sulcal length/mean depth and surface area the former, and the surfer area the latter, for the three cohorts.

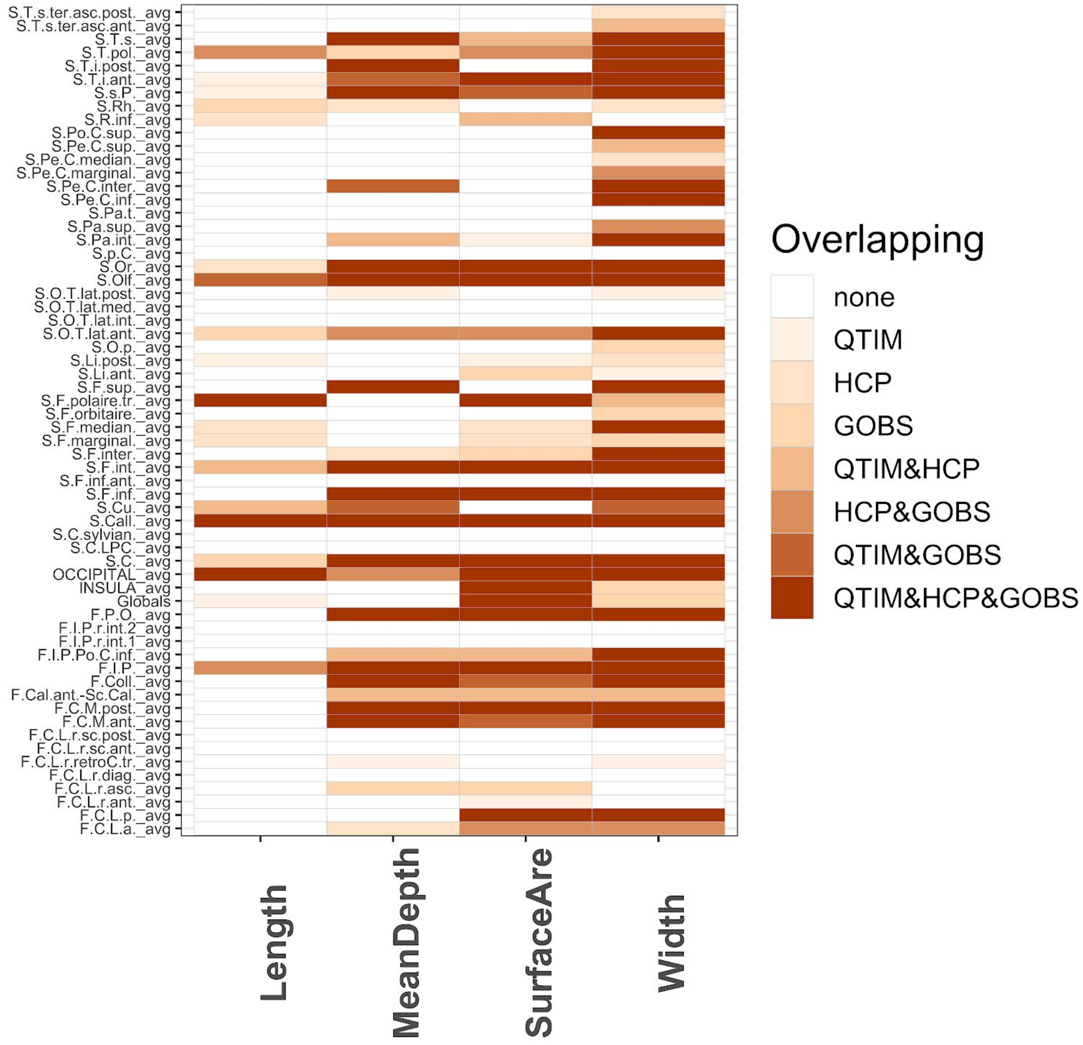


**Figure S11:** Sulci showing univariate $h^2$ overlap between QTIM HCP and GOBS **univariate h2** for the **bilaterally averaged sulci.** Only the *Bonferroni* corrected results are reported. Among others, regions like the *right subcallosal sulcus* (S.Call._right) and the *right parieto-occipital fissure* (F.P.O._right) show significant heritability for sulcal length/mean depth and surface area the former, and the surfer area the latter, for the three cohorts.

Heritability across sulci for each cohort and for Meta and Mega analysis

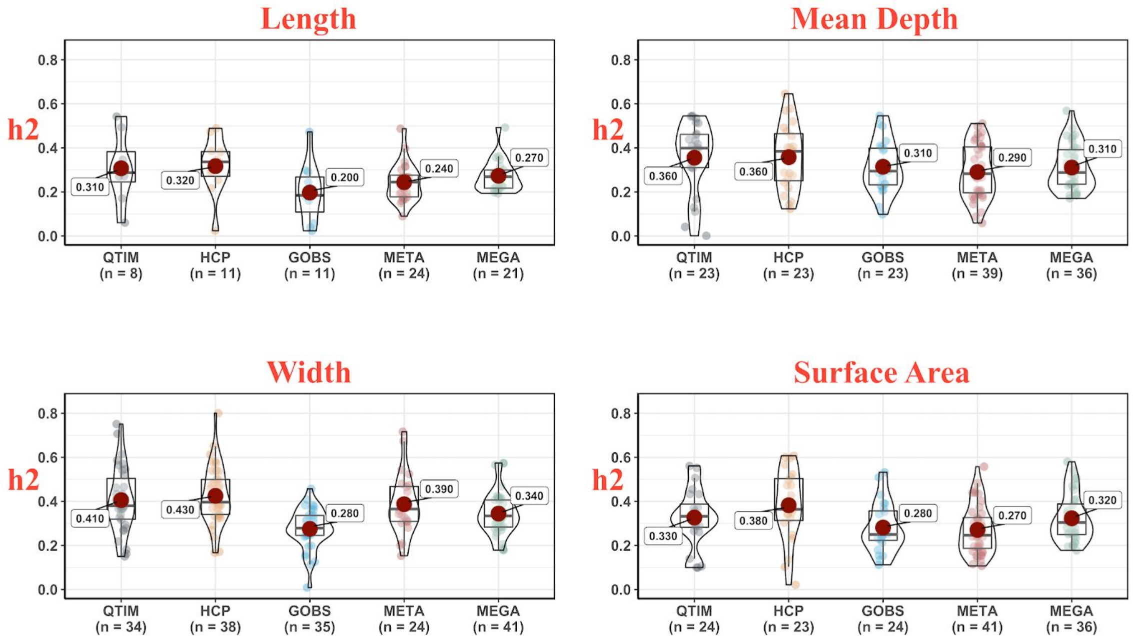


**Figure S12:** Violin/box plots of sulcal-based heritability (h2) for the bilaterally average sulcal descriptors, in QTIM, HCP, GOBS and for the meta- and mega-analyses. The average $h^2$ value is reported with the number of sulci surviving *Bonferroni* correction between parentheses.

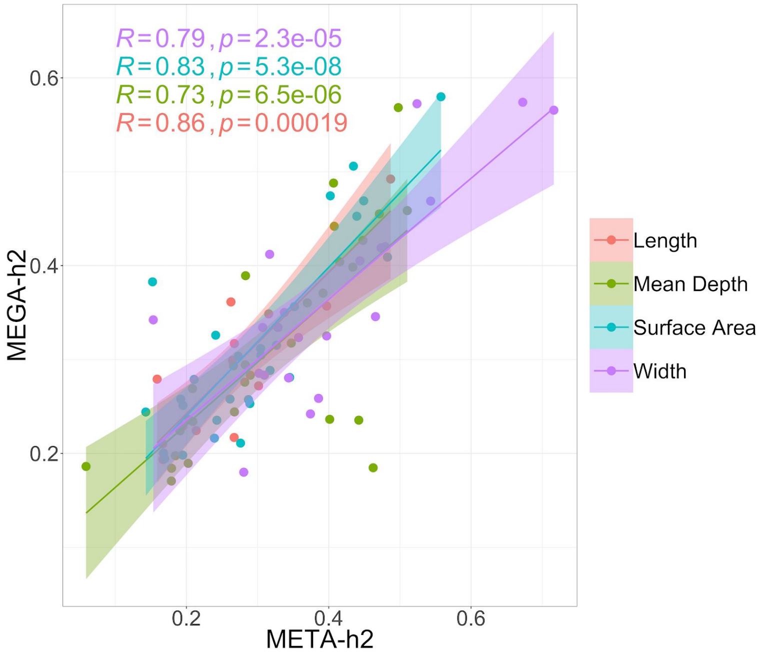


**Figure S13:** Scatter plots with Pearson's correlation coefficients estimated between Meta and Mega analysis, for sulcal length, mean depth, surface area, and width.

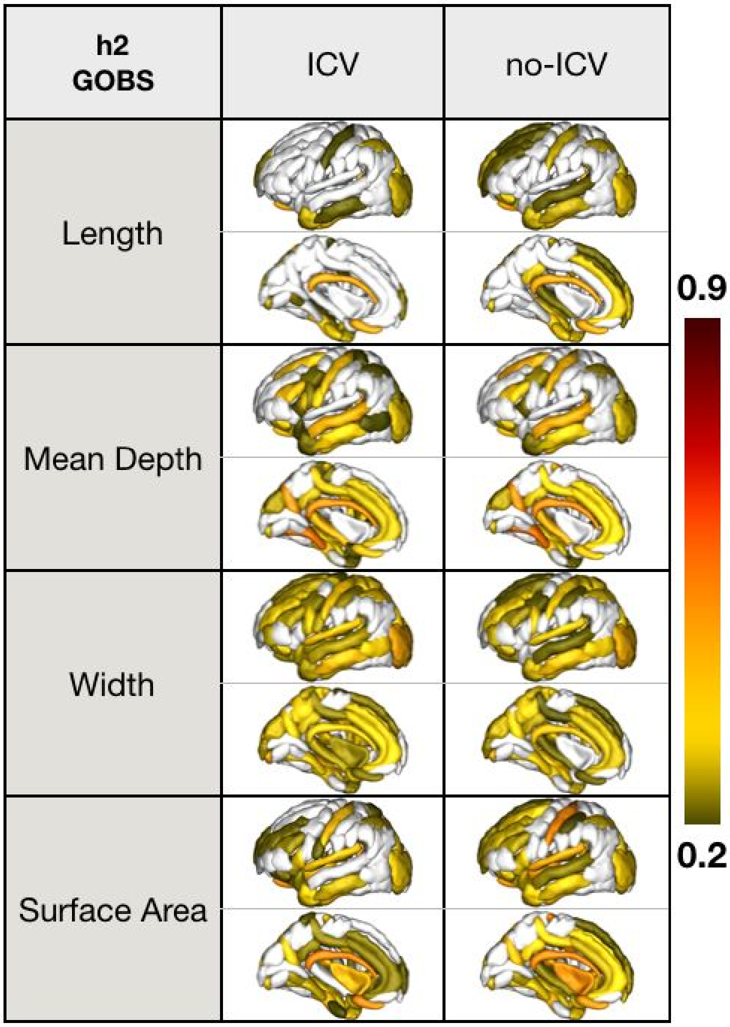


**Figure S14:** Sulcal-based h2 for length, mean depth, surface are and width, for GOBS, controlling for ICV (left) and without ICV (right). The genetic influence over many sulcal lengths appears to be driven by ICV; as surface area is also a function of the length, this pattern is also seen there. Covarying for ICV has relatively no effect on average sulcal depth or width.

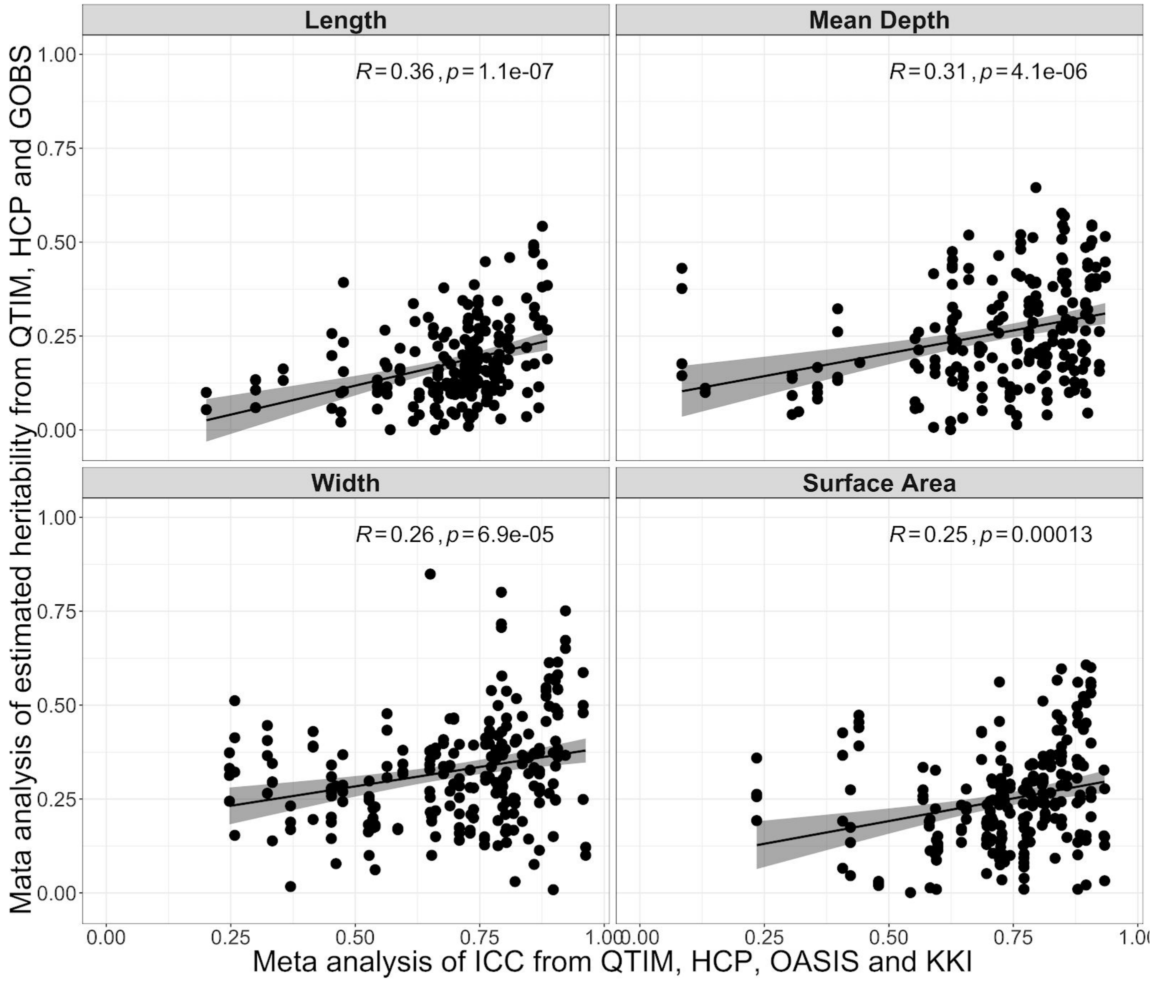


**Figure S15:** Scatter plots with Pearson's correlation coefficients estimated between Meta-h2 and Meta-ICC analysis, for bilaterally averaged measures of sulcal length, mean depth, surface area, and width.

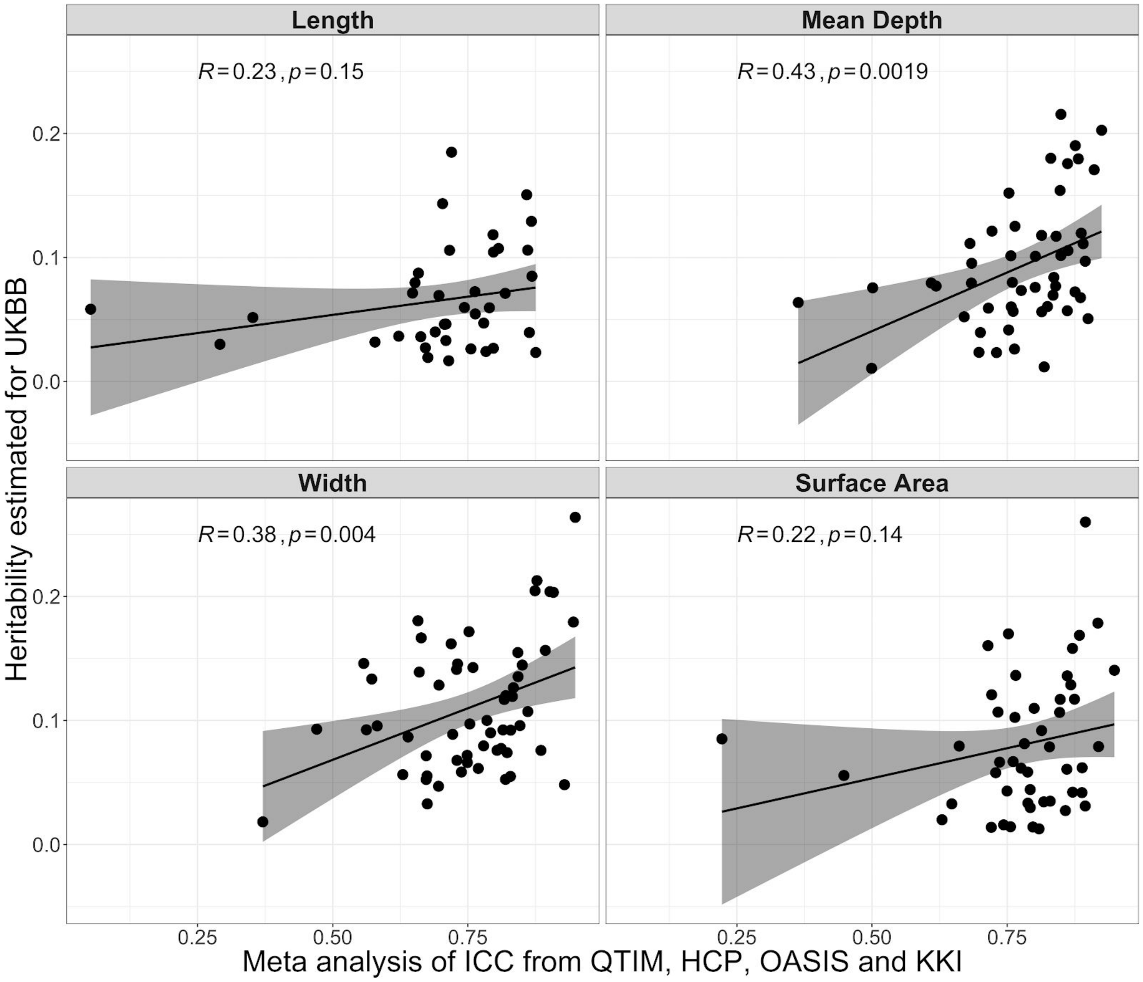


**Figure S16:** Scatter plots with Pearson's correlation coefficients estimated between h2 estimated in UKBB and Meta-ICC analysis, for bilaterally averaged measures of sulcal length, mean depth, surface area, and width.

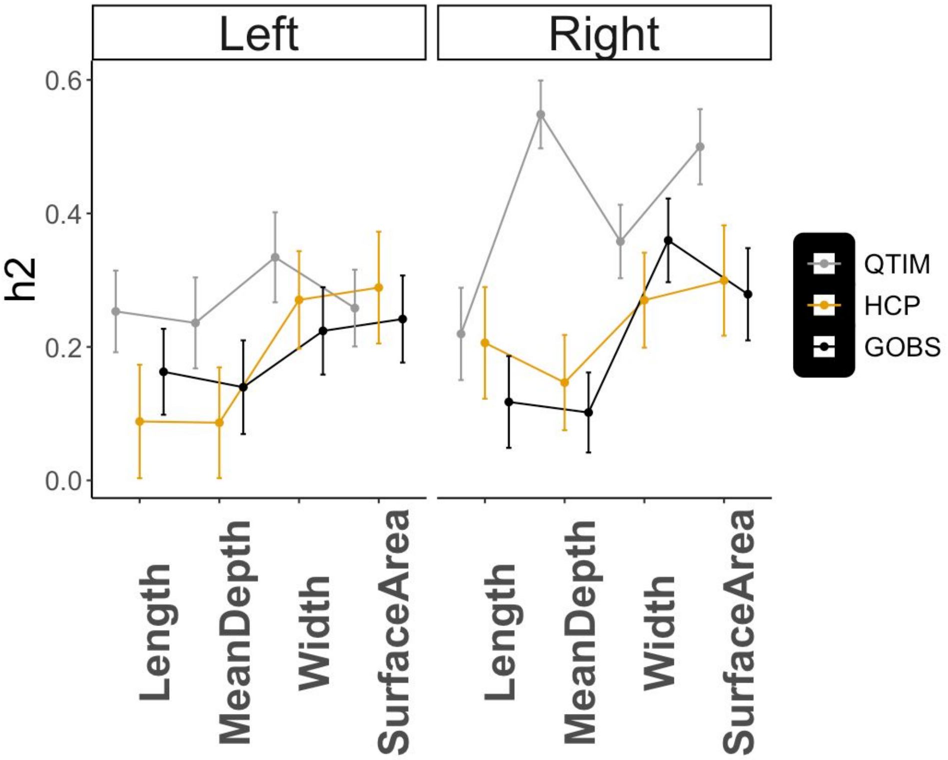


**Figure S17:** Univariate heritability ($h^2$) with standard error of global sulcal shape descriptor, for left and right hemisphere. The heritability was computed on the sum of sulcal length, mean depth, width and surface area across left and right sulci. QTIM, HCP and GOBS show similar trend across descriptor and hemispheres, except QTIM which seems to have higher heritability for the right hemisphere comparing to the left one.

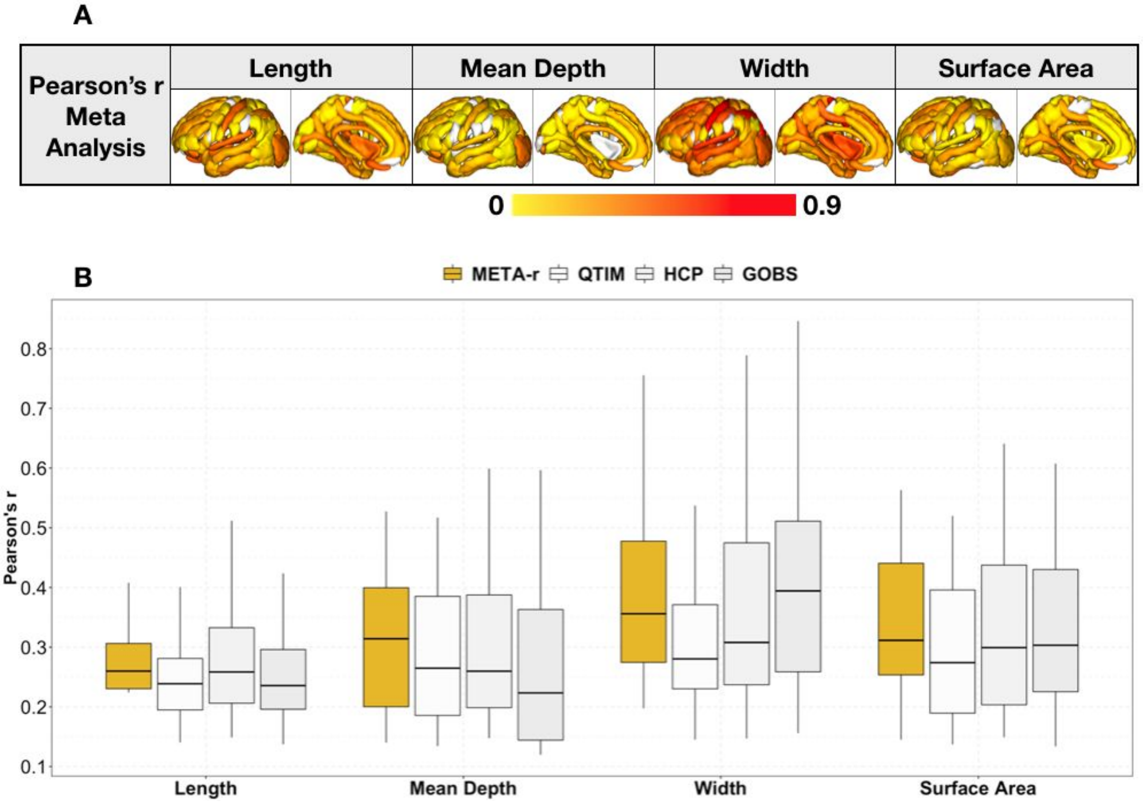


**Figure S18:** A) **Meta-analysis** of Pearson's correlation between left and right sulcal length, mean depth, surface area and width. B) Boxplots of Pearson's correlation between left and right sulcal measures for HCP, QTIM, GOBS and for the meta analysis (META-r) as mapped in A).

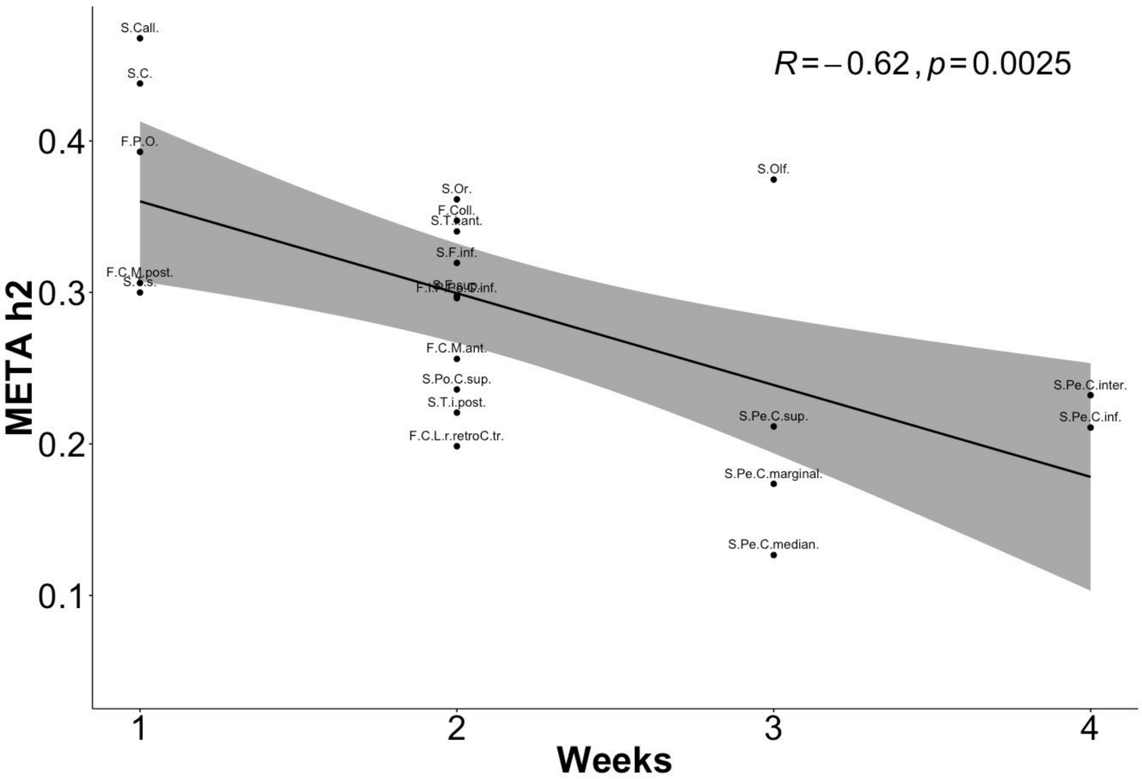


**Figure S19:** Pearson's correlation between heritability ($h^2$) and the appearance of sulci (in weeks). From Dubois et al. 2018 we grouped the sulci in four groups, 26.7w ("1"), 31.0w ("2"), 34.0w ("3") and 35.7w ("4") [2]. $h^2$ here has been computed as the average of the heritability of sulca length, mean depth, width and surface area, as estimated by the meta analysis of the bilaterally averages shape measures. The negative correlation (r=-0.62, p=0.0025) suggests that sulci appearing early in brain development are those showing higher estimated heritability.

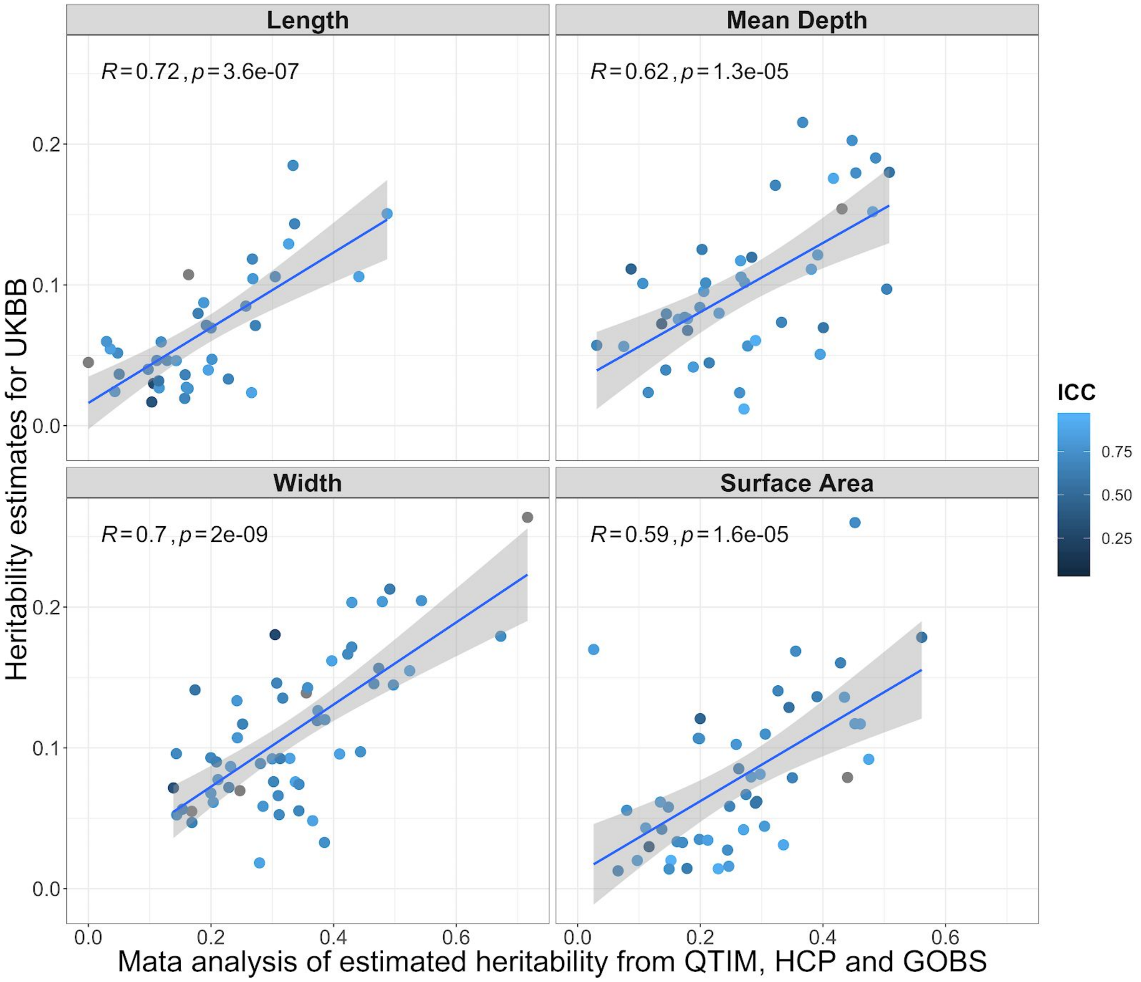


**Figure S20:** Pearson's correlation between heritability ($h^2$) estimated from UKBB sulcal shape descriptors and the meta analysis of heritability estimated from QTIM, HCP and GOBS. The scatter points are colored based on the ICC computed pooling together 4 test-retest. The results reported here refer to the bilaterally averaged measures of sulcal length, mean depth, width and surface area.

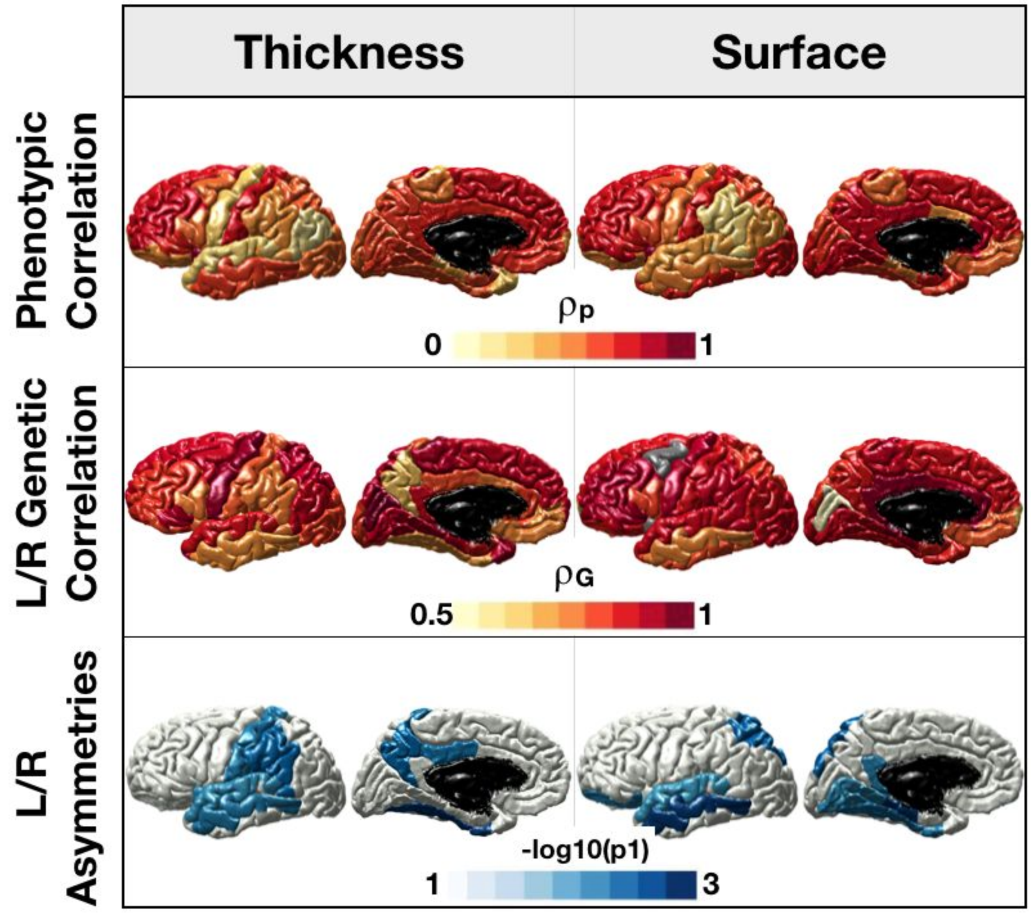


**Figure S21:** Top and middle rows show respectively the meta-analysis of bilaterally phenotypic correlation and the genetic correlation (equation 1 in main text ) between left and right thickness and surface values extracted from FreeSurfer ROIs (Desikan-Killiany atlas [3] ). For the meta-analysis only QTIM and HCP datasets have been used. The bottom row shows those regions for which the 95% confidence interval for genetic correlation did not include 1, identifying regions that show potential genetic asymmetries (p1 is the p-value when testing for differences from 1); our results may extend previous findings for cortical thickness of temporal lobe and postcentral gyrus and cortical surface of the temporal lobe and superior parietal lobe [4,5]. The results reported are *Bonferroni* corrected.

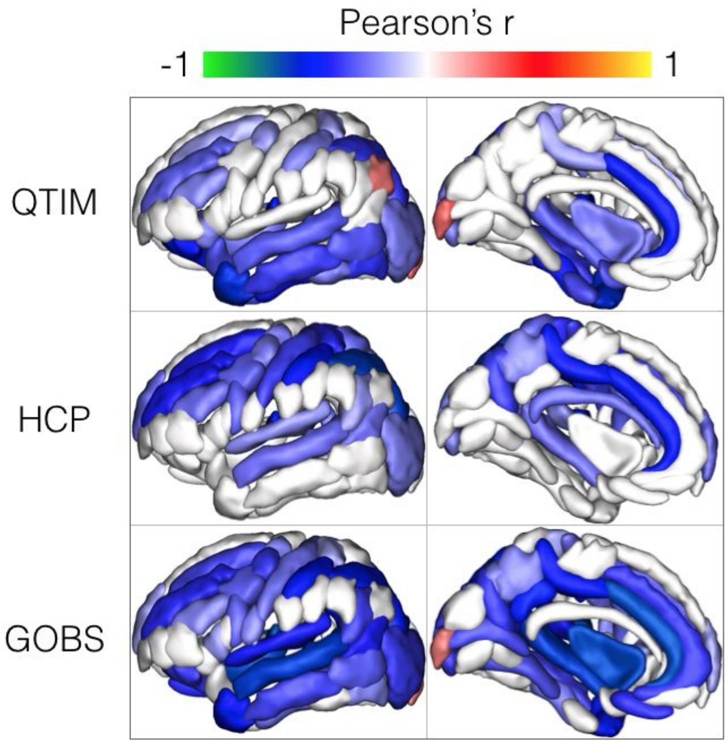


**Figure S22:** Pearson's correlation between sulcal width and grey matter thickness (GM), for QTIM, HCP and GOBS. For each sulcus the surrounding grey matter thickness has been estimated using BrainVISA Morphologist pipeline. A negative correlation between GM thickness and sulcal width suggests a differential trajectories of these two measures in brain development. For the cohorts analyzed here most of the sulci show a negative correlation (colorbar blue to green) between -0.2 and -0.6. Further investigation and longitudinal designs may better disentangle GM and sulcal with trajectories.
