## Supplementary Tables for "The reliability and heritability of cortical folds and their genetic correlations across hemispheres"

**Table SA.** Sulci nomenclature: BrainVISA sulcal labels are reported along with the anatomical nomenclature and the corresponding brain area.

| Brain regions | Sulcal Labels | Anatomical nomenclature |
|---|---|---|
| | **F.C.L.a** | Anterior lateral fissure |
| | **F.C.L.r.ant.** | Anterior ramus of lateral fissure |
| | **F.C.L.r.diag.** | Diagonal ramus of lateral fissure |
| | **F.C.L.r.sc.ant** | Anterior sub-central ramus of the lateral fissure |
| | **F.C.L.p.** | Posterior lateral fissure |
| | **F.C.L.r.asc.** | Ascending ramus of the lateral fissure |
| | **F.C.L.r.retroC.tr.** | Retro central transverse ramus of the lateral fissure |
| | **F.C.L.r.sc.post.** | Posterior sub-central ramus of the lateral fissure |
| **Lateral Fissures** | **S.C.sylvian** | Central sylvian sulcus |
| | **F.Cal.ant.-Sc.Cal.** | Calcarine fissure |
| **Calcarine** | **S.O.p.** | Occipito-polar sulcus |
| **Occipital Lobe** | **OCCIPITAL** | Occipital lobe |
| **Central Areas** | **S.C.** | Central sulcus |
| | **S.Call.** | Subcallosal sulcus |
| | **S.C.LPC.** | Paracentral lobule central sulcus |
| | **S.p.C.** | Paracentral sulcus |
| | **F.C.M.ant.** | Calloso-marginal ramus of the lateral fissure |
| | **F.C.M.post.** | Calloso-marginal posterior fissure |
| | **S.F.int.** | Internal frontal sulcus |
| **Internal Frontal Areas** | **S.R.inf.** | Inferior rostral sulcus |
| | **S.F.inf.** | Inferior frontal sulcus |
| | **S.F.marginal.** | Marginal frontal sulcus |
| | **S.F.orbitaire.** | Orbital frontal sulcus |
| | **S.F.sup.** | Superior frontal sulcus |
| | **S.F.median.** | Median frontal sulcus |
| | **S.Or.** | Orbital sulcus |
| | **S.F.inf.ant.** | Anterior inferior frontal sulcus |
| | **S.F.inter.** | Intermediate frontal sulcus |
| | **S.F.polaire.tr.** | Polar frontal sulcus |
| **Frontal Lobe** | **S.Olf.** | Olfactory sulcus |
| | **S.Li.ant.** | Anterior intralingual sulcus |
| | **S.s.P.** | Sub-parietal sulcus |
| | **S.Pa.int.** | Internal parietal sulcus |
| | **F.P.O.** | Parieto-occipital fissure |
| | **S.Cu.** | Cuneal sulcus |
| **Posterior Internal Regions** | **S.Li.post.** | Posterior intra-lingual sulcus |
| | **S.Pa.sup.** | Superior parietal sulcus |
| | **S.GSM.** | Sulcus of the supra-marginal gyrus |
| | **S.Po.C.sup.** | Superior postcentral sulcus |
| | **F.I.P.** | Intraparietal sulcus |
| | **F.I.P.Po.C.inf.** | Superior postcentral intraparietal superior sulcus |
| | **F.I.P.r.int.1** | Primary intermediate ramus of the intraparietal sulcus |
| | **F.I.P.r.int.2** | Secondary intermediate ramus of the intraparietal sulcus |
| **Parietal Lobe** | **S.Pa.t.** | Transverse parietal sulcus |
| | **S.Pe.C.inf.** | Inferior precentral sulcus |
| | **S.Pe.C.marginal.** | Marginal precentral sulcus |
| | **S.Pe.C.sup.** | Superior precentral sulcus |
| | **S.Pe.C.inter.** | Intermediate precentral sulcus |
| **Pre-central areas** | **S.Pe.C.median.** | Median precentral sulcus |
| | **S.T.i.ant.** | Anterior inferior temporal sulcus |
| | **S.T.pol.** | Polar temporal sulcus |
| | **S.T.s.** | Superior temporal sulcus |

| | | |
|---|---|---|
| External temporal lobe | **S.T.s.ter.asc.ant.** | Anterior terminal ascending branch of the superior temporal sulcus |
| | **S.T.s.ter.asc.post.** | Posterior terminal ascending branch of the superior temporal sulcus |
| | **S.T.i.post.** | Posterior inferior temporal sulcus |
| INSULA | **INSULA** | Insula |
| Basal Temporal Lobe | **S.O.T.lat.int.** | Internal occipito-temporal lateral sulcus |
| | **S.O.T.lat.post.** | Posterior occipito-temporal lateral sulcus |
| | **F.Coll.** | Collateral fissure |
| | **S.Rh.** | Rhinal sulcus |
| | **S.O.T.lat.ant.** | Anterior occipito-temporal lateral sulcus |
| | **S.O.T.lat.med.** | Median occipito-temporal lateral sulcus |

**Table S3:** ICC for the left hemisphere sulcal descriptors (yellow: ICC > 0.75; orange: ICC > 0.9)

| Sulci | Descriptor | QTIM | QTIM_SE | HCP | HCP_SE | OASIS | OASIS_SE | KKI | KKI_SE | Meta-SE-ICC |
|---|---|---|---|---|---|---|---|---|---|---|
| F.C.L.a._left | Length | 0.26 | 0.15 | 0.12 | 0.1 | 0.62 | 0.22 | 0.33 | 0.19 | 0.24 |
| F.C.L.a._left | MeanDepth | 0.14 | 0.11 | 0.16 | 0.12 | 0.71 | 0.21 | 0.15 | 0.14 | 0.21 |
| F.C.L.a._left | Width | 0.34 | 0.15 | 0.58 | 0.15 | 0.61 | 0.21 | 0.74 | 0.19 | 0.55 |
| F.C.L.a._left | SurfaceArea | 0.14 | 0.11 | 0.24 | 0.14 | 0.38 | 0.21 | 0.68 | 0.19 | 0.28 |
| F.C.L.p._left | Length | 0.63 | 0.15 | 0.75 | 0.14 | 0.61 | 0.2 | 0.8 | 0.17 | 0.71 |
| F.C.L.p._left | MeanDepth | 0.45 | 0.16 | 0.7 | 0.15 | 0.66 | 0.21 | 0.46 | 0.19 | 0.57 |
| F.C.L.p._left | Width | 0.74 | 0.15 | 0.79 | 0.14 | 0.85 | 0.18 | 0.93 | 0.15 | 0.82 |
| F.C.L.p._left | SurfaceArea | 0.5 | 0.16 | 0.81 | 0.14 | 0.83 | 0.18 | 0.84 | 0.17 | 0.74 |
| F.C.L.r.ant._left | Length | 0.5 | 0.17 | 0.52 | 0.16 | 0.43 | 0.21 | 0.82 | 0.17 | 0.57 |
| F.C.L.r.ant._left | MeanDepth | 0.32 | 0.15 | 0.54 | 0.15 | 0.69 | 0.2 | 0.62 | 0.19 | 0.52 |
| F.C.L.r.ant._left | Width | 0.58 | 0.16 | 0.75 | 0.14 | 0.52 | 0.21 | 0.93 | 0.15 | 0.72 |
| F.C.L.r.ant._left | SurfaceArea | 0.59 | 0.16 | 0.48 | 0.15 | 0.73 | 0.2 | 0.91 | 0.16 | 0.67 |
| F.C.L.r.asc._left | Length | 0.57 | 0.16 | 0.82 | 0.13 | 0.88 | 0.17 | 0.77 | 0.19 | 0.76 |
| F.C.L.r.asc._left | MeanDepth | 0.71 | 0.15 | 0.16 | 0.12 | 0.85 | 0.19 | 0.79 | 0.18 | 0.53 |
| F.C.L.r.asc._left | Width | 0.32 | 0.15 | 0.87 | 0.13 | 0.44 | 0.21 | 0.94 | 0.15 | 0.69 |
| F.C.L.r.asc._left | SurfaceArea | 0.57 | 0.15 | 0.63 | 0.15 | 0.75 | 0.19 | 0.86 | 0.17 | 0.69 |
| F.C.L.r.diag._left | Length | - | - | - | - | - | - | - | - | - |
| F.C.L.r.diag._left | MeanDepth | - | - | - | - | - | - | - | - | - |
| F.C.L.r.diag._left | Width | - | - | - | - | - | - | - | - | - |
| F.C.L.r.diag._left | SurfaceArea | - | - | - | - | - | - | - | - | - |
| F.C.L.r.retroC.tr._left | Length | 0.75 | 0.16 | 0.65 | 0.16 | 0.44 | 0.21 | 0.68 | 0.21 | 0.65 |
| F.C.L.r.retroC.tr._left | MeanDepth | 0.73 | 0.16 | 0.67 | 0.15 | 0.92 | 0.17 | 0.79 | 0.18 | 0.77 |
| F.C.L.r.retroC.tr._left | Width | 0.76 | 0.16 | 0.64 | 0.16 | 0.88 | 0.18 | 0.63 | 0.2 | 0.73 |
| F.C.L.r.retroC.tr._left | SurfaceArea | 0.85 | 0.14 | 0.7 | 0.15 | 0.86 | 0.18 | 0.87 | 0.17 | 0.82 |
| F.C.L.r.sc.ant._left | Length | - | - | - | - | - | - | - | - | - |
| F.C.L.r.sc.ant._left | MeanDepth | - | - | - | - | - | - | - | - | - |
| F.C.L.r.sc.ant._left | Width | - | - | - | - | - | - | - | - | - |
| F.C.L.r.sc.ant._left | SurfaceArea | - | - | - | - | - | - | - | - | - |
| F.C.L.r.sc.post._left | Length | - | - | - | - | - | - | - | - | - |
| F.C.L.r.sc.post._left | MeanDepth | 0.72 | 0.18 | 0.68 | 0.17 | 0.34 | 0.22 | 0.47 | 0.21 | 0.58 |
| F.C.L.r.sc.post._left | Width | 0.9 | 0.15 | 0.83 | 0.16 | 0.91 | 0.19 | 0.95 | 0.16 | 0.89 |
| F.C.L.r.sc.post._left | SurfaceArea | 0.73 | 0.18 | 0.63 | 0.17 | 0.93 | 0.19 | 0.55 | 0.22 | 0.72 |
| F.C.M.ant._left | Length | 0.61 | 0.16 | 0.66 | 0.15 | 0.77 | 0.19 | 0.63 | 0.19 | 0.66 |
| F.C.M.ant._left | MeanDepth | 0.62 | 0.16 | 0.69 | 0.15 | 0.91 | 0.16 | 0.77 | 0.18 | 0.74 |
| F.C.M.ant._left | Width | 0.62 | 0.16 | 0.58 | 0.16 | 0.67 | 0.2 | 0.57 | 0.2 | 0.61 |
| F.C.M.ant._left | SurfaceArea | 0.65 | 0.16 | 0.67 | 0.15 | 0.83 | 0.19 | 0.76 | 0.18 | 0.71 |
| F.C.M.post._left | Length | 0.5 | 0.16 | 0.48 | 0.15 | 0.43 | 0.2 | 0.7 | 0.19 | 0.52 |
| F.C.M.post._left | MeanDepth | 0.71 | 0.15 | 0.87 | 0.12 | 0.91 | 0.16 | 0.86 | 0.16 | 0.84 |
| F.C.M.post._left | Width | 0.87 | 0.13 | 0.8 | 0.13 | 0.8 | 0.19 | 0.82 | 0.17 | 0.83 |
| F.C.M.post._left | SurfaceArea | 0.66 | 0.15 | 0.58 | 0.15 | 0.8 | 0.19 | 0.9 | 0.16 | 0.72 |
| F.Cal.ant.-Sc.Cal._left | Length | 0.68 | 0.15 | 0.62 | 0.15 | 0.38 | 0.2 | 0.41 | 0.19 | 0.55 |
| F.Cal.ant.-Sc.Cal._left | MeanDepth | 0.55 | 0.15 | 0.73 | 0.15 | 0.8 | 0.19 | 0.57 | 0.19 | 0.66 |
| F.Cal.ant.-Sc.Cal._left | Width | 0.65 | 0.15 | 0.6 | 0.15 | 0.58 | 0.2 | 0.82 | 0.18 | 0.66 |
| F.Cal.ant.-Sc.Cal._left | SurfaceArea | 0.81 | 0.14 | 0.88 | 0.12 | 0.84 | 0.18 | 0.86 | 0.16 | 0.85 |
| F.Coll._left | Length | 0.89 | 0.13 | 0.51 | 0.15 | 0.29 | 0.18 | 0.57 | 0.2 | 0.63 |
| F.Coll._left | MeanDepth | 0.91 | 0.12 | 0.86 | 0.13 | 0.61 | 0.2 | 0.92 | 0.15 | 0.86 |
| F.Coll._left | Width | 0.74 | 0.14 | 0.78 | 0.14 | 0.14 | 0.13 | 0.71 | 0.19 | 0.56 |
| F.Coll._left | SurfaceArea | 0.95 | 0.11 | 0.61 | 0.15 | 0.48 | 0.2 | 0.62 | 0.19 | 0.75 |
| F.I.P._left | Length | 0.79 | 0.14 | 0.61 | 0.15 | 0.86 | 0.18 | 0.42 | 0.19 | 0.69 |
| F.I.P._left | MeanDepth | 0.88 | 0.13 | 0.82 | 0.13 | 0.69 | 0.2 | 0.88 | 0.16 | 0.83 |
| F.I.P._left | Width | 0.92 | 0.12 | 0.95 | 0.11 | 0.94 | 0.15 | 0.94 | 0.15 | 0.94 |
| F.I.P._left | SurfaceArea | 0.83 | 0.13 | 0.76 | 0.14 | 0.89 | 0.17 | 0.74 | 0.18 | 0.8 |
| F.I.P.Po.C.inf._left | Length | 0.84 | 0.13 | 0.75 | 0.14 | 0.67 | 0.21 | 0.52 | 0.2 | 0.73 |
| F.I.P.Po.C.inf._left | MeanDepth | 0.86 | 0.13 | 0.84 | 0.13 | 0.71 | 0.2 | 0.84 | 0.17 | 0.83 |
| F.I.P.Po.C.inf._left | Width | 0.88 | 0.13 | 0.9 | 0.12 | 0.87 | 0.18 | 0.94 | 0.15 | 0.9 |
| F.I.P.Po.C.inf._left | SurfaceArea | 0.83 | 0.14 | 0.83 | 0.13 | 0.73 | 0.2 | 0.44 | 0.2 | 0.76 |
| F.I.P.r.int.1_left | Length | 0.6 | 0.16 | 0.78 | 0.14 | 0.8 | 0.2 | 0.42 | 0.19 | 0.67 |
| F.I.P.r.int.1_left | MeanDepth | 0.81 | 0.14 | 0.73 | 0.15 | 0.95 | 0.16 | 0.82 | 0.17 | 0.82 |
| F.I.P.r.int.1_left | Width | 0.82 | 0.14 | 0.8 | 0.14 | 0.86 | 0.2 | 0.55 | 0.2 | 0.77 |
| F.I.P.r.int.1_left | SurfaceArea | 0.71 | 0.15 | 0.72 | 0.15 | 0.9 | 0.18 | 0.72 | 0.19 | 0.76 |
| F.I.P.r.int.2_left | Length | 0.3 | 0.16 | 0.46 | 0.17 | 0.72 | 0.22 | 0.17 | 0.14 | 0.36 |

| | | | | | | | | | | |
|---|---|---|---|---|---|---|---|---|---|---|
| F.I.P.r.int.2_left | MeanDepth | 0.33 | 0.16 | 0.69 | 0.16 | 0.86 | 0.19 | 0.71 | 0.19 | 0.62 |
| F.I.P.r.int.2_left | Width | 0.24 | 0.15 | 0.65 | 0.16 | 0.85 | 0.2 | 0.77 | 0.18 | 0.58 |
| F.I.P.r.int.2_left | SurfaceArea | 0.14 | 0.12 | 0.62 | 0.17 | 0.98 | 0.15 | 0.63 | 0.19 | 0.53 |
| F.P.O._left | Length | 0.69 | 0.15 | 0.72 | 0.15 | 0.88 | 0.17 | 0.35 | 0.19 | 0.68 |
| F.P.O._left | MeanDepth | 0.88 | 0.13 | 0.75 | 0.14 | 0.79 | 0.19 | 0.63 | 0.19 | 0.78 |
| F.P.O._left | Width | 0.92 | 0.12 | 0.86 | 0.13 | 0.81 | 0.19 | 0.84 | 0.17 | 0.87 |
| F.P.O._left | SurfaceArea | 0.69 | 0.15 | 0.78 | 0.14 | 0.92 | 0.16 | 0.45 | 0.19 | 0.73 |
| INSULA_left | Length | - | - | - | - | - | - | - | - | - |
| INSULA_left | MeanDepth | 0.24 | 0.14 | 0.04 | 0.05 | 0.49 | 0.21 | 0.27 | 0.17 | 0.1 |
| INSULA_left | Width | 0.28 | 0.15 | 0.48 | 0.15 | 0.46 | 0.2 | 0.75 | 0.18 | 0.47 |
| INSULA_left | SurfaceArea | 0.36 | 0.15 | 0.63 | 0.15 | 0.5 | 0.21 | 0.78 | 0.18 | 0.56 |
| OCCIPITAL_left | Length | 0.83 | 0.13 | 0.85 | 0.13 | 0.66 | 0.2 | 0.8 | 0.17 | 0.81 |
| OCCIPITAL_left | MeanDepth | 0.85 | 0.13 | 0.87 | 0.12 | 0.82 | 0.19 | 0.79 | 0.18 | 0.84 |
| OCCIPITAL_left | Width | 0.5 | 0.16 | 0.86 | 0.13 | 0.78 | 0.19 | 0.92 | 0.15 | 0.78 |
| OCCIPITAL_left | SurfaceArea | 0.86 | 0.13 | 0.88 | 0.12 | 0.73 | 0.2 | 0.74 | 0.19 | 0.83 |
| S.C._left | Length | 0.72 | 0.15 | 0.62 | 0.15 | 0.94 | 0.15 | 0.79 | 0.18 | 0.76 |
| S.C._left | MeanDepth | 0.68 | 0.15 | 0.88 | 0.12 | 0.62 | 0.2 | 0.88 | 0.16 | 0.79 |
| S.C._left | Width | 0.93 | 0.12 | 0.94 | 0.11 | 0.95 | 0.15 | 0.95 | 0.14 | 0.94 |
| S.C._left | SurfaceArea | 0.86 | 0.13 | 0.86 | 0.13 | 0.98 | 0.13 | 0.9 | 0.16 | 0.9 |
| S.C.LPC._left | Length | 0.85 | 0.18 | - | - | 0.51 | 0.24 | 0.75 | 0.21 | 0.73 |
| S.C.LPC._left | MeanDepth | 0.92 | 0.16 | - | - | 0.89 | 0.2 | 0.93 | 0.17 | 0.92 |
| S.C.LPC._left | Width | 0.73 | 0.2 | - | - | 0.8 | 0.23 | 0.96 | 0.16 | 0.85 |
| S.C.LPC._left | SurfaceArea | - | - | - | - | 0.65 | 0.24 | 0.88 | 0.2 | 0.78 |
| S.C.sylvian._left | Length | 0.54 | 0.21 | - | - | - | - | 0.9 | 0.19 | 0.73 |
| S.C.sylvian._left | MeanDepth | 0.93 | 0.16 | - | - | 0.75 | 0.24 | 0.9 | 0.18 | 0.88 |
| S.C.sylvian._left | Width | - | - | - | - | 0.93 | 0.2 | 0.84 | 0.19 | 0.88 |
| S.C.sylvian._left | SurfaceArea | - | - | - | - | 0.75 | 0.25 | 0.94 | 0.17 | 0.88 |
| S.Call._left | Length | 0.67 | 0.15 | 0.72 | 0.14 | 0.77 | 0.2 | 0.81 | 0.17 | 0.74 |
| S.Call._left | MeanDepth | 0.83 | 0.14 | 0.7 | 0.15 | 0.91 | 0.17 | 0.71 | 0.19 | 0.79 |
| S.Call._left | Width | 0.43 | 0.16 | 0.26 | 0.15 | 0.65 | 0.21 | 0.27 | 0.17 | 0.37 |
| S.Call._left | SurfaceArea | 0.67 | 0.15 | 0.71 | 0.14 | 0.75 | 0.2 | 0.78 | 0.18 | 0.72 |
| S.Cu._left | Length | 0.76 | 0.14 | 0.68 | 0.15 | 0.6 | 0.2 | 0.59 | 0.19 | 0.68 |
| S.Cu._left | MeanDepth | 0.73 | 0.15 | 0.66 | 0.15 | 0.28 | 0.18 | 0.95 | 0.14 | 0.7 |
| S.Cu._left | Width | 0.87 | 0.13 | 0.72 | 0.14 | 0.9 | 0.17 | 0.83 | 0.17 | 0.83 |
| S.Cu._left | SurfaceArea | 0.67 | 0.15 | 0.74 | 0.14 | 0.62 | 0.2 | 0.38 | 0.19 | 0.63 |
| S.F.inf._left | Length | 0.48 | 0.16 | 0.7 | 0.15 | 0.74 | 0.2 | 0.93 | 0.15 | 0.72 |
| S.F.inf._left | MeanDepth | 0.89 | 0.13 | 0.9 | 0.12 | 0.86 | 0.18 | 0.92 | 0.15 | 0.89 |
| S.F.inf._left | Width | 0.51 | 0.16 | 0.75 | 0.14 | 0.73 | 0.2 | 0.86 | 0.17 | 0.71 |
| S.F.inf._left | SurfaceArea | 0.8 | 0.14 | 0.82 | 0.13 | 0.73 | 0.2 | 0.96 | 0.13 | 0.84 |
| S.F.inf.ant._left | Length | 0.54 | 0.16 | 0.61 | 0.15 | 0.37 | 0.2 | 0.86 | 0.16 | 0.62 |
| S.F.inf.ant._left | MeanDepth | 0.67 | 0.16 | 0.68 | 0.15 | 0.59 | 0.21 | 0.77 | 0.18 | 0.68 |
| S.F.inf.ant._left | Width | 0.14 | 0.12 | 0.77 | 0.14 | 0.84 | 0.19 | 0.84 | 0.17 | 0.56 |
| S.F.inf.ant._left | SurfaceArea | 0.64 | 0.16 | 0.52 | 0.15 | 0.35 | 0.2 | 0.9 | 0.15 | 0.63 |
| S.F.int._left | Length | 0.71 | 0.15 | 0.88 | 0.12 | 0.86 | 0.18 | 0.81 | 0.17 | 0.82 |
| S.F.int._left | MeanDepth | 0.82 | 0.14 | 0.87 | 0.13 | 0.84 | 0.18 | 0.9 | 0.15 | 0.86 |
| S.F.int._left | Width | 0.7 | 0.15 | 0.43 | 0.15 | 0.64 | 0.2 | 0.71 | 0.18 | 0.61 |
| S.F.int._left | SurfaceArea | 0.78 | 0.14 | 0.91 | 0.12 | 0.87 | 0.18 | 0.91 | 0.15 | 0.87 |
| S.F.inter._left | Length | 0.77 | 0.14 | 0.82 | 0.13 | 0.66 | 0.2 | 0.91 | 0.15 | 0.81 |
| S.F.inter._left | MeanDepth | 0.75 | 0.15 | 0.88 | 0.12 | 0.59 | 0.2 | 0.72 | 0.18 | 0.77 |
| S.F.inter._left | Width | 0.88 | 0.13 | 0.79 | 0.14 | 0.7 | 0.2 | 0.67 | 0.19 | 0.79 |
| S.F.inter._left | SurfaceArea | 0.84 | 0.13 | 0.9 | 0.12 | 0.45 | 0.2 | 0.87 | 0.16 | 0.82 |
| S.F.marginal._left | Length | 0.33 | 0.15 | 0.74 | 0.14 | 0.74 | 0.2 | 0.81 | 0.17 | 0.64 |
| S.F.marginal._left | MeanDepth | 0.73 | 0.15 | 0.7 | 0.14 | 0.78 | 0.2 | 0.63 | 0.19 | 0.71 |
| S.F.marginal._left | Width | 0.46 | 0.16 | 0.62 | 0.15 | 0.48 | 0.22 | 0.79 | 0.18 | 0.59 |
| S.F.marginal._left | SurfaceArea | 0.48 | 0.16 | 0.65 | 0.15 | 0.77 | 0.2 | 0.78 | 0.18 | 0.65 |
| S.F.median._left | Length | 0.88 | 0.13 | 0.84 | 0.13 | 0.77 | 0.19 | 0.8 | 0.17 | 0.84 |
| S.F.median._left | MeanDepth | 0.91 | 0.12 | 0.53 | 0.15 | 0.85 | 0.18 | 0.75 | 0.18 | 0.78 |
| S.F.median._left | Width | 0.87 | 0.13 | 0.9 | 0.12 | 0.82 | 0.19 | 0.81 | 0.18 | 0.86 |
| S.F.median._left | SurfaceArea | 0.89 | 0.12 | 0.88 | 0.12 | 0.87 | 0.17 | 0.71 | 0.18 | 0.86 |
| S.F.orbitaire._left | Length | 0.5 | 0.16 | 0.69 | 0.16 | 0.72 | 0.21 | 0.57 | 0.2 | 0.61 |
| S.F.orbitaire._left | MeanDepth | 0.53 | 0.16 | 0.88 | 0.13 | 0.82 | 0.2 | 0.77 | 0.18 | 0.76 |
| S.F.orbitaire._left | Width | 0.15 | 0.12 | 0.87 | 0.13 | 0.52 | 0.22 | 0.68 | 0.19 | 0.51 |
| S.F.orbitaire._left | SurfaceArea | 0.74 | 0.15 | 0.83 | 0.14 | 0.79 | 0.2 | 0.55 | 0.19 | 0.74 |

| | | | | | | | | | | |
|---|---|---|---|---|---|---|---|---|---|---|
| S.F.polaire.tr._left | Length | 0.62 | 0.15 | 0.66 | 0.15 | 0.3 | 0.19 | 0.68 | 0.19 | 0.58 |
| S.F.polaire.tr._left | MeanDepth | 0.64 | 0.15 | 0.69 | 0.15 | 0.39 | 0.2 | 0.57 | 0.19 | 0.6 |
| S.F.polaire.tr._left | Width | 0.67 | 0.15 | 0.79 | 0.14 | 0.64 | 0.21 | 0.94 | 0.14 | 0.78 |
| S.F.polaire.tr._left | SurfaceArea | 0.62 | 0.15 | 0.6 | 0.15 | 0.19 | 0.16 | 0.69 | 0.19 | 0.51 |
| S.F.sup._left | Length | 0.43 | 0.16 | 0.55 | 0.15 | 0.66 | 0.2 | 0.85 | 0.16 | 0.61 |
| S.F.sup._left | MeanDepth | 0.92 | 0.12 | 0.84 | 0.13 | 0.81 | 0.19 | 0.9 | 0.16 | 0.88 |
| S.F.sup._left | Width | 0.85 | 0.13 | 0.94 | 0.11 | 0.86 | 0.18 | 0.79 | 0.18 | 0.88 |
| S.F.sup._left | SurfaceArea | 0.57 | 0.15 | 0.74 | 0.14 | 0.89 | 0.17 | 0.69 | 0.19 | 0.72 |
| S.GSM._left | Length | 0.55 | 0.18 | - | - | 0.92 | 0.2 | 0.92 | 0.18 | 0.79 |
| S.GSM._left | MeanDepth | 0.79 | 0.16 | - | - | 0.99 | 0.16 | 0.89 | 0.18 | 0.89 |
| S.GSM._left | Width | 0.81 | 0.16 | - | - | 0.87 | 0.21 | 0.77 | 0.2 | 0.81 |
| S.GSM._left | SurfaceArea | 0.74 | 0.17 | - | - | 0.86 | 0.23 | 0.93 | 0.17 | 0.84 |
| S.Li.ant._left | Length | 0.64 | 0.16 | 0.7 | 0.16 | 0.46 | 0.21 | 0.76 | 0.19 | 0.65 |
| S.Li.ant._left | MeanDepth | 0.75 | 0.15 | 0.9 | 0.13 | 0.52 | 0.21 | 0.75 | 0.19 | 0.77 |
| S.Li.ant._left | Width | 0.69 | 0.15 | 0.63 | 0.16 | 0.66 | 0.21 | 0.89 | 0.17 | 0.72 |
| S.Li.ant._left | SurfaceArea | 0.79 | 0.15 | 0.76 | 0.15 | 0.42 | 0.21 | 0.89 | 0.17 | 0.75 |
| S.Li.post._left | Length | 0.58 | 0.15 | 0.71 | 0.15 | 0.5 | 0.2 | 0.63 | 0.19 | 0.62 |
| S.Li.post._left | MeanDepth | 0.56 | 0.15 | 0.41 | 0.15 | 0.66 | 0.2 | 0.91 | 0.15 | 0.64 |
| S.Li.post._left | Width | 0.51 | 0.16 | 0.81 | 0.14 | 0.83 | 0.19 | 0.89 | 0.16 | 0.76 |
| S.Li.post._left | SurfaceArea | 0.67 | 0.15 | 0.67 | 0.15 | 0.49 | 0.2 | 0.8 | 0.17 | 0.67 |
| S.O.p._left | Length | 0.59 | 0.19 | 0.58 | 0.18 | 0.61 | 0.23 | 0.24 | 0.17 | 0.48 |
| S.O.p._left | MeanDepth | 0.13 | 0.12 | 0.72 | 0.17 | 0.83 | 0.21 | 0.89 | 0.18 | 0.5 |
| S.O.p._left | Width | 0.36 | 0.18 | - | - | 0.68 | 0.23 | 0.86 | 0.19 | 0.62 |
| S.O.p._left | SurfaceArea | 0.7 | 0.18 | 0.55 | 0.18 | 0.78 | 0.22 | 0.72 | 0.2 | 0.68 |
| S.O.T.lat.ant._left | Length | 0.72 | 0.15 | 0.92 | 0.12 | 0.84 | 0.18 | 0.94 | 0.14 | 0.87 |
| S.O.T.lat.ant._left | MeanDepth | 0.79 | 0.14 | 0.89 | 0.12 | 0.63 | 0.2 | 0.96 | 0.14 | 0.85 |
| S.O.T.lat.ant._left | Width | 0.49 | 0.16 | 0.65 | 0.15 | 0.75 | 0.2 | 0.9 | 0.16 | 0.69 |
| S.O.T.lat.ant._left | SurfaceArea | 0.86 | 0.13 | 0.96 | 0.1 | 0.91 | 0.16 | 0.98 | 0.12 | 0.94 |
| S.O.T.lat.int._left | Length | 0.62 | 0.16 | 0.82 | 0.16 | 0.78 | 0.2 | 0.89 | 0.17 | 0.78 |
| S.O.T.lat.int._left | MeanDepth | 0.66 | 0.16 | 0.44 | 0.17 | 0.82 | 0.19 | 0.64 | 0.2 | 0.63 |
| S.O.T.lat.int._left | Width | 0.6 | 0.17 | 0.55 | 0.18 | 0.95 | 0.16 | 0.84 | 0.18 | 0.74 |
| S.O.T.lat.int._left | SurfaceArea | 0.44 | 0.17 | 0.75 | 0.17 | 0.94 | 0.16 | 0.92 | 0.16 | 0.77 |
| S.O.T.lat.med._left | Length | 0.29 | 0.16 | 0.5 | 0.16 | 0.28 | 0.19 | 0.6 | 0.2 | 0.41 |
| S.O.T.lat.med._left | MeanDepth | 0.6 | 0.17 | 0.7 | 0.15 | 0.96 | 0.16 | 0.82 | 0.18 | 0.77 |
| S.O.T.lat.med._left | Width | 0.25 | 0.15 | 0.7 | 0.15 | 0.67 | 0.22 | 0.81 | 0.19 | 0.57 |
| S.O.T.lat.med._left | SurfaceArea | 0.37 | 0.16 | 0.78 | 0.14 | 0.62 | 0.22 | 0.72 | 0.19 | 0.63 |
| S.O.T.lat.post._left | Length | 0.67 | 0.15 | 0.65 | 0.15 | 0.87 | 0.19 | 0.84 | 0.17 | 0.74 |
| S.O.T.lat.post._left | MeanDepth | 0.88 | 0.13 | 0.77 | 0.14 | 0.86 | 0.18 | 0.93 | 0.15 | 0.86 |
| S.O.T.lat.post._left | Width | 0.31 | 0.15 | 0.75 | 0.14 | 0.74 | 0.2 | 0.89 | 0.17 | 0.66 |
| S.O.T.lat.post._left | SurfaceArea | 0.8 | 0.14 | 0.71 | 0.14 | 0.81 | 0.19 | 0.9 | 0.15 | 0.8 |
| S.Olf._left | Length | 0.42 | 0.16 | 0.72 | 0.14 | 0.8 | 0.19 | 0.69 | 0.18 | 0.64 |
| S.Olf._left | MeanDepth | 0.6 | 0.16 | 0.89 | 0.12 | 0.72 | 0.2 | 0.82 | 0.17 | 0.78 |
| S.Olf._left | Width | 0.43 | 0.16 | 0.54 | 0.15 | 0.81 | 0.19 | 0.83 | 0.17 | 0.63 |
| S.Olf._left | SurfaceArea | 0.62 | 0.15 | 0.89 | 0.12 | 0.95 | 0.15 | 0.82 | 0.17 | 0.83 |
| S.Or._left | Length | 0.68 | 0.15 | 0.83 | 0.13 | 0.81 | 0.19 | 0.87 | 0.16 | 0.8 |
| S.Or._left | MeanDepth | 0.8 | 0.14 | 0.46 | 0.15 | 0.94 | 0.16 | 0.94 | 0.15 | 0.79 |
| S.Or._left | Width | 0.55 | 0.16 | 0.68 | 0.15 | 0.74 | 0.19 | 0.87 | 0.17 | 0.7 |
| S.Or._left | SurfaceArea | 0.71 | 0.15 | 0.82 | 0.13 | 0.85 | 0.18 | 0.92 | 0.15 | 0.82 |
| S.p.C._left | Length | 0.54 | 0.17 | 0.82 | 0.14 | 0.86 | 0.18 | 0.68 | 0.2 | 0.73 |
| S.p.C._left | MeanDepth | 0.65 | 0.16 | 0.86 | 0.14 | 0.67 | 0.21 | 0.75 | 0.19 | 0.75 |
| S.p.C._left | Width | 0.58 | 0.17 | 0.9 | 0.13 | 0.82 | 0.2 | 0.83 | 0.18 | 0.79 |
| S.p.C._left | SurfaceArea | 0.41 | 0.17 | 0.87 | 0.14 | 0.87 | 0.19 | 0.69 | 0.19 | 0.72 |
| S.Pa.int._left | Length | 0.54 | 0.16 | 0.53 | 0.15 | 0.78 | 0.19 | 0.75 | 0.18 | 0.63 |
| S.Pa.int._left | MeanDepth | 0.75 | 0.15 | 0.73 | 0.15 | 0.88 | 0.18 | 0.88 | 0.16 | 0.8 |
| S.Pa.int._left | Width | 0.69 | 0.15 | 0.86 | 0.13 | 0.76 | 0.2 | 0.87 | 0.16 | 0.8 |
| S.Pa.int._left | SurfaceArea | 0.58 | 0.16 | 0.74 | 0.15 | 0.83 | 0.18 | 0.66 | 0.19 | 0.7 |
| S.Pa.sup._left | Length | 0.49 | 0.19 | 0.83 | 0.15 | 0.92 | 0.18 | 0.73 | 0.21 | 0.76 |
| S.Pa.sup._left | MeanDepth | 0.83 | 0.16 | 0.85 | 0.15 | 0.95 | 0.17 | 0.79 | 0.2 | 0.86 |
| S.Pa.sup._left | Width | 0.71 | 0.18 | 0.73 | 0.16 | 0.95 | 0.17 | 0.78 | 0.2 | 0.79 |
| S.Pa.sup._left | SurfaceArea | 0.42 | 0.19 | 0.83 | 0.15 | 0.97 | 0.16 | 0.88 | 0.18 | 0.8 |
| S.Pa.t._left | Length | - | - | - | - | 0.93 | 0.21 | - | - | - |
| S.Pa.t._left | MeanDepth | 0.9 | 0.16 | 0.9 | 0.15 | 0.93 | 0.18 | 0.67 | 0.21 | 0.87 |
| S.Pa.t._left | Width | 0.88 | 0.17 | 0.43 | 0.18 | 0.72 | 0.22 | 0.63 | 0.21 | 0.67 |

| | | | | | | | | | | |
|---|---|---|---|---|---|---|---|---|---|---|
| S.Pa.t._left | SurfaceArea | 0.73 | 0.2 | 0.77 | 0.17 | 0.99 | 0.14 | 0.85 | 0.2 | 0.86 |
| S.Pe.C.inf._left | Length | 0.63 | 0.16 | 0.71 | 0.15 | 0.64 | 0.22 | 0.88 | 0.18 | 0.71 |
| S.Pe.C.inf._left | MeanDepth | 0.76 | 0.15 | 0.81 | 0.14 | 0.9 | 0.18 | 0.69 | 0.2 | 0.8 |
| S.Pe.C.inf._left | Width | 0.59 | 0.16 | 0.82 | 0.14 | 0.63 | 0.21 | 0.86 | 0.19 | 0.74 |
| S.Pe.C.inf._left | SurfaceArea | 0.69 | 0.16 | 0.82 | 0.14 | 0.85 | 0.2 | 0.89 | 0.18 | 0.8 |
| S.Pe.C.inter._left | Length | 0.68 | 0.15 | 0.62 | 0.15 | 0.67 | 0.2 | 0.67 | 0.19 | 0.66 |
| S.Pe.C.inter._left | MeanDepth | 0.57 | 0.16 | 0.65 | 0.15 | 0.75 | 0.19 | 0.75 | 0.18 | 0.67 |
| S.Pe.C.inter._left | Width | 0.66 | 0.15 | 0.72 | 0.14 | 0.63 | 0.2 | 0.86 | 0.16 | 0.72 |
| S.Pe.C.inter._left | SurfaceArea | 0.7 | 0.15 | 0.78 | 0.14 | 0.59 | 0.2 | 0.51 | 0.19 | 0.68 |
| S.Pe.C.marginal._left | Length | 0.43 | 0.19 | 0.55 | 0.17 | - | - | 0.91 | 0.2 | 0.61 |
| S.Pe.C.marginal._left | MeanDepth | 0.9 | 0.14 | 0.75 | 0.15 | 0.86 | 0.24 | 0.84 | 0.19 | 0.83 |
| S.Pe.C.marginal._left | Width | 0.73 | 0.17 | 0.84 | 0.14 | 0.36 | 0.24 | 0.21 | 0.17 | 0.59 |
| S.Pe.C.marginal._left | SurfaceArea | 0.41 | 0.18 | 0.68 | 0.16 | 0.31 | 0.23 | 0.69 | 0.22 | 0.54 |
| S.Pe.C.median._left | Length | 0.73 | 0.16 | 0.81 | 0.14 | 0.21 | 0.17 | 0.64 | 0.2 | 0.62 |
| S.Pe.C.median._left | MeanDepth | 0.65 | 0.16 | 0.86 | 0.13 | 0.86 | 0.18 | 0.84 | 0.18 | 0.8 |
| S.Pe.C.median._left | Width | 0.74 | 0.15 | 0.86 | 0.13 | 0.95 | 0.16 | 0.74 | 0.19 | 0.83 |
| S.Pe.C.median._left | SurfaceArea | 0.62 | 0.16 | 0.85 | 0.13 | 0.54 | 0.21 | 0.65 | 0.2 | 0.71 |
| S.Pe.C.sup._left | Length | 0.77 | 0.16 | 0.56 | 0.15 | 0.86 | 0.19 | 0.78 | 0.19 | 0.72 |
| S.Pe.C.sup._left | MeanDepth | 0.86 | 0.15 | 0.53 | 0.15 | 0.49 | 0.22 | 0.55 | 0.21 | 0.64 |
| S.Pe.C.sup._left | Width | 0.86 | 0.15 | 0.78 | 0.14 | 0.8 | 0.2 | 0.86 | 0.18 | 0.82 |
| S.Pe.C.sup._left | SurfaceArea | 0.87 | 0.14 | 0.59 | 0.15 | 0.79 | 0.21 | 0.81 | 0.19 | 0.76 |
| S.Po.C.sup._left | Length | 0.87 | 0.13 | 0.74 | 0.14 | 0.67 | 0.2 | 0.45 | 0.19 | 0.73 |
| S.Po.C.sup._left | MeanDepth | 0.91 | 0.12 | 0.88 | 0.12 | 0.8 | 0.19 | 0.65 | 0.19 | 0.84 |
| S.Po.C.sup._left | Width | 0.84 | 0.14 | 0.89 | 0.12 | 0.92 | 0.17 | 0.88 | 0.16 | 0.88 |
| S.Po.C.sup._left | SurfaceArea | 0.89 | 0.13 | 0.81 | 0.13 | 0.72 | 0.2 | 0.47 | 0.19 | 0.77 |
| S.R.inf._left | Length | 0.62 | 0.16 | 0.8 | 0.16 | 0.98 | 0.16 | 0.83 | 0.18 | 0.81 |
| S.R.inf._left | MeanDepth | 0.79 | 0.15 | 0.9 | 0.14 | 0.92 | 0.2 | 0.95 | 0.15 | 0.89 |
| S.R.inf._left | Width | 0.46 | 0.17 | 0.73 | 0.17 | 0.78 | 0.23 | 0.71 | 0.2 | 0.65 |
| S.R.inf._left | SurfaceArea | 0.79 | 0.15 | 0.66 | 0.17 | 0.98 | 0.16 | 0.94 | 0.15 | 0.85 |
| S.Rh._left | Length | 0.66 | 0.16 | 0.37 | 0.16 | 0.25 | 0.19 | 0.5 | 0.2 | 0.46 |
| S.Rh._left | MeanDepth | 0.75 | 0.15 | 0.7 | 0.15 | 0.6 | 0.23 | 0.83 | 0.17 | 0.73 |
| S.Rh._left | Width | 0.57 | 0.16 | 0.56 | 0.16 | 0.76 | 0.22 | 0.69 | 0.19 | 0.62 |
| S.Rh._left | SurfaceArea | 0.78 | 0.15 | 0.7 | 0.15 | 0.23 | 0.18 | 0.71 | 0.19 | 0.63 |
| S.s.P._left | Length | 0.81 | 0.14 | 0.8 | 0.14 | 0.85 | 0.18 | 0.87 | 0.16 | 0.83 |
| S.s.P._left | MeanDepth | 0.92 | 0.12 | 0.89 | 0.12 | 0.82 | 0.18 | 0.96 | 0.13 | 0.91 |
| S.s.P._left | Width | 0.91 | 0.12 | 0.81 | 0.13 | 0.3 | 0.19 | 0.89 | 0.16 | 0.79 |
| S.s.P._left | SurfaceArea | 0.93 | 0.12 | 0.91 | 0.12 | 0.9 | 0.17 | 0.93 | 0.15 | 0.92 |
| S.T.i.ant._left | Length | 0.35 | 0.15 | 0.95 | 0.11 | 0.86 | 0.18 | 0.85 | 0.17 | 0.79 |
| S.T.i.ant._left | MeanDepth | 0.47 | 0.16 | 0.93 | 0.12 | 0.78 | 0.2 | 0.83 | 0.17 | 0.78 |
| S.T.i.ant._left | Width | 0.39 | 0.15 | 0.82 | 0.13 | 0.58 | 0.21 | 0.83 | 0.17 | 0.67 |
| S.T.i.ant._left | SurfaceArea | 0.49 | 0.16 | 0.97 | 0.1 | 0.85 | 0.18 | 0.89 | 0.16 | 0.85 |
| S.T.i.post._left | Length | 0.75 | 0.14 | 0.76 | 0.14 | 0.45 | 0.2 | 0.54 | 0.19 | 0.67 |
| S.T.i.post._left | MeanDepth | 0.8 | 0.14 | 0.38 | 0.15 | 0.73 | 0.2 | 0.83 | 0.17 | 0.68 |
| S.T.i.post._left | Width | 0.36 | 0.15 | 0.76 | 0.14 | 0.4 | 0.2 | 0.85 | 0.16 | 0.62 |
| S.T.i.post._left | SurfaceArea | 0.71 | 0.15 | 0.52 | 0.15 | 0.3 | 0.18 | 0.51 | 0.19 | 0.53 |
| S.T.pol._left | Length | 0.42 | 0.15 | 0.73 | 0.14 | 0.8 | 0.19 | 0.47 | 0.19 | 0.61 |
| S.T.pol._left | MeanDepth | 0.45 | 0.16 | 0.88 | 0.12 | 0.84 | 0.19 | 0.66 | 0.19 | 0.73 |
| S.T.pol._left | Width | 0.51 | 0.16 | 0.61 | 0.15 | 0.65 | 0.21 | 0.82 | 0.17 | 0.64 |
| S.T.pol._left | SurfaceArea | 0.49 | 0.16 | 0.76 | 0.14 | 0.82 | 0.19 | 0.48 | 0.19 | 0.64 |
| S.T.s._left | Length | 0.65 | 0.15 | 0.76 | 0.14 | 0.82 | 0.19 | 0.76 | 0.18 | 0.74 |
| S.T.s._left | MeanDepth | 0.74 | 0.14 | 0.83 | 0.13 | 0.72 | 0.2 | 0.85 | 0.17 | 0.79 |
| S.T.s._left | Width | 0.54 | 0.16 | 0.74 | 0.14 | 0.59 | 0.2 | 0.84 | 0.18 | 0.68 |
| S.T.s._left | SurfaceArea | 0.61 | 0.15 | 0.76 | 0.14 | 0.89 | 0.17 | 0.86 | 0.17 | 0.77 |
| S.T.s.ter.asc.ant._left | Length | 0.54 | 0.16 | 0.54 | 0.15 | 0.78 | 0.2 | 0.7 | 0.18 | 0.62 |
| S.T.s.ter.asc.ant._left | MeanDepth | 0.71 | 0.15 | 0.87 | 0.13 | 0.76 | 0.21 | 0.76 | 0.18 | 0.79 |
| S.T.s.ter.asc.ant._left | Width | 0.85 | 0.14 | 0.69 | 0.15 | 0.73 | 0.21 | 0.17 | 0.14 | 0.6 |
| S.T.s.ter.asc.ant._left | SurfaceArea | 0.63 | 0.16 | 0.57 | 0.15 | 0.91 | 0.18 | 0.6 | 0.19 | 0.67 |
| S.T.s.ter.asc.post._left | Length | 0.33 | 0.15 | 0.72 | 0.14 | 0.73 | 0.2 | 0.42 | 0.19 | 0.55 |
| S.T.s.ter.asc.post._left | MeanDepth | 0.73 | 0.15 | 0.35 | 0.15 | 0.78 | 0.2 | 0.57 | 0.19 | 0.59 |
| S.T.s.ter.asc.post._left | Width | 0.21 | 0.14 | 0.65 | 0.15 | 0.89 | 0.17 | 0.64 | 0.19 | 0.55 |
| S.T.s.ter.asc.post._left | SurfaceArea | 0.5 | 0.16 | 0.76 | 0.14 | 0.73 | 0.2 | 0.56 | 0.19 | 0.64 |

**Table S4:** ICC for the right hemisphere sulcal descriptors (yellow: ICC > 0.75; orange: ICC > 0.9)

| Sulci | Descriptor | QTIM | QTIM_SE | HCP | HCP_SE | OASIS | OASIS_SE | KKI | KKI_SE | Meta-SE-ICC |
|---|---|---|---|---|---|---|---|---|---|---|
| F.C.L.a._right | Length | 0.04 | 0.05 | 0.51 | 0.16 | 0.18 | 0.15 | 0.48 | 0.2 | 0.11 |
| F.C.L.a._right | MeanDepth | 0.44 | 0.16 | 0.52 | 0.16 | 0.15 | 0.14 | - | - | 0.35 |
| F.C.L.a._right | Width | 0.39 | 0.16 | 0.68 | 0.15 | 0.84 | 0.19 | 0.86 | 0.17 | 0.67 |
| F.C.L.a._right | SurfaceArea | 0.25 | 0.14 | 0.63 | 0.15 | 0.03 | 0.05 | 0.56 | 0.2 | 0.12 |
| F.C.L.p._right | Length | 0.72 | 0.15 | 0.76 | 0.14 | 0.79 | 0.19 | 0.8 | 0.17 | 0.76 |
| F.C.L.p._right | MeanDepth | 0.17 | 0.12 | 0.56 | 0.15 | 0.72 | 0.2 | 0.71 | 0.18 | 0.46 |
| F.C.L.p._right | Width | 0.84 | 0.13 | 0.88 | 0.12 | 0.79 | 0.19 | 0.93 | 0.15 | 0.87 |
| F.C.L.p._right | SurfaceArea | 0.65 | 0.16 | 0.89 | 0.12 | 0.9 | 0.17 | 0.95 | 0.14 | 0.86 |
| F.C.L.r.ant._right | Length | 0.58 | 0.18 | 0.61 | 0.17 | 0.85 | 0.21 | 0.75 | 0.2 | 0.68 |
| F.C.L.r.ant._right | MeanDepth | 0.63 | 0.17 | 0.83 | 0.15 | 0.75 | 0.22 | 0.7 | 0.19 | 0.74 |
| F.C.L.r.ant._right | Width | 0.61 | 0.17 | 0.71 | 0.16 | 0.34 | 0.21 | 0.87 | 0.17 | 0.67 |
| F.C.L.r.ant._right | SurfaceArea | 0.71 | 0.16 | 0.75 | 0.16 | 0.9 | 0.19 | 0.91 | 0.16 | 0.81 |
| F.C.L.r.asc._right | Length | 0.6 | 0.16 | 0.74 | 0.15 | 0.72 | 0.21 | 0.76 | 0.19 | 0.7 |
| F.C.L.r.asc._right | MeanDepth | 0.77 | 0.15 | 0.75 | 0.15 | 0.78 | 0.19 | 0.88 | 0.16 | 0.79 |
| F.C.L.r.asc._right | Width | 0.51 | 0.16 | 0.82 | 0.14 | 0.9 | 0.17 | 0.64 | 0.19 | 0.73 |
| F.C.L.r.asc._right | SurfaceArea | 0.72 | 0.15 | 0.78 | 0.14 | 0.86 | 0.18 | 0.91 | 0.15 | 0.81 |
| F.C.L.r.diag._right | Length | - | - | - | - | - | - | - | - | - |
| F.C.L.r.diag._right | MeanDepth | - | - | - | - | - | - | - | - | - |
| F.C.L.r.diag._right | Width | - | - | - | - | - | - | - | - | - |
| F.C.L.r.diag._right | SurfaceArea | - | - | - | - | - | - | - | - | - |
| F.C.L.r.retroC.tr._right | Length | 0.73 | 0.17 | 0.9 | 0.13 | 0.82 | 0.2 | 0.88 | 0.19 | 0.84 |
| F.C.L.r.retroC.tr._right | MeanDepth | 0.55 | 0.17 | 0.84 | 0.14 | 0.96 | 0.15 | 0.96 | 0.15 | 0.85 |
| F.C.L.r.retroC.tr._right | Width | 0.76 | 0.16 | 0.61 | 0.17 | 0.95 | 0.17 | 0.9 | 0.18 | 0.8 |
| F.C.L.r.retroC.tr._right | SurfaceArea | 0.83 | 0.16 | 0.89 | 0.13 | 0.88 | 0.18 | 0.97 | 0.15 | 0.89 |
| F.C.L.r.sc.ant._right | Length | - | - | - | - | - | - | - | - | - |
| F.C.L.r.sc.ant._right | MeanDepth | - | - | - | - | - | - | - | - | - |
| F.C.L.r.sc.ant._right | Width | - | - | - | - | - | - | - | - | - |
| F.C.L.r.sc.ant._right | SurfaceArea | - | - | - | - | - | - | - | - | - |
| F.C.L.r.sc.post._right | Length | - | - | - | - | - | - | - | - | - |
| F.C.L.r.sc.post._right | MeanDepth | 0.59 | 0.17 | 0.62 | 0.16 | 0.66 | 0.23 | 0.55 | 0.2 | 0.6 |
| F.C.L.r.sc.post._right | Width | 0.79 | 0.15 | 0.85 | 0.14 | 0.65 | 0.23 | 0.79 | 0.18 | 0.79 |
| F.C.L.r.sc.post._right | SurfaceArea | 0.8 | 0.15 | 0.75 | 0.15 | 0.17 | 0.16 | 0.4 | 0.2 | 0.55 |
| F.C.M.ant._right | Length | 0.58 | 0.16 | 0.68 | 0.15 | 0.79 | 0.19 | 0.93 | 0.15 | 0.74 |
| F.C.M.ant._right | MeanDepth | 0.74 | 0.15 | 0.9 | 0.12 | 0.57 | 0.21 | 0.93 | 0.15 | 0.82 |
| F.C.M.ant._right | Width | 0.66 | 0.16 | 0.22 | 0.14 | 0.79 | 0.19 | 0.51 | 0.2 | 0.5 |
| F.C.M.ant._right | SurfaceArea | 0.62 | 0.16 | 0.84 | 0.13 | 0.8 | 0.19 | 0.97 | 0.13 | 0.83 |
| F.C.M.post._right | Length | 0.47 | 0.16 | 0.67 | 0.15 | 0.42 | 0.2 | 0.65 | 0.19 | 0.57 |
| F.C.M.post._right | MeanDepth | 0.93 | 0.12 | 0.84 | 0.13 | 0.94 | 0.15 | 0.83 | 0.17 | 0.89 |
| F.C.M.post._right | Width | 0.92 | 0.12 | 0.8 | 0.14 | 0.72 | 0.2 | 0.9 | 0.16 | 0.85 |
| F.C.M.post._right | SurfaceArea | 0.58 | 0.16 | 0.77 | 0.14 | 0.56 | 0.21 | 0.84 | 0.17 | 0.7 |
| F.Cal.ant.-Sc.Cal._right | Length | 0.43 | 0.16 | 0.68 | 0.14 | 0.53 | 0.21 | 0.53 | 0.2 | 0.55 |
| F.Cal.ant.-Sc.Cal._right | MeanDepth | 0.37 | 0.16 | 0.9 | 0.12 | 0.93 | 0.16 | 0.78 | 0.18 | 0.77 |
| F.Cal.ant.-Sc.Cal._right | Width | 0.4 | 0.16 | 0.85 | 0.13 | 0.7 | 0.2 | 0.89 | 0.16 | 0.72 |
| F.Cal.ant.-Sc.Cal._right | SurfaceArea | 0.2 | 0.13 | 0.9 | 0.12 | 0.88 | 0.17 | 0.86 | 0.16 | 0.68 |
| F.Coll._right | Length | 0.76 | 0.14 | 0.54 | 0.15 | 0.33 | 0.2 | 0.85 | 0.17 | 0.65 |
| F.Coll._right | MeanDepth | 0.85 | 0.13 | 0.82 | 0.13 | 0.75 | 0.19 | 0.86 | 0.16 | 0.83 |
| F.Coll._right | Width | 0.59 | 0.15 | 0.52 | 0.16 | 0.42 | 0.2 | 0.73 | 0.19 | 0.57 |
| F.Coll._right | SurfaceArea | 0.81 | 0.14 | 0.63 | 0.15 | 0.73 | 0.2 | 0.89 | 0.16 | 0.77 |
| F.I.P._right | Length | 0.82 | 0.14 | 0.75 | 0.14 | 0.45 | 0.21 | 0.78 | 0.18 | 0.73 |
| F.I.P._right | MeanDepth | 0.79 | 0.14 | 0.68 | 0.15 | 0.71 | 0.2 | 0.94 | 0.14 | 0.79 |
| F.I.P._right | Width | 0.91 | 0.12 | 0.96 | 0.1 | 0.89 | 0.17 | 0.75 | 0.18 | 0.9 |
| F.I.P._right | SurfaceArea | 0.82 | 0.14 | 0.8 | 0.14 | 0.77 | 0.19 | 0.77 | 0.18 | 0.79 |
| F.I.P.Po.C.inf._right | Length | 0.68 | 0.15 | 0.8 | 0.14 | 0.39 | 0.2 | 0.93 | 0.15 | 0.75 |
| F.I.P.Po.C.inf._right | MeanDepth | 0.73 | 0.15 | 0.53 | 0.15 | 0.91 | 0.16 | 0.91 | 0.16 | 0.76 |
| F.I.P.Po.C.inf._right | Width | 0.86 | 0.13 | 0.86 | 0.13 | 0.83 | 0.19 | 0.9 | 0.16 | 0.86 |
| F.I.P.Po.C.inf._right | SurfaceArea | 0.68 | 0.15 | 0.74 | 0.14 | 0.51 | 0.2 | 0.95 | 0.14 | 0.76 |
| F.I.P.r.int.1_right | Length | 0.78 | 0.15 | 0.53 | 0.16 | 0.86 | 0.19 | 0.67 | 0.19 | 0.7 |
| F.I.P.r.int.1_right | MeanDepth | 0.44 | 0.16 | 0.73 | 0.14 | 0.87 | 0.19 | 0.31 | 0.18 | 0.59 |
| F.I.P.r.int.1_right | Width | 0.44 | 0.16 | 0.74 | 0.14 | 0.6 | 0.22 | 0.66 | 0.19 | 0.62 |
| F.I.P.r.int.1_right | SurfaceArea | 0.78 | 0.15 | 0.5 | 0.16 | 0.91 | 0.17 | 0.58 | 0.19 | 0.7 |
| F.I.P.r.int.2_right | Length | 0.82 | 0.17 | 0.84 | 0.15 | - | - | 0.64 | 0.24 | 0.79 |

| | | | | | | | | | | |
|---|---|---|---|---|---|---|---|---|---|---|
| F.I.P.r.int.2_right | MeanDepth | 0.75 | 0.17 | 0.88 | 0.14 | 0.71 | 0.26 | 0.94 | 0.18 | 0.84 |
| F.I.P.r.int.2_right | Width | 0.77 | 0.18 | 0.8 | 0.15 | - | - | 0.57 | 0.24 | 0.75 |
| F.I.P.r.int.2_right | SurfaceArea | 0.76 | 0.17 | 0.88 | 0.14 | - | - | 0.7 | 0.24 | 0.81 |
| F.P.O._right | Length | 0.28 | 0.15 | 0.67 | 0.15 | 0.58 | 0.2 | 0.79 | 0.17 | 0.56 |
| F.P.O._right | MeanDepth | 0.27 | 0.15 | 0.87 | 0.12 | 0.3 | 0.19 | 0.53 | 0.2 | 0.55 |
| F.P.O._right | Width | 0.75 | 0.15 | 0.87 | 0.12 | 0.86 | 0.18 | 0.88 | 0.16 | 0.84 |
| F.P.O._right | SurfaceArea | 0.68 | 0.15 | 0.91 | 0.12 | 0.88 | 0.17 | 0.88 | 0.16 | 0.84 |
| INSULA_right | Length | - | - | - | - | - | - | - | - | - |
| INSULA_right | MeanDepth | 0.07 | 0.08 | 0.49 | 0.15 | 0.14 | 0.13 | 0.17 | 0.14 | 0.16 |
| INSULA_right | Width | 0.26 | 0.15 | 0.49 | 0.15 | 0.67 | 0.2 | 0.83 | 0.17 | 0.52 |
| INSULA_right | SurfaceArea | 0.56 | 0.16 | 0.71 | 0.14 | 0.66 | 0.2 | 0.67 | 0.19 | 0.65 |
| OCCIPITAL_right | Length | 0.77 | 0.14 | 0.92 | 0.11 | 0.74 | 0.19 | 0.79 | 0.17 | 0.83 |
| OCCIPITAL_right | MeanDepth | 0.86 | 0.13 | 0.78 | 0.14 | 0.66 | 0.2 | 0.86 | 0.16 | 0.81 |
| OCCIPITAL_right | Width | 0.77 | 0.14 | 0.89 | 0.12 | 0.74 | 0.19 | 0.88 | 0.16 | 0.84 |
| OCCIPITAL_right | SurfaceArea | 0.77 | 0.14 | 0.89 | 0.12 | 0.82 | 0.18 | 0.83 | 0.17 | 0.83 |
| S.C._right | Length | 0.7 | 0.15 | 0.85 | 0.13 | 0.86 | 0.18 | 0.75 | 0.18 | 0.79 |
| S.C._right | MeanDepth | 0.79 | 0.14 | 0.71 | 0.14 | 0.79 | 0.19 | 0.82 | 0.17 | 0.77 |
| S.C._right | Width | 0.92 | 0.12 | 0.94 | 0.11 | 0.97 | 0.14 | 0.95 | 0.14 | 0.94 |
| S.C._right | SurfaceArea | 0.87 | 0.13 | 0.97 | 0.1 | 0.97 | 0.14 | 0.91 | 0.15 | 0.94 |
| S.C.LPC._right | Length | - | - | - | - | - | - | - | - | - |
| S.C.LPC._right | MeanDepth | - | - | - | - | - | - | - | - | - |
| S.C.LPC._right | Width | - | - | - | - | - | - | - | - | - |
| S.C.LPC._right | SurfaceArea | - | - | - | - | - | - | - | - | - |
| S.C.sylvian._right | Length | 0.76 | 0.19 | 0.89 | 0.15 | 0.87 | 0.19 | 0.65 | 0.21 | 0.82 |
| S.C.sylvian._right | MeanDepth | 0.84 | 0.16 | 0.95 | 0.13 | 0.98 | 0.14 | 0.72 | 0.19 | 0.9 |
| S.C.sylvian._right | Width | 0.69 | 0.18 | 0.94 | 0.13 | 0.91 | 0.18 | 0.77 | 0.18 | 0.85 |
| S.C.sylvian._right | SurfaceArea | 0.93 | 0.14 | 0.96 | 0.13 | 0.97 | 0.15 | 0.92 | 0.16 | 0.95 |
| S.Call._right | Length | 0.85 | 0.13 | 0.85 | 0.13 | 0.65 | 0.21 | 0.86 | 0.16 | 0.82 |
| S.Call._right | MeanDepth | 0.78 | 0.14 | 0.89 | 0.12 | 0.85 | 0.18 | 0.86 | 0.17 | 0.85 |
| S.Call._right | Width | 0.55 | 0.15 | 0.36 | 0.15 | 0.37 | 0.2 | 0.23 | 0.16 | 0.38 |
| S.Call._right | SurfaceArea | 0.92 | 0.12 | 0.9 | 0.12 | 0.7 | 0.2 | 0.92 | 0.15 | 0.89 |
| S.Cu._right | Length | 0.37 | 0.16 | 0.75 | 0.14 | 0.64 | 0.2 | 0.76 | 0.18 | 0.63 |
| S.Cu._right | MeanDepth | 0.55 | 0.16 | 0.79 | 0.14 | 0.71 | 0.2 | 0.64 | 0.19 | 0.68 |
| S.Cu._right | Width | 0.52 | 0.16 | 0.6 | 0.15 | 0.78 | 0.19 | 0.97 | 0.13 | 0.73 |
| S.Cu._right | SurfaceArea | 0.51 | 0.16 | 0.81 | 0.13 | 0.76 | 0.19 | 0.85 | 0.17 | 0.73 |
| S.F.inf._right | Length | 0.51 | 0.16 | 0.8 | 0.14 | 0.73 | 0.2 | 0.72 | 0.18 | 0.69 |
| S.F.inf._right | MeanDepth | 0.72 | 0.15 | 0.97 | 0.1 | 0.85 | 0.18 | 0.86 | 0.16 | 0.88 |
| S.F.inf._right | Width | 0.71 | 0.15 | 0.85 | 0.13 | 0.67 | 0.2 | 0.83 | 0.17 | 0.78 |
| S.F.inf._right | SurfaceArea | 0.44 | 0.16 | 0.93 | 0.11 | 0.71 | 0.2 | 0.61 | 0.19 | 0.73 |
| S.F.inf.ant._right | Length | 0.36 | 0.15 | 0.48 | 0.16 | 0.64 | 0.2 | 0.49 | 0.19 | 0.47 |
| S.F.inf.ant._right | MeanDepth | 0.59 | 0.16 | 0.88 | 0.12 | 0.73 | 0.2 | 0.42 | 0.19 | 0.7 |
| S.F.inf.ant._right | Width | 0.64 | 0.15 | 0.78 | 0.14 | 0.58 | 0.21 | 0.83 | 0.17 | 0.73 |
| S.F.inf.ant._right | SurfaceArea | 0.37 | 0.16 | 0.8 | 0.14 | 0.59 | 0.2 | 0.45 | 0.19 | 0.58 |
| S.F.int._right | Length | 0.7 | 0.15 | 0.81 | 0.14 | 0.79 | 0.19 | 0.89 | 0.16 | 0.8 |
| S.F.int._right | MeanDepth | 0.7 | 0.15 | 0.78 | 0.14 | 0.86 | 0.18 | 0.93 | 0.15 | 0.81 |
| S.F.int._right | Width | 0.59 | 0.15 | 0.68 | 0.14 | 0.7 | 0.2 | 0.79 | 0.18 | 0.68 |
| S.F.int._right | SurfaceArea | 0.67 | 0.15 | 0.78 | 0.14 | 0.74 | 0.2 | 0.94 | 0.14 | 0.79 |
| S.F.inter._right | Length | 0.37 | 0.16 | 0.71 | 0.14 | 0.5 | 0.21 | 0.37 | 0.19 | 0.51 |
| S.F.inter._right | MeanDepth | 0.57 | 0.15 | 0.8 | 0.14 | 0.86 | 0.18 | 0.92 | 0.15 | 0.79 |
| S.F.inter._right | Width | 0.52 | 0.16 | 0.77 | 0.14 | 0.74 | 0.2 | 0.72 | 0.18 | 0.69 |
| S.F.inter._right | SurfaceArea | 0.45 | 0.16 | 0.77 | 0.14 | 0.65 | 0.2 | 0.57 | 0.2 | 0.62 |
| S.F.marginal._right | Length | 0.65 | 0.15 | 0.68 | 0.15 | 0.55 | 0.21 | 0.68 | 0.19 | 0.65 |
| S.F.marginal._right | MeanDepth | 0.69 | 0.15 | 0.72 | 0.14 | 0.5 | 0.2 | 0.57 | 0.2 | 0.64 |
| S.F.marginal._right | Width | 0.49 | 0.16 | 0.72 | 0.14 | 0.53 | 0.21 | 0.72 | 0.19 | 0.62 |
| S.F.marginal._right | SurfaceArea | 0.54 | 0.16 | 0.73 | 0.14 | 0.34 | 0.19 | 0.83 | 0.17 | 0.63 |
| S.F.median._right | Length | 0.76 | 0.14 | 0.87 | 0.12 | 0.8 | 0.19 | 0.83 | 0.17 | 0.82 |
| S.F.median._right | MeanDepth | 0.92 | 0.12 | 0.83 | 0.13 | 0.76 | 0.2 | 0.76 | 0.18 | 0.84 |
| S.F.median._right | Width | 0.83 | 0.14 | 0.86 | 0.13 | 0.83 | 0.19 | 0.89 | 0.16 | 0.85 |
| S.F.median._right | SurfaceArea | 0.76 | 0.14 | 0.9 | 0.12 | 0.76 | 0.19 | 0.85 | 0.17 | 0.83 |
| S.F.orbitaire._right | Length | 0.37 | 0.16 | 0.56 | 0.16 | 0.79 | 0.2 | 0.82 | 0.18 | 0.61 |
| S.F.orbitaire._right | MeanDepth | 0.57 | 0.17 | 0.79 | 0.14 | 0.93 | 0.17 | 0.85 | 0.18 | 0.78 |
| S.F.orbitaire._right | Width | 0.66 | 0.17 | 0.68 | 0.16 | 0.71 | 0.21 | 0.83 | 0.19 | 0.71 |
| S.F.orbitaire._right | SurfaceArea | 0.37 | 0.16 | 0.6 | 0.16 | 0.87 | 0.19 | 0.89 | 0.17 | 0.67 |

| | | | | | | | | | | |
|---|---|---|---|---|---|---|---|---|---|---|
| S.F.polaire.tr._right | Length | 0.42 | 0.15 | 0.62 | 0.15 | 0.63 | 0.2 | 0.66 | 0.19 | 0.57 |
| S.F.polaire.tr._right | MeanDepth | 0.72 | 0.15 | 0.48 | 0.15 | 0.81 | 0.19 | 0.88 | 0.16 | 0.71 |
| S.F.polaire.tr._right | Width | 0.38 | 0.15 | 0.84 | 0.13 | 0.88 | 0.17 | 0.94 | 0.15 | 0.77 |
| S.F.polaire.tr._right | SurfaceArea | 0.44 | 0.16 | 0.55 | 0.15 | 0.63 | 0.21 | 0.77 | 0.18 | 0.58 |
| S.F.sup._right | Length | 0.77 | 0.14 | 0.75 | 0.14 | 0.52 | 0.21 | 0.78 | 0.18 | 0.73 |
| S.F.sup._right | MeanDepth | 0.83 | 0.13 | 0.93 | 0.11 | 0.72 | 0.2 | 0.95 | 0.14 | 0.88 |
| S.F.sup._right | Width | 0.62 | 0.15 | 0.91 | 0.12 | 0.74 | 0.2 | 0.85 | 0.17 | 0.8 |
| S.F.sup._right | SurfaceArea | 0.78 | 0.14 | 0.8 | 0.13 | 0.51 | 0.21 | 0.81 | 0.18 | 0.75 |
| S.Li.ant._right | Length | 0.56 | 0.17 | 0.67 | 0.15 | 0.77 | 0.2 | 0.85 | 0.17 | 0.71 |
| S.Li.ant._right | MeanDepth | 0.89 | 0.13 | 0.95 | 0.11 | 0.93 | 0.17 | 0.3 | 0.18 | 0.83 |
| S.Li.ant._right | Width | 0.73 | 0.16 | 0.88 | 0.13 | 0.85 | 0.2 | 0.81 | 0.18 | 0.82 |
| S.Li.ant._right | SurfaceArea | 0.74 | 0.16 | 0.64 | 0.16 | 0.88 | 0.18 | 0.75 | 0.18 | 0.74 |
| S.Li.post._right | Length | 0.77 | 0.14 | 0.55 | 0.15 | 0.5 | 0.2 | 0.61 | 0.19 | 0.63 |
| S.Li.post._right | MeanDepth | 0.67 | 0.15 | 0.61 | 0.15 | 0.64 | 0.2 | 0.72 | 0.18 | 0.66 |
| S.Li.post._right | Width | 0.72 | 0.15 | 0.75 | 0.15 | 0.84 | 0.18 | 0.81 | 0.18 | 0.77 |
| S.Li.post._right | SurfaceArea | 0.81 | 0.14 | 0.54 | 0.15 | 0.49 | 0.21 | 0.79 | 0.17 | 0.68 |
| S.O.p._right | Length | 0.42 | 0.16 | 0.55 | 0.16 | 0.78 | 0.2 | 0.5 | 0.2 | 0.55 |
| S.O.p._right | MeanDepth | 0.51 | 0.16 | 0.51 | 0.16 | 0.74 | 0.2 | 0.46 | 0.2 | 0.55 |
| S.O.p._right | Width | 0.52 | 0.16 | 0.58 | 0.16 | 0.67 | 0.21 | 0.96 | 0.14 | 0.71 |
| S.O.p._right | SurfaceArea | 0.55 | 0.16 | 0.46 | 0.16 | 0.86 | 0.18 | 0.33 | 0.19 | 0.55 |
| S.O.T.lat.ant._right | Length | 0.54 | 0.16 | 0.78 | 0.14 | 0.54 | 0.21 | 0.72 | 0.18 | 0.66 |
| S.O.T.lat.ant._right | MeanDepth | 0.91 | 0.12 | 0.85 | 0.13 | 0.86 | 0.18 | 0.88 | 0.16 | 0.88 |
| S.O.T.lat.ant._right | Width | 0.67 | 0.16 | 0.7 | 0.15 | 0.67 | 0.21 | 0.89 | 0.16 | 0.74 |
| S.O.T.lat.ant._right | SurfaceArea | 0.88 | 0.13 | 0.9 | 0.12 | 0.8 | 0.19 | 0.87 | 0.16 | 0.87 |
| S.O.T.lat.int._right | Length | 0.77 | 0.17 | 0.19 | 0.14 | 0.62 | 0.22 | 0.69 | 0.2 | 0.51 |
| S.O.T.lat.int._right | MeanDepth | 0.69 | 0.17 | 0.78 | 0.16 | 0.62 | 0.22 | 0.77 | 0.19 | 0.72 |
| S.O.T.lat.int._right | Width | 0.46 | 0.18 | 0.79 | 0.16 | 0.73 | 0.21 | 0.67 | 0.2 | 0.67 |
| S.O.T.lat.int._right | SurfaceArea | 0.67 | 0.17 | 0.63 | 0.17 | 0.46 | 0.22 | 0.81 | 0.19 | 0.66 |
| S.O.T.lat.med._right | Length | 0.21 | 0.14 | 0.81 | 0.14 | - | - | 0.87 | 0.16 | 0.61 |
| S.O.T.lat.med._right | MeanDepth | 0.71 | 0.16 | 0.95 | 0.11 | 0.38 | 0.21 | 0.91 | 0.16 | 0.82 |
| S.O.T.lat.med._right | Width | 0.59 | 0.17 | 0.79 | 0.14 | 0.61 | 0.22 | 0.76 | 0.18 | 0.7 |
| S.O.T.lat.med._right | SurfaceArea | 0.29 | 0.16 | 0.89 | 0.13 | 0.14 | 0.13 | 0.9 | 0.16 | 0.56 |
| S.O.T.lat.post._right | Length | 0.58 | 0.15 | 0.66 | 0.15 | 0.35 | 0.19 | 0.33 | 0.18 | 0.51 |
| S.O.T.lat.post._right | MeanDepth | 0.82 | 0.14 | 0.78 | 0.14 | 0.63 | 0.2 | 0.8 | 0.17 | 0.78 |
| S.O.T.lat.post._right | Width | 0.56 | 0.16 | 0.72 | 0.15 | 0.51 | 0.22 | 0.79 | 0.17 | 0.66 |
| S.O.T.lat.post._right | SurfaceArea | 0.6 | 0.15 | 0.67 | 0.15 | 0.58 | 0.2 | 0.46 | 0.19 | 0.59 |
| S.Olf._right | Length | 0.61 | 0.15 | 0.73 | 0.14 | 0.34 | 0.2 | 0.68 | 0.19 | 0.61 |
| S.Olf._right | MeanDepth | 0.74 | 0.14 | 0.9 | 0.12 | 0.81 | 0.19 | 0.86 | 0.16 | 0.84 |
| S.Olf._right | Width | 0.58 | 0.16 | 0.64 | 0.15 | 0.85 | 0.18 | 0.45 | 0.19 | 0.63 |
| S.Olf._right | SurfaceArea | 0.77 | 0.14 | 0.9 | 0.12 | 0.48 | 0.21 | 0.82 | 0.17 | 0.79 |
| S.Or._right | Length | 0.49 | 0.16 | 0.82 | 0.13 | 0.83 | 0.18 | 0.84 | 0.17 | 0.74 |
| S.Or._right | MeanDepth | 0.48 | 0.16 | 0.49 | 0.15 | 0.89 | 0.17 | 0.93 | 0.15 | 0.7 |
| S.Or._right | Width | 0.61 | 0.15 | 0.8 | 0.14 | 0.8 | 0.19 | 0.73 | 0.18 | 0.74 |
| S.Or._right | SurfaceArea | 0.67 | 0.15 | 0.88 | 0.12 | 0.94 | 0.15 | 0.87 | 0.16 | 0.84 |
| S.p.C._right | Length | 0.56 | 0.17 | 0.6 | 0.17 | 0.82 | 0.2 | 0.93 | 0.15 | 0.73 |
| S.p.C._right | MeanDepth | 0.47 | 0.16 | 0.67 | 0.16 | 0.94 | 0.16 | 0.86 | 0.17 | 0.73 |
| S.p.C._right | Width | 0.4 | 0.17 | 0.88 | 0.14 | 0.86 | 0.19 | 0.83 | 0.18 | 0.75 |
| S.p.C._right | SurfaceArea | 0.51 | 0.17 | 0.81 | 0.15 | 0.98 | 0.14 | 0.96 | 0.14 | 0.84 |
| S.Pa.int._right | Length | 0.25 | 0.14 | 0.7 | 0.15 | 0.86 | 0.18 | 0.64 | 0.19 | 0.57 |
| S.Pa.int._right | MeanDepth | 0.46 | 0.16 | 0.82 | 0.14 | 0.89 | 0.17 | 0.89 | 0.16 | 0.76 |
| S.Pa.int._right | Width | 0.68 | 0.15 | 0.66 | 0.15 | 0.9 | 0.17 | 0.84 | 0.17 | 0.76 |
| S.Pa.int._right | SurfaceArea | 0.31 | 0.15 | 0.83 | 0.14 | 0.81 | 0.19 | 0.79 | 0.18 | 0.67 |
| S.Pa.sup._right | Length | 0.61 | 0.17 | 0.52 | 0.16 | 0.68 | 0.23 | 0.59 | 0.2 | 0.59 |
| S.Pa.sup._right | MeanDepth | 0.77 | 0.15 | 0.5 | 0.16 | 0.9 | 0.19 | 0.91 | 0.16 | 0.76 |
| S.Pa.sup._right | Width | 0.68 | 0.16 | 0.94 | 0.12 | 0.86 | 0.2 | 0.88 | 0.17 | 0.85 |
| S.Pa.sup._right | SurfaceArea | 0.58 | 0.16 | 0.46 | 0.16 | 0.55 | 0.24 | 0.88 | 0.17 | 0.62 |
| S.Pa.t._right | Length | 0.56 | 0.18 | - | - | 0.9 | 0.19 | 0.78 | 0.2 | 0.74 |
| S.Pa.t._right | MeanDepth | 0.76 | 0.15 | 0.56 | 0.16 | 0.73 | 0.21 | 0.83 | 0.18 | 0.72 |
| S.Pa.t._right | Width | 0.87 | 0.14 | 0.56 | 0.16 | 0.67 | 0.22 | 0.73 | 0.19 | 0.72 |
| S.Pa.t._right | SurfaceArea | 0.77 | 0.15 | 0.77 | 0.15 | 0.96 | 0.16 | 0.86 | 0.17 | 0.83 |
| S.Pe.C.inf._right | Length | 0.65 | 0.16 | 0.7 | 0.16 | 0.61 | 0.22 | 0.86 | 0.18 | 0.71 |
| S.Pe.C.inf._right | MeanDepth | 0.59 | 0.16 | 0.79 | 0.15 | 0.84 | 0.19 | 0.57 | 0.2 | 0.7 |
| S.Pe.C.inf._right | Width | 0.5 | 0.16 | 0.81 | 0.15 | 0.55 | 0.22 | 0.89 | 0.17 | 0.71 |

| | | | | | | | | | | |
|---|---|---|---|---|---|---|---|---|---|---|
| S.Pe.C.inf._right | SurfaceArea | 0.68 | 0.15 | 0.69 | 0.16 | 0.68 | 0.21 | 0.88 | 0.17 | 0.74 |
| S.Pe.C.inter._right | Length | 0.52 | 0.16 | 0.83 | 0.13 | 0.49 | 0.2 | 0.6 | 0.19 | 0.65 |
| S.Pe.C.inter._right | MeanDepth | 0.68 | 0.15 | 0.76 | 0.14 | 0.61 | 0.2 | 0.41 | 0.19 | 0.65 |
| S.Pe.C.inter._right | Width | 0.71 | 0.15 | 0.93 | 0.11 | 0.85 | 0.18 | 0.84 | 0.17 | 0.85 |
| S.Pe.C.inter._right | SurfaceArea | 0.53 | 0.16 | 0.82 | 0.13 | 0.48 | 0.2 | 0.55 | 0.2 | 0.64 |
| S.Pe.C.marginal._right | Length | 0.36 | 0.17 | 0.7 | 0.15 | 0.34 | 0.2 | 0.78 | 0.19 | 0.56 |
| S.Pe.C.marginal._right | MeanDepth | 0.6 | 0.16 | 0.8 | 0.14 | 0.34 | 0.2 | 0.79 | 0.18 | 0.66 |
| S.Pe.C.marginal._right | Width | 0.52 | 0.16 | 0.93 | 0.12 | 0.69 | 0.21 | 0.88 | 0.16 | 0.79 |
| S.Pe.C.marginal._right | SurfaceArea | 0.55 | 0.16 | 0.76 | 0.15 | 0.22 | 0.17 | 0.84 | 0.17 | 0.6 |
| S.Pe.C.median._right | Length | 0.7 | 0.16 | 0.84 | 0.14 | 0.77 | 0.2 | 0.52 | 0.2 | 0.73 |
| S.Pe.C.median._right | MeanDepth | 0.62 | 0.16 | 0.89 | 0.13 | 0.87 | 0.17 | 0.86 | 0.17 | 0.82 |
| S.Pe.C.median._right | Width | 0.73 | 0.15 | 0.91 | 0.13 | 0.88 | 0.18 | 0.91 | 0.16 | 0.86 |
| S.Pe.C.median._right | SurfaceArea | 0.53 | 0.16 | 0.87 | 0.13 | 0.88 | 0.18 | 0.67 | 0.2 | 0.75 |
| S.Pe.C.sup._right | Length | 0.89 | 0.14 | 0.74 | 0.14 | 0.2 | 0.17 | 0.82 | 0.17 | 0.69 |
| S.Pe.C.sup._right | MeanDepth | 0.87 | 0.14 | 0.89 | 0.12 | 0.33 | 0.21 | 0.87 | 0.16 | 0.81 |
| S.Pe.C.sup._right | Width | 0.89 | 0.14 | 0.92 | 0.12 | 0.66 | 0.23 | 0.76 | 0.18 | 0.85 |
| S.Pe.C.sup._right | SurfaceArea | 0.9 | 0.13 | 0.8 | 0.14 | 0.07 | 0.08 | 0.85 | 0.17 | 0.45 |
| S.Po.C.sup._right | Length | 0.62 | 0.16 | 0.72 | 0.15 | 0.5 | 0.21 | 0.93 | 0.15 | 0.72 |
| S.Po.C.sup._right | MeanDepth | 0.91 | 0.13 | 0.91 | 0.12 | 0.65 | 0.21 | 0.86 | 0.17 | 0.87 |
| S.Po.C.sup._right | Width | 0.82 | 0.15 | 0.92 | 0.12 | 0.72 | 0.2 | 0.91 | 0.16 | 0.86 |
| S.Po.C.sup._right | SurfaceArea | 0.76 | 0.15 | 0.78 | 0.14 | 0.41 | 0.2 | 0.93 | 0.15 | 0.76 |
| S.R.inf._right | Length | 0.31 | 0.15 | 0.62 | 0.16 | 0.8 | 0.19 | 0.88 | 0.16 | 0.64 |
| S.R.inf._right | MeanDepth | 0.52 | 0.16 | 0.91 | 0.13 | 0.88 | 0.18 | 0.92 | 0.15 | 0.82 |
| S.R.inf._right | Width | 0.68 | 0.16 | 0.5 | 0.17 | 0.66 | 0.21 | 0.44 | 0.19 | 0.57 |
| S.R.inf._right | SurfaceArea | 0.41 | 0.16 | 0.59 | 0.17 | 0.75 | 0.2 | 0.95 | 0.14 | 0.69 |
| S.Rh._right | Length | 0.45 | 0.16 | 0.48 | 0.16 | 0.68 | 0.21 | 0.58 | 0.19 | 0.53 |
| S.Rh._right | MeanDepth | 0.2 | 0.13 | 0.9 | 0.12 | 0.84 | 0.19 | 0.68 | 0.19 | 0.64 |
| S.Rh._right | Width | 0.17 | 0.13 | 0.75 | 0.15 | 0.77 | 0.2 | 0.77 | 0.19 | 0.54 |
| S.Rh._right | SurfaceArea | 0.31 | 0.15 | 0.82 | 0.14 | 0.88 | 0.18 | 0.5 | 0.2 | 0.63 |
| S.s.P._right | Length | 0.52 | 0.16 | 0.61 | 0.15 | 0.81 | 0.19 | 0.61 | 0.19 | 0.62 |
| S.s.P._right | MeanDepth | 0.83 | 0.13 | 0.24 | 0.14 | 0.73 | 0.2 | 0.78 | 0.18 | 0.62 |
| S.s.P._right | Width | 0.81 | 0.14 | 0.79 | 0.14 | 0.32 | 0.19 | 0.8 | 0.17 | 0.72 |
| S.s.P._right | SurfaceArea | 0.72 | 0.15 | 0.78 | 0.14 | 0.66 | 0.2 | 0.85 | 0.17 | 0.76 |
| S.T.i.ant._right | Length | 0.68 | 0.15 | 0.79 | 0.13 | 0.78 | 0.2 | 0.66 | 0.19 | 0.73 |
| S.T.i.ant._right | MeanDepth | 0.81 | 0.14 | 0.93 | 0.11 | 0.78 | 0.19 | 0.93 | 0.15 | 0.88 |
| S.T.i.ant._right | Width | 0.63 | 0.15 | 0.79 | 0.14 | 0.63 | 0.2 | 0.77 | 0.18 | 0.72 |
| S.T.i.ant._right | SurfaceArea | 0.78 | 0.14 | 0.89 | 0.12 | 0.82 | 0.18 | 0.74 | 0.18 | 0.82 |
| S.T.i.post._right | Length | 0.77 | 0.14 | 0.72 | 0.14 | 0.81 | 0.19 | 0.75 | 0.18 | 0.76 |
| S.T.i.post._right | MeanDepth | 0.66 | 0.15 | 0.83 | 0.13 | 0.79 | 0.19 | 0.63 | 0.19 | 0.74 |
| S.T.i.post._right | Width | 0.68 | 0.15 | 0.81 | 0.13 | 0.57 | 0.21 | 0.73 | 0.18 | 0.72 |
| S.T.i.post._right | SurfaceArea | 0.74 | 0.14 | 0.78 | 0.14 | 0.82 | 0.18 | 0.86 | 0.16 | 0.79 |
| S.T.pol._right | Length | 0.71 | 0.15 | 0.71 | 0.14 | 0.88 | 0.17 | 0.45 | 0.19 | 0.7 |
| S.T.pol._right | MeanDepth | 0.61 | 0.15 | 0.47 | 0.16 | 0.62 | 0.21 | 0.76 | 0.18 | 0.6 |
| S.T.pol._right | Width | 0.18 | 0.13 | 0.6 | 0.15 | 0.84 | 0.19 | 0.84 | 0.18 | 0.54 |
| S.T.pol._right | SurfaceArea | 0.75 | 0.14 | 0.7 | 0.15 | 0.93 | 0.16 | 0.64 | 0.19 | 0.76 |
| S.T.s._right | Length | 0.57 | 0.15 | 0.86 | 0.13 | 0.86 | 0.18 | 0.81 | 0.17 | 0.78 |
| S.T.s._right | MeanDepth | 0.79 | 0.14 | 0.89 | 0.12 | 0.71 | 0.2 | 0.82 | 0.18 | 0.82 |
| S.T.s._right | Width | 0.58 | 0.16 | 0.74 | 0.14 | 0.73 | 0.2 | 0.85 | 0.17 | 0.72 |
| S.T.s._right | SurfaceArea | 0.68 | 0.15 | 0.93 | 0.11 | 0.94 | 0.15 | 0.76 | 0.18 | 0.85 |
| S.T.s.ter.asc.ant._right | Length | 0.63 | 0.16 | 0.75 | 0.14 | 0.59 | 0.21 | 0.26 | 0.17 | 0.58 |
| S.T.s.ter.asc.ant._right | MeanDepth | 0.69 | 0.16 | 0.74 | 0.14 | 0.8 | 0.19 | 0.23 | 0.16 | 0.61 |
| S.T.s.ter.asc.ant._right | Width | 0.64 | 0.16 | 0.76 | 0.14 | 0.42 | 0.21 | 0.61 | 0.19 | 0.64 |
| S.T.s.ter.asc.ant._right | SurfaceArea | 0.66 | 0.16 | 0.93 | 0.11 | 0.64 | 0.21 | 0.26 | 0.17 | 0.7 |
| S.T.s.ter.asc.post._right | Length | 0.69 | 0.15 | 0.83 | 0.13 | 0.61 | 0.21 | 0.47 | 0.19 | 0.69 |
| S.T.s.ter.asc.post._right | MeanDepth | 0.81 | 0.14 | 0.75 | 0.14 | 0.78 | 0.19 | 0.61 | 0.19 | 0.75 |
| S.T.s.ter.asc.post._right | Width | 0.58 | 0.16 | 0.81 | 0.13 | 0.71 | 0.2 | 0.55 | 0.19 | 0.68 |
| S.T.s.ter.asc.post._right | SurfaceArea | 0.81 | 0.14 | 0.9 | 0.12 | 0.85 | 0.18 | 0.76 | 0.18 | 0.84 |

**Table S5:** ICC for the bilaterally average sulcal descriptors (yellow: ICC > 0.75; orange: ICC > 0.9)

| Sulci | Descriptor | QTIM | QTIM_SE | HCP | HCP_SE | OASIS | OASIS_SE | KKI | KKI_SE | Meta-SE-ICC |
|---|---|---|---|---|---|---|---|---|---|---|
| F.C.L.a. | Length | 0.18 | 0.13 | 0.38 | 0.15 | 0.25 | 0.18 | 0.46 | 0.2 | 0.29 |
| F.C.L.a. | MeanDepth | 0.3 | 0.15 | 0.14 | 0.11 | 0.5 | 0.22 | - | - | - |
| F.C.L.a. | Width | 0.41 | 0.16 | 0.58 | 0.15 | 0.88 | 0.18 | 0.85 | 0.17 | 0.66 |
| F.C.L.a. | SurfaceArea | 0.37 | 0.15 | 0.45 | 0.15 | 0.06 | 0.08 | 0.64 | 0.2 | 0.22 |
| F.C.L.p. | Length | 0.71 | 0.15 | 0.77 | 0.14 | 0.81 | 0.19 | 0.82 | 0.17 | 0.77 |
| F.C.L.p. | MeanDepth | 0.18 | 0.13 | 0.62 | 0.15 | 0.81 | 0.19 | 0.7 | 0.18 | 0.5 |
| F.C.L.p. | Width | 0.78 | 0.14 | 0.88 | 0.12 | 0.9 | 0.17 | 0.95 | 0.14 | 0.87 |
| F.C.L.p. | SurfaceArea | 0.75 | 0.15 | 0.85 | 0.13 | 0.92 | 0.16 | 0.89 | 0.16 | 0.85 |
| F.C.L.r.ant. | Length | 0.6 | 0.17 | 0.54 | 0.17 | 0.46 | 0.23 | 0.89 | 0.16 | 0.65 |
| F.C.L.r.ant. | MeanDepth | 0.45 | 0.17 | 0.82 | 0.15 | 0.8 | 0.21 | 0.73 | 0.19 | 0.7 |
| F.C.L.r.ant. | Width | 0.55 | 0.17 | 0.68 | 0.16 | 0.51 | 0.23 | 0.92 | 0.16 | 0.7 |
| F.C.L.r.ant. | SurfaceArea | 0.64 | 0.17 | 0.68 | 0.16 | 0.68 | 0.23 | 0.94 | 0.15 | 0.76 |
| F.C.L.r.asc. | Length | 0.62 | 0.16 | 0.81 | 0.14 | 0.93 | 0.16 | 0.81 | 0.18 | 0.79 |
| F.C.L.r.asc. | MeanDepth | 0.77 | 0.15 | 0.36 | 0.15 | 0.8 | 0.19 | 0.86 | 0.17 | 0.68 |
| F.C.L.r.asc. | Width | 0.66 | 0.15 | 0.87 | 0.13 | 0.79 | 0.19 | 0.91 | 0.16 | 0.81 |
| F.C.L.r.asc. | SurfaceArea | 0.73 | 0.15 | 0.74 | 0.15 | 0.84 | 0.18 | 0.9 | 0.16 | 0.79 |
| F.C.L.r.diag. | Length | - | - | - | - | - | - | - | - | - |
| F.C.L.r.diag. | MeanDepth | - | - | - | - | - | - | - | - | - |
| F.C.L.r.diag. | Width | - | - | - | - | - | - | - | - | - |
| F.C.L.r.diag. | SurfaceArea | - | - | - | - | - | - | - | - | - |
| F.C.L.r.retroC.tr. | Length | 0.81 | 0.16 | 0.84 | 0.15 | 0.67 | 0.22 | 0.81 | 0.21 | 0.8 |
| F.C.L.r.retroC.tr. | MeanDepth | 0.74 | 0.17 | 0.77 | 0.16 | 0.8 | 0.21 | 0.72 | 0.22 | 0.76 |
| F.C.L.r.retroC.tr. | Width | 0.77 | 0.17 | 0.73 | 0.17 | 0.79 | 0.21 | 0.8 | 0.21 | 0.77 |
| F.C.L.r.retroC.tr. | SurfaceArea | 0.83 | 0.16 | 0.79 | 0.16 | 0.92 | 0.18 | 0.85 | 0.21 | 0.85 |
| F.C.L.r.sc.ant. | Length | - | - | - | - | - | - | - | - | - |
| F.C.L.r.sc.ant. | MeanDepth | - | - | - | - | - | - | - | - | - |
| F.C.L.r.sc.ant. | Width | - | - | - | - | - | - | - | - | - |
| F.C.L.r.sc.ant. | SurfaceArea | - | - | - | - | - | - | - | - | - |
| F.C.L.r.sc.post. | Length | - | - | - | - | - | - | - | - | - |
| F.C.L.r.sc.post. | MeanDepth | 0.31 | 0.18 | - | - | 0.82 | 0.24 | 0.51 | 0.22 | 0.5 |
| F.C.L.r.sc.post. | Width | 0.71 | 0.19 | - | - | 0.83 | 0.23 | 0.93 | 0.17 | 0.83 |
| F.C.L.r.sc.post. | SurfaceArea | 0.88 | 0.16 | - | - | 0.93 | 0.21 | 0.37 | 0.21 | 0.75 |
| F.C.M.ant. | Length | 0.54 | 0.16 | 0.76 | 0.14 | 0.7 | 0.2 | 0.75 | 0.18 | 0.69 |
| F.C.M.ant. | MeanDepth | 0.7 | 0.16 | 0.83 | 0.13 | 0.62 | 0.21 | 0.87 | 0.17 | 0.78 |
| F.C.M.ant. | Width | 0.81 | 0.15 | 0.6 | 0.15 | 0.76 | 0.19 | 0.45 | 0.2 | 0.67 |
| F.C.M.ant. | SurfaceArea | 0.6 | 0.17 | 0.85 | 0.13 | 0.82 | 0.19 | 0.85 | 0.17 | 0.79 |
| F.C.M.post. | Length | 0.53 | 0.16 | 0.62 | 0.15 | 0.52 | 0.21 | 0.82 | 0.17 | 0.62 |
| F.C.M.post. | MeanDepth | 0.83 | 0.13 | 0.89 | 0.12 | 0.95 | 0.15 | 0.91 | 0.15 | 0.89 |
| F.C.M.post. | Width | 0.9 | 0.12 | 0.85 | 0.13 | 0.81 | 0.19 | 0.93 | 0.15 | 0.88 |
| F.C.M.post. | SurfaceArea | 0.69 | 0.15 | 0.76 | 0.14 | 0.8 | 0.19 | 0.91 | 0.16 | 0.78 |
| F.Cal.ant.-Sc.Cal. | Length | 0.67 | 0.15 | 0.77 | 0.14 | 0.61 | 0.2 | 0.53 | 0.2 | 0.67 |
| F.Cal.ant.-Sc.Cal. | MeanDepth | 0.64 | 0.15 | 0.92 | 0.12 | 0.92 | 0.16 | 0.67 | 0.19 | 0.81 |
| F.Cal.ant.-Sc.Cal. | Width | 0.56 | 0.16 | 0.58 | 0.15 | 0.73 | 0.2 | 0.92 | 0.15 | 0.7 |
| F.Cal.ant.-Sc.Cal. | SurfaceArea | 0.7 | 0.15 | 0.95 | 0.11 | 0.87 | 0.17 | 0.91 | 0.15 | 0.87 |
| F.Coll. | Length | 0.88 | 0.13 | 0.66 | 0.15 | 0.48 | 0.2 | 0.61 | 0.19 | 0.71 |
| F.Coll. | MeanDepth | 0.92 | 0.12 | 0.89 | 0.12 | 0.83 | 0.18 | 0.91 | 0.15 | 0.89 |
| F.Coll. | Width | 0.7 | 0.15 | 0.71 | 0.14 | 0.43 | 0.2 | 0.72 | 0.19 | 0.66 |
| F.Coll. | SurfaceArea | 0.91 | 0.12 | 0.82 | 0.13 | 0.68 | 0.2 | 0.79 | 0.18 | 0.83 |
| F.I.P. | Length | 0.86 | 0.13 | 0.79 | 0.14 | 0.85 | 0.18 | 0.71 | 0.18 | 0.81 |
| F.I.P. | MeanDepth | 0.85 | 0.13 | 0.85 | 0.13 | 0.81 | 0.19 | 0.93 | 0.15 | 0.86 |
| F.I.P. | Width | 0.93 | 0.12 | 0.97 | 0.1 | 0.93 | 0.16 | 0.92 | 0.15 | 0.95 |
| F.I.P. | SurfaceArea | 0.91 | 0.12 | 0.89 | 0.12 | 0.94 | 0.15 | 0.81 | 0.17 | 0.89 |
| F.I.P.Po.C.inf. | Length | 0.86 | 0.13 | 0.82 | 0.13 | 0.32 | 0.2 | 0.63 | 0.2 | 0.73 |
| F.I.P.Po.C.inf. | MeanDepth | 0.8 | 0.14 | 0.74 | 0.14 | 0.86 | 0.18 | 0.9 | 0.16 | 0.82 |
| F.I.P.Po.C.inf. | Width | 0.91 | 0.12 | 0.89 | 0.12 | 0.84 | 0.19 | 0.95 | 0.14 | 0.9 |
| F.I.P.Po.C.inf. | SurfaceArea | 0.83 | 0.14 | 0.89 | 0.12 | 0.66 | 0.21 | 0.71 | 0.19 | 0.81 |
| F.I.P.r.int.1 | Length | 0.81 | 0.15 | 0.73 | 0.15 | 0.82 | 0.21 | 0.7 | 0.19 | 0.76 |
| F.I.P.r.int.1 | MeanDepth | 0.74 | 0.16 | 0.74 | 0.15 | 0.84 | 0.2 | 0.33 | 0.18 | 0.67 |
| F.I.P.r.int.1 | Width | 0.76 | 0.16 | 0.78 | 0.14 | 0.78 | 0.22 | 0.8 | 0.18 | 0.78 |
| F.I.P.r.int.1 | SurfaceArea | 0.83 | 0.15 | 0.65 | 0.16 | 0.87 | 0.2 | 0.57 | 0.2 | 0.74 |
| F.I.P.r.int.2 | Length | 0.37 | 0.19 | - | - | 0.03 | 0.04 | 0.72 | 0.23 | 0.05 |

| | | | | | | | | | | |
|---|---|---|---|---|---|---|---|---|---|---|
| F.I.P.r.int.2 | MeanDepth | 0.67 | 0.19 | 0.92 | 0.14 | 0.72 | 0.26 | 0.85 | 0.21 | 0.82 |
| F.I.P.r.int.2 | Width | 0.8 | 0.18 | 0.71 | 0.17 | 0.95 | 0.2 | 0.66 | 0.24 | 0.79 |
| F.I.P.r.int.2 | SurfaceArea | 0.21 | 0.16 | - | - | - | - | 0.8 | 0.22 | - |
| F.P.O. | Length | 0.61 | 0.15 | 0.73 | 0.14 | 0.75 | 0.19 | 0.53 | 0.19 | 0.66 |
| F.P.O. | MeanDepth | 0.55 | 0.16 | 0.88 | 0.13 | 0.82 | 0.18 | 0.69 | 0.18 | 0.75 |
| F.P.O. | Width | 0.89 | 0.13 | 0.91 | 0.12 | 0.86 | 0.18 | 0.89 | 0.16 | 0.89 |
| F.P.O. | SurfaceArea | 0.8 | 0.14 | 0.87 | 0.13 | 0.89 | 0.18 | 0.62 | 0.19 | 0.81 |
| INSULA | Length | - | - | - | - | - | - | - | - | - |
| INSULA | MeanDepth | 0.22 | 0.14 | 0.5 | 0.15 | 0.43 | 0.2 | 0.36 | 0.18 | 0.36 |
| INSULA | Width | 0.61 | 0.15 | 0.56 | 0.15 | 0.63 | 0.2 | 0.87 | 0.17 | 0.66 |
| INSULA | SurfaceArea | 0.6 | 0.15 | 0.77 | 0.14 | 0.64 | 0.2 | 0.82 | 0.17 | 0.71 |
| OCCIPITAL | Length | 0.85 | 0.13 | 0.92 | 0.12 | 0.74 | 0.19 | 0.85 | 0.17 | 0.86 |
| OCCIPITAL | MeanDepth | 0.85 | 0.13 | 0.81 | 0.13 | 0.85 | 0.18 | 0.86 | 0.16 | 0.84 |
| OCCIPITAL | Width | 0.57 | 0.16 | 0.92 | 0.11 | 0.89 | 0.17 | 0.93 | 0.15 | 0.84 |
| OCCIPITAL | SurfaceArea | 0.87 | 0.13 | 0.91 | 0.12 | 0.83 | 0.18 | 0.86 | 0.16 | 0.87 |
| S.C. | Length | 0.7 | 0.15 | 0.8 | 0.13 | 0.93 | 0.16 | 0.83 | 0.17 | 0.81 |
| S.C. | MeanDepth | 0.83 | 0.13 | 0.89 | 0.12 | 0.71 | 0.2 | 0.89 | 0.16 | 0.85 |
| S.C. | Width | 0.94 | 0.11 | 0.95 | 0.11 | 0.97 | 0.14 | 0.93 | 0.15 | 0.95 |
| S.C. | SurfaceArea | 0.84 | 0.13 | 0.93 | 0.11 | 0.98 | 0.13 | 0.91 | 0.15 | 0.92 |
| S.C.LPC. | Length | - | - | - | - | - | - | - | - | - |
| S.C.LPC. | MeanDepth | - | - | - | - | - | - | - | - | - |
| S.C.LPC. | Width | - | - | - | - | - | - | - | - | - |
| S.C.LPC. | SurfaceArea | - | - | - | - | - | - | - | - | - |
| S.C.sylvian. | Length | - | - | - | - | 0.92 | 0.21 | 0.82 | 0.2 | 0.86 |
| S.C.sylvian. | MeanDepth | - | - | - | - | 0.86 | 0.23 | 0.79 | 0.2 | 0.82 |
| S.C.sylvian. | Width | - | - | - | - | 0.95 | 0.19 | 0.91 | 0.18 | 0.93 |
| S.C.sylvian. | SurfaceArea | - | - | - | - | 0.95 | 0.2 | 0.6 | 0.22 | 0.8 |
| S.Call. | Length | 0.85 | 0.13 | 0.83 | 0.13 | 0.93 | 0.16 | 0.85 | 0.17 | 0.86 |
| S.Call. | MeanDepth | 0.85 | 0.13 | 0.88 | 0.12 | 0.71 | 0.2 | 0.79 | 0.17 | 0.83 |
| S.Call. | Width | 0.64 | 0.15 | 0.49 | 0.16 | 0.63 | 0.21 | 0.5 | 0.2 | 0.56 |
| S.Call. | SurfaceArea | 0.94 | 0.11 | 0.9 | 0.12 | 0.94 | 0.16 | 0.88 | 0.16 | 0.92 |
| S.Cu. | Length | 0.56 | 0.16 | 0.67 | 0.15 | 0.82 | 0.18 | 0.69 | 0.19 | 0.68 |
| S.Cu. | MeanDepth | 0.73 | 0.15 | 0.72 | 0.14 | 0.71 | 0.2 | 0.88 | 0.17 | 0.76 |
| S.Cu. | Width | 0.77 | 0.14 | 0.77 | 0.14 | 0.77 | 0.2 | 0.91 | 0.16 | 0.8 |
| S.Cu. | SurfaceArea | 0.5 | 0.16 | 0.73 | 0.14 | 0.87 | 0.17 | 0.73 | 0.19 | 0.7 |
| S.F.inf. | Length | 0.52 | 0.16 | 0.73 | 0.15 | 0.78 | 0.19 | 0.85 | 0.17 | 0.71 |
| S.F.inf. | MeanDepth | 0.89 | 0.13 | 0.96 | 0.1 | 0.9 | 0.17 | 0.92 | 0.15 | 0.92 |
| S.F.inf. | Width | 0.6 | 0.16 | 0.84 | 0.13 | 0.62 | 0.2 | 0.86 | 0.17 | 0.75 |
| S.F.inf. | SurfaceArea | 0.68 | 0.15 | 0.9 | 0.12 | 0.68 | 0.21 | 0.83 | 0.17 | 0.8 |
| S.F.inf.ant. | Length | 0.41 | 0.16 | 0.66 | 0.15 | 0.6 | 0.21 | 0.66 | 0.19 | 0.58 |
| S.F.inf.ant. | MeanDepth | 0.52 | 0.16 | 0.77 | 0.14 | 0.72 | 0.2 | 0.78 | 0.18 | 0.7 |
| S.F.inf.ant. | Width | 0.45 | 0.16 | 0.86 | 0.13 | 0.77 | 0.2 | 0.78 | 0.18 | 0.73 |
| S.F.inf.ant. | SurfaceArea | 0.62 | 0.16 | 0.71 | 0.14 | 0.6 | 0.21 | 0.61 | 0.19 | 0.65 |
| S.F.int. | Length | 0.78 | 0.14 | 0.9 | 0.12 | 0.89 | 0.17 | 0.91 | 0.16 | 0.87 |
| S.F.int. | MeanDepth | 0.78 | 0.14 | 0.9 | 0.12 | 0.81 | 0.19 | 0.89 | 0.16 | 0.85 |
| S.F.int. | Width | 0.65 | 0.15 | 0.66 | 0.15 | 0.74 | 0.19 | 0.86 | 0.17 | 0.72 |
| S.F.int. | SurfaceArea | 0.78 | 0.14 | 0.92 | 0.11 | 0.87 | 0.17 | 0.94 | 0.14 | 0.88 |
| S.F.inter. | Length | 0.53 | 0.16 | 0.87 | 0.12 | 0.77 | 0.19 | 0.8 | 0.17 | 0.76 |
| S.F.inter. | MeanDepth | 0.81 | 0.14 | 0.86 | 0.13 | 0.78 | 0.19 | 0.87 | 0.16 | 0.84 |
| S.F.inter. | Width | 0.78 | 0.14 | 0.79 | 0.14 | 0.72 | 0.2 | 0.65 | 0.19 | 0.75 |
| S.F.inter. | SurfaceArea | 0.67 | 0.15 | 0.92 | 0.12 | 0.81 | 0.19 | 0.86 | 0.16 | 0.83 |
| S.F.marginal. | Length | 0.48 | 0.16 | 0.8 | 0.14 | 0.63 | 0.21 | 0.83 | 0.17 | 0.7 |
| S.F.marginal. | MeanDepth | 0.71 | 0.15 | 0.67 | 0.15 | 0.61 | 0.21 | 0.71 | 0.19 | 0.68 |
| S.F.marginal. | Width | 0.49 | 0.16 | 0.64 | 0.15 | 0.15 | 0.14 | 0.74 | 0.18 | 0.47 |
| S.F.marginal. | SurfaceArea | 0.54 | 0.16 | 0.81 | 0.13 | 0.55 | 0.21 | 0.88 | 0.16 | 0.72 |
| S.F.median. | Length | 0.86 | 0.13 | 0.91 | 0.12 | 0.82 | 0.18 | 0.88 | 0.16 | 0.88 |
| S.F.median. | MeanDepth | 0.89 | 0.13 | 0.68 | 0.14 | 0.86 | 0.18 | 0.77 | 0.18 | 0.8 |
| S.F.median. | Width | 0.86 | 0.13 | 0.89 | 0.12 | 0.92 | 0.17 | 0.88 | 0.17 | 0.89 |
| S.F.median. | SurfaceArea | 0.88 | 0.13 | 0.91 | 0.12 | 0.85 | 0.18 | 0.89 | 0.16 | 0.89 |
| S.F.orbitaire. | Length | 0.48 | 0.17 | 0.48 | 0.17 | 0.77 | 0.22 | 0.66 | 0.2 | 0.57 |
| S.F.orbitaire. | MeanDepth | 0.32 | 0.16 | 0.85 | 0.14 | 0.97 | 0.16 | 0.84 | 0.18 | 0.75 |
| S.F.orbitaire. | Width | 0.31 | 0.16 | 0.78 | 0.15 | 0.79 | 0.21 | 0.85 | 0.18 | 0.66 |
| S.F.orbitaire. | SurfaceArea | 0.63 | 0.17 | 0.64 | 0.16 | 0.88 | 0.2 | 0.8 | 0.19 | 0.73 |

| | | | | | | | | | | |
|---|---|---|---|---|---|---|---|---|---|---|
| S.F.polaire.tr. | Length | 0.65 | 0.15 | 0.76 | 0.14 | 0.73 | 0.2 | 0.74 | 0.18 | 0.72 |
| S.F.polaire.tr. | MeanDepth | 0.68 | 0.15 | 0.58 | 0.15 | 0.8 | 0.19 | 0.79 | 0.17 | 0.7 |
| S.F.polaire.tr. | Width | 0.58 | 0.16 | 0.85 | 0.13 | 0.86 | 0.18 | 0.96 | 0.14 | 0.82 |
| S.F.polaire.tr. | SurfaceArea | 0.59 | 0.16 | 0.68 | 0.15 | 0.62 | 0.21 | 0.76 | 0.18 | 0.66 |
| S.F.sup. | Length | 0.64 | 0.16 | 0.71 | 0.14 | 0.81 | 0.19 | 0.88 | 0.16 | 0.75 |
| S.F.sup. | MeanDepth | 0.89 | 0.12 | 0.91 | 0.12 | 0.79 | 0.19 | 0.95 | 0.14 | 0.9 |
| S.F.sup. | Width | 0.77 | 0.14 | 0.94 | 0.11 | 0.7 | 0.2 | 0.85 | 0.17 | 0.85 |
| S.F.sup. | SurfaceArea | 0.68 | 0.15 | 0.87 | 0.12 | 0.83 | 0.18 | 0.88 | 0.16 | 0.82 |
| S.Li.ant. | Length | 0.57 | - | 0.63 | - | 0.79 | - | 0.65 | - | 0.65 |
| S.Li.ant. | MeanDepth | 0.81 | - | 0.96 | - | 0.68 | - | 0.76 | - | 0.85 |
| S.Li.ant. | Width | 0.8 | - | 0.75 | - | 0.72 | - | 0.88 | - | 0.79 |
| S.Li.ant. | SurfaceArea | 0.76 | - | 0.72 | - | 0.75 | - | 0.82 | - | 0.76 |
| S.Li.post. | Length | 0.75 | 0.17 | 0.52 | 0.17 | 0.78 | 0.21 | 0.61 | 0.2 | 0.66 |
| S.Li.post. | MeanDepth | 0.79 | 0.15 | 0.62 | 0.12 | 0.77 | 0.22 | 0.85 | 0.19 | 0.75 |
| S.Li.post. | Width | 0.68 | 0.16 | 0.84 | 0.16 | 0.85 | 0.22 | 0.91 | 0.17 | 0.82 |
| S.Li.post. | SurfaceArea | 0.83 | 0.16 | 0.58 | 0.17 | 0.81 | 0.21 | 0.76 | 0.18 | 0.74 |
| S.O.p. | Length | 0.62 | 0.14 | - | 0.15 | 0.81 | 0.2 | 0.48 | 0.17 | 0.64 |
| S.O.p. | MeanDepth | 0.23 | 0.12 | - | 0.13 | 0.89 | 0.18 | 0.92 | 0.15 | 0.62 |
| S.O.p. | Width | 0.11 | 0.15 | - | 0.14 | 0.68 | 0.2 | 0.9 | 0.18 | 0.37 |
| S.O.p. | SurfaceArea | 0.66 | 0.13 | - | 0.14 | 0.88 | 0.2 | 0.61 | 0.17 | 0.72 |
| S.O.T.lat.ant. | Length | 0.83 | 0.15 | 0.91 | 0.16 | 0.82 | 0.19 | 0.89 | 0.19 | 0.87 |
| S.O.T.lat.ant. | MeanDepth | 0.91 | 0.14 | 0.91 | 0.15 | 0.85 | 0.19 | 0.94 | 0.17 | 0.91 |
| S.O.T.lat.ant. | Width | 0.7 | 0.16 | 0.71 | 0.14 | 0.66 | 0.18 | 0.9 | 0.15 | 0.75 |
| S.O.T.lat.ant. | SurfaceArea | 0.94 | 0.14 | 0.96 | 0.16 | 0.92 | 0.19 | 0.96 | 0.18 | 0.95 |
| S.O.T.lat.int. | Length | 0.63 | 0.14 | - | 0.12 | 0.59 | 0.18 | 0.87 | 0.16 | 0.71 |
| S.O.T.lat.int. | MeanDepth | 0.68 | 0.12 | - | 0.12 | 0.67 | 0.18 | 0.7 | 0.14 | 0.68 |
| S.O.T.lat.int. | Width | 0.73 | 0.15 | - | 0.15 | 0.79 | 0.21 | 0.48 | 0.16 | 0.67 |
| S.O.T.lat.int. | SurfaceArea | 0.73 | 0.11 | - | 0.11 | 0.69 | 0.16 | 0.93 | 0.14 | 0.81 |
| S.O.T.lat.med. | Length | 0.27 | 0.19 | 0.86 | - | 0.09 | 0.22 | 0.42 | 0.18 | 0.35 |
| S.O.T.lat.med. | MeanDepth | 0.67 | 0.19 | 0.88 | - | 0.63 | 0.22 | 0.89 | 0.2 | 0.8 |
| S.O.T.lat.med. | Width | 0.31 | 0.19 | 0.86 | - | 0.64 | 0.21 | 0.83 | 0.21 | 0.67 |
| S.O.T.lat.med. | SurfaceArea | 0.31 | 0.19 | 0.75 | - | 0.16 | 0.22 | 0.61 | 0.16 | 0.45 |
| S.O.T.lat.post. | Length | 0.81 | 0.16 | 0.7 | 0.14 | 0.47 | 0.1 | 0.79 | 0.2 | 0.72 |
| S.O.T.lat.post. | MeanDepth | 0.91 | 0.17 | 0.83 | 0.13 | 0.84 | 0.24 | 0.91 | 0.17 | 0.88 |
| S.O.T.lat.post. | Width | 0.72 | 0.17 | 0.79 | 0.14 | 0.65 | 0.24 | 0.8 | 0.18 | 0.75 |
| S.O.T.lat.post. | SurfaceArea | 0.83 | 0.17 | 0.81 | 0.15 | 0.59 | 0.15 | 0.85 | 0.2 | 0.79 |
| S.Olf. | Length | 0.58 | 0.18 | 0.82 | - | 0.47 | 0.22 | 0.87 | 0.22 | 0.72 |
| S.Olf. | MeanDepth | 0.77 | 0.16 | 0.91 | - | 0.92 | 0.2 | 0.9 | 0.18 | 0.88 |
| S.Olf. | Width | 0.75 | 0.11 | 0.61 | - | 0.86 | 0.23 | 0.69 | 0.18 | 0.72 |
| S.Olf. | SurfaceArea | 0.79 | 0.18 | 0.94 | - | 0.84 | 0.2 | 0.95 | 0.22 | 0.89 |
| S.Or. | Length | 0.57 | 0.16 | 0.89 | 0.13 | 0.75 | 0.21 | 0.9 | 0.16 | 0.8 |
| S.Or. | MeanDepth | 0.74 | 0.14 | 0.36 | 0.12 | 0.94 | 0.16 | 0.87 | 0.15 | 0.72 |
| S.Or. | Width | 0.73 | 0.15 | 0.91 | 0.15 | 0.75 | 0.18 | 0.87 | 0.19 | 0.83 |
| S.Or. | SurfaceArea | 0.77 | 0.14 | 0.85 | 0.11 | 0.91 | 0.19 | 0.93 | 0.14 | 0.86 |
| S.p.C. | Length | 0.7 | 0.15 | 0.81 | 0.13 | 0.94 | 0.19 | 0.81 | 0.19 | 0.81 |
| S.p.C. | MeanDepth | 0.71 | 0.15 | 0.91 | 0.15 | 0.81 | 0.2 | 0.82 | 0.19 | 0.82 |
| S.p.C. | Width | 0.75 | 0.16 | 0.94 | 0.14 | 0.76 | 0.17 | 0.88 | 0.19 | 0.85 |
| S.p.C. | SurfaceArea | 0.72 | 0.15 | 0.93 | 0.13 | 0.9 | 0.18 | 0.82 | 0.18 | 0.85 |
| S.Pa.int. | Length | 0.52 | 0.16 | 0.8 | 0.12 | 0.81 | 0.2 | 0.72 | 0.15 | 0.71 |
| S.Pa.int. | MeanDepth | 0.66 | 0.15 | 0.92 | 0.15 | 0.93 | 0.15 | 0.92 | 0.16 | 0.86 |
| S.Pa.int. | Width | 0.68 | 0.15 | 0.87 | 0.12 | 0.86 | 0.19 | 0.91 | 0.16 | 0.83 |
| S.Pa.int. | SurfaceArea | 0.5 | 0.14 | 0.86 | 0.13 | 0.83 | 0.16 | 0.77 | 0.15 | 0.74 |
| S.Pa.sup. | Length | 0.6 | 0.16 | 0.68 | 0.14 | 0.76 | 0.19 | 0.67 | 0.18 | 0.67 |
| S.Pa.sup. | MeanDepth | 0.82 | 0.15 | 0.66 | 0.12 | 0.79 | 0.16 | 0.92 | 0.15 | 0.79 |
| S.Pa.sup. | Width | 0.63 | 0.15 | 0.89 | 0.13 | 0.89 | 0.18 | 0.86 | 0.16 | 0.83 |
| S.Pa.sup. | SurfaceArea | 0.6 | 0.16 | - | 0.13 | 0.81 | 0.18 | 0.93 | 0.18 | 0.79 |
| S.Pa.t. | Length | - | 0.2 | - | 0.18 | 0.89 | 0.23 | 0.67 | 0.23 | 0.78 |
| S.Pa.t. | MeanDepth | 0.87 | 0.18 | - | 0.18 | 0.87 | 0.22 | 0.84 | 0.19 | 0.86 |
| S.Pa.t. | Width | 0.88 | 0.2 | - | 0.15 | 0.59 | 0.2 | 0.6 | 0.2 | 0.73 |
| S.Pa.t. | SurfaceArea | - | 0.2 | - | - | 0.9 | 0.22 | 0.85 | 0.18 | 0.87 |
| S.Pe.C.inf. | Length | 0.78 | - | 0.81 | - | 0.57 | 0.22 | 0.91 | 0.22 | 0.79 |
| S.Pe.C.inf. | MeanDepth | 0.63 | 0.17 | 0.81 | - | 0.86 | 0.22 | 0.77 | 0.19 | 0.76 |
| S.Pe.C.inf. | Width | 0.29 | 0.17 | 0.84 | - | 0.69 | 0.25 | 0.92 | 0.22 | 0.67 |

| | | | | | | | | | | |
|---|---|---|---|---|---|---|---|---|---|---|
| S.Pe.C.inf. | SurfaceArea | 0.73 | - | 0.76 | - | 0.72 | 0.21 | 0.8 | 0.19 | 0.75 |
| S.Pe.C.inter. | Length | 0.64 | 0.15 | 0.85 | 0.15 | 0.69 | 0.23 | 0.72 | 0.18 | 0.74 |
| S.Pe.C.inter. | MeanDepth | 0.68 | 0.16 | 0.6 | 0.15 | 0.81 | 0.2 | 0.82 | 0.21 | 0.72 |
| S.Pe.C.inter. | Width | 0.82 | 0.16 | 0.85 | 0.15 | 0.85 | 0.23 | 0.86 | 0.18 | 0.84 |
| S.Pe.C.inter. | SurfaceArea | 0.6 | 0.16 | 0.85 | 0.16 | 0.61 | 0.22 | 0.65 | 0.2 | 0.7 |
| S.Pe.C.marginal. | Length | 0.64 | 0.16 | 0.63 | 0.13 | - | 0.2 | 0.8 | 0.18 | 0.68 |
| S.Pe.C.marginal. | MeanDepth | 0.68 | 0.15 | 0.8 | 0.15 | - | 0.19 | 0.82 | 0.17 | 0.76 |
| S.Pe.C.marginal. | Width | 0.76 | 0.14 | 0.85 | 0.13 | - | 0.18 | 0.82 | 0.17 | 0.81 |
| S.Pe.C.marginal. | SurfaceArea | 0.7 | 0.16 | 0.72 | 0.13 | - | 0.2 | 0.79 | 0.19 | 0.73 |
| S.Pe.C.median. | Length | 0.74 | 0.18 | 0.75 | 0.17 | 0.45 | - | 0.49 | 0.21 | 0.64 |
| S.Pe.C.median. | MeanDepth | 0.64 | 0.17 | 0.87 | 0.16 | 0.9 | - | 0.84 | 0.2 | 0.81 |
| S.Pe.C.median. | Width | 0.85 | 0.17 | 0.91 | 0.15 | 0.77 | - | 0.86 | 0.2 | 0.86 |
| S.Pe.C.median. | SurfaceArea | 0.67 | 0.18 | 0.84 | 0.16 | 0.4 | - | 0.38 | 0.21 | 0.63 |
| S.Pe.C.sup. | Length | 0.79 | 0.16 | 0.7 | 0.16 | 0.93 | 0.21 | 0.79 | 0.21 | 0.79 |
| S.Pe.C.sup. | MeanDepth | 0.85 | 0.17 | 0.73 | 0.14 | 0.92 | 0.17 | 0.76 | 0.19 | 0.81 |
| S.Pe.C.sup. | Width | 0.74 | 0.15 | 0.88 | 0.13 | 0.87 | 0.2 | 0.73 | 0.18 | 0.82 |
| S.Pe.C.sup. | SurfaceArea | 0.89 | 0.17 | 0.76 | 0.14 | 0.87 | 0.2 | 0.83 | 0.2 | 0.83 |
| S.Po.C.sup. | Length | 0.77 | 0.17 | 0.75 | 0.15 | 0.49 | 0.2 | 0.71 | 0.19 | 0.71 |
| S.Po.C.sup. | MeanDepth | 0.94 | 0.16 | 0.87 | 0.15 | 0.74 | 0.2 | 0.74 | 0.19 | 0.86 |
| S.Po.C.sup. | Width | 0.91 | 0.17 | 0.92 | 0.13 | 0.86 | 0.22 | 0.92 | 0.2 | 0.91 |
| S.Po.C.sup. | SurfaceArea | 0.85 | 0.15 | 0.77 | 0.14 | 0.47 | 0.22 | 0.79 | 0.18 | 0.76 |
| S.R.inf. | Length | 0.63 | 0.15 | - | 0.15 | 0.95 | 0.21 | 0.9 | 0.19 | 0.82 |
| S.R.inf. | MeanDepth | 0.8 | 0.12 | - | 0.13 | 0.93 | 0.2 | 0.94 | 0.18 | 0.89 |
| S.R.inf. | Width | 0.65 | 0.13 | - | 0.12 | 0.66 | 0.18 | 0.57 | 0.15 | 0.63 |
| S.R.inf. | SurfaceArea | 0.8 | 0.14 | - | 0.14 | 0.94 | 0.21 | 0.95 | 0.18 | 0.89 |
| S.Rh. | Length | 0.7 | 0.17 | 0.71 | - | 0.52 | 0.19 | 0.6 | 0.16 | 0.66 |
| S.Rh. | MeanDepth | 0.5 | 0.15 | 0.94 | - | 0.76 | 0.2 | 0.6 | 0.15 | 0.76 |
| S.Rh. | Width | 0.33 | 0.17 | 0.56 | - | 0.79 | 0.26 | 0.75 | 0.2 | 0.57 |
| S.Rh. | SurfaceArea | 0.71 | 0.15 | 0.84 | - | 0.5 | 0.2 | 0.72 | 0.15 | 0.73 |
| S.s.P. | Length | 0.76 | 0.17 | 0.76 | 0.16 | 0.83 | 0.17 | 0.79 | 0.19 | 0.78 |
| S.s.P. | MeanDepth | 0.88 | 0.16 | 0.71 | 0.14 | 0.77 | 0.2 | 0.94 | 0.19 | 0.83 |
| S.s.P. | Width | 0.9 | 0.16 | 0.86 | 0.14 | 0.58 | 0.2 | 0.75 | 0.18 | 0.82 |
| S.s.P. | SurfaceArea | 0.91 | 0.17 | 0.76 | 0.14 | 0.82 | 0.18 | 0.93 | 0.19 | 0.86 |
| S.T.i.ant. | Length | 0.63 | 0.16 | 0.89 | 0.15 | 0.85 | 0.23 | 0.78 | 0.2 | 0.8 |
| S.T.i.ant. | MeanDepth | 0.72 | 0.17 | 0.96 | 0.12 | 0.85 | 0.22 | 0.92 | 0.2 | 0.89 |
| S.T.i.ant. | Width | 0.66 | 0.16 | 0.82 | 0.16 | 0.55 | 0.21 | 0.81 | 0.2 | 0.73 |
| S.T.i.ant. | SurfaceArea | 0.73 | 0.16 | 0.95 | 0.14 | 0.86 | 0.23 | 0.86 | 0.19 | 0.87 |
| S.T.i.post. | Length | 0.77 | 0.15 | 0.85 | 0.12 | 0.59 | 0.18 | 0.71 | 0.18 | 0.76 |
| S.T.i.post. | MeanDepth | 0.74 | 0.15 | 0.69 | 0.1 | 0.71 | 0.18 | 0.79 | 0.15 | 0.73 |
| S.T.i.post. | Width | 0.58 | 0.15 | 0.83 | 0.13 | 0.15 | 0.21 | 0.83 | 0.18 | 0.58 |
| S.T.i.post. | SurfaceArea | 0.74 | 0.15 | 0.82 | 0.11 | 0.45 | 0.18 | 0.77 | 0.16 | 0.73 |
| S.T.pol. | Length | 0.66 | 0.14 | 0.79 | 0.13 | 0.88 | 0.2 | 0.41 | 0.18 | 0.7 |
| S.T.pol. | MeanDepth | 0.61 | 0.14 | 0.69 | 0.14 | 0.89 | 0.2 | 0.91 | 0.17 | 0.76 |
| S.T.pol. | Width | 0.21 | 0.15 | 0.65 | 0.13 | 0.78 | 0.14 | 0.83 | 0.17 | 0.56 |
| S.T.pol. | SurfaceArea | 0.68 | 0.14 | 0.8 | 0.13 | 0.95 | 0.2 | 0.55 | 0.18 | 0.77 |
| S.T.s. | Length | 0.73 | 0.15 | 0.9 | 0.14 | 0.9 | 0.18 | 0.87 | 0.19 | 0.85 |
| S.T.s. | MeanDepth | 0.83 | 0.15 | 0.94 | 0.15 | 0.87 | 0.18 | 0.84 | 0.15 | 0.88 |
| S.T.s. | Width | 0.63 | 0.14 | 0.76 | 0.15 | 0.66 | 0.2 | 0.93 | 0.17 | 0.76 |
| S.T.s. | SurfaceArea | 0.72 | 0.15 | 0.91 | 0.13 | 0.86 | 0.16 | 0.91 | 0.2 | 0.86 |
| S.T.s.ter.asc.ant. | Length | 0.75 | 0.14 | 0.7 | 0.12 | 0.58 | 0.17 | 0.55 | 0.17 | 0.67 |
| S.T.s.ter.asc.ant. | MeanDepth | 0.82 | 0.13 | 0.89 | 0.11 | 0.9 | 0.17 | 0.69 | 0.17 | 0.84 |
| S.T.s.ter.asc.ant. | Width | 0.63 | 0.15 | 0.72 | 0.14 | 0.76 | 0.2 | 0.41 | 0.15 | 0.64 |
| S.T.s.ter.asc.ant. | SurfaceArea | 0.83 | 0.15 | 0.82 | 0.12 | 0.66 | 0.18 | 0.46 | 0.15 | 0.74 |
| S.T.s.ter.asc.post. | Length | 0.67 | 0.15 | 0.82 | 0.15 | 0.81 | 0.22 | 0.63 | 0.19 | 0.74 |
| S.T.s.ter.asc.post. | MeanDepth | 0.63 | 0.14 | 0.64 | 0.12 | 0.58 | 0.19 | 0.57 | 0.18 | 0.61 |
| S.T.s.ter.asc.post. | Width | 0.48 | 0.16 | 0.82 | 0.14 | 0.9 | 0.21 | 0.73 | 0.19 | 0.74 |
| S.T.s.ter.asc.post. | SurfaceArea | 0.67 | 0.14 | 0.86 | 0.14 | 0.84 | 0.22 | 0.7 | 0.19 | 0.78 |

**Table S6:** Bland-Altman analysis: estimation of **bias b** (mean difference between test and retest) in % for left, right and averaged sulcal shape descriptors. **Mean (SD)** of b across cohorts is reported. b estimates the differences between scan and rescan. While the ICC estimates the relation between within-subject variance and between-subjects variance, b represent a subject-based index. If scan and rescan are perfectly reliable, b should be equal to zero. Here we highlight those cases showing b greater than 0.1 as in 2. Sulcal-based values of b are reported in Supplementary Table_S7-S8 separately for each cohort and sulcal descriptor.

| **bias (*b*)** | **Length** | | | **Mean Depth** | | | **Width** | | | **Surface Area** | | |
|---|---|---|---|---|---|---|---|---|---|---|---|---|
| | left | right | avg | left | right | avg | left | right | avg | left | right | avg |
| **QTIM** | 5e-04 (0.013) | 7e-04 (0.007) | 0.003 (0.11) | -6e-04 (0.004) | -7e-04 (0.004) | -1e-03 (0.03) | -2e-05 (2e-04) | -1e-05 (1e-04) | -0.011 (0.06) | -9e-03 (0.02) | -7e-04 (0.02) | -0.017 (0.04) |
| **HCP** | 2e-04 (0.007) | 1e-02 (0.125) | 0.003 (0.04) | 6e-05 (0.002) | -2e-04 (0.002) | 2e-04 (0.02) | 2e-05 (1e-04) | -6e-06 (9e-05) | 0.005 (0.03) | -6e-03 (0.02) | -7e-03 (0.02) | -0.009 (0.03) |
| **OASIS** | -1e-03 (0.008) | -8e-03 (0.066) | -0.006 (0.06) | -5e-04 (0.004) | -1e-04 (0.004) | -2e-03 (0.03) | -3e-05 (2e-04) | -1e-05 (2e-04) | -0.012 (0.07) | 5e-04 (0.02) | 1e-03 (0.01) | 0.007 (0.04) |
| **KKI** | -6e-03 (0.071) | 1e-01 (0.960) | 0.043 (0.27) | -6e-04 (0.003) | -3e-04 (0.003) | -6e-03 (0.02) | -2e-05 (1e-04) | 2e-05 (1e-04) | 0.005 (0.10) | -5e-04 (0.03) | 9e-03 (0.03) | -0.005 (0.05) |

**Table S7:** bias (*b*) values for the left hemisphere sulcal descriptors
(red: abs(b) > 0.1 or NaN; orange: abs(b) > 1)

| Sulci | Descriptor | OASIS | KKI | QTIM | HCP |
|---|---|---|---|---|---|
| F.C.L.a. | Length | -7.20E-04 | -5.50E-04 | -1.90E-03 | 1.00E-03 |
| F.C.L.a. | MeanDepth | -3.80E-03 | -1.00E-02 | -3.70E-03 | -1.30E-03 |
| F.C.L.a. | Width | -4.20E-03 | 5.90E-03 | -1.10E-02 | 1.90E-02 |
| F.C.L.a. | SurfaceArea | -4.10E-06 | -4.30E-05 | -1.10E-04 | 5.10E-05 |
| F.C.L.p. | Length | -2.10E-04 | 2.50E-05 | -7.70E-05 | 6.40E-06 |
| F.C.L.p. | MeanDepth | 1.40E-03 | -1.20E-03 | 9.70E-04 | 8.00E-04 |
| F.C.L.p. | Width | -9.70E-03 | 7.50E-03 | -3.60E-02 | -3.10E-03 |
| F.C.L.p. | SurfaceArea | -7.20E-07 | 1.10E-06 | -4.70E-06 | -3.70E-06 |
| F.C.L.r.ant. | Length | 5.30E-04 | 1.40E-03 | 3.40E-03 | -1.20E-03 |
| F.C.L.r.ant. | MeanDepth | -9.40E-04 | -2.50E-03 | -3.00E-03 | -1.60E-03 |
| F.C.L.r.ant. | Width | 1.30E-02 | 1.90E-02 | -9.00E-03 | 2.50E-02 |
| F.C.L.r.ant. | SurfaceArea | -1.00E-05 | -4.00E-06 | 4.00E-05 | -8.80E-05 |
| F.C.L.r.asc. | Length | -8.50E-04 | 1.80E-04 | -2.70E-03 | -6.60E-04 |
| F.C.L.r.asc. | MeanDepth | -4.60E-03 | -3.40E-03 | 6.70E-04 | 7.20E-04 |
| F.C.L.r.asc. | Width | 1.40E-02 | 1.90E-02 | -4.20E-02 | -4.20E-03 |
| F.C.L.r.asc. | SurfaceArea | -7.10E-05 | -6.00E-05 | -9.80E-05 | -1.90E-05 |
| F.C.L.r.diag. | Length | 2.00E-03 | -1.40E-03 | -2.30E-03 | 1.10E-04 |
| F.C.L.r.diag. | MeanDepth | 6.30E-03 | 2.50E-04 | 3.20E-03 | 4.30E-04 |
| F.C.L.r.diag. | Width | -2.90E-02 | -9.70E-03 | 2.00E-03 | -2.40E-03 |
| F.C.L.r.diag. | SurfaceArea | 1.60E-04 | -5.30E-05 | 2.40E-05 | 1.70E-05 |
| F.C.L.r.retroC.tr. | Length | 9.00E-03 | -3.00E-03 | 2.80E-04 | 2.60E-03 |
| F.C.L.r.retroC.tr. | MeanDepth | -1.20E-02 | -3.30E-03 | 4.20E-03 | -2.10E-03 |
| F.C.L.r.retroC.tr. | Width | 3.00E-02 | -1.50E-02 | -5.50E-02 | -1.90E-02 |
| F.C.L.r.retroC.tr. | SurfaceArea | 9.00E-05 | -5.70E-05 | 8.60E-05 | -4.50E-05 |
| F.C.L.r.sc.ant. | Length | -1.00E-02 | 1.50E-01 | -3.30E-03 | - |
| F.C.L.r.sc.ant. | MeanDepth | 1.80E-03 | -4.90E-05 | 5.30E-05 | 0.00E+00 |
| F.C.L.r.sc.ant. | Width | 2.30E-03 | -2.40E-02 | -2.40E-03 | 0.00E+00 |
| F.C.L.r.sc.ant. | SurfaceArea | 3.60E-05 | 2.50E-04 | -1.00E-05 | 0.00E+00 |
| F.C.L.r.sc.post. | Length | -6.00E-02 | 3.70E-02 | -3.50E-02 | 3.40E-02 |
| F.C.L.r.sc.post. | MeanDepth | 1.30E-03 | -1.90E-03 | 3.20E-03 | 2.40E-03 |
| F.C.L.r.sc.post. | Width | -4.80E-02 | 1.30E-02 | 4.20E-02 | 6.90E-02 |
| F.C.L.r.sc.post. | SurfaceArea | 2.20E-04 | -6.10E-05 | 1.70E-04 | -8.10E-05 |
| F.C.M.ant. | Length | -1.60E-04 | -9.60E-04 | 1.50E-04 | 1.90E-04 |
| F.C.M.ant. | MeanDepth | 3.70E-03 | -6.00E-03 | 1.50E-03 | -1.30E-03 |
| F.C.M.ant. | Width | 4.40E-03 | -1.10E-02 | -3.20E-03 | 2.40E-02 |
| F.C.M.ant. | SurfaceArea | -2.00E-05 | -7.30E-05 | 2.30E-05 | 2.40E-05 |
| F.C.M.post. | Length | 3.50E-05 | 6.40E-04 | -5.40E-05 | -6.10E-04 |
| F.C.M.post. | MeanDepth | -4.90E-04 | -1.90E-03 | 1.80E-03 | 1.70E-03 |
| F.C.M.post. | Width | -1.20E-02 | -2.00E-02 | -1.80E-02 | -9.40E-03 |
| F.C.M.post. | SurfaceArea | 3.40E-06 | 2.20E-05 | 2.30E-06 | -2.30E-05 |
| F.Cal.ant.-Sc.Cal. | Length | -3.30E-04 | 2.40E-04 | 1.10E-04 | 5.00E-04 |
| F.Cal.ant.-Sc.Cal. | MeanDepth | 7.60E-04 | -1.30E-03 | 6.80E-04 | -2.50E-03 |
| F.Cal.ant.-Sc.Cal. | Width | -3.00E-02 | 2.90E-02 | -3.60E-02 | -1.30E-02 |
| F.Cal.ant.-Sc.Cal. | SurfaceArea | -5.30E-06 | -6.00E-06 | 7.20E-06 | 1.70E-05 |
| F.Coll. | Length | 8.10E-04 | 3.50E-04 | -3.00E-05 | -2.00E-04 |
| F.Coll. | MeanDepth | -4.20E-04 | -7.50E-04 | 2.40E-04 | -1.60E-04 |
| F.Coll. | Width | 1.20E-02 | 4.50E-02 | -2.30E-02 | -2.80E-02 |
| F.Coll. | SurfaceArea | 3.70E-05 | 2.10E-05 | -4.50E-06 | -1.10E-05 |
| F.I.P. | Length | -2.10E-04 | 2.90E-04 | 8.10E-06 | -5.80E-05 |
| F.I.P. | MeanDepth | 1.50E-03 | -6.90E-04 | 6.60E-04 | 2.40E-04 |
| F.I.P. | Width | -6.00E-03 | -5.70E-03 | -1.90E-02 | -6.50E-03 |
| F.I.P. | SurfaceArea | -6.40E-06 | 6.70E-06 | 3.80E-06 | -1.40E-06 |
| F.I.P.Po.C.inf. | Length | 1.60E-03 | 1.00E-03 | -1.80E-04 | 1.40E-04 |
| F.I.P.Po.C.inf. | MeanDepth | 1.60E-03 | -1.10E-03 | -2.70E-03 | 5.70E-04 |

| | | | | | |
|---|---|---|---|---|---|
| F.I.P.Po.C.inf. | Width | 5.50E-04 | 9.70E-04 | -7.40E-03 | -1.80E-03 |
| F.I.P.Po.C.inf. | SurfaceArea | 6.10E-05 | 3.10E-05 | -2.30E-05 | 1.50E-05 |
| F.I.P.r.int.1 | Length | 2.30E-03 | 1.10E-03 | 8.00E-04 | 5.50E-04 |
| F.I.P.r.int.1 | MeanDepth | 4.30E-03 | 3.70E-03 | 3.40E-03 | 4.50E-03 |
| F.I.P.r.int.1 | Width | -2.90E-02 | -3.70E-02 | -1.20E-02 | -3.40E-03 |
| F.I.P.r.int.1 | SurfaceArea | 1.40E-04 | 9.80E-05 | 3.80E-05 | 6.40E-05 |
| F.I.P.r.int.2 | Length | 3.40E-03 | -1.40E-03 | -8.60E-03 | -5.20E-04 |
| F.I.P.r.int.2 | MeanDepth | 2.50E-03 | -4.00E-03 | -1.20E-02 | -8.70E-04 |
| F.I.P.r.int.2 | Width | 1.70E-02 | -8.00E-03 | 1.70E-02 | 2.10E-02 |
| F.I.P.r.int.2 | SurfaceArea | 3.20E-04 | -2.20E-04 | -8.50E-04 | 1.20E-04 |
| F.P.O. | Length | 1.50E-04 | -1.90E-03 | -3.80E-04 | -9.40E-06 |
| F.P.O. | MeanDepth | -7.70E-04 | -5.00E-04 | -1.20E-03 | -5.10E-04 |
| F.P.O. | Width | -1.10E-02 | -3.00E-02 | -1.70E-02 | 6.60E-03 |
| F.P.O. | SurfaceArea | 4.00E-06 | -4.00E-05 | -1.60E-05 | 1.10E-05 |
| INSULA | Length | 2.90E-03 | -5.30E-01 | 9.40E-02 | -4.30E-02 |
| INSULA | MeanDepth | 1.60E-03 | -2.00E-03 | -8.40E-04 | 6.50E-04 |
| INSULA | Width | 2.60E-03 | -5.10E-03 | 9.50E-02 | -1.40E-02 |
| INSULA | SurfaceArea | -4.50E-06 | -3.70E-07 | 1.10E-05 | 1.10E-06 |
| OCCIPITAL | Length | 3.00E-04 | -3.80E-06 | 7.30E-05 | -1.10E-04 |
| OCCIPITAL | MeanDepth | -1.80E-03 | -7.30E-04 | 6.60E-04 | 1.10E-03 |
| OCCIPITAL | Width | 1.70E-02 | 1.10E-02 | -2.20E-02 | -2.00E-02 |
| OCCIPITAL | SurfaceArea | 1.20E-05 | -1.70E-06 | 1.10E-05 | -6.20E-06 |
| S.C. | Length | 1.10E-04 | 1.80E-04 | -2.90E-04 | 5.90E-05 |
| S.C. | MeanDepth | -6.50E-05 | -2.60E-04 | 3.00E-04 | 3.20E-04 |
| S.C. | Width | 1.10E-03 | -6.20E-03 | -1.40E-02 | -1.60E-03 |
| S.C. | SurfaceArea | 2.20E-08 | 6.00E-06 | -1.30E-05 | 4.60E-06 |
| S.C.LPC. | Length | -7.90E-03 | -1.20E-03 | -9.60E-04 | -1.60E-04 |
| S.C.LPC. | MeanDepth | -6.90E-03 | -1.20E-03 | -3.20E-03 | 6.90E-04 |
| S.C.LPC. | Width | -9.40E-03 | -6.50E-03 | 1.80E-03 | -2.00E-02 |
| S.C.LPC. | SurfaceArea | -8.20E-04 | -3.20E-04 | -1.20E-04 | 1.20E-05 |
| S.C.sylvian. | Length | -5.60E-03 | 1.80E-03 | -1.60E-03 | 2.30E-03 |
| S.C.sylvian. | MeanDepth | 1.10E-02 | -1.80E-03 | 2.00E-03 | 4.90E-03 |
| S.C.sylvian. | Width | -1.30E-03 | -2.10E-02 | -2.30E-02 | -3.90E-02 |
| S.C.sylvian. | SurfaceArea | -1.30E-04 | 2.50E-05 | -2.20E-04 | 4.60E-04 |
| S.Call. | Length | -1.30E-04 | 3.10E-04 | -1.40E-05 | -1.30E-03 |
| S.Call. | MeanDepth | -2.40E-03 | -1.20E-02 | -4.80E-03 | -1.30E-03 |
| S.Call. | Width | 1.20E-02 | -1.40E-02 | -2.20E-02 | 3.60E-02 |
| S.Call. | SurfaceArea | -5.80E-05 | -2.10E-05 | -5.20E-05 | -1.30E-04 |
| S.Cu. | Length | -1.30E-03 | -1.20E-03 | -1.50E-04 | 1.20E-03 |
| S.Cu. | MeanDepth | 4.70E-03 | 2.60E-03 | -5.70E-03 | -4.90E-03 |
| S.Cu. | Width | -3.60E-03 | 4.70E-02 | 7.20E-03 | -3.90E-02 |
| S.Cu. | SurfaceArea | -1.30E-04 | -5.10E-05 | -8.40E-05 | 5.40E-06 |
| S.F.inf. | Length | -7.00E-04 | -2.80E-04 | 1.80E-04 | -9.70E-04 |
| S.F.inf. | MeanDepth | -2.00E-03 | 2.10E-04 | -7.20E-05 | 1.70E-03 |
| S.F.inf. | Width | 5.90E-03 | -6.00E-03 | -2.50E-02 | 2.00E-03 |
| S.F.inf. | SurfaceArea | -3.80E-05 | -7.10E-06 | 1.10E-05 | -1.50E-05 |
| S.F.inf.ant. | Length | -3.70E-03 | -1.10E-03 | -7.30E-04 | 1.60E-03 |
| S.F.inf.ant. | MeanDepth | -6.50E-04 | 2.70E-03 | 1.10E-04 | -1.60E-03 |
| S.F.inf.ant. | Width | -7.50E-04 | -2.20E-02 | 3.30E-03 | -5.90E-03 |
| S.F.inf.ant. | SurfaceArea | -2.60E-04 | -4.90E-05 | -5.80E-05 | 6.60E-05 |
| S.F.int. | Length | -3.70E-05 | 1.10E-04 | -8.50E-05 | 1.90E-05 |
| S.F.int. | MeanDepth | 6.10E-04 | 4.40E-04 | -1.00E-03 | 1.50E-04 |
| S.F.int. | Width | 7.80E-04 | -5.30E-03 | -8.90E-03 | -1.40E-02 |
| S.F.int. | SurfaceArea | 6.80E-06 | 9.50E-06 | -7.00E-06 | 5.90E-06 |
| S.F.inter. | Length | 4.20E-04 | 3.20E-04 | 1.90E-04 | -7.40E-05 |
| S.F.inter. | MeanDepth | 6.70E-04 | 1.20E-03 | -5.10E-04 | -8.10E-04 |
| S.F.inter. | Width | 2.10E-03 | -1.80E-02 | 9.50E-04 | -2.00E-03 |

| | | | | | |
|---|---|---|---|---|---|
| S.F.inter. | SurfaceArea | 2.30E-05 | 1.10E-05 | 8.80E-06 | -2.50E-06 |
| S.F.marginal. | Length | -2.00E-03 | -5.20E-04 | -6.10E-04 | 6.70E-04 |
| S.F.marginal. | MeanDepth | -8.60E-03 | -1.40E-04 | 2.20E-04 | 1.50E-03 |
| S.F.marginal. | Width | 3.50E-02 | 3.30E-02 | -1.20E-02 | -2.10E-02 |
| S.F.marginal. | SurfaceArea | -2.00E-04 | -2.20E-05 | -4.90E-05 | 5.40E-05 |
| S.F.median. | Length | -1.20E-03 | -5.40E-04 | -5.70E-04 | 5.30E-04 |
| S.F.median. | MeanDepth | -9.60E-04 | 4.50E-03 | -3.60E-03 | 2.10E-03 |
| S.F.median. | Width | 6.00E-03 | -1.40E-02 | -2.40E-03 | 8.90E-03 |
| S.F.median. | SurfaceArea | -9.90E-05 | 2.60E-06 | -8.40E-05 | 5.70E-05 |
| S.F.orbitaire. | Length | -1.60E-03 | -5.10E-03 | 4.10E-03 | -1.70E-04 |
| S.F.orbitaire. | MeanDepth | 9.20E-03 | -2.70E-03 | 8.90E-03 | -2.20E-03 |
| S.F.orbitaire. | Width | -2.30E-02 | 1.10E-02 | -3.40E-02 | 5.30E-04 |
| S.F.orbitaire. | SurfaceArea | 1.10E-04 | -4.20E-04 | 3.90E-04 | -8.30E-05 |
| S.F.polaire.tr. | Length | -8.10E-04 | -4.30E-04 | -1.90E-03 | -7.90E-04 |
| S.F.polaire.tr. | MeanDepth | -2.20E-03 | -2.10E-03 | -4.90E-03 | -1.30E-03 |
| S.F.polaire.tr. | Width | 3.70E-03 | -1.50E-02 | -4.50E-03 | 2.60E-03 |
| S.F.polaire.tr. | SurfaceArea | -9.20E-05 | -4.10E-05 | -1.80E-04 | -5.20E-05 |
| S.F.sup. | Length | 2.80E-04 | -2.00E-04 | 1.10E-04 | -1.20E-04 |
| S.F.sup. | MeanDepth | -2.40E-03 | -1.10E-03 | 6.90E-04 | -9.00E-04 |
| S.F.sup. | Width | 1.20E-02 | -5.30E-03 | -8.60E-03 | -5.20E-03 |
| S.F.sup. | SurfaceArea | 2.30E-06 | -9.60E-06 | 2.70E-06 | -7.70E-06 |
| S.GSM. | Length | 1.80E-03 | -5.70E-04 | -2.70E-03 | 1.20E-03 |
| S.GSM. | MeanDepth | 4.20E-03 | -1.40E-03 | -1.30E-03 | 1.40E-03 |
| S.GSM. | Width | 7.60E-03 | -9.60E-03 | -8.00E-03 | -1.60E-02 |
| S.GSM. | SurfaceArea | 1.90E-04 | -7.70E-05 | -2.50E-04 | 6.10E-05 |
| S.Li.ant. | Length | 3.70E-03 | 1.50E-03 | 1.10E-03 | 9.20E-04 |
| S.Li.ant. | MeanDepth | -4.60E-03 | -2.00E-03 | 2.20E-03 | 1.40E-03 |
| S.Li.ant. | Width | -5.60E-02 | 7.70E-02 | -3.50E-02 | -1.10E-02 |
| S.Li.ant. | SurfaceArea | 1.30E-04 | 7.50E-05 | 6.80E-05 | 3.30E-05 |
| S.Li.post. | Length | 7.60E-04 | -3.80E-03 | -3.20E-04 | 3.30E-03 |
| S.Li.post. | MeanDepth | -6.10E-04 | 2.50E-03 | 8.20E-03 | 4.50E-03 |
| S.Li.post. | Width | 1.00E-02 | -3.60E-03 | -4.00E-02 | -1.20E-02 |
| S.Li.post. | SurfaceArea | 1.20E-05 | -2.50E-04 | 1.00E-04 | 2.70E-04 |
| S.O.p. | Length | -3.60E-03 | -2.30E-03 | -1.50E-04 | -1.00E-03 |
| S.O.p. | MeanDepth | -5.20E-03 | -3.50E-04 | 1.40E-02 | -2.00E-03 |
| S.O.p. | Width | 6.00E-02 | -3.70E-02 | -6.20E-02 | -1.20E-02 |
| S.O.p. | SurfaceArea | -1.30E-04 | -1.50E-04 | 3.20E-04 | -1.00E-04 |
| S.O.T.lat.ant. | Length | -1.10E-03 | -2.50E-05 | 1.70E-04 | 5.60E-04 |
| S.O.T.lat.ant. | MeanDepth | 5.70E-03 | 2.20E-03 | 5.00E-03 | 1.40E-03 |
| S.O.T.lat.ant. | Width | -2.60E-03 | 1.30E-03 | -5.20E-03 | -2.30E-02 |
| S.O.T.lat.ant. | SurfaceArea | -3.40E-05 | -4.30E-06 | 1.50E-05 | 3.30E-05 |
| S.O.T.lat.int. | Length | 2.10E-03 | 1.60E-03 | -7.30E-04 | 3.40E-03 |
| S.O.T.lat.int. | MeanDepth | -1.10E-02 | 5.50E-03 | -5.80E-03 | -1.30E-03 |
| S.O.T.lat.int. | Width | 2.30E-02 | -5.10E-02 | -1.00E-02 | -3.80E-02 |
| S.O.T.lat.int. | SurfaceArea | -6.30E-05 | 1.30E-04 | -3.60E-05 | 1.80E-04 |
| S.O.T.lat.med. | Length | -1.10E-03 | 2.70E-03 | 3.10E-03 | -4.10E-03 |
| S.O.T.lat.med. | MeanDepth | -4.10E-03 | -4.60E-04 | 3.20E-03 | -2.20E-03 |
| S.O.T.lat.med. | Width | -6.20E-03 | 2.40E-02 | -2.30E-02 | 3.90E-02 |
| S.O.T.lat.med. | SurfaceArea | -1.10E-04 | 1.00E-04 | 2.20E-04 | -2.20E-04 |
| S.O.T.lat.post. | Length | -1.40E-04 | 4.00E-04 | 4.70E-05 | 8.80E-04 |
| S.O.T.lat.post. | MeanDepth | 2.70E-03 | -8.00E-04 | 2.60E-03 | 9.40E-04 |
| S.O.T.lat.post. | Width | -1.20E-02 | 1.80E-02 | -2.70E-02 | -3.90E-02 |
| S.O.T.lat.post. | SurfaceArea | 2.60E-05 | 1.40E-05 | 1.20E-05 | 4.50E-05 |
| S.Olf. | Length | 6.30E-04 | -5.40E-04 | 3.90E-03 | 2.60E-04 |
| S.Olf. | MeanDepth | -6.90E-03 | 2.90E-04 | -7.70E-03 | 3.10E-04 |
| S.Olf. | Width | 1.20E-03 | -1.30E-02 | 5.70E-03 | -3.50E-02 |
| S.Olf. | SurfaceArea | -2.10E-05 | -4.70E-05 | 1.20E-04 | 4.10E-05 |

| | | | | | |
|---|---|---|---|---|---|
| S.Or. | Length | 2.80E-04 | 2.70E-04 | -3.10E-04 | -1.50E-04 |
| S.Or. | MeanDepth | 2.50E-03 | -1.30E-03 | -8.50E-03 | 1.50E-03 |
| S.Or. | Width | -2.10E-03 | -1.00E-02 | 2.30E-02 | 8.00E-03 |
| S.Or. | SurfaceArea | 2.60E-05 | 3.00E-06 | -5.20E-05 | -2.60E-06 |
| S.p.C. | Length | -4.00E-03 | 8.20E-04 | 3.00E-03 | 3.00E-04 |
| S.p.C. | MeanDepth | -8.10E-03 | 6.10E-03 | 3.50E-03 | -3.60E-03 |
| S.p.C. | Width | 1.80E-02 | -3.10E-03 | -2.60E-02 | 7.70E-03 |
| S.p.C. | SurfaceArea | -6.60E-04 | 1.60E-04 | 4.90E-04 | -1.40E-05 |
| S.Pa.int. | Length | -8.30E-04 | 5.20E-04 | -7.10E-05 | 7.00E-05 |
| S.Pa.int. | MeanDepth | 6.50E-04 | 4.50E-03 | 9.10E-04 | 9.30E-04 |
| S.Pa.int. | Width | -2.60E-03 | 1.50E-02 | -1.70E-02 | 1.60E-03 |
| S.Pa.int. | SurfaceArea | -4.00E-05 | 7.30E-05 | 2.50E-05 | 2.10E-05 |
| S.Pa.sup. | Length | -2.00E-03 | -2.40E-03 | 1.60E-03 | 4.80E-04 |
| S.Pa.sup. | MeanDepth | 1.20E-03 | -5.40E-03 | -1.00E-03 | 6.80E-04 |
| S.Pa.sup. | Width | -1.80E-03 | 2.60E-03 | -8.00E-03 | -8.40E-03 |
| S.Pa.sup. | SurfaceArea | -5.20E-05 | -1.40E-04 | 4.90E-05 | 4.00E-05 |
| S.Pa.t. | Length | 4.50E-03 | 1.90E-02 | -9.90E-03 | 6.70E-04 |
| S.Pa.t. | MeanDepth | -8.00E-04 | -1.40E-03 | 2.50E-03 | 7.60E-04 |
| S.Pa.t. | Width | 4.20E-03 | 1.30E-01 | -3.00E-03 | -7.00E-02 |
| S.Pa.t. | SurfaceArea | 1.70E-04 | 2.60E-04 | -1.70E-04 | 2.80E-05 |
| S.Pe.C.inf. | Length | 1.80E-03 | 1.60E-03 | -1.80E-03 | -1.80E-04 |
| S.Pe.C.inf. | MeanDepth | -1.10E-03 | 2.20E-03 | 2.90E-04 | 3.60E-04 |
| S.Pe.C.inf. | Width | -7.60E-03 | -2.30E-02 | -1.10E-03 | 1.20E-03 |
| S.Pe.C.inf. | SurfaceArea | 1.30E-05 | 2.10E-05 | -1.80E-05 | -1.30E-06 |
| S.Pe.C.inter. | Length | -1.10E-03 | 1.30E-03 | -6.00E-04 | -6.20E-04 |
| S.Pe.C.inter. | MeanDepth | 2.00E-04 | 4.20E-03 | -5.20E-03 | -1.00E-03 |
| S.Pe.C.inter. | Width | -2.70E-04 | -3.10E-02 | 4.30E-03 | -4.20E-03 |
| S.Pe.C.inter. | SurfaceArea | -2.30E-05 | 6.10E-05 | -6.00E-05 | -2.40E-05 |
| S.Pe.C.marginal. | Length | -9.10E-03 | 6.40E-03 | -2.70E-03 | 5.10E-03 |
| S.Pe.C.marginal. | MeanDepth | -2.90E-03 | 2.00E-03 | -5.10E-03 | 6.90E-04 |
| S.Pe.C.marginal. | Width | -2.00E-02 | -1.30E-02 | 6.20E-03 | -6.40E-03 |
| S.Pe.C.marginal. | SurfaceArea | -3.50E-04 | 3.00E-04 | -1.90E-04 | 1.70E-04 |
| S.Pe.C.median. | Length | -5.00E-03 | -2.90E-03 | -2.60E-03 | 7.40E-05 |
| S.Pe.C.median. | MeanDepth | 2.10E-03 | -1.00E-03 | -7.90E-04 | -1.20E-03 |
| S.Pe.C.median. | Width | 9.20E-03 | 7.70E-04 | -1.60E-02 | 2.50E-03 |
| S.Pe.C.median. | SurfaceArea | -1.90E-04 | -1.20E-04 | -5.70E-05 | -9.00E-06 |
| S.Pe.C.sup. | Length | 4.90E-03 | 9.00E-04 | -8.20E-04 | 8.50E-04 |
| S.Pe.C.sup. | MeanDepth | 2.10E-03 | 2.70E-03 | 1.30E-03 | 1.40E-03 |
| S.Pe.C.sup. | Width | 1.90E-02 | -2.10E-03 | -1.80E-02 | -7.10E-04 |
| S.Pe.C.sup. | SurfaceArea | 2.40E-04 | 5.20E-05 | -7.80E-06 | 4.40E-05 |
| S.Po.C.sup. | Length | 1.00E-03 | 4.40E-04 | -2.00E-04 | -3.10E-04 |
| S.Po.C.sup. | MeanDepth | -1.90E-03 | -5.30E-03 | -3.90E-03 | -7.80E-04 |
| S.Po.C.sup. | Width | -5.20E-03 | -1.40E-03 | -6.80E-03 | 5.00E-03 |
| S.Po.C.sup. | SurfaceArea | 6.00E-06 | -4.90E-05 | -8.80E-06 | -2.40E-05 |
| S.R.inf. | Length | -1.60E-03 | -1.60E-03 | -3.40E-04 | 1.80E-03 |
| S.R.inf. | MeanDepth | 2.20E-03 | 1.80E-03 | -8.50E-04 | 9.40E-04 |
| S.R.inf. | Width | 4.10E-03 | -3.00E-02 | 1.90E-03 | -6.30E-02 |
| S.R.inf. | SurfaceArea | -1.60E-04 | -1.50E-04 | 1.30E-06 | 2.90E-04 |
| S.Rh. | Length | 3.80E-03 | -2.00E-03 | -3.80E-03 | 1.60E-03 |
| S.Rh. | MeanDepth | -3.10E-03 | -1.20E-03 | -1.20E-02 | 2.60E-03 |
| S.Rh. | Width | -5.50E-04 | 2.50E-02 | 3.90E-02 | -8.90E-04 |
| S.Rh. | SurfaceArea | 1.60E-04 | -1.60E-04 | -5.10E-04 | 2.00E-04 |
| S.s.P. | Length | 4.70E-04 | -3.80E-04 | -7.60E-05 | 8.90E-04 |
| S.s.P. | MeanDepth | 1.00E-05 | 3.60E-04 | -1.30E-03 | -1.40E-03 |
| S.s.P. | Width | -1.30E-02 | -5.20E-02 | 9.80E-04 | -4.30E-03 |
| S.s.P. | SurfaceArea | 3.40E-05 | -1.40E-05 | -2.70E-05 | 5.70E-05 |
| S.T.i.ant. | Length | 7.00E-05 | -1.70E-04 | -4.60E-04 | -2.00E-05 |

| | | | | | |
|---|---|---|---|---|---|
| S.T.i.ant. | MeanDepth | 9.80E-04 | -8.50E-04 | -4.20E-03 | 4.80E-05 |
| S.T.i.ant. | Width | -3.30E-03 | 1.40E-02 | 4.40E-03 | -1.90E-02 |
| S.T.i.ant. | SurfaceArea | 6.20E-06 | -2.00E-05 | -5.70E-05 | 3.00E-06 |
| S.T.i.post. | Length | -5.80E-05 | 6.80E-05 | 3.90E-05 | -4.50E-04 |
| S.T.i.post. | MeanDepth | -1.90E-03 | 2.00E-03 | -3.00E-03 | 1.40E-03 |
| S.T.i.post. | Width | 3.00E-03 | -5.80E-03 | -2.20E-02 | 3.30E-03 |
| S.T.i.post. | SurfaceArea | -1.50E-05 | 2.10E-05 | -9.30E-06 | 6.20E-08 |
| S.T.pol. | Length | -3.80E-04 | 8.30E-04 | 1.90E-03 | -4.30E-04 |
| S.T.pol. | MeanDepth | 1.50E-03 | 3.70E-03 | -5.20E-03 | -1.00E-03 |
| S.T.pol. | Width | 1.70E-03 | 4.20E-03 | 8.30E-03 | 2.00E-03 |
| S.T.pol. | SurfaceArea | -3.00E-05 | 7.30E-05 | 1.20E-04 | -5.70E-05 |
| S.T.s. | Length | -4.40E-06 | -3.20E-05 | -6.70E-05 | -1.10E-04 |
| S.T.s. | MeanDepth | 1.70E-03 | 1.50E-04 | 1.20E-05 | -8.00E-04 |
| S.T.s. | Width | 3.10E-03 | 2.50E-04 | -1.50E-02 | -1.40E-02 |
| S.T.s. | SurfaceArea | 5.10E-06 | -6.10E-07 | -6.10E-06 | -4.60E-06 |
| S.T.s.ter.asc.ant. | Length | -7.10E-05 | -6.50E-03 | 2.10E-03 | 8.60E-04 |
| S.T.s.ter.asc.ant. | MeanDepth | -2.50E-04 | -5.30E-03 | -1.30E-03 | 9.30E-04 |
| S.T.s.ter.asc.ant. | Width | 1.50E-02 | -4.00E-02 | -1.40E-02 | -2.80E-02 |
| S.T.s.ter.asc.ant. | SurfaceArea | -3.30E-05 | -3.30E-04 | 5.60E-05 | 4.70E-05 |
| S.T.s.ter.asc.post. | Length | -1.50E-03 | 3.10E-03 | 3.00E-04 | 8.60E-05 |
| S.T.s.ter.asc.post. | MeanDepth | -6.70E-03 | -5.80E-04 | -1.40E-04 | -3.90E-03 |
| S.T.s.ter.asc.post. | Width | 8.90E-03 | 3.80E-02 | -2.40E-02 | 1.30E-02 |
| S.T.s.ter.asc.post. | SurfaceArea | -2.10E-05 | 1.10E-04 | 3.90E-05 | -2.90E-05 |

**Table S8:** bias (*b*) values for the right hemisphere sulcal descriptors (red: abs(*b*) > 0.1 or NaN; orange: abs(*b*) > 1)

| Sulci | Descriptor | OASIS | KKI | QTIM | HCP |
|---|---|---|---|---|---|
| F.C.L.a. | Length | -7.40E-04 | -7.50E-04 | -5.20E-04 | -5.00E-04 |
| F.C.L.a. | MeanDepth | -3.30E-04 | -3.50E-03 | 1.10E-03 | -4.50E-03 |
| F.C.L.a. | Width | -5.00E-05 | -3.60E-05 | -2.00E-05 | -1.80E-05 |
| F.C.L.a. | SurfaceArea | -1.40E-03 | 2.70E-03 | -2.90E-03 | 1.30E-02 |
| F.C.L.p. | Length | 9.80E-05 | 6.80E-05 | 5.90E-05 | -2.60E-05 |
| F.C.L.p. | MeanDepth | 1.70E-03 | -2.20E-03 | -8.30E-05 | 1.40E-03 |
| F.C.L.p. | Width | 4.20E-06 | -2.60E-06 | 3.80E-06 | 2.80E-07 |
| F.C.L.p. | SurfaceArea | -6.50E-03 | 6.70E-03 | 1.70E-03 | -2.10E-03 |
| F.C.L.r.ant. | Length | -3.70E-03 | -2.10E-03 | -3.10E-03 | 6.40E-04 |
| F.C.L.r.ant. | MeanDepth | 2.10E-03 | 2.50E-04 | 9.80E-04 | -4.20E-04 |
| F.C.L.r.ant. | Width | -3.90E-05 | 2.90E-05 | -8.70E-05 | 2.50E-05 |
| F.C.L.r.ant. | SurfaceArea | -2.70E-02 | 2.10E-02 | -1.60E-02 | -1.50E-02 |
| F.C.L.r.asc. | Length | 9.10E-04 | 6.20E-04 | -6.60E-04 | 3.90E-04 |
| F.C.L.r.asc. | MeanDepth | -9.90E-04 | 1.20E-03 | 9.20E-04 | -5.80E-03 |
| F.C.L.r.asc. | Width | 1.90E-05 | 1.40E-05 | 5.20E-06 | -1.10E-05 |
| F.C.L.r.asc. | SurfaceArea | -6.30E-04 | -8.40E-03 | -6.60E-03 | -1.40E-02 |
| F.C.L.r.diag. | Length | 2.40E-03 | 1.50E-03 | -8.50E-04 | -1.90E-03 |
| F.C.L.r.diag. | MeanDepth | 1.90E-03 | 4.60E-03 | -2.00E-03 | 1.80E-03 |
| F.C.L.r.diag. | Width | 4.50E-05 | 1.90E-04 | -2.60E-05 | -3.80E-05 |
| F.C.L.r.diag. | SurfaceArea | 1.10E-02 | -2.50E-02 | 1.40E-02 | -5.30E-03 |
| F.C.L.r.retroC.tr. | Length | 9.60E-04 | -2.40E-03 | -3.00E-03 | 1.30E-03 |
| F.C.L.r.retroC.tr. | MeanDepth | 5.50E-04 | -3.60E-03 | 3.30E-03 | -1.70E-03 |
| F.C.L.r.retroC.tr. | Width | -5.80E-06 | -1.30E-04 | -1.10E-04 | -4.30E-06 |
| F.C.L.r.retroC.tr. | SurfaceArea | -4.20E-02 | -2.30E-02 | 1.90E-02 | -1.30E-02 |
| F.C.L.r.sc.ant. | Length | - | - | - | -3.30E-01 |
| F.C.L.r.sc.ant. | MeanDepth | 6.80E-04 | 6.60E-04 | 1.00E-03 | 2.50E-04 |
| F.C.L.r.sc.ant. | Width | 2.10E-05 | -1.30E-04 | 4.30E-05 | -7.20E-06 |
| F.C.L.r.sc.ant. | SurfaceArea | -6.90E-03 | 2.00E-02 | 2.50E-02 | -1.10E-03 |
| F.C.L.r.sc.post. | Length | -5.10E-01 | 7.40E+00 | 0.00E+00 | 9.10E-01 |
| F.C.L.r.sc.post. | MeanDepth | -1.30E-03 | -2.10E-03 | 2.80E-04 | 2.60E-03 |
| F.C.L.r.sc.post. | Width | 3.30E-04 | 6.20E-04 | -3.30E-04 | -4.30E-06 |
| F.C.L.r.sc.post. | SurfaceArea | -1.30E-03 | 5.20E-03 | 2.60E-03 | -8.80E-04 |
| F.C.M.ant. | Length | -1.40E-04 | 3.90E-04 | 5.50E-04 | -9.10E-07 |
| F.C.M.ant. | MeanDepth | 2.60E-03 | -7.50E-04 | 1.30E-03 | 1.90E-03 |
| F.C.M.ant. | Width | 1.10E-06 | 1.40E-05 | 5.10E-05 | 8.70E-06 |
| F.C.M.ant. | SurfaceArea | 3.80E-04 | -3.60E-02 | -2.20E-02 | -1.20E-02 |
| F.C.M.post. | Length | 5.00E-04 | -5.10E-04 | 1.00E-04 | 2.00E-04 |
| F.C.M.post. | MeanDepth | -2.40E-03 | 9.90E-04 | -7.50E-04 | -1.10E-03 |
| F.C.M.post. | Width | 1.40E-05 | -3.00E-05 | 3.10E-06 | 4.30E-06 |
| F.C.M.post. | SurfaceArea | 1.00E-03 | -4.00E-03 | -9.20E-03 | -9.30E-03 |
| F.Cal.ant.-Sc.Cal. | Length | -4.30E-04 | -3.30E-05 | 5.00E-04 | 2.60E-04 |
| F.Cal.ant.-Sc.Cal. | MeanDepth | -4.70E-04 | -2.80E-04 | 1.80E-04 | -2.40E-04 |
| F.Cal.ant.-Sc.Cal. | Width | -3.40E-05 | -3.50E-06 | 2.10E-05 | 4.50E-06 |

| | | | | | |
|---|---|---|---|---|---|
| F.Cal.ant.-Sc.Cal. | SurfaceArea | -8.60E-03 | 3.80E-02 | -3.00E-02 | -2.20E-02 |
| F.Coll. | Length | 1.70E-04 | 3.50E-04 | -4.40E-04 | 2.30E-04 |
| F.Coll. | MeanDepth | 1.30E-03 | 8.60E-04 | -1.30E-03 | -9.00E-04 |
| F.Coll. | Width | 1.30E-05 | 1.40E-05 | -3.40E-05 | 1.20E-05 |
| F.Coll. | SurfaceArea | 3.70E-03 | 2.10E-02 | 4.60E-03 | 1.20E-02 |
| F.I.P. | Length | 1.60E-04 | -1.90E-04 | -7.50E-05 | -1.50E-04 |
| F.I.P. | MeanDepth | -4.60E-04 | -3.00E-04 | -2.30E-03 | 5.90E-04 |
| F.I.P. | Width | 3.40E-06 | -6.40E-06 | -3.70E-06 | -3.00E-06 |
| F.I.P. | SurfaceArea | 4.60E-03 | -1.20E-02 | -1.90E-03 | -2.70E-03 |
| F.I.P.Po.C.inf. | Length | 8.00E-04 | -2.10E-04 | 4.20E-04 | 3.60E-05 |
| F.I.P.Po.C.inf. | MeanDepth | -1.80E-04 | 9.60E-04 | -6.10E-04 | 9.60E-04 |
| F.I.P.Po.C.inf. | Width | 4.00E-05 | 2.50E-06 | 5.80E-06 | 4.30E-06 |
| F.I.P.Po.C.inf. | SurfaceArea | 1.70E-04 | -1.40E-02 | -6.10E-03 | -1.10E-02 |
| F.I.P.r.int.1 | Length | 8.70E-05 | -1.00E-03 | -9.60E-04 | 9.00E-05 |
| F.I.P.r.int.1 | MeanDepth | -5.30E-03 | -3.80E-03 | 2.90E-03 | -4.70E-03 |
| F.I.P.r.int.1 | Width | -9.20E-05 | -1.30E-04 | -8.00E-06 | -5.60E-05 |
| F.I.P.r.int.1 | SurfaceArea | -1.10E-02 | -2.00E-02 | 2.20E-02 | -3.00E-02 |
| F.I.P.r.int.2 | Length | 7.50E-03 | 4.40E-03 | 8.60E-04 | 6.90E-04 |
| F.I.P.r.int.2 | MeanDepth | 6.90E-03 | -2.50E-03 | -7.70E-03 | -5.40E-03 |
| F.I.P.r.int.2 | Width | 5.80E-04 | 2.20E-04 | -5.30E-05 | -4.50E-05 |
| F.I.P.r.int.2 | SurfaceArea | -5.70E-03 | 4.50E-02 | 4.60E-02 | 4.60E-02 |
| F.P.O. | Length | 7.50E-04 | -8.30E-04 | -4.30E-05 | -8.30E-04 |
| F.P.O. | MeanDepth | -1.90E-03 | -1.70E-04 | 2.60E-03 | -1.80E-04 |
| F.P.O. | Width | 1.10E-05 | 4.30E-06 | 7.40E-06 | -1.50E-05 |
| F.P.O. | SurfaceArea | 5.10E-03 | 6.40E-03 | -2.40E-02 | -5.00E-03 |
| INSULA | Length | 1.30E-02 | 2.80E-01 | 5.00E-02 | 1.70E-02 |
| INSULA | MeanDepth | 3.50E-04 | -7.80E-04 | -2.50E-04 | -1.10E-03 |
| INSULA | Width | 2.30E-06 | 1.50E-06 | 2.90E-06 | 6.60E-07 |
| INSULA | SurfaceArea | 2.90E-02 | 2.60E-02 | 6.40E-02 | 8.50E-03 |
| OCCIPITAL | Length | -1.80E-04 | -2.00E-05 | 1.70E-04 | 1.20E-05 |
| OCCIPITAL | MeanDepth | -1.50E-03 | 1.40E-03 | 1.00E-03 | 8.60E-04 |
| OCCIPITAL | Width | -1.20E-05 | 4.20E-06 | 1.20E-05 | 9.00E-07 |
| OCCIPITAL | SurfaceArea | 9.30E-03 | 2.80E-02 | -1.80E-02 | 7.30E-03 |
| S.C. | Length | -6.00E-05 | 1.60E-04 | -1.10E-04 | -1.70E-05 |
| S.C. | MeanDepth | -4.60E-19 | -6.00E-04 | -5.80E-04 | -3.80E-05 |
| S.C. | Width | 2.50E-07 | 2.70E-06 | -7.50E-06 | 1.70E-06 |
| S.C. | SurfaceArea | 5.40E-03 | -3.00E-03 | -4.70E-03 | -7.70E-03 |
| S.C.LPC. | Length | -1.90E-03 | 7.00E-03 | 1.40E-03 | -3.20E-04 |
| S.C.LPC. | MeanDepth | 2.70E-03 | 1.40E-03 | -3.90E-03 | 3.60E-04 |
| S.C.LPC. | Width | -9.60E-05 | 5.10E-04 | -3.40E-05 | -1.30E-05 |
| S.C.LPC. | SurfaceArea | 3.50E-03 | -5.30E-03 | 1.10E-02 | -1.00E-03 |
| S.C.sylvian. | Length | -3.30E-03 | -4.60E-04 | -3.20E-03 | 1.10E-03 |
| S.C.sylvian. | MeanDepth | -5.10E-04 | -4.80E-03 | -1.80E-05 | -7.80E-05 |
| S.C.sylvian. | Width | -1.70E-04 | -2.50E-04 | -1.30E-04 | -6.80E-06 |
| S.C.sylvian. | SurfaceArea | 7.80E-04 | 5.70E-02 | 2.50E-02 | 2.60E-02 |
| S.Call. | Length | 3.40E-04 | -2.10E-04 | 8.50E-05 | 1.80E-04 |

| | | | | | |
|---|---|---|---|---|---|
| S.Call. | MeanDepth | 2.00E-03 | 9.90E-04 | 2.00E-03 | 2.30E-03 |
| S.Call. | Width | 4.80E-05 | -1.90E-05 | 2.20E-05 | 2.90E-05 |
| S.Call. | SurfaceArea | -5.20E-03 | -9.40E-03 | -2.10E-03 | -9.50E-02 |
| S.Cu. | Length | 6.90E-04 | 1.50E-03 | 4.40E-06 | -4.60E-05 |
| S.Cu. | MeanDepth | 2.10E-03 | -6.00E-03 | -8.30E-04 | 2.90E-04 |
| S.Cu. | Width | 6.40E-05 | 7.50E-05 | -5.60E-06 | 5.70E-06 |
| S.Cu. | SurfaceArea | -6.40E-03 | 1.70E-02 | -4.80E-03 | -3.90E-02 |
| S.F.inf. | Length | 1.40E-04 | -6.60E-04 | 1.30E-04 | 3.10E-04 |
| S.F.inf. | MeanDepth | -8.30E-04 | 3.80E-04 | 2.40E-03 | 1.50E-03 |
| S.F.inf. | Width | -1.00E-05 | -4.20E-05 | 2.00E-05 | 1.60E-05 |
| S.F.inf. | SurfaceArea | 1.70E-02 | 1.60E-03 | -6.90E-03 | -1.80E-02 |
| S.F.inf.ant. | Length | 6.40E-04 | -2.20E-03 | 1.10E-04 | 4.60E-04 |
| S.F.inf.ant. | MeanDepth | -1.10E-03 | -3.70E-03 | -2.30E-03 | 2.10E-04 |
| S.F.inf.ant. | Width | 4.70E-05 | -1.70E-04 | -2.20E-05 | 4.10E-05 |
| S.F.inf.ant. | SurfaceArea | -1.60E-02 | -1.80E-03 | -9.10E-03 | -8.90E-03 |
| S.F.int. | Length | -7.50E-05 | 1.00E-05 | -3.10E-04 | -3.40E-06 |
| S.F.int. | MeanDepth | -3.70E-04 | 2.20E-03 | -5.50E-04 | -1.10E-03 |
| S.F.int. | Width | -9.50E-06 | 5.60E-06 | -2.50E-05 | -5.70E-06 |
| S.F.int. | SurfaceArea | -1.70E-03 | -1.30E-02 | -7.00E-03 | -1.70E-02 |
| S.F.inter. | Length | 4.40E-04 | 5.20E-06 | 1.30E-04 | 1.00E-04 |
| S.F.inter. | MeanDepth | -1.40E-04 | -4.00E-04 | -1.80E-03 | 1.00E-03 |
| S.F.inter. | Width | 2.00E-05 | -6.40E-06 | 1.00E-06 | 1.10E-05 |
| S.F.inter. | SurfaceArea | 2.60E-03 | 8.50E-03 | -1.20E-02 | -3.80E-03 |
| S.F.marginal. | Length | -3.70E-03 | 3.90E-04 | -1.60E-03 | -1.90E-04 |
| S.F.marginal. | MeanDepth | -5.80E-03 | 3.40E-03 | -3.40E-04 | -1.40E-03 |
| S.F.marginal. | Width | -3.00E-04 | 5.10E-05 | -1.50E-04 | -3.00E-05 |
| S.F.marginal. | SurfaceArea | 1.80E-02 | 1.70E-02 | -4.00E-03 | 8.90E-03 |
| S.F.median. | Length | -1.30E-03 | 9.00E-05 | 8.50E-04 | 1.10E-03 |
| S.F.median. | MeanDepth | -6.00E-03 | 2.30E-03 | -1.30E-03 | -9.50E-04 |
| S.F.median. | Width | -1.10E-04 | 3.90E-05 | 6.30E-05 | 1.00E-04 |
| S.F.median. | SurfaceArea | -1.20E-03 | -4.30E-05 | -7.30E-03 | -1.20E-03 |
| S.F.orbitaire. | Length | -7.80E-04 | -3.30E-04 | 6.70E-04 | 3.70E-04 |
| S.F.orbitaire. | MeanDepth | -3.40E-03 | -7.50E-03 | 4.50E-04 | -1.40E-03 |
| S.F.orbitaire. | Width | -2.30E-04 | -1.00E-04 | 8.30E-05 | 4.30E-05 |
| S.F.orbitaire. | SurfaceArea | 1.30E-03 | -5.00E-03 | 5.70E-03 | -1.40E-02 |
| S.F.polaire.tr. | Length | 9.60E-05 | 3.40E-04 | 1.20E-03 | -5.60E-04 |
| S.F.polaire.tr. | MeanDepth | 1.50E-03 | 2.10E-04 | 4.20E-03 | -5.00E-03 |
| S.F.polaire.tr. | Width | -1.10E-05 | 3.80E-05 | 8.90E-05 | -1.00E-04 |
| S.F.polaire.tr. | SurfaceArea | -8.60E-03 | -2.20E-02 | -1.50E-02 | 6.50E-03 |
| S.F.sup. | Length | -1.30E-04 | -4.50E-04 | -2.10E-04 | -1.30E-04 |
| S.F.sup. | MeanDepth | 6.20E-04 | 1.80E-03 | 6.10E-04 | 3.40E-04 |
| S.F.sup. | Width | -4.50E-06 | -1.60E-05 | -8.20E-06 | -3.30E-06 |
| S.F.sup. | SurfaceArea | 2.30E-03 | -6.70E-03 | -1.20E-02 | -4.30E-03 |
| S.Li.ant. | Length | -4.80E-03 | -8.60E-05 | -2.70E-03 | -2.10E-03 |
| S.Li.ant. | MeanDepth | 7.40E-04 | -2.50E-03 | 1.70E-03 | 4.40E-04 |
| S.Li.ant. | Width | -3.20E-04 | 7.40E-05 | -1.00E-04 | -1.80E-04 |

| | | | | | |
|---|---|---|---|---|---|
| S.Li.ant. | SurfaceArea | 2.20E-03 | 1.60E-01 | -4.20E-02 | 1.70E-02 |
| S.Li.post. | Length | -2.80E-03 | 1.40E-04 | 2.20E-03 | -1.80E-03 |
| S.Li.post. | MeanDepth | -7.00E-03 | -7.00E-03 | -7.50E-03 | -1.80E-03 |
| S.Li.post. | Width | -2.70E-04 | -6.30E-05 | 7.60E-05 | -1.40E-04 |
| S.Li.post. | SurfaceArea | 1.50E-02 | 1.60E-02 | 2.00E-03 | 3.40E-02 |
| S.O.p. | Length | 6.20E-03 | -2.50E-03 | -5.20E-03 | 3.10E-03 |
| S.O.p. | MeanDepth | 6.20E-03 | 2.50E-03 | -3.60E-05 | 1.40E-03 |
| S.O.p. | Width | 3.60E-04 | -1.60E-04 | -2.50E-04 | 1.50E-04 |
| S.O.p. | SurfaceArea | -1.20E-02 | 1.50E-02 | 6.10E-03 | -3.00E-02 |
| S.O.T.lat.ant. | Length | 1.00E-03 | 1.00E-03 | 5.10E-04 | 7.20E-04 |
| S.O.T.lat.ant. | MeanDepth | -2.80E-04 | 9.50E-04 | -3.70E-03 | 1.20E-03 |
| S.O.T.lat.ant. | Width | 3.10E-05 | 3.20E-05 | -4.70E-05 | 4.00E-05 |
| S.O.T.lat.ant. | SurfaceArea | 1.30E-02 | 1.20E-02 | 4.60E-02 | -3.50E-03 |
| S.O.T.lat.int. | Length | -6.20E-03 | -3.50E-03 | 1.70E-03 | -4.80E-03 |
| S.O.T.lat.int. | MeanDepth | 1.70E-02 | 4.30E-03 | -3.00E-03 | -3.40E-03 |
| S.O.T.lat.int. | Width | -2.40E-04 | -9.90E-05 | 8.80E-05 | -4.30E-04 |
| S.O.T.lat.int. | SurfaceArea | -1.50E-02 | -2.40E-02 | -1.40E-02 | -5.30E-03 |
| S.O.T.lat.med. | Length | -8.30E-04 | 9.20E-04 | -3.50E-03 | 6.60E-04 |
| S.O.T.lat.med. | MeanDepth | -1.40E-02 | 1.50E-03 | -7.70E-03 | 4.40E-04 |
| S.O.T.lat.med. | Width | -2.50E-04 | 8.50E-05 | -3.60E-04 | 3.70E-05 |
| S.O.T.lat.med. | SurfaceArea | 4.00E-02 | 4.20E-02 | 2.10E-02 | -4.50E-02 |
| S.O.T.lat.post. | Length | 2.30E-03 | 2.20E-04 | 2.00E-03 | 5.70E-04 |
| S.O.T.lat.post. | MeanDepth | 4.40E-04 | 1.60E-03 | 3.70E-03 | -1.60E-03 |
| S.O.T.lat.post. | Width | 1.30E-04 | -2.30E-05 | 1.20E-04 | 3.10E-05 |
| S.O.T.lat.post. | SurfaceArea | 2.30E-02 | 6.50E-02 | 4.40E-03 | 1.50E-02 |
| S.Olf. | Length | 1.20E-04 | 4.80E-05 | 7.40E-04 | 1.20E-04 |
| S.Olf. | MeanDepth | 1.50E-03 | -5.70E-04 | 2.40E-03 | 1.80E-03 |
| S.Olf. | Width | 3.20E-05 | 2.60E-05 | 1.20E-04 | 4.60E-05 |
| S.Olf. | SurfaceArea | -1.60E-02 | 5.30E-02 | -9.90E-03 | -6.80E-02 |
| S.Or. | Length | -1.70E-04 | -1.20E-04 | -9.50E-04 | 1.10E-04 |
| S.Or. | MeanDepth | -1.50E-04 | -1.10E-03 | 1.70E-03 | 1.70E-03 |
| S.Or. | Width | -1.20E-06 | -2.20E-05 | -6.30E-05 | 2.10E-05 |
| S.Or. | SurfaceArea | 4.80E-03 | 5.10E-03 | -4.30E-03 | -1.40E-02 |
| S.p.C. | Length | -1.70E-03 | 6.00E-04 | 8.80E-04 | 1.40E-03 |
| S.p.C. | MeanDepth | 2.80E-03 | -4.40E-03 | -7.50E-03 | 1.20E-04 |
| S.p.C. | Width | -7.10E-05 | -3.10E-05 | -7.30E-05 | 1.70E-04 |
| S.p.C. | SurfaceArea | -4.40E-03 | 5.90E-03 | 1.30E-02 | -1.50E-03 |
| S.Pa.int. | Length | -1.40E-03 | 3.40E-05 | 3.70E-04 | 1.70E-03 |
| S.Pa.int. | MeanDepth | -2.60E-03 | -1.60E-03 | -7.00E-03 | 6.50E-03 |
| S.Pa.int. | Width | -1.20E-04 | -4.80E-05 | -6.30E-05 | 8.60E-05 |
| S.Pa.int. | SurfaceArea | 4.10E-03 | -2.50E-02 | -1.40E-02 | 2.10E-02 |
| S.Pa.sup. | Length | 5.70E-04 | 1.50E-03 | 3.10E-03 | -9.70E-04 |
| S.Pa.sup. | MeanDepth | -2.70E-03 | 5.50E-03 | 9.30E-04 | -1.60E-03 |
| S.Pa.sup. | Width | 4.70E-05 | 2.20E-04 | 2.80E-04 | -1.10E-04 |
| S.Pa.sup. | SurfaceArea | 3.00E-03 | -3.10E-03 | -2.30E-02 | 1.20E-02 |
| S.Pa.t. | Length | -1.80E-03 | 6.00E-03 | -9.00E-03 | -2.60E-04 |

| | | | | | |
|---|---|---|---|---|---|
| S.Pa.t. | MeanDepth | 1.90E-03 | -6.20E-03 | -4.80E-05 | -1.10E-03 |
| S.Pa.t. | Width | -1.80E-05 | 1.50E-05 | -1.40E-04 | 4.00E-05 |
| S.Pa.t. | SurfaceArea | 1.10E-02 | 2.90E-02 | 2.90E-02 | 3.20E-02 |
| S.Pe.C.inf. | Length | -7.30E-04 | 2.00E-04 | 5.00E-04 | -5.70E-04 |
| S.Pe.C.inf. | MeanDepth | 8.40E-04 | 1.90E-03 | 3.00E-03 | -5.70E-04 |
| S.Pe.C.inf. | Width | -2.00E-05 | 1.60E-05 | 5.40E-05 | -1.70E-05 |
| S.Pe.C.inf. | SurfaceArea | -2.90E-03 | -7.20E-03 | -2.40E-02 | 1.00E-03 |
| S.Pe.C.inter. | Length | 4.30E-04 | -3.00E-05 | 5.60E-05 | 7.00E-05 |
| S.Pe.C.inter. | MeanDepth | -7.20E-04 | -2.80E-04 | -1.50E-03 | -5.80E-05 |
| S.Pe.C.inter. | Width | 1.10E-05 | -7.00E-06 | -1.20E-05 | 1.50E-06 |
| S.Pe.C.inter. | SurfaceArea | 9.70E-03 | -8.70E-03 | -4.20E-03 | -1.20E-02 |
| S.Pe.C.marginal. | Length | 4.50E-03 | 5.10E-03 | 5.00E-04 | -5.20E-04 |
| S.Pe.C.marginal. | MeanDepth | 5.40E-03 | 3.70E-03 | 2.10E-06 | 1.10E-03 |
| S.Pe.C.marginal. | Width | 1.90E-04 | 2.30E-04 | 2.50E-05 | 1.80E-05 |
| S.Pe.C.marginal. | SurfaceArea | -2.10E-02 | -1.40E-03 | -2.90E-02 | 3.00E-03 |
| S.Pe.C.median. | Length | -1.90E-03 | 2.30E-03 | 2.20E-03 | 3.30E-04 |
| S.Pe.C.median. | MeanDepth | -2.90E-03 | 4.20E-03 | -2.60E-03 | 1.80E-04 |
| S.Pe.C.median. | Width | -1.90E-04 | 2.40E-04 | 8.60E-05 | 4.30E-05 |
| S.Pe.C.median. | SurfaceArea | 1.10E-02 | -7.90E-03 | 1.50E-03 | -1.70E-03 |
| S.Pe.C.sup. | Length | -4.90E-03 | -3.70E-04 | 7.40E-05 | -1.80E-03 |
| S.Pe.C.sup. | MeanDepth | -5.10E-03 | 3.30E-03 | -2.30E-03 | 7.70E-05 |
| S.Pe.C.sup. | Width | -2.40E-04 | -4.00E-06 | -1.00E-05 | -5.20E-05 |
| S.Pe.C.sup. | SurfaceArea | -6.90E-03 | 3.80E-02 | -1.50E-02 | -3.40E-02 |
| S.Po.C.sup. | Length | -3.20E-03 | -4.70E-04 | 3.70E-04 | 1.30E-03 |
| S.Po.C.sup. | MeanDepth | -1.40E-03 | 3.10E-03 | 2.50E-03 | 2.90E-03 |
| S.Po.C.sup. | Width | -1.60E-04 | -1.40E-05 | -1.30E-05 | 7.80E-05 |
| S.Po.C.sup. | SurfaceArea | 3.80E-03 | -8.00E-03 | -9.50E-03 | -1.00E-02 |
| S.R.inf. | Length | 3.40E-03 | 9.20E-05 | 4.50E-03 | -3.00E-04 |
| S.R.inf. | MeanDepth | -5.70E-03 | 1.80E-03 | 1.20E-02 | 1.50E-03 |
| S.R.inf. | Width | 3.10E-04 | -6.00E-06 | 6.60E-04 | 7.80E-05 |
| S.R.inf. | SurfaceArea | 9.90E-03 | -5.40E-02 | -2.50E-02 | -1.60E-02 |
| S.Rh. | Length | -5.40E-05 | -5.30E-04 | -4.60E-04 | -3.30E-03 |
| S.Rh. | MeanDepth | 3.50E-03 | -5.30E-03 | -9.70E-03 | -2.70E-03 |
| S.Rh. | Width | -2.50E-06 | -1.60E-04 | -3.50E-04 | -3.10E-04 |
| S.Rh. | SurfaceArea | 3.90E-03 | 8.80E-02 | 1.10E-02 | 1.90E-02 |
| S.s.P. | Length | 2.40E-04 | 9.00E-04 | 6.00E-04 | -8.30E-05 |
| S.s.P. | MeanDepth | -4.20E-03 | -1.10E-03 | -3.00E-03 | 3.20E-03 |
| S.s.P. | Width | -3.40E-05 | 5.70E-06 | 3.60E-05 | 5.90E-05 |
| S.s.P. | SurfaceArea | 1.90E-02 | 6.80E-02 | -1.60E-02 | -6.50E-02 |
| S.T.i.ant. | Length | -3.80E-04 | -2.70E-04 | -1.80E-04 | 1.70E-04 |
| S.T.i.ant. | MeanDepth | -2.30E-04 | -5.70E-06 | -1.60E-04 | 1.10E-03 |
| S.T.i.ant. | Width | -2.90E-05 | -1.50E-05 | -1.60E-05 | 1.20E-05 |
| S.T.i.ant. | SurfaceArea | 7.00E-03 | 4.40E-03 | 5.20E-03 | -1.60E-02 |
| S.T.i.post. | Length | -9.40E-04 | -3.70E-04 | -7.70E-04 | 1.90E-04 |
| S.T.i.post. | MeanDepth | -1.30E-03 | -3.80E-04 | -1.80E-03 | -4.30E-04 |
| S.T.i.post. | Width | -4.20E-05 | -2.40E-05 | -2.70E-05 | 1.30E-05 |

| | | | | | |
|---|---|---|---|---|---|
| S.T.i.post. | SurfaceArea | 1.20E-02 | 5.20E-02 | 1.70E-02 | -7.40E-03 |
| S.T.pol. | Length | 2.80E-04 | 2.50E-03 | 8.00E-04 | 2.10E-04 |
| S.T.pol. | MeanDepth | 3.80E-03 | -3.90E-04 | -1.40E-02 | -2.30E-03 |
| S.T.pol. | Width | 6.10E-06 | 1.60E-04 | -6.50E-05 | 2.00E-06 |
| S.T.pol. | SurfaceArea | -1.00E-02 | -3.00E-02 | 1.80E-02 | -1.40E-02 |
| S.T.s. | Length | 3.20E-05 | 4.00E-04 | -3.20E-04 | 2.00E-04 |
| S.T.s. | MeanDepth | -3.80E-04 | -1.50E-03 | 1.30E-03 | -4.80E-04 |
| S.T.s. | Width | -5.70E-08 | 9.40E-06 | -5.80E-06 | 6.50E-06 |
| S.T.s. | SurfaceArea | 1.10E-02 | 1.20E-02 | 4.70E-03 | -1.80E-02 |
| S.T.s.ter.asc.ant. | Length | -2.70E-03 | 9.30E-06 | 2.80E-04 | 6.40E-04 |
| S.T.s.ter.asc.ant. | MeanDepth | 1.30E-04 | -3.40E-03 | 7.60E-04 | -3.80E-04 |
| S.T.s.ter.asc.ant. | Width | -1.30E-04 | -3.60E-05 | 1.10E-05 | 2.50E-05 |
| S.T.s.ter.asc.ant. | SurfaceArea | 5.90E-03 | -2.90E-02 | -2.20E-03 | -6.70E-03 |
| S.T.s.ter.asc.post. | Length | 2.00E-03 | 6.40E-04 | 2.50E-04 | 6.90E-04 |
| S.T.s.ter.asc.post. | MeanDepth | 2.00E-03 | 1.90E-03 | 1.70E-03 | 2.60E-04 |
| S.T.s.ter.asc.post. | Width | 9.00E-05 | 3.00E-05 | 2.50E-06 | 1.30E-05 |
| S.T.s.ter.asc.post. | SurfaceArea | 6.50E-03 | -1.30E-02 | -1.10E-02 | 2.60E-02 |

**Table S9:** bias (*b*) values for the bilaterally average sulcal descriptors (red: abs(b) > 0.1 or NaN; orange: abs(b) > 1)

| Sulci | Descriptor | OASIS | KKI | QTIM | HCP |
|---|---|---|---|---|---|
| F.C.L.a. | Length | -3.20E-02 | -5.00E-02 | -9.10E-02 | 1.50E-02 |
| F.C.L.a. | MeanDepth | 8.70E-03 | -8.30E-02 | -1.80E-02 | -3.50E-02 |
| F.C.L.a. | Width | -2.00E-02 | 1.20E-02 | -2.50E-02 | 5.20E-02 |
| F.C.L.a. | SurfaceArea | -2.60E-02 | -7.90E-02 | -1.20E-01 | 2.90E-02 |
| F.C.L.p. | Length | -8.10E-03 | 4.80E-03 | -1.70E-03 | -1.00E-03 |
| F.C.L.p. | MeanDepth | 2.80E-02 | -3.10E-02 | 9.50E-03 | 2.20E-02 |
| F.C.L.p. | Width | -1.50E-02 | 1.00E-02 | -2.70E-02 | -3.60E-03 |
| F.C.L.p. | SurfaceArea | 6.80E-03 | -2.20E-03 | -5.50E-03 | -8.60E-03 |
| F.C.L.r.ant. | Length | 1.30E-02 | 7.10E-03 | 3.10E-02 | -1.90E-03 |
| F.C.L.r.ant. | MeanDepth | 2.40E-03 | -1.60E-02 | -4.20E-03 | -1.70E-02 |
| F.C.L.r.ant. | Width | -3.30E-04 | 3.00E-02 | -3.80E-02 | -1.50E-03 |
| F.C.L.r.ant. | SurfaceArea | -1.70E-02 | 5.20E-03 | 2.80E-02 | -6.40E-03 |
| F.C.L.r.asc. | Length | -1.20E-04 | 8.90E-03 | -3.00E-02 | -1.40E-02 |
| F.C.L.r.asc. | MeanDepth | -3.20E-02 | -1.10E-02 | 1.30E-02 | -4.00E-02 |
| F.C.L.r.asc. | Width | 1.70E-02 | 5.30E-03 | -5.40E-02 | -1.90E-02 |
| F.C.L.r.asc. | SurfaceArea | -1.60E-02 | -1.40E-02 | -1.20E-02 | -1.70E-02 |
| F.C.L.r.diag. | Length | -2.00E-02 | -7.30E-02 | -6.40E-02 | -2.00E-03 |
| F.C.L.r.diag. | MeanDepth | 3.50E-02 | 1.20E-02 | 3.50E-02 | 1.60E-02 |
| F.C.L.r.diag. | Width | -1.10E-02 | -6.20E-02 | -9.30E-04 | -1.70E-02 |
| F.C.L.r.diag. | SurfaceArea | -2.30E-02 | -1.70E-02 | 1.60E-03 | 1.90E-02 |
| F.C.L.r.retroC.tr. | Length | 8.80E-02 | -6.60E-02 | -3.50E-02 | 5.50E-02 |
| F.C.L.r.retroC.tr. | MeanDepth | -6.30E-02 | -4.40E-02 | 4.00E-02 | -1.50E-02 |
| F.C.L.r.retroC.tr. | Width | -4.10E-02 | -5.00E-02 | -4.70E-02 | -7.40E-03 |
| F.C.L.r.retroC.tr. | SurfaceArea | 2.10E-02 | -4.70E-02 | -1.30E-02 | -7.30E-04 |
| F.C.L.r.sc.ant. | Length | -1.20E-01 | 2.00E+00 | 9.20E-03 | 0.00E+00 |
| F.C.L.r.sc.ant. | MeanDepth | 3.70E-02 | -2.50E-02 | 1.50E-01 | 0.00E+00 |
| F.C.L.r.sc.ant. | Width | 8.80E-03 | -2.90E-01 | 5.70E-02 | 0.00E+00 |
| F.C.L.r.sc.ant. | SurfaceArea | 1.20E-03 | 6.70E-01 | 3.10E-02 | 0.00E+00 |
| F.C.L.r.sc.post. | Length | 1.70E-02 | 3.00E-01 | -1.90E-02 | 1.60E-01 |
| F.C.L.r.sc.post. | MeanDepth | 5.20E-03 | -3.50E-02 | 2.70E-02 | 5.50E-02 |
| F.C.L.r.sc.post. | Width | -4.30E-02 | 2.20E-02 | 2.40E-02 | 3.00E-02 |
| F.C.L.r.sc.post. | SurfaceArea | 4.80E-02 | 6.40E-02 | -2.10E-02 | 1.40E-03 |
| F.C.M.ant. | Length | -1.30E-02 | -2.70E-02 | 3.40E-02 | 8.60E-03 |
| F.C.M.ant. | MeanDepth | 2.70E-02 | -2.80E-02 | 1.50E-02 | 2.80E-03 |
| F.C.M.ant. | Width | 7.60E-03 | -3.10E-02 | -5.80E-02 | 1.60E-02 |
| F.C.M.ant. | SurfaceArea | -1.30E-02 | -3.90E-02 | 5.00E-02 | 2.20E-02 |
| F.C.M.post. | Length | 2.30E-02 | 5.20E-03 | 2.30E-03 | -2.30E-02 |
| F.C.M.post. | MeanDepth | -1.50E-02 | -5.70E-03 | 5.80E-03 | 3.40E-03 |
| F.C.M.post. | Width | -1.50E-02 | -2.20E-02 | -3.30E-02 | -1.60E-02 |
| F.C.M.post. | SurfaceArea | 1.50E-02 | -5.30E-03 | 4.80E-03 | -2.10E-02 |
| F.Cal.ant.-Sc.Cal. | Length | -3.10E-02 | 1.00E-02 | 2.50E-02 | 3.40E-02 |
| F.Cal.ant.-Sc.Cal. | MeanDepth | 2.10E-03 | -9.70E-03 | 5.80E-03 | -2.00E-02 |
| F.Cal.ant.-Sc.Cal. | Width | -3.30E-02 | 3.20E-02 | -5.80E-02 | -1.90E-02 |

| | | | | | |
|---|---|---|---|---|---|
| F.Cal.ant.-Sc.Cal. | SurfaceArea | -4.00E-02 | -1.20E-02 | 3.20E-02 | 2.50E-02 |
| F.Coll. | Length | 5.60E-02 | 3.40E-02 | -2.30E-02 | -9.40E-04 |
| F.Coll. | MeanDepth | 5.20E-03 | 9.40E-04 | -5.90E-03 | -6.20E-03 |
| F.Coll. | Width | 1.70E-02 | 2.80E-02 | -2.40E-02 | -8.50E-03 |
| F.Coll. | SurfaceArea | 5.60E-02 | 3.70E-02 | -3.90E-02 | -2.30E-03 |
| F.I.P. | Length | -1.70E-04 | 3.70E-03 | -7.00E-03 | -2.50E-02 |
| F.I.P. | MeanDepth | 7.30E-03 | -6.70E-03 | -1.10E-02 | 6.20E-03 |
| F.I.P. | Width | -1.40E-03 | -1.80E-02 | -2.50E-02 | -7.50E-03 |
| F.I.P. | SurfaceArea | -6.10E-03 | -2.70E-03 | -6.70E-04 | -1.40E-02 |
| F.I.P.Po.C.inf. | Length | 8.90E-02 | 3.80E-02 | 8.00E-03 | 7.10E-03 |
| F.I.P.Po.C.inf. | MeanDepth | 9.40E-03 | 1.80E-03 | -2.30E-02 | 1.10E-02 |
| F.I.P.Po.C.inf. | Width | 8.80E-04 | -1.50E-02 | -1.60E-02 | -1.20E-02 |
| F.I.P.Po.C.inf. | SurfaceArea | 9.90E-02 | 3.70E-02 | -1.90E-02 | 2.10E-02 |
| F.I.P.r.int.1 | Length | 3.00E-02 | -9.70E-03 | -1.10E-02 | 2.50E-03 |
| F.I.P.r.int.1 | MeanDepth | -7.90E-03 | -6.00E-04 | 3.00E-02 | 3.50E-05 |
| F.I.P.r.int.1 | Width | -7.50E-02 | -6.00E-02 | 7.70E-03 | 1.80E-02 |
| F.I.P.r.int.1 | SurfaceArea | 7.20E-04 | -2.80E-02 | 3.40E-03 | -1.50E-02 |
| F.I.P.r.int.2 | Length | 1.20E-01 | 3.30E-02 | -4.20E-02 | -3.50E-02 |
| F.I.P.r.int.2 | MeanDepth | 6.70E-02 | -5.60E-02 | -7.50E-02 | -4.10E-03 |
| F.I.P.r.int.2 | Width | 1.60E-03 | 5.50E-02 | 5.10E-02 | 4.90E-02 |
| F.I.P.r.int.2 | SurfaceArea | 2.10E-01 | -3.00E-02 | -1.40E-01 | -2.40E-02 |
| F.P.O. | Length | 2.80E-02 | -8.30E-02 | -1.40E-02 | -2.70E-02 |
| F.P.O. | MeanDepth | -2.20E-02 | -5.10E-03 | 1.50E-02 | -5.40E-03 |
| F.P.O. | Width | -7.30E-03 | -1.70E-02 | -3.40E-02 | 1.30E-03 |
| F.P.O. | SurfaceArea | 1.90E-02 | -3.30E-02 | -6.80E-03 | -9.10E-03 |
| INSULA | Length | 1.30E-01 | 3.90E-01 | 7.00E-01 | -1.40E-01 |
| INSULA | MeanDepth | 2.50E-02 | -3.30E-02 | -1.40E-02 | -5.40E-03 |
| INSULA | Width | 2.70E-02 | 1.30E-02 | 1.10E-01 | -3.60E-03 |
| INSULA | SurfaceArea | -5.10E-03 | 2.80E-03 | 3.00E-02 | 4.30E-03 |
| OCCIPITAL | Length | 3.90E-03 | -3.10E-03 | 2.80E-02 | -8.70E-03 |
| OCCIPITAL | MeanDepth | -1.40E-02 | 3.10E-03 | 7.10E-03 | 9.00E-03 |
| OCCIPITAL | Width | 3.00E-02 | 2.90E-02 | -5.60E-02 | -7.70E-03 |
| OCCIPITAL | SurfaceArea | -1.10E-02 | 6.10E-03 | 4.50E-02 | -7.30E-03 |
| S.C. | Length | 3.60E-03 | 2.20E-02 | -2.50E-02 | 3.30E-03 |
| S.C. | MeanDepth | -4.80E-04 | -6.00E-03 | -2.30E-03 | 2.20E-03 |
| S.C. | Width | 8.30E-03 | -1.00E-02 | -2.30E-02 | -8.40E-03 |
| S.C. | SurfaceArea | 4.40E-04 | 1.50E-02 | -3.40E-02 | 1.20E-02 |
| S.C.LPC. | Length | -2.40E-01 | 8.70E-02 | -5.70E-02 | -7.30E-02 |
| S.C.LPC. | MeanDepth | -4.30E-02 | -8.30E-03 | -2.20E-17 | 1.70E-02 |
| S.C.LPC. | Width | 1.60E-01 | 5.60E-03 | 5.30E-02 | -8.90E-03 |
| S.C.LPC. | SurfaceArea | -2.60E-01 | 5.80E-02 | -6.20E-02 | -1.80E-02 |
| S.C.sylvian. | Length | -3.00E-02 | -4.10E-02 | -9.50E-02 | -6.90E-03 |
| S.C.sylvian. | MeanDepth | 6.30E-02 | -2.40E-02 | 2.90E-02 | 1.40E-02 |
| S.C.sylvian. | Width | -6.00E-03 | -2.60E-02 | -1.10E-01 | -4.50E-02 |
| S.C.sylvian. | SurfaceArea | -4.70E-03 | -6.60E-02 | -7.20E-02 | 1.50E-02 |
| S.Call. | Length | 6.40E-04 | 1.70E-03 | 4.60E-03 | -3.00E-02 |

| | | | | | |
|---|---|---|---|---|---|
| S.Call. | MeanDepth | -5.00E-03 | -2.90E-02 | -7.20E-03 | 3.30E-03 |
| S.Call. | Width | 2.90E-02 | -2.30E-02 | -6.40E-02 | -6.80E-02 |
| S.Call. | SurfaceArea | -1.40E-02 | -2.10E-02 | -3.00E-03 | -2.00E-02 |
| S.Cu. | Length | -9.30E-03 | 2.80E-02 | -3.30E-03 | 2.10E-02 |
| S.Cu. | MeanDepth | 2.40E-02 | -1.40E-02 | -2.30E-02 | -1.80E-02 |
| S.Cu. | Width | -1.40E-02 | 4.10E-02 | 5.40E-03 | -4.30E-02 |
| S.Cu. | SurfaceArea | -1.50E-02 | 2.10E-02 | -2.70E-02 | 3.60E-03 |
| S.F.inf. | Length | -2.50E-02 | -2.90E-02 | 9.30E-03 | -2.90E-02 |
| S.F.inf. | MeanDepth | -1.80E-02 | 3.80E-03 | 1.60E-02 | 2.20E-02 |
| S.F.inf. | Width | 2.50E-02 | -3.70E-03 | -3.60E-02 | -1.70E-02 |
| S.F.inf. | SurfaceArea | -4.30E-02 | -3.90E-02 | 2.30E-02 | -3.30E-03 |
| S.F.inf.ant. | Length | -1.80E-02 | -8.20E-02 | -3.20E-02 | 4.20E-02 |
| S.F.inf.ant. | MeanDepth | -7.30E-03 | -4.30E-03 | -1.70E-02 | -6.00E-03 |
| S.F.inf.ant. | Width | -2.50E-02 | -2.10E-02 | -1.10E-02 | -8.90E-03 |
| S.F.inf.ant. | SurfaceArea | -2.20E-02 | -8.80E-02 | -5.10E-02 | 3.80E-02 |
| S.F.int. | Length | -1.00E-02 | 1.10E-02 | -3.80E-02 | 1.20E-03 |
| S.F.int. | MeanDepth | 8.50E-04 | 8.70E-03 | -5.30E-03 | -3.00E-03 |
| S.F.int. | Width | -1.90E-03 | -1.70E-02 | -3.20E-02 | -3.00E-02 |
| S.F.int. | SurfaceArea | -1.90E-03 | 1.90E-02 | -3.80E-02 | -5.60E-04 |
| S.F.inter. | Length | 5.80E-02 | 2.10E-02 | 2.10E-02 | 5.30E-04 |
| S.F.inter. | MeanDepth | 2.60E-03 | 3.50E-03 | -1.20E-02 | 1.00E-03 |
| S.F.inter. | Width | 5.90E-03 | -1.20E-02 | -1.70E-02 | -5.00E-03 |
| S.F.inter. | SurfaceArea | 6.00E-02 | 4.20E-03 | 1.30E-02 | 9.10E-03 |
| S.F.marginal. | Length | -1.40E-01 | 2.60E-03 | -4.40E-02 | 1.10E-02 |
| S.F.marginal. | MeanDepth | -6.00E-02 | 1.10E-02 | 1.20E-02 | 2.10E-04 |
| S.F.marginal. | Width | 1.10E-01 | 4.50E-02 | -4.40E-02 | -8.80E-03 |
| S.F.marginal. | SurfaceArea | -1.80E-01 | 1.20E-02 | -4.20E-02 | 8.20E-03 |
| S.F.median. | Length | -9.00E-02 | -1.90E-02 | 1.20E-02 | 5.10E-02 |
| S.F.median. | MeanDepth | -2.20E-02 | 2.20E-02 | -1.60E-02 | 4.00E-03 |
| S.F.median. | Width | 1.30E-02 | -2.30E-02 | -2.90E-02 | 1.20E-02 |
| S.F.median. | SurfaceArea | -9.00E-02 | 1.90E-02 | -4.70E-03 | 6.10E-02 |
| S.F.orbitaire. | Length | -3.70E-02 | -9.30E-02 | 8.00E-02 | 3.40E-03 |
| S.F.orbitaire. | MeanDepth | 7.70E-03 | -3.20E-02 | 3.80E-02 | -1.50E-02 |
| S.F.orbitaire. | Width | -2.80E-02 | -3.60E-03 | -4.50E-02 | -2.20E-02 |
| S.F.orbitaire. | SurfaceArea | -4.70E-02 | -1.10E-01 | 1.10E-01 | -1.00E-02 |
| S.F.polaire.tr. | Length | -1.80E-02 | -1.80E-03 | -1.00E-02 | -3.90E-02 |
| S.F.polaire.tr. | MeanDepth | -1.50E-03 | -5.50E-03 | -2.70E-03 | -2.20E-02 |
| S.F.polaire.tr. | Width | -8.60E-03 | -4.60E-02 | -4.90E-02 | 1.10E-02 |
| S.F.polaire.tr. | SurfaceArea | -3.00E-02 | 1.70E-03 | -2.80E-02 | -5.80E-02 |
| S.F.sup. | Length | 1.00E-02 | -4.20E-02 | -7.10E-03 | -1.70E-02 |
| S.F.sup. | MeanDepth | -1.20E-02 | 4.80E-03 | 8.00E-03 | -4.00E-03 |
| S.F.sup. | Width | 1.90E-02 | -1.50E-02 | -2.90E-02 | -9.20E-03 |
| S.F.sup. | SurfaceArea | -2.50E-03 | -4.20E-02 | -7.80E-03 | -1.80E-02 |
| S.Li.ant. | Length | - | - | - | - |
| S.Li.ant. | MeanDepth | - | - | - | - |
| S.Li.ant. | Width | - | - | - | - |

| | | | | | |
|---|---|---|---|---|---|
| S.Li.ant. | SurfaceArea | - | - | - | - |
| S.Li.post. | Length | 2.80E-02 | 2.10E-02 | 1.50E-03 | 2.30E-03 |
| S.Li.post. | MeanDepth | -5.30E-03 | 8.70E-03 | 1.90E-02 | 1.40E-02 |
| S.Li.post. | Width | -8.80E-02 | 2.10E-02 | -7.90E-02 | -1.40E-02 |
| S.Li.post. | SurfaceArea | 1.60E-02 | 4.10E-02 | 6.20E-03 | -7.50E-03 |
| S.O.p. | Length | 7.60E-02 | 1.90E-02 | 7.10E-02 | 4.50E-02 |
| S.O.p. | MeanDepth | 1.80E-02 | 2.80E-03 | 3.00E-02 | -3.00E-03 |
| S.O.p. | Width | 1.90E-02 | 5.30E-02 | -5.00E-02 | -1.50E-02 |
| S.O.p. | SurfaceArea | 9.20E-02 | -4.30E-03 | 8.60E-02 | 4.60E-02 |
| S.O.T.lat.ant. | Length | -4.60E-02 | -6.40E-02 | 3.90E-02 | 3.30E-02 |
| S.O.T.lat.ant. | MeanDepth | -2.80E-02 | -1.80E-02 | -1.30E-03 | 1.00E-02 |
| S.O.T.lat.ant. | Width | 3.00E-02 | 5.50E-03 | -6.40E-02 | 1.60E-02 |
| S.O.T.lat.ant. | SurfaceArea | -8.50E-02 | -8.40E-02 | 5.10E-02 | 3.90E-02 |
| S.O.T.lat.int. | Length | -9.90E-03 | 2.60E-02 | 1.90E-02 | 3.60E-02 |
| S.O.T.lat.int. | MeanDepth | 2.30E-02 | 1.30E-02 | 2.20E-03 | 1.20E-02 |
| S.O.T.lat.int. | Width | 2.50E-02 | 9.80E-03 | 9.00E-02 | -1.30E-02 |
| S.O.T.lat.int. | SurfaceArea | -5.00E-03 | 1.10E-02 | -1.70E-02 | 3.70E-02 |
| S.O.T.lat.med. | Length | -3.20E-02 | -1.40E-02 | 3.00E-03 | -2.00E-02 |
| S.O.T.lat.med. | MeanDepth | 3.20E-02 | 3.00E-02 | -1.80E-02 | -3.50E-02 |
| S.O.T.lat.med. | Width | -1.10E-02 | -5.60E-02 | -4.20E-02 | -1.10E-02 |
| S.O.T.lat.med. | SurfaceArea | -3.70E-02 | 1.90E-02 | 1.90E-02 | -5.40E-02 |
| S.O.T.lat.post. | Length | -5.40E-02 | 6.90E-02 | -5.50E-02 | -5.50E-02 |
| S.O.T.lat.post. | MeanDepth | -7.60E-02 | 1.20E-02 | -4.00E-02 | -1.30E-02 |
| S.O.T.lat.post. | Width | 7.50E-02 | 2.70E-02 | 5.70E-03 | 3.60E-03 |
| S.O.T.lat.post. | SurfaceArea | -1.30E-01 | 6.20E-02 | -1.10E-01 | -5.20E-02 |
| S.Olf. | Length | 3.40E-02 | -9.50E-02 | -7.70E-02 | 2.00E-02 |
| S.Olf. | MeanDepth | 9.60E-03 | 2.20E-03 | 6.00E-02 | 2.10E-03 |
| S.Olf. | Width | 1.20E-01 | -2.50E-02 | -6.80E-02 | -3.90E-02 |
| S.Olf. | SurfaceArea | 5.20E-02 | -1.10E-01 | -2.70E-03 | 2.10E-02 |
| S.Or. | Length | 2.10E-02 | -1.40E-02 | 9.80E-02 | 1.10E-02 |
| S.Or. | MeanDepth | -1.80E-02 | -1.30E-03 | -1.90E-02 | 8.00E-03 |
| S.Or. | Width | -2.10E-02 | 3.20E-02 | -1.20E-02 | -1.30E-01 |
| S.Or. | SurfaceArea | 8.30E-03 | -6.80E-03 | 7.10E-02 | 3.00E-02 |
| S.p.C. | Length | 4.30E-02 | 8.40E-02 | 1.30E-02 | 2.30E-02 |
| S.p.C. | MeanDepth | -2.50E-02 | 8.40E-03 | 9.80E-03 | -2.30E-02 |
| S.p.C. | Width | 1.40E-02 | 1.90E-02 | -3.60E-02 | 2.00E-02 |
| S.p.C. | SurfaceArea | 6.70E-02 | 8.70E-02 | 2.10E-02 | -4.90E-03 |
| S.Pa.int. | Length | 8.20E-03 | 1.10E-02 | -6.10E-02 | -2.80E-03 |
| S.Pa.int. | MeanDepth | 7.40E-03 | -8.70E-03 | -2.50E-02 | 1.20E-02 |
| S.Pa.int. | Width | 2.60E-03 | -2.70E-03 | 3.70E-02 | -6.20E-03 |
| S.Pa.int. | SurfaceArea | 1.60E-02 | -1.20E-02 | -7.50E-02 | 1.30E-02 |
| S.Pa.sup. | Length | -5.40E-02 | 2.00E-02 | 6.00E-03 | 1.90E-02 |
| S.Pa.sup. | MeanDepth | -7.50E-03 | 1.30E-02 | -2.50E-02 | 2.10E-02 |
| S.Pa.sup. | Width | 1.10E-03 | -1.30E-02 | -4.50E-02 | 1.40E-02 |
| S.Pa.sup. | SurfaceArea | -5.80E-02 | 2.20E-02 | -1.10E-02 | -5.10E-03 |
| S.Pa.t. | Length | -3.00E-02 | -5.90E-02 | 2.80E-02 | -3.00E-02 |

| | | | | | |
|---|---|---|---|---|---|
| S.Pa.t. | MeanDepth | -9.30E-03 | -7.30E-03 | -1.80E-02 | 4.30E-04 |
| S.Pa.t. | Width | 1.30E-02 | 9.30E-03 | -4.80E-02 | -4.70E-03 |
| S.Pa.t. | SurfaceArea | -1.10E-02 | -2.30E-02 | 6.10E-03 | -3.70E-02 |
| S.Pe.C.inf. | Length | 2.80E-02 | 2.10E-01 | -2.30E-01 | 2.40E-02 |
| S.Pe.C.inf. | MeanDepth | -3.60E-03 | -8.30E-03 | 1.10E-02 | 9.80E-03 |
| S.Pe.C.inf. | Width | 2.20E-02 | 9.90E-02 | 2.60E-02 | -3.30E-02 |
| S.Pe.C.inf. | SurfaceArea | 4.20E-02 | 7.70E-02 | -1.40E-01 | 5.20E-02 |
| S.Pe.C.inter. | Length | -6.30E-03 | 2.90E-02 | -2.00E-02 | -2.10E-02 |
| S.Pe.C.inter. | MeanDepth | -1.30E-03 | 1.00E-02 | 2.40E-02 | 1.50E-03 |
| S.Pe.C.inter. | Width | -2.10E-02 | -3.30E-02 | -3.20E-02 | -7.90E-03 |
| S.Pe.C.inter. | SurfaceArea | -2.70E-02 | 1.10E-02 | 2.30E-02 | -9.30E-03 |
| S.Pe.C.marginal. | Length | -5.90E-03 | 3.20E-02 | 1.20E-03 | -8.40E-03 |
| S.Pe.C.marginal. | MeanDepth | -3.60E-03 | 2.60E-02 | -4.20E-02 | -8.30E-03 |
| S.Pe.C.marginal. | Width | 1.10E-02 | -3.80E-02 | -4.00E-03 | -1.40E-02 |
| S.Pe.C.marginal. | SurfaceArea | -1.40E-03 | 3.50E-02 | -3.00E-02 | -1.10E-02 |
| S.Pe.C.median. | Length | 6.00E-02 | 7.50E-02 | -4.80E-02 | 3.40E-02 |
| S.Pe.C.median. | MeanDepth | -7.20E-03 | 5.00E-02 | -3.80E-02 | 1.60E-02 |
| S.Pe.C.median. | Width | -5.10E-02 | -4.40E-02 | -1.70E-02 | -5.80E-03 |
| S.Pe.C.median. | SurfaceArea | 6.20E-02 | 1.00E-01 | -7.30E-02 | 5.50E-02 |
| S.Pe.C.sup. | Length | -1.10E-01 | -4.00E-02 | -3.60E-03 | 1.00E-02 |
| S.Pe.C.sup. | MeanDepth | 5.20E-04 | -2.90E-03 | -2.80E-02 | -8.40E-03 |
| S.Pe.C.sup. | Width | 3.50E-02 | 4.30E-03 | -9.00E-03 | 5.70E-03 |
| S.Pe.C.sup. | SurfaceArea | -1.20E-01 | -2.40E-02 | -1.30E-02 | 5.30E-03 |
| S.Po.C.sup. | Length | 8.50E-04 | -6.70E-03 | -7.00E-03 | -2.70E-02 |
| S.Po.C.sup. | MeanDepth | -3.30E-02 | 3.50E-02 | -1.80E-02 | 9.00E-03 |
| S.Po.C.sup. | Width | 2.60E-02 | 4.80E-02 | -3.20E-02 | -3.50E-02 |
| S.Po.C.sup. | SurfaceArea | -1.20E-02 | 8.00E-03 | -9.80E-03 | -1.40E-02 |
| S.R.inf. | Length | -4.40E-02 | -3.20E-03 | 8.30E-03 | 2.60E-02 |
| S.R.inf. | MeanDepth | -1.90E-02 | -7.40E-03 | -3.20E-03 | 1.30E-02 |
| S.R.inf. | Width | -3.20E-03 | -1.50E-02 | -2.90E-02 | -9.40E-03 |
| S.R.inf. | SurfaceArea | -8.40E-02 | -3.00E-02 | -8.40E-03 | 3.30E-02 |
| S.Rh. | Length | 6.50E-03 | -7.30E-03 | 6.30E-02 | 5.50E-02 |
| S.Rh. | MeanDepth | -6.60E-03 | 5.60E-03 | 1.70E-02 | 5.60E-03 |
| S.Rh. | Width | 2.70E-02 | -5.30E-02 | -3.60E-02 | -2.60E-02 |
| S.Rh. | SurfaceArea | 5.60E-03 | -1.60E-02 | 8.80E-02 | 8.00E-02 |
| S.s.P. | Length | -6.40E-02 | 1.20E-02 | 5.90E-02 | 4.10E-02 |
| S.s.P. | MeanDepth | -1.90E-02 | 9.40E-03 | -4.60E-03 | 8.10E-03 |
| S.s.P. | Width | 3.50E-02 | -8.80E-03 | -6.20E-02 | 2.00E-02 |
| S.s.P. | SurfaceArea | -9.20E-02 | 3.00E-02 | 8.40E-02 | 5.50E-02 |
| S.T.i.ant. | Length | 9.30E-02 | -4.70E-02 | -9.70E-02 | -9.60E-03 |
| S.T.i.ant. | MeanDepth | 3.90E-03 | -2.50E-02 | -8.10E-02 | -6.20E-03 |
| S.T.i.ant. | Width | 8.30E-03 | 1.10E-01 | 1.10E-01 | 6.60E-03 |
| S.T.i.ant. | SurfaceArea | 7.10E-02 | -8.10E-02 | -2.20E-01 | -1.10E-02 |
| S.T.i.post. | Length | -2.10E-02 | -2.70E-02 | -3.30E-02 | 9.00E-03 |
| S.T.i.post. | MeanDepth | 3.30E-03 | -3.80E-03 | -1.70E-02 | 5.10E-03 |
| S.T.i.post. | Width | 6.40E-03 | 1.40E-02 | 2.20E-02 | -1.90E-02 |

| | | | | | |
|---|---|---|---|---|---|
| S.T.i.post. | SurfaceArea | -2.60E-02 | -3.60E-02 | -5.40E-02 | 1.40E-02 |
| S.T.pol. | Length | -3.70E-02 | -1.10E-02 | -2.70E-02 | -7.50E-03 |
| S.T.pol. | MeanDepth | -1.90E-02 | 9.20E-03 | -2.70E-02 | 5.90E-03 |
| S.T.pol. | Width | 1.40E-02 | 2.80E-02 | -2.20E-02 | -9.20E-04 |
| S.T.pol. | SurfaceArea | -5.40E-02 | -7.30E-05 | -3.60E-02 | 1.30E-02 |
| S.T.s. | Length | -1.10E-03 | 8.80E-02 | 7.90E-02 | -7.90E-03 |
| S.T.s. | MeanDepth | 2.10E-02 | 1.20E-02 | -6.30E-02 | -1.30E-02 |
| S.T.s. | Width | -2.20E-02 | -1.70E-02 | 7.30E-02 | -9.40E-03 |
| S.T.s. | SurfaceArea | -6.70E-03 | 9.40E-02 | 2.60E-02 | -2.50E-02 |
| S.T.s.ter.asc.ant. | Length | 1.60E-03 | 2.00E-02 | -2.50E-02 | 2.90E-03 |
| S.T.s.ter.asc.ant. | MeanDepth | 7.40E-03 | -1.10E-02 | 1.10E-02 | -9.80E-03 |
| S.T.s.ter.asc.ant. | Width | 1.60E-02 | 7.80E-03 | -1.10E-02 | -2.00E-02 |
| S.T.s.ter.asc.ant. | SurfaceArea | 1.10E-02 | 1.60E-02 | -2.40E-02 | 2.20E-03 |
| S.T.s.ter.asc.post. | Length | -5.40E-02 | -1.20E-01 | 5.40E-02 | 3.30E-02 |
| S.T.s.ter.asc.post. | MeanDepth | 8.40E-03 | -5.00E-02 | -2.00E-03 | 5.20E-03 |
| S.T.s.ter.asc.post. | Width | 5.90E-03 | -5.60E-02 | -2.10E-02 | -2.70E-02 |
| S.T.s.ter.asc.post. | SurfaceArea | -7.00E-02 | -1.70E-01 | 4.50E-02 | 4.40E-02 |

**Table S10:** heritability eatimates for left hemisphere sulcal descriptors (yellow: bonferroni corrected for 123*4 comparisons) in QTIM.

| Sulci | Descriptor | h2 | pval | SE |
|---|---|---|---|---|
| F.C.L.a. | Length | 0.09 | 8.06E-02 | 0.07 |
| F.C.L.a. | MeanDepth | 0.03 | 3.17E-01 | 0.07 |
| F.C.L.a. | Width | 0.12 | 3.42E-02 | 0.06 |
| F.C.L.a. | SurfaceArea | 0.04 | 2.93E-01 | 0.07 |
| F.C.L.p. | Length | 0.09 | 8.39E-02 | 0.06 |
| F.C.L.p. | MeanDepth | 0.01 | 4.67E-01 | 0.06 |
| F.C.L.p. | Width | 0.32 | 2.00E-07 | 0.06 |
| F.C.L.p. | SurfaceArea | 0.10 | 6.35E-02 | 0.06 |
| F.C.L.r.ant. | Length | 0.03 | 3.15E-01 | 0.07 |
| F.C.L.r.ant. | MeanDepth | 0.01 | 4.57E-01 | 0.07 |
| F.C.L.r.ant. | Width | 0.15 | 1.24E-02 | 0.07 |
| F.C.L.r.ant. | SurfaceArea | 0.07 | 1.45E-01 | 0.07 |
| F.C.L.r.asc. | Length | - | - | - |
| F.C.L.r.asc. | MeanDepth | 0.08 | 9.96E-02 | 0.06 |
| F.C.L.r.asc. | Width | 0.15 | 1.37E-02 | 0.07 |
| F.C.L.r.asc. | SurfaceArea | 0.15 | 1.43E-02 | 0.07 |
| F.C.L.r.diag. | Length | 0.13 | 1.65E-01 | 0.14 |
| F.C.L.r.diag. | MeanDepth | 0.00 | 4.99E-01 | 0.12 |
| F.C.L.r.diag. | Width | 0.02 | 4.59E-01 | 0.15 |
| F.C.L.r.diag. | SurfaceArea | 0.07 | 2.82E-01 | 0.12 |
| F.C.L.r.retroC.tr. | Length | - | - | - |
| F.C.L.r.retroC.tr. | MeanDepth | 0.06 | 2.22E-01 | 0.08 |
| F.C.L.r.retroC.tr. | Width | 0.25 | 2.14E-04 | 0.07 |
| F.C.L.r.retroC.tr. | SurfaceArea | 0.01 | 4.30E-01 | 0.07 |
| F.C.L.r.sc.ant. | Length | 0.25 | 1.28E-01 | 0.21 |
| F.C.L.r.sc.ant. | MeanDepth | - | - | - |
| F.C.L.r.sc.ant. | Width | - | - | - |
| F.C.L.r.sc.ant. | SurfaceArea | 0.34 | 9.80E-02 | 0.24 |
| F.C.L.r.sc.post. | Length | 0.04 | 3.25E-01 | 0.08 |
| F.C.L.r.sc.post. | MeanDepth | 0.28 | 2.04E-01 | 0.33 |
| F.C.L.r.sc.post. | Width | 0.69 | 3.05E-02 | 0.20 |
| F.C.L.r.sc.post. | SurfaceArea | 0.12 | 6.01E-02 | 0.08 |
| F.C.M.ant. | Length | 0.20 | 1.33E-03 | 0.06 |
| F.C.M.ant. | MeanDepth | - | - | - |
| F.C.M.ant. | Width | 0.13 | 3.24E-02 | 0.07 |
| F.C.M.ant. | SurfaceArea | - | - | - |
| F.C.M.post. | Length | 0.31 | 9.00E-07 | 0.06 |
| F.C.M.post. | MeanDepth | 0.11 | 3.71E-02 | 0.06 |
| F.C.M.post. | Width | 0.25 | 3.92E-05 | 0.06 |
| F.C.M.post. | SurfaceArea | 0.15 | 6.59E-03 | 0.06 |
| F.Cal.ant.-Sc.Cal. | Length | 0.15 | 2.16E-02 | 0.07 |
| F.Cal.ant.-Sc.Cal. | MeanDepth | 0.07 | 1.38E-01 | 0.07 |
| F.Cal.ant.-Sc.Cal. | Width | 0.14 | 2.17E-02 | 0.07 |

| | | | | |
|---|---|---|---|---|
| F.Cal.ant.-Sc.Cal. | SurfaceArea | 0.24 | 2.90E-04 | 0.07 |
| F.Coll. | Length | 0.22 | 6.65E-04 | 0.06 |
| F.Coll. | MeanDepth | 0.02 | 4.10E-01 | 0.07 |
| F.Coll. | Width | 0.11 | 4.25E-02 | 0.06 |
| F.Coll. | SurfaceArea | 0.15 | 2.12E-02 | 0.07 |
| F.I.P. | Length | 0.15 | 1.10E-02 | 0.07 |
| F.I.P. | MeanDepth | - | - | - |
| F.I.P. | Width | 0.43 | 5.49E-11 | 0.06 |
| F.I.P. | SurfaceArea | 0.00 | 4.99E-01 | 0.07 |
| F.I.P.Po.C.inf. | Length | 0.13 | 2.21E-02 | 0.07 |
| F.I.P.Po.C.inf. | MeanDepth | - | - | - |
| F.I.P.Po.C.inf. | Width | 0.25 | 8.70E-05 | 0.06 |
| F.I.P.Po.C.inf. | SurfaceArea | 0.07 | 1.39E-01 | 0.07 |
| F.I.P.r.int.1 | Length | - | - | - |
| F.I.P.r.int.1 | MeanDepth | - | - | - |
| F.I.P.r.int.1 | Width | 0.10 | 5.70E-02 | 0.06 |
| F.I.P.r.int.1 | SurfaceArea | 0.02 | 3.98E-01 | 0.06 |
| F.I.P.r.int.2 | Length | 0.18 | 5.36E-03 | 0.07 |
| F.I.P.r.int.2 | MeanDepth | - | - | - |
| F.I.P.r.int.2 | Width | 0.09 | 9.54E-02 | 0.07 |
| F.I.P.r.int.2 | SurfaceArea | 0.05 | 2.46E-01 | 0.07 |
| F.P.O. | Length | 0.28 | 1.01E-05 | 0.06 |
| F.P.O. | MeanDepth | 0.10 | 6.51E-02 | 0.07 |
| F.P.O. | Width | 0.15 | 1.12E-02 | 0.06 |
| F.P.O. | SurfaceArea | 0.37 | 1.24E-08 | 0.06 |
| INSULA | Length | 0.04 | 2.71E-01 | 0.06 |
| INSULA | MeanDepth | 0.11 | 2.46E-01 | 0.16 |
| INSULA | Width | 0.55 | 2.85E-03 | 0.15 |
| INSULA | SurfaceArea | 0.21 | 7.80E-04 | 0.06 |
| OCCIPITAL | Length | 0.13 | 3.77E-02 | 0.07 |
| OCCIPITAL | MeanDepth | 0.20 | 6.12E-04 | 0.06 |
| OCCIPITAL | Width | 0.26 | 1.17E-04 | 0.07 |
| OCCIPITAL | SurfaceArea | 0.26 | 4.87E-05 | 0.06 |
| S.C. | Length | 0.18 | 2.02E-03 | 0.06 |
| S.C. | MeanDepth | - | - | - |
| S.C. | Width | 0.41 | 3.72E-11 | 0.06 |
| S.C. | SurfaceArea | 0.07 | 1.54E-01 | 0.07 |
| S.C.LPC. | Length | 0.09 | 1.69E-01 | 0.10 |
| S.C.LPC. | MeanDepth | 0.13 | 7.36E-02 | 0.09 |
| S.C.LPC. | Width | 0.17 | 3.47E-02 | 0.09 |
| S.C.LPC. | SurfaceArea | 0.13 | 7.78E-02 | 0.09 |
| S.C.sylvian. | Length | 0.29 | 2.63E-04 | 0.08 |
| S.C.sylvian. | MeanDepth | 0.15 | 5.49E-02 | 0.09 |
| S.C.sylvian. | Width | 0.22 | 9.33E-03 | 0.09 |
| S.C.sylvian. | SurfaceArea | 0.23 | 3.65E-03 | 0.08 |
| S.Call. | Length | 0.25 | 1.98E-04 | 0.07 |

| | | | | |
|---|---|---|---|---|
| S.Call. | MeanDepth | 0.10 | 6.35E-02 | 0.07 |
| S.Call. | Width | 0.23 | 5.34E-04 | 0.07 |
| S.Call. | SurfaceArea | 0.13 | 2.61E-02 | 0.07 |
| S.Cu. | Length | 0.11 | 6.78E-02 | 0.07 |
| S.Cu. | MeanDepth | - | - | - |
| S.Cu. | Width | 0.17 | 6.94E-03 | 0.07 |
| S.Cu. | SurfaceArea | - | - | - |
| S.F.inf. | Length | 0.18 | 2.28E-03 | 0.06 |
| S.F.inf. | MeanDepth | 0.02 | 3.55E-01 | 0.06 |
| S.F.inf. | Width | 0.25 | 7.90E-05 | 0.06 |
| S.F.inf. | SurfaceArea | 0.06 | 1.97E-01 | 0.07 |
| S.F.inf.ant. | Length | - | - | - |
| S.F.inf.ant. | MeanDepth | - | - | - |
| S.F.inf.ant. | Width | - | - | - |
| S.F.inf.ant. | SurfaceArea | - | - | - |
| S.F.int. | Length | 0.15 | 1.39E-02 | 0.07 |
| S.F.int. | MeanDepth | 0.07 | 1.60E-01 | 0.07 |
| S.F.int. | Width | 0.06 | 2.10E-01 | 0.07 |
| S.F.int. | SurfaceArea | 0.08 | 1.35E-01 | 0.07 |
| S.F.inter. | Length | 0.11 | 5.02E-02 | 0.07 |
| S.F.inter. | MeanDepth | 0.03 | 3.25E-01 | 0.07 |
| S.F.inter. | Width | 0.31 | 1.00E-06 | 0.06 |
| S.F.inter. | SurfaceArea | 0.04 | 2.83E-01 | 0.07 |
| S.F.marginal. | Length | 0.03 | 3.06E-01 | 0.07 |
| S.F.marginal. | MeanDepth | 0.19 | 3.31E-03 | 0.07 |
| S.F.marginal. | Width | 0.04 | 2.72E-01 | 0.07 |
| S.F.marginal. | SurfaceArea | 0.19 | 2.35E-03 | 0.07 |
| S.F.median. | Length | 0.13 | 2.72E-02 | 0.07 |
| S.F.median. | MeanDepth | 0.18 | 2.92E-03 | 0.06 |
| S.F.median. | Width | 0.16 | 1.15E-02 | 0.07 |
| S.F.median. | SurfaceArea | 0.17 | 4.04E-03 | 0.07 |
| S.F.orbitaire. | Length | 0.06 | 1.84E-01 | 0.06 |
| S.F.orbitaire. | MeanDepth | - | - | - |
| S.F.orbitaire. | Width | - | - | - |
| S.F.orbitaire. | SurfaceArea | - | - | - |
| S.F.polaire.tr. | Length | 0.07 | 1.51E-01 | 0.07 |
| S.F.polaire.tr. | MeanDepth | 0.21 | 1.60E-03 | 0.07 |
| S.F.polaire.tr. | Width | 0.14 | 2.29E-02 | 0.07 |
| S.F.polaire.tr. | SurfaceArea | 0.18 | 6.01E-03 | 0.07 |
| S.F.sup. | Length | 0.15 | 5.58E-03 | 0.06 |
| S.F.sup. | MeanDepth | - | - | - |
| S.F.sup. | Width | 0.43 | 1.50E-11 | 0.06 |
| S.F.sup. | SurfaceArea | 0.03 | 3.41E-01 | 0.07 |
| S.GSM. | Length | 0.13 | 6.00E-02 | 0.08 |
| S.GSM. | MeanDepth | 0.02 | 4.31E-01 | 0.09 |
| S.GSM. | Width | 0.15 | 4.46E-02 | 0.09 |

| | | | | |
|---|---|---|---|---|
| S.GSM. | SurfaceArea | 0.03 | 3.45E-01 | 0.09 |
| S.Li.ant. | Length | 0.14 | 1.54E-02 | 0.07 |
| S.Li.ant. | MeanDepth | - | - | - |
| S.Li.ant. | Width | 0.12 | 4.57E-02 | 0.07 |
| S.Li.ant. | SurfaceArea | 0.00 | 4.82E-01 | 0.07 |
| S.Li.post. | Length | 0.06 | 1.86E-01 | 0.07 |
| S.Li.post. | MeanDepth | 0.17 | 7.06E-03 | 0.07 |
| S.Li.post. | Width | 0.06 | 1.70E-01 | 0.07 |
| S.Li.post. | SurfaceArea | 0.16 | 8.85E-03 | 0.07 |
| S.O.p. | Length | 0.16 | 1.83E-02 | 0.07 |
| S.O.p. | MeanDepth | 0.05 | 2.42E-01 | 0.08 |
| S.O.p. | Width | 0.11 | 9.58E-02 | 0.08 |
| S.O.p. | SurfaceArea | 0.12 | 5.68E-02 | 0.07 |
| S.O.T.lat.ant. | Length | 0.08 | 1.13E-01 | 0.06 |
| S.O.T.lat.ant. | MeanDepth | 0.06 | 1.81E-01 | 0.06 |
| S.O.T.lat.ant. | Width | 0.07 | 1.28E-01 | 0.06 |
| S.O.T.lat.ant. | SurfaceArea | 0.10 | 5.89E-02 | 0.06 |
| S.O.T.lat.int. | Length | 0.08 | 1.40E-01 | 0.07 |
| S.O.T.lat.int. | MeanDepth | 0.06 | 1.91E-01 | 0.07 |
| S.O.T.lat.int. | Width | 0.16 | 1.14E-02 | 0.07 |
| S.O.T.lat.int. | SurfaceArea | 0.08 | 1.34E-01 | 0.07 |
| S.O.T.lat.med. | Length | 0.06 | 1.95E-01 | 0.07 |
| S.O.T.lat.med. | MeanDepth | 0.02 | 3.79E-01 | 0.07 |
| S.O.T.lat.med. | Width | 0.05 | 2.46E-01 | 0.07 |
| S.O.T.lat.med. | SurfaceArea | 0.00 | 4.92E-01 | 0.07 |
| S.O.T.lat.post. | Length | 0.17 | 6.73E-03 | 0.07 |
| S.O.T.lat.post. | MeanDepth | 0.04 | 2.50E-01 | 0.06 |
| S.O.T.lat.post. | Width | 0.06 | 1.69E-01 | 0.07 |
| S.O.T.lat.post. | SurfaceArea | 0.08 | 1.04E-01 | 0.06 |
| S.Olf. | Length | 0.16 | 8.58E-03 | 0.07 |
| S.Olf. | MeanDepth | 0.05 | 2.21E-01 | 0.07 |
| S.Olf. | Width | 0.04 | 2.94E-01 | 0.07 |
| S.Olf. | SurfaceArea | 0.09 | 8.99E-02 | 0.07 |
| S.Or. | Length | 0.14 | 1.64E-02 | 0.06 |
| S.Or. | MeanDepth | 0.05 | 2.21E-01 | 0.06 |
| S.Or. | Width | 0.17 | 5.00E-03 | 0.07 |
| S.Or. | SurfaceArea | 0.07 | 1.48E-01 | 0.07 |
| S.p.C. | Length | 0.01 | 4.47E-01 | 0.08 |
| S.p.C. | MeanDepth | 0.16 | 9.55E-03 | 0.07 |
| S.p.C. | Width | 0.09 | 1.30E-01 | 0.08 |
| S.p.C. | SurfaceArea | 0.11 | 7.06E-02 | 0.07 |
| S.Pa.int. | Length | 0.20 | 1.55E-03 | 0.07 |
| S.Pa.int. | MeanDepth | 0.14 | 2.65E-02 | 0.07 |
| S.Pa.int. | Width | 0.32 | 2.90E-06 | 0.07 |
| S.Pa.int. | SurfaceArea | 0.16 | 1.13E-02 | 0.07 |
| S.Pa.sup. | Length | 0.08 | 1.47E-01 | 0.08 |

| | | | | |
|---|---|---|---|---|
| S.Pa.sup. | MeanDepth | 0.03 | 3.52E-01 | 0.08 |
| S.Pa.sup. | Width | 0.24 | 1.23E-03 | 0.08 |
| S.Pa.sup. | SurfaceArea | 0.03 | 3.44E-01 | 0.08 |
| S.Pa.t. | Length | 0.06 | 2.52E-01 | 0.09 |
| S.Pa.t. | MeanDepth | 0.03 | 3.83E-01 | 0.12 |
| S.Pa.t. | Width | 0.09 | 2.01E-01 | 0.11 |
| S.Pa.t. | SurfaceArea | 0.10 | 1.15E-01 | 0.08 |
| S.Pe.C.inf. | Length | 0.08 | 1.30E-01 | 0.07 |
| S.Pe.C.inf. | MeanDepth | 0.02 | 3.75E-01 | 0.08 |
| S.Pe.C.inf. | Width | 0.24 | 5.10E-04 | 0.07 |
| S.Pe.C.inf. | SurfaceArea | 0.04 | 2.89E-01 | 0.07 |
| S.Pe.C.inter. | Length | 0.11 | 7.45E-02 | 0.07 |
| S.Pe.C.inter. | MeanDepth | 0.15 | 1.24E-02 | 0.07 |
| S.Pe.C.inter. | Width | 0.12 | 3.54E-02 | 0.07 |
| S.Pe.C.inter. | SurfaceArea | 0.18 | 5.04E-03 | 0.07 |
| S.Pe.C.marginal. | Length | 0.07 | 2.03E-01 | 0.09 |
| S.Pe.C.marginal. | MeanDepth | 0.06 | 2.09E-01 | 0.08 |
| S.Pe.C.marginal. | Width | 0.18 | 1.38E-02 | 0.08 |
| S.Pe.C.marginal. | SurfaceArea | 0.05 | 2.66E-01 | 0.07 |
| S.Pe.C.median. | Length | 0.12 | 5.27E-02 | 0.07 |
| S.Pe.C.median. | MeanDepth | 0.20 | 3.50E-03 | 0.07 |
| S.Pe.C.median. | Width | 0.08 | 1.39E-01 | 0.07 |
| S.Pe.C.median. | SurfaceArea | 0.15 | 2.06E-02 | 0.07 |
| S.Pe.C.sup. | Length | 0.13 | 4.35E-02 | 0.07 |
| S.Pe.C.sup. | MeanDepth | 0.04 | 3.09E-01 | 0.08 |
| S.Pe.C.sup. | Width | 0.14 | 2.42E-02 | 0.07 |
| S.Pe.C.sup. | SurfaceArea | 0.05 | 2.59E-01 | 0.07 |
| S.Po.C.sup. | Length | 0.08 | 1.46E-01 | 0.07 |
| S.Po.C.sup. | MeanDepth | - | - | - |
| S.Po.C.sup. | Width | 0.20 | 2.03E-03 | 0.07 |
| S.Po.C.sup. | SurfaceArea | - | - | - |
| S.R.inf. | Length | 0.05 | 2.56E-01 | 0.08 |
| S.R.inf. | MeanDepth | 0.15 | 1.39E-02 | 0.07 |
| S.R.inf. | Width | 0.13 | 5.16E-02 | 0.08 |
| S.R.inf. | SurfaceArea | 0.13 | 2.65E-02 | 0.07 |
| S.Rh. | Length | 0.02 | 3.64E-01 | 0.07 |
| S.Rh. | MeanDepth | 0.15 | 1.00E-02 | 0.07 |
| S.Rh. | Width | - | - | - |
| S.Rh. | SurfaceArea | 0.12 | 3.27E-02 | 0.07 |
| S.s.P. | Length | 0.28 | 4.45E-05 | 0.07 |
| S.s.P. | MeanDepth | 0.09 | 8.08E-02 | 0.07 |
| S.s.P. | Width | 0.15 | 1.66E-02 | 0.07 |
| S.s.P. | SurfaceArea | 0.16 | 9.09E-03 | 0.07 |
| S.T.i.ant. | Length | 0.07 | 1.55E-01 | 0.07 |
| S.T.i.ant. | MeanDepth | 0.08 | 1.17E-01 | 0.07 |
| S.T.i.ant. | Width | 0.11 | 5.71E-02 | 0.07 |

| | | | | |
|---|---|---|---|---|
| S.T.i.ant. | SurfaceArea | 0.07 | 1.57E-01 | 0.07 |
| S.T.i.post. | Length | 0.16 | 9.24E-03 | 0.07 |
| S.T.i.post. | MeanDepth | - | - | - |
| S.T.i.post. | Width | 0.22 | 8.16E-04 | 0.07 |
| S.T.i.post. | SurfaceArea | 0.03 | 3.57E-01 | 0.07 |
| S.T.pol. | Length | 0.00 | 4.83E-01 | 0.07 |
| S.T.pol. | MeanDepth | 0.04 | 2.79E-01 | 0.08 |
| S.T.pol. | Width | 0.17 | 9.28E-03 | 0.07 |
| S.T.pol. | SurfaceArea | 0.12 | 5.77E-02 | 0.08 |
| S.T.s. | Length | 0.27 | 1.96E-05 | 0.06 |
| S.T.s. | MeanDepth | - | - | - |
| S.T.s. | Width | 0.29 | 6.00E-06 | 0.06 |
| S.T.s. | SurfaceArea | 0.03 | 3.16E-01 | 0.07 |
| S.T.s.ter.asc.ant. | Length | - | - | - |
| S.T.s.ter.asc.ant. | MeanDepth | - | - | - |
| S.T.s.ter.asc.ant. | Width | 0.17 | 7.00E-03 | 0.07 |
| S.T.s.ter.asc.ant. | SurfaceArea | - | - | - |
| S.T.s.ter.asc.post. | Length | 0.10 | 6.66E-02 | 0.07 |
| S.T.s.ter.asc.post. | MeanDepth | 0.06 | 1.94E-01 | 0.07 |
| S.T.s.ter.asc.post. | Width | 0.18 | 7.10E-03 | 0.07 |
| S.T.s.ter.asc.post. | SurfaceArea | 0.11 | 4.78E-02 | 0.07 |
| Globals | Length | 0.25 | 3.46E-05 | 0.06 |
| Globals | MeanDepth | 0.24 | 4.44E-04 | 0.07 |
| Globals | Width | 0.33 | 4.00E-08 | 0.06 |
| Globals | SurfaceArea | 0.26 | 1.33E-04 | 0.07 |

**Table S11:** heritability eatimates for right hemisphere sulcal descriptors (yellow: bonferroni corrected for 123*4 comparisons) in QTIM.

| Sulci | Descriptor | h2 | pval | SE |
|---|---|---|---|---|
| F.C.L.a. | Length | 0.15 | 1.29E-02 | 0.06 |
| F.C.L.a. | MeanDepth | 0.02 | 3.83E-01 | 0.07 |
| F.C.L.a. | Width | 0.13 | 3.48E-02 | 0.07 |
| F.C.L.a. | SurfaceArea | 0.09 | 1.04E-01 | 0.07 |
| F.C.L.p. | Length | 0.20 | 2.34E-03 | 0.07 |
| F.C.L.p. | MeanDepth | 0.10 | 4.80E-02 | 0.06 |
| F.C.L.p. | Width | 0.24 | 2.22E-05 | 0.06 |
| F.C.L.p. | SurfaceArea | 0.26 | 3.19E-05 | 0.06 |
| F.C.L.r.ant. | Length | - | - | - |
| F.C.L.r.ant. | MeanDepth | 0.12 | 6.86E-02 | 0.08 |
| F.C.L.r.ant. | Width | 0.15 | 3.95E-02 | 0.09 |
| F.C.L.r.ant. | SurfaceArea | 0.19 | 4.13E-03 | 0.07 |
| F.C.L.r.asc. | Length | 0.16 | 9.69E-03 | 0.07 |
| F.C.L.r.asc. | MeanDepth | - | - | - |
| F.C.L.r.asc. | Width | 0.06 | 2.07E-01 | 0.07 |
| F.C.L.r.asc. | SurfaceArea | 0.10 | 7.56E-02 | 0.07 |
| F.C.L.r.diag. | Length | 0.09 | 2.21E-01 | 0.11 |
| F.C.L.r.diag. | MeanDepth | 0.33 | 5.29E-03 | 0.12 |
| F.C.L.r.diag. | Width | - | - | - |
| F.C.L.r.diag. | SurfaceArea | 0.18 | 8.22E-02 | 0.12 |
| F.C.L.r.retroC.tr. | Length | 0.18 | 4.88E-03 | 0.07 |
| F.C.L.r.retroC.tr. | MeanDepth | 0.12 | 3.96E-02 | 0.07 |
| F.C.L.r.retroC.tr. | Width | 0.12 | 5.64E-02 | 0.07 |
| F.C.L.r.retroC.tr. | SurfaceArea | 0.17 | 7.51E-03 | 0.07 |
| F.C.L.r.sc.ant. | Length | 0.61 | 2.05E-02 | 0.21 |
| F.C.L.r.sc.ant. | MeanDepth | - | - | - |
| F.C.L.r.sc.ant. | Width | - | - | - |
| F.C.L.r.sc.ant. | SurfaceArea | 0.48 | 2.51E-02 | 0.20 |
| F.C.L.r.sc.post. | Length | 0.04 | 3.00E-01 | 0.08 |
| F.C.L.r.sc.post. | MeanDepth | 0.91 | 1.14E-04 | 0.06 |
| F.C.L.r.sc.post. | Width | 0.30 | 2.61E-01 | 0.46 |
| F.C.L.r.sc.post. | SurfaceArea | 0.19 | 4.43E-03 | 0.07 |
| F.C.M.ant. | Length | 0.22 | 3.57E-04 | 0.06 |
| F.C.M.ant. | MeanDepth | 0.10 | 7.71E-02 | 0.07 |
| F.C.M.ant. | Width | 0.27 | 1.93E-05 | 0.06 |
| F.C.M.ant. | SurfaceArea | 0.19 | 1.88E-03 | 0.06 |
| F.C.M.post. | Length | 0.18 | 5.20E-03 | 0.07 |
| F.C.M.post. | MeanDepth | 0.08 | 1.19E-01 | 0.06 |
| F.C.M.post. | Width | 0.33 | 7.00E-07 | 0.06 |
| F.C.M.post. | SurfaceArea | 0.16 | 5.61E-03 | 0.06 |
| F.Cal.ant.-Sc.Cal. | Length | 0.12 | 3.42E-02 | 0.06 |
| F.Cal.ant.-Sc.Cal. | MeanDepth | 0.19 | 1.77E-03 | 0.06 |
| F.Cal.ant.-Sc.Cal. | Width | 0.10 | 6.44E-02 | 0.07 |

| | | | | |
|---|---|---|---|---|
| F.Cal.ant.-Sc.Cal. | SurfaceArea | 0.32 | 5.00E-07 | 0.06 |
| F.Coll. | Length | 0.28 | 3.30E-06 | 0.06 |
| F.Coll. | MeanDepth | 0.21 | 6.46E-04 | 0.06 |
| F.Coll. | Width | 0.22 | 3.77E-04 | 0.06 |
| F.Coll. | SurfaceArea | 0.22 | 3.89E-04 | 0.06 |
| F.I.P. | Length | 0.23 | 1.96E-04 | 0.06 |
| F.I.P. | MeanDepth | 0.20 | 1.39E-03 | 0.06 |
| F.I.P. | Width | 0.44 | 1.16E-13 | 0.05 |
| F.I.P. | SurfaceArea | 0.29 | 7.50E-06 | 0.06 |
| F.I.P.Po.C.inf. | Length | 0.18 | 2.23E-03 | 0.06 |
| F.I.P.Po.C.inf. | MeanDepth | 0.13 | 2.47E-02 | 0.06 |
| F.I.P.Po.C.inf. | Width | 0.27 | 1.56E-05 | 0.06 |
| F.I.P.Po.C.inf. | SurfaceArea | 0.14 | 1.42E-02 | 0.07 |
| F.I.P.r.int.1 | Length | - | - | - |
| F.I.P.r.int.1 | MeanDepth | 0.17 | 8.13E-03 | 0.07 |
| F.I.P.r.int.1 | Width | 0.07 | 1.98E-01 | 0.08 |
| F.I.P.r.int.1 | SurfaceArea | 0.11 | 5.88E-02 | 0.07 |
| F.I.P.r.int.2 | Length | - | - | - |
| F.I.P.r.int.2 | MeanDepth | - | - | - |
| F.I.P.r.int.2 | Width | 0.05 | 3.18E-01 | 0.10 |
| F.I.P.r.int.2 | SurfaceArea | 0.07 | 2.23E-01 | 0.09 |
| F.P.O. | Length | 0.10 | 5.19E-02 | 0.06 |
| F.P.O. | MeanDepth | 0.15 | 1.37E-02 | 0.06 |
| F.P.O. | Width | 0.25 | 9.55E-05 | 0.06 |
| F.P.O. | SurfaceArea | 0.23 | 2.39E-04 | 0.06 |
| INSULA | Length | 0.11 | 4.19E-02 | 0.06 |
| INSULA | MeanDepth | 0.07 | 3.14E-01 | 0.15 |
| INSULA | Width | 0.18 | 1.28E-01 | 0.15 |
| INSULA | SurfaceArea | 0.25 | 1.34E-04 | 0.07 |
| OCCIPITAL | Length | 0.22 | 1.20E-03 | 0.07 |
| OCCIPITAL | MeanDepth | 0.32 | 5.00E-07 | 0.06 |
| OCCIPITAL | Width | 0.25 | 1.34E-04 | 0.07 |
| OCCIPITAL | SurfaceArea | 0.32 | 6.00E-07 | 0.06 |
| S.C. | Length | 0.25 | 4.00E-05 | 0.06 |
| S.C. | MeanDepth | 0.22 | 5.18E-04 | 0.07 |
| S.C. | Width | 0.44 | 8.73E-13 | 0.05 |
| S.C. | SurfaceArea | 0.27 | 2.92E-05 | 0.06 |
| S.C.LPC. | Length | - | - | - |
| S.C.LPC. | MeanDepth | 0.03 | 4.52E-01 | 0.23 |
| S.C.LPC. | Width | 0.49 | 1.20E-02 | 0.17 |
| S.C.LPC. | SurfaceArea | 0.15 | 2.39E-01 | 0.21 |
| S.C.sylvian. | Length | 0.19 | 1.15E-02 | 0.08 |
| S.C.sylvian. | MeanDepth | 0.23 | 4.30E-03 | 0.08 |
| S.C.sylvian. | Width | 0.27 | 1.67E-03 | 0.09 |
| S.C.sylvian. | SurfaceArea | 0.23 | 2.12E-03 | 0.08 |
| S.Call. | Length | 0.37 | 4.51E-09 | 0.06 |

| | | | | |
|---|---|---|---|---|
| S.Call. | MeanDepth | 0.35 | 1.00E-07 | 0.06 |
| S.Call. | Width | 0.30 | 2.10E-06 | 0.06 |
| S.Call. | SurfaceArea | 0.40 | 1.50E-09 | 0.06 |
| S.Cu. | Length | 0.13 | 2.50E-02 | 0.07 |
| S.Cu. | MeanDepth | 0.04 | 2.98E-01 | 0.07 |
| S.Cu. | Width | 0.09 | 1.05E-01 | 0.07 |
| S.Cu. | SurfaceArea | - | - | - |
| S.F.inf. | Length | 0.34 | 1.00E-07 | 0.06 |
| S.F.inf. | MeanDepth | - | - | - |
| S.F.inf. | Width | 0.25 | 8.11E-05 | 0.06 |
| S.F.inf. | SurfaceArea | 0.15 | 1.39E-02 | 0.07 |
| S.F.inf.ant. | Length | 0.17 | 6.55E-03 | 0.07 |
| S.F.inf.ant. | MeanDepth | 0.21 | 2.52E-04 | 0.06 |
| S.F.inf.ant. | Width | 0.17 | 4.11E-03 | 0.06 |
| S.F.inf.ant. | SurfaceArea | 0.19 | 1.00E-03 | 0.06 |
| S.F.int. | Length | 0.17 | 6.04E-03 | 0.07 |
| S.F.int. | MeanDepth | 0.34 | 2.00E-07 | 0.06 |
| S.F.int. | Width | 0.33 | 1.00E-07 | 0.06 |
| S.F.int. | SurfaceArea | 0.35 | 2.00E-07 | 0.06 |
| S.F.inter. | Length | 0.10 | 5.37E-02 | 0.06 |
| S.F.inter. | MeanDepth | 0.14 | 1.72E-02 | 0.06 |
| S.F.inter. | Width | 0.42 | 9.76E-12 | 0.05 |
| S.F.inter. | SurfaceArea | 0.11 | 3.69E-02 | 0.06 |
| S.F.margi-l. | Length | 0.13 | 2.61E-02 | 0.07 |
| S.F.margi-l. | MeanDepth | 0.24 | 2.45E-04 | 0.07 |
| S.F.margi-l. | Width | 0.12 | 5.44E-02 | 0.07 |
| S.F.margi-l. | SurfaceArea | 0.22 | 5.40E-04 | 0.07 |
| S.F.median. | Length | 0.21 | 4.46E-04 | 0.06 |
| S.F.median. | MeanDepth | 0.16 | 3.43E-03 | 0.06 |
| S.F.median. | Width | 0.27 | 5.30E-06 | 0.06 |
| S.F.median. | SurfaceArea | 0.20 | 4.98E-04 | 0.06 |
| S.F.orbitaire. | Length | - | - | - |
| S.F.orbitaire. | MeanDepth | 0.16 | 1.35E-02 | 0.07 |
| S.F.orbitaire. | Width | 0.14 | 3.46E-02 | 0.08 |
| S.F.orbitaire. | SurfaceArea | 0.13 | 4.40E-02 | 0.07 |
| S.F.polaire.tr. | Length | 0.15 | 1.30E-02 | 0.07 |
| S.F.polaire.tr. | MeanDepth | - | - | - |
| S.F.polaire.tr. | Width | 0.19 | 1.70E-03 | 0.06 |
| S.F.polaire.tr. | SurfaceArea | - | - | - |
| S.F.sup. | Length | 0.17 | 3.47E-03 | 0.06 |
| S.F.sup. | MeanDepth | - | - | - |
| S.F.sup. | Width | 0.33 | 1.84E-08 | 0.06 |
| S.F.sup. | SurfaceArea | 0.16 | 6.62E-03 | 0.06 |
| S.Li.ant. | Length | 0.09 | 1.13E-01 | 0.08 |
| S.Li.ant. | MeanDepth | 0.13 | 2.97E-02 | 0.07 |
| S.Li.ant. | Width | 0.07 | 1.56E-01 | 0.07 |

| | | | | |
|---|---|---|---|---|
| S.Li.ant. | SurfaceArea | 0.09 | 9.11E-02 | 0.07 |
| S.Li.post. | Length | 0.07 | 1.54E-01 | 0.06 |
| S.Li.post. | MeanDepth | 0.14 | 1.90E-02 | 0.07 |
| S.Li.post. | Width | 0.25 | 1.62E-04 | 0.07 |
| S.Li.post. | SurfaceArea | 0.14 | 2.03E-02 | 0.07 |
| S.O.p. | Length | 0.13 | 2.46E-02 | 0.07 |
| S.O.p. | MeanDepth | 0.07 | 1.57E-01 | 0.07 |
| S.O.p. | Width | 0.21 | 1.15E-03 | 0.07 |
| S.O.p. | SurfaceArea | 0.11 | 5.05E-02 | 0.07 |
| S.O.T.lat.ant. | Length | 0.14 | 2.56E-02 | 0.07 |
| S.O.T.lat.ant. | MeanDepth | 0.11 | 4.57E-02 | 0.06 |
| S.O.T.lat.ant. | Width | 0.37 | 1.46E-09 | 0.05 |
| S.O.T.lat.ant. | SurfaceArea | 0.12 | 3.94E-02 | 0.07 |
| S.O.T.lat.int. | Length | 0.11 | 7.92E-02 | 0.08 |
| S.O.T.lat.int. | MeanDepth | 0.21 | 1.62E-03 | 0.07 |
| S.O.T.lat.int. | Width | 0.23 | 1.64E-03 | 0.07 |
| S.O.T.lat.int. | SurfaceArea | 0.21 | 1.59E-03 | 0.07 |
| S.O.T.lat.med. | Length | 0.23 | 8.00E-04 | 0.07 |
| S.O.T.lat.med. | MeanDepth | - | - | - |
| S.O.T.lat.med. | Width | 0.25 | 4.65E-04 | 0.07 |
| S.O.T.lat.med. | SurfaceArea | 0.12 | 5.70E-02 | 0.08 |
| S.O.T.lat.post. | Length | 0.09 | 9.50E-02 | 0.07 |
| S.O.T.lat.post. | MeanDepth | 0.04 | 2.90E-01 | 0.07 |
| S.O.T.lat.post. | Width | 0.08 | 1.13E-01 | 0.07 |
| S.O.T.lat.post. | SurfaceArea | 0.09 | 1.14E-01 | 0.07 |
| S.Olf. | Length | 0.23 | 1.13E-04 | 0.06 |
| S.Olf. | MeanDepth | 0.20 | 1.79E-03 | 0.07 |
| S.Olf. | Width | 0.25 | 5.95E-05 | 0.06 |
| S.Olf. | SurfaceArea | 0.23 | 2.47E-04 | 0.06 |
| S.Or. | Length | 0.14 | 1.90E-02 | 0.07 |
| S.Or. | MeanDepth | 0.23 | 2.40E-04 | 0.06 |
| S.Or. | Width | 0.22 | 2.64E-04 | 0.06 |
| S.Or. | SurfaceArea | 0.25 | 8.35E-05 | 0.06 |
| S.p.C. | Length | 0.19 | 5.80E-03 | 0.07 |
| S.p.C. | MeanDepth | 0.15 | 2.90E-02 | 0.08 |
| S.p.C. | Width | 0.19 | 6.23E-03 | 0.07 |
| S.p.C. | SurfaceArea | 0.19 | 9.33E-03 | 0.08 |
| S.Pa.int. | Length | 0.05 | 2.33E-01 | 0.07 |
| S.Pa.int. | MeanDepth | 0.06 | 1.88E-01 | 0.07 |
| S.Pa.int. | Width | 0.15 | 1.36E-02 | 0.07 |
| S.Pa.int. | SurfaceArea | 0.07 | 1.54E-01 | 0.06 |
| S.Pa.sup. | Length | 0.04 | 2.82E-01 | 0.07 |
| S.Pa.sup. | MeanDepth | 0.05 | 2.11E-01 | 0.07 |
| S.Pa.sup. | Width | 0.13 | 2.98E-02 | 0.07 |
| S.Pa.sup. | SurfaceArea | 0.02 | 3.88E-01 | 0.07 |
| S.Pa.t. | Length | 0.13 | 4.01E-02 | 0.07 |

| | | | | |
|---|---|---|---|---|
| S.Pa.t. | MeanDepth | 0.02 | 4.19E-01 | 0.09 |
| S.Pa.t. | Width | 0.27 | 6.25E-04 | 0.08 |
| S.Pa.t. | SurfaceArea | 0.12 | 5.06E-02 | 0.07 |
| S.Pe.C.inf. | Length | 0.07 | 1.57E-01 | 0.07 |
| S.Pe.C.inf. | MeanDepth | 0.22 | 9.05E-04 | 0.07 |
| S.Pe.C.inf. | Width | 0.28 | 4.39E-05 | 0.07 |
| S.Pe.C.inf. | SurfaceArea | 0.21 | 1.27E-03 | 0.07 |
| S.Pe.C.inter. | Length | 0.09 | 7.91E-02 | 0.07 |
| S.Pe.C.inter. | MeanDepth | 0.27 | 5.86E-05 | 0.07 |
| S.Pe.C.inter. | Width | 0.19 | 1.22E-03 | 0.06 |
| S.Pe.C.inter. | SurfaceArea | 0.31 | 2.00E-06 | 0.06 |
| S.Pe.C.margi-l. | Length | 0.13 | 3.01E-02 | 0.07 |
| S.Pe.C.margi-l. | MeanDepth | 0.08 | 1.51E-01 | 0.08 |
| S.Pe.C.margi-l. | Width | 0.29 | 3.17E-05 | 0.07 |
| S.Pe.C.margi-l. | SurfaceArea | 0.11 | 7.70E-02 | 0.08 |
| S.Pe.C.median. | Length | 0.05 | 2.59E-01 | 0.08 |
| S.Pe.C.median. | MeanDepth | 0.03 | 3.52E-01 | 0.07 |
| S.Pe.C.median. | Width | 0.12 | 6.56E-02 | 0.08 |
| S.Pe.C.median. | SurfaceArea | 0.06 | 2.12E-01 | 0.07 |
| S.Pe.C.sup. | Length | 0.18 | 4.85E-03 | 0.07 |
| S.Pe.C.sup. | MeanDepth | - | - | - |
| S.Pe.C.sup. | Width | 0.13 | 3.77E-02 | 0.07 |
| S.Pe.C.sup. | SurfaceArea | 0.09 | 1.14E-01 | 0.07 |
| S.Po.C.sup. | Length | 0.27 | 5.14E-05 | 0.07 |
| S.Po.C.sup. | MeanDepth | 0.02 | 4.09E-01 | 0.07 |
| S.Po.C.sup. | Width | 0.26 | 1.21E-04 | 0.07 |
| S.Po.C.sup. | SurfaceArea | 0.08 | 1.35E-01 | 0.07 |
| S.R.inf. | Length | - | - | - |
| S.R.inf. | MeanDepth | 0.04 | 2.75E-01 | 0.07 |
| S.R.inf. | Width | 0.14 | 1.80E-02 | 0.07 |
| S.R.inf. | SurfaceArea | - | - | - |
| S.Rh. | Length | 0.17 | 1.27E-02 | 0.08 |
| S.Rh. | MeanDepth | 0.14 | 3.86E-02 | 0.08 |
| S.Rh. | Width | 0.23 | 2.02E-03 | 0.08 |
| S.Rh. | SurfaceArea | 0.16 | 2.23E-02 | 0.08 |
| S.s.P. | Length | 0.18 | 3.65E-03 | 0.07 |
| S.s.P. | MeanDepth | 0.16 | 1.58E-02 | 0.07 |
| S.s.P. | Width | 0.12 | 3.38E-02 | 0.07 |
| S.s.P. | SurfaceArea | 0.23 | 4.76E-04 | 0.07 |
| S.T.i.ant. | Length | 0.14 | 1.54E-02 | 0.06 |
| S.T.i.ant. | MeanDepth | 0.21 | 8.64E-04 | 0.06 |
| S.T.i.ant. | Width | 0.32 | 1.00E-07 | 0.06 |
| S.T.i.ant. | SurfaceArea | 0.25 | 4.55E-05 | 0.06 |
| S.T.i.post. | Length | 0.12 | 2.95E-02 | 0.07 |
| S.T.i.post. | MeanDepth | 0.11 | 5.15E-02 | 0.07 |
| S.T.i.post. | Width | 0.12 | 3.63E-02 | 0.07 |

| | | | | |
|---|---|---|---|---|
| S.T.i.post. | SurfaceArea | 0.11 | 5.32E-02 | 0.07 |
| S.T.pol. | Length | 0.26 | 1.74E-04 | 0.07 |
| S.T.pol. | MeanDepth | 0.10 | 7.11E-02 | 0.07 |
| S.T.pol. | Width | 0.14 | 2.79E-02 | 0.07 |
| S.T.pol. | SurfaceArea | 0.15 | 1.35E-02 | 0.07 |
| S.T.s. | Length | 0.13 | 2.40E-02 | 0.07 |
| S.T.s. | MeanDepth | 0.07 | 1.50E-01 | 0.07 |
| S.T.s. | Width | 0.20 | 1.32E-03 | 0.07 |
| S.T.s. | SurfaceArea | 0.20 | 1.58E-03 | 0.07 |
| S.T.s.ter.asc.ant. | Length | 0.01 | 4.66E-01 | 0.07 |
| S.T.s.ter.asc.ant. | MeanDepth | 0.07 | 1.58E-01 | 0.07 |
| S.T.s.ter.asc.ant. | Width | 0.21 | 1.45E-03 | 0.07 |
| S.T.s.ter.asc.ant. | SurfaceArea | 0.03 | 3.25E-01 | 0.07 |
| S.T.s.ter.asc.post. | Length | 0.05 | 2.10E-01 | 0.07 |
| S.T.s.ter.asc.post. | MeanDepth | 0.03 | 3.17E-01 | 0.07 |
| S.T.s.ter.asc.post. | Width | 0.17 | 5.11E-03 | 0.06 |
| S.T.s.ter.asc.post. | SurfaceArea | 0.07 | 1.32E-01 | 0.07 |
| Globals | Length | 0.22 | 1.03E-03 | 0.07 |
| Globals | MeanDepth | 0.55 | 6.84E-17 | 0.05 |
| Globals | Width | 0.36 | 6.66E-09 | 0.06 |
| Globals | SurfaceArea | 0.50 | 1.01E-13 | 0.06 |

**Table S12:** heritability eatimates for bilaterally average sulcal descriptors (yellow: bonferroni corrected for 61*4 comparisons) in QTIM.

| Sulci | Descriptor | h2 | pval | SE |
|---|---|---|---|---|
| F.C.L.a. | Length | 0.13 | 4.70E-02 | 0.07 |
| F.C.L.a. | MeanDepth | 0.15 | 2.60E-02 | 0.07 |
| F.C.L.a. | Width | 0.25 | 4.90E-04 | 0.07 |
| F.C.L.a. | SurfaceArea | 0.19 | 5.30E-03 | 0.08 |
| F.C.L.p. | Length | 0.17 | 1.10E-02 | 0.06 |
| F.C.L.p. | MeanDepth | 0.21 | 6.70E-04 | 0.07 |
| F.C.L.p. | Width | 0.61 | 7.40E-19 | 0.05 |
| F.C.L.p. | SurfaceArea | 0.37 | 1.90E-06 | 0.07 |
| F.C.L.r.ant. | Length | 0.27 | 2.90E-03 | 0.09 |
| F.C.L.r.ant. | MeanDepth | 0.14 | 7.40E-02 | 0.07 |
| F.C.L.r.ant. | Width | 0.23 | 2.40E-03 | 0.08 |
| F.C.L.r.ant. | SurfaceArea | 0.33 | 1.40E-05 | 0.09 |
| F.C.L.r.asc. | Length | 0.07 | 2.20E-01 | 0.08 |
| F.C.L.r.asc. | MeanDepth | 0.17 | 1.60E-02 | 0.08 |
| F.C.L.r.asc. | Width | 0.25 | 4.20E-04 | 0.07 |
| F.C.L.r.asc. | SurfaceArea | 0.28 | 2.60E-04 | 0.09 |
| F.C.L.r.diag. | Length | 0.30 | 2.20E-01 | 0.25 |
| F.C.L.r.diag. | MeanDepth | 0.01 | 4.90E-01 | 0.27 |
| F.C.L.r.diag. | Width | 0.10 | 5.00E-01 | 0.31 |
| F.C.L.r.diag. | SurfaceArea | 0.20 | 2.50E-01 | 0.34 |
| F.C.L.r.retroC.tr. | Length | 0.10 | 1.50E-01 | 0.08 |
| F.C.L.r.retroC.tr. | MeanDepth | 0.40 | 2.60E-06 | 0.08 |
| F.C.L.r.retroC.tr. | Width | 0.38 | 1.50E-05 | 0.08 |
| F.C.L.r.retroC.tr. | SurfaceArea | 0.27 | 4.60E-04 | 0.10 |
| F.C.L.r.sc.ant. | Length | - | - | - |
| F.C.L.r.sc.ant. | MeanDepth | - | - | - |
| F.C.L.r.sc.ant. | Width | - | - | - |
| F.C.L.r.sc.ant. | SurfaceArea | 0.10 | 5.00E-01 | - |
| F.C.L.r.sc.post. | Length | - | - | - |
| F.C.L.r.sc.post. | MeanDepth | - | 1.00E+00 | 0.11 |
| F.C.L.r.sc.post. | Width | - | 1.00E+00 | - |
| F.C.L.r.sc.post. | SurfaceArea | 0.02 | 4.30E-01 | - |
| F.C.M.ant. | Length | 0.10 | 9.20E-02 | 0.06 |
| F.C.M.ant. | MeanDepth | 0.46 | 4.40E-13 | 0.06 |
| F.C.M.ant. | Width | 0.38 | 1.00E-07 | 0.06 |
| F.C.M.ant. | SurfaceArea | 0.29 | 9.90E-06 | 0.07 |
| F.C.M.post. | Length | 0.06 | 2.10E-01 | 0.07 |
| F.C.M.post. | MeanDepth | 0.40 | 3.20E-08 | 0.07 |
| F.C.M.post. | Width | 0.57 | 3.70E-16 | 0.05 |
| F.C.M.post. | SurfaceArea | 0.31 | 9.90E-06 | 0.07 |
| F.Cal.ant.-Sc.Cal. | Length | 0.23 | 1.10E-03 | 0.05 |
| F.Cal.ant.-Sc.Cal. | MeanDepth | 0.53 | - | 0.06 |
| F.Cal.ant.-Sc.Cal. | Width | 0.37 | 1.60E-08 | 0.06 |

| | | | | |
|---|---|---|---|---|
| F.Cal.ant.-Sc.Cal. | SurfaceArea | 0.51 | - | 0.07 |
| F.Coll. | Length | 0.22 | 1.30E-03 | 0.05 |
| F.Coll. | MeanDepth | 0.54 | 2.10E-14 | 0.06 |
| F.Coll. | Width | 0.36 | 3.80E-08 | 0.06 |
| F.Coll. | SurfaceArea | 0.43 | 2.00E-09 | 0.07 |
| F.I.P. | Length | 0.23 | 4.40E-04 | 0.05 |
| F.I.P. | MeanDepth | 0.55 | 4.70E-17 | 0.06 |
| F.I.P. | Width | 0.75 | 5.70E-31 | 0.03 |
| F.I.P. | SurfaceArea | 0.37 | 3.20E-08 | 0.07 |
| F.I.P.Po.C.inf. | Length | 0.22 | 6.50E-04 | 0.06 |
| F.I.P.Po.C.inf. | MeanDepth | 0.34 | 4.00E-07 | 0.07 |
| F.I.P.Po.C.inf. | Width | 0.59 | 1.90E-18 | 0.05 |
| F.I.P.Po.C.inf. | SurfaceArea | 0.26 | 1.20E-04 | 0.07 |
| F.I.P.r.int.1 | Length | 0.29 | 5.40E-04 | 0.08 |
| F.I.P.r.int.1 | MeanDepth | 0.10 | 1.20E-01 | 0.08 |
| F.I.P.r.int.1 | Width | 0.18 | 2.00E-02 | 0.09 |
| F.I.P.r.int.1 | SurfaceArea | 0.19 | 1.10E-02 | 0.08 |
| F.I.P.r.int.2 | Length | - | 5.00E-01 | - |
| F.I.P.r.int.2 | MeanDepth | - | 5.00E-01 | 0.13 |
| F.I.P.r.int.2 | Width | 0.28 | 1.90E-02 | 0.13 |
| F.I.P.r.int.2 | SurfaceArea | - | 5.00E-01 | - |
| F.P.O. | Length | 0.26 | 3.80E-04 | 0.06 |
| F.P.O. | MeanDepth | 0.41 | 5.60E-09 | 0.06 |
| F.P.O. | Width | 0.54 | 2.40E-15 | 0.05 |
| F.P.O. | SurfaceArea | 0.43 | 5.90E-11 | 0.07 |
| INSULA | Length | 0.10 | 5.00E-01 | - |
| INSULA | MeanDepth | - | 1.00E+00 | 0.07 |
| INSULA | Width | 0.85 | 6.30E-02 | 0.16 |
| INSULA | SurfaceArea | 0.36 | 2.20E-06 | - |
| OCCIPITAL | Length | 0.54 | 1.30E-15 | 0.08 |
| OCCIPITAL | MeanDepth | 0.17 | 1.50E-02 | 0.05 |
| OCCIPITAL | Width | 0.55 | 8.30E-17 | 0.05 |
| OCCIPITAL | SurfaceArea | 0.56 | 4.60E-16 | 0.05 |
| S.C. | Length | 0.13 | 3.50E-02 | 0.07 |
| S.C. | MeanDepth | 0.44 | 3.80E-09 | 0.06 |
| S.C. | Width | 0.71 | 5.20E-29 | 0.04 |
| S.C. | SurfaceArea | 0.45 | 3.20E-11 | 0.07 |
| S.C.LPC. | Length | 0.10 | 5.00E-01 | - |
| S.C.LPC. | MeanDepth | 0.10 | 4.20E-01 | 0.46 |
| S.C.LPC. | Width | 0.10 | 4.00E-01 | - |
| S.C.LPC. | SurfaceArea | 0.33 | 2.80E-01 | 0.42 |
| S.C.sylvian. | Length | 0.25 | 1.50E-02 | 0.14 |
| S.C.sylvian. | MeanDepth | 0.29 | 2.80E-02 | 0.10 |
| S.C.sylvian. | Width | 0.41 | 1.50E-03 | 0.12 |
| S.C.sylvian. | SurfaceArea | 0.28 | 6.40E-03 | 0.11 |
| S.Call. | Length | 0.49 | 1.10E-14 | 0.06 |

| | | | | |
|---|---|---|---|---|
| S.Call. | MeanDepth | 0.46 | 5.70E-12 | 0.05 |
| S.Call. | Width | 0.27 | 1.40E-04 | 0.07 |
| S.Call. | SurfaceArea | 0.55 | 1.20E-17 | 0.05 |
| S.Cu. | Length | 0.27 | 1.00E-04 | 0.07 |
| S.Cu. | MeanDepth | 0.31 | 1.10E-05 | 0.07 |
| S.Cu. | Width | 0.37 | 1.00E-06 | 0.07 |
| S.Cu. | SurfaceArea | 0.16 | 1.60E-02 | 0.07 |
| S.F.inf. | Length | 0.04 | 2.90E-01 | 0.06 |
| S.F.inf. | MeanDepth | 0.52 | 1.00E-13 | 0.07 |
| S.F.inf. | Width | 0.40 | 2.30E-09 | 0.06 |
| S.F.inf. | SurfaceArea | 0.30 | 1.70E-05 | 0.07 |
| S.F.inf.ant. | Length | 0.10 | 1.10E-01 | 0.08 |
| S.F.inf.ant. | MeanDepth | 0.05 | 2.50E-01 | 0.08 |
| S.F.inf.ant. | Width | 0.21 | 1.40E-03 | 0.07 |
| S.F.inf.ant. | SurfaceArea | 0.17 | 1.60E-02 | 0.08 |
| S.F.int. | Length | 0.30 | 9.40E-06 | 0.06 |
| S.F.int. | MeanDepth | 0.42 | 3.10E-09 | 0.07 |
| S.F.int. | Width | 0.43 | 2.50E-10 | 0.06 |
| S.F.int. | SurfaceArea | 0.31 | 1.50E-05 | 0.07 |
| S.F.inter. | Length | 0.21 | 1.10E-03 | 0.06 |
| S.F.inter. | MeanDepth | 0.23 | 2.70E-04 | 0.07 |
| S.F.inter. | Width | 0.47 | 2.90E-12 | 0.06 |
| S.F.inter. | SurfaceArea | 0.24 | 2.80E-04 | 0.07 |
| S.F.marginal. | Length | 0.15 | 2.30E-02 | 0.07 |
| S.F.marginal. | MeanDepth | 0.10 | 1.40E-01 | 0.07 |
| S.F.marginal. | Width | 0.17 | 1.40E-02 | 0.08 |
| S.F.marginal. | SurfaceArea | 0.14 | 2.40E-02 | 0.07 |
| S.F.median. | Length | 0.27 | 8.00E-04 | 0.07 |
| S.F.median. | MeanDepth | 0.14 | 2.60E-02 | 0.08 |
| S.F.median. | Width | 0.37 | 3.00E-07 | 0.07 |
| S.F.median. | SurfaceArea | 0.25 | 1.10E-03 | 0.08 |
| S.F.orbitaire. | Length | 0.06 | 2.40E-01 | 0.07 |
| S.F.orbitaire. | MeanDepth | 0.17 | 1.40E-02 | 0.07 |
| S.F.orbitaire. | Width | 0.19 | 4.50E-03 | 0.07 |
| S.F.orbitaire. | SurfaceArea | 0.11 | 6.50E-02 | 0.08 |
| S.F.polaire.tr. | Length | 0.35 | 2.30E-06 | 0.07 |
| S.F.polaire.tr. | MeanDepth | 0.22 | 1.00E-03 | 0.07 |
| S.F.polaire.tr. | Width | 0.37 | 2.00E-07 | 0.06 |
| S.F.polaire.tr. | SurfaceArea | 0.33 | 3.70E-06 | 0.07 |
| S.F.sup. | Length | - | 5.00E-01 | 0.07 |
| S.F.sup. | MeanDepth | 0.43 | 7.90E-09 | 0.08 |
| S.F.sup. | Width | 0.57 | 6.90E-17 | 0.05 |
| S.F.sup. | SurfaceArea | 0.21 | 3.30E-03 | - |
| S.Li.ant. | Length | 0.17 | 3.20E-02 | 0.09 |
| S.Li.ant. | MeanDepth | 0.33 | 4.10E-04 | 0.09 |
| S.Li.ant. | Width | 0.40 | 6.20E-06 | 0.08 |

| | | | | |
|---|---|---|---|---|
| S.Li.ant. | SurfaceArea | 0.29 | 9.90E-04 | 0.09 |
| S.Li.post. | Length | 0.28 | 8.30E-05 | 0.08 |
| S.Li.post. | MeanDepth | 0.16 | 2.40E-02 | 0.07 |
| S.Li.post. | Width | 0.19 | 9.10E-03 | 0.08 |
| S.Li.post. | SurfaceArea | 0.28 | 9.40E-05 | 0.07 |
| S.O.p. | Length | 0.20 | 1.30E-02 | 0.08 |
| S.O.p. | MeanDepth | 0.26 | 1.30E-03 | 0.09 |
| S.O.p. | Width | 0.27 | 1.20E-03 | 0.08 |
| S.O.p. | SurfaceArea | 0.22 | 7.20E-03 | 0.09 |
| S.O.T.lat.ant. | Length | 0.16 | 1.90E-02 | 0.07 |
| S.O.T.lat.ant. | MeanDepth | 0.18 | 9.20E-03 | 0.07 |
| S.O.T.lat.ant. | Width | 0.29 | 2.70E-05 | 0.07 |
| S.O.T.lat.ant. | SurfaceArea | 0.19 | 5.00E-03 | 0.07 |
| S.O.T.lat.int. | Length | 0.10 | 2.50E-01 | 0.09 |
| S.O.T.lat.int. | MeanDepth | 0.13 | 8.00E-02 | 0.09 |
| S.O.T.lat.int. | Width | 0.15 | 4.10E-02 | 0.09 |
| S.O.T.lat.int. | SurfaceArea | 0.09 | 4.10E-01 | 0.09 |
| S.O.T.lat.med. | Length | - | 5.00E-01 | 0.09 |
| S.O.T.lat.med. | MeanDepth | 0.04 | 3.20E-01 | 0.09 |
| S.O.T.lat.med. | Width | 0.18 | 2.00E-02 | 0.09 |
| S.O.T.lat.med. | SurfaceArea | 0.01 | 4.60E-01 | - |
| S.O.T.lat.post. | Length | - | 5.00E-01 | 0.07 |
| S.O.T.lat.post. | MeanDepth | 0.34 | 2.10E-06 | 0.07 |
| S.O.T.lat.post. | Width | 0.33 | 1.00E-06 | 0.06 |
| S.O.T.lat.post. | SurfaceArea | 0.14 | 2.90E-02 | - |
| S.Olf. | Length | 0.30 | 1.60E-05 | 0.07 |
| S.Olf. | MeanDepth | 0.36 | 5.00E-07 | 0.07 |
| S.Olf. | Width | 0.28 | 3.50E-05 | 0.07 |
| S.Olf. | SurfaceArea | 0.34 | 2.10E-06 | 0.07 |
| S.Or. | Length | 0.24 | 1.70E-03 | 0.06 |
| S.Or. | MeanDepth | 0.43 | 2.10E-10 | 0.06 |
| S.Or. | Width | 0.42 | 1.00E-08 | 0.06 |
| S.Or. | SurfaceArea | 0.46 | 1.60E-09 | 0.08 |
| S.p.C. | Length | 0.15 | 3.50E-02 | 0.09 |
| S.p.C. | MeanDepth | - | 5.00E-01 | 0.09 |
| S.p.C. | Width | 0.15 | 4.50E-02 | 0.09 |
| S.p.C. | SurfaceArea | 0.10 | 1.70E-01 | 0.08 |
| S.Pa.int. | Length | 0.25 | 4.20E-04 | 0.07 |
| S.Pa.int. | MeanDepth | 0.30 | 1.20E-05 | 0.07 |
| S.Pa.int. | Width | 0.41 | 1.90E-09 | 0.06 |
| S.Pa.int. | SurfaceArea | 0.34 | 9.00E-07 | 0.07 |
| S.Pa.sup. | Length | 0.01 | 5.00E-01 | 0.09 |
| S.Pa.sup. | MeanDepth | 0.03 | 3.60E-01 | - |
| S.Pa.sup. | Width | 0.22 | 6.00E-03 | 0.09 |
| S.Pa.sup. | SurfaceArea | - | 5.00E-01 | 0.10 |
| S.Pa.t. | Length | - | 5.00E-01 | 0.15 |

| | | | | |
|---|---|---|---|---|
| S.Pa.t. | MeanDepth | 0.42 | 9.20E-03 | 0.09 |
| S.Pa.t. | Width | 0.32 | 2.70E-02 | 0.15 |
| S.Pa.t. | SurfaceArea | 0.10 | 1.50E-01 | - |
| S.Pe.C.inf. | Length | 0.11 | 7.60E-02 | 0.09 |
| S.Pe.C.inf. | MeanDepth | 0.17 | 3.00E-02 | 0.08 |
| S.Pe.C.inf. | Width | 0.41 | 9.60E-09 | 0.06 |
| S.Pe.C.inf. | SurfaceArea | 0.14 | 3.80E-02 | 0.08 |
| S.Pe.C.inter. | Length | 0.13 | 2.90E-02 | 0.07 |
| S.Pe.C.inter. | MeanDepth | 0.38 | 7.00E-07 | 0.07 |
| S.Pe.C.inter. | Width | 0.37 | 2.80E-08 | 0.06 |
| S.Pe.C.inter. | SurfaceArea | 0.17 | 7.00E-03 | 0.07 |
| S.Pe.C.marginal. | Length | 0.10 | 4.30E-01 | 0.10 |
| S.Pe.C.marginal. | MeanDepth | 0.18 | 4.50E-02 | 0.09 |
| S.Pe.C.marginal. | Width | 0.27 | 4.10E-03 | 0.10 |
| S.Pe.C.marginal. | SurfaceArea | 0.10 | 1.30E-01 | 0.10 |
| S.Pe.C.median. | Length | 0.04 | 3.30E-01 | 0.08 |
| S.Pe.C.median. | MeanDepth | 0.10 | 2.60E-01 | 0.09 |
| S.Pe.C.median. | Width | 0.16 | 3.00E-02 | 0.09 |
| S.Pe.C.median. | SurfaceArea | 0.04 | 3.30E-01 | 0.10 |
| S.Pe.C.sup. | Length | 0.10 | 9.80E-02 | 0.07 |
| S.Pe.C.sup. | MeanDepth | 0.27 | 2.50E-04 | 0.08 |
| S.Pe.C.sup. | Width | 0.39 | 2.00E-07 | 0.07 |
| S.Pe.C.sup. | SurfaceArea | 0.19 | 7.60E-03 | 0.08 |
| S.Po.C.sup. | Length | 0.10 | 8.70E-02 | 0.07 |
| S.Po.C.sup. | MeanDepth | 0.14 | 3.40E-02 | 0.07 |
| S.Po.C.sup. | Width | 0.51 | 5.40E-12 | 0.06 |
| S.Po.C.sup. | SurfaceArea | 0.11 | 7.10E-02 | 0.07 |
| S.R.inf. | Length | 0.28 | 2.60E-04 | 0.08 |
| S.R.inf. | MeanDepth | 0.23 | 2.20E-03 | 0.07 |
| S.R.inf. | Width | 0.16 | 3.70E-02 | 0.09 |
| S.R.inf. | SurfaceArea | 0.32 | 3.60E-05 | 0.08 |
| S.Rh. | Length | 0.06 | 2.20E-01 | 0.08 |
| S.Rh. | MeanDepth | 0.11 | 8.90E-02 | 0.08 |
| S.Rh. | Width | 0.22 | 3.60E-03 | 0.08 |
| S.Rh. | SurfaceArea | 0.10 | 1.40E-01 | 0.08 |
| S.s.P. | Length | 0.27 | 1.60E-04 | 0.05 |
| S.s.P. | MeanDepth | 0.51 | 8.30E-14 | 0.07 |
| S.s.P. | Width | 0.46 | 3.90E-11 | 0.06 |
| S.s.P. | SurfaceArea | 0.34 | 4.60E-06 | 0.07 |
| S.T.i.ant. | Length | 0.35 | 1.00E-07 | 0.06 |
| S.T.i.ant. | MeanDepth | 0.31 | 9.00E-07 | 0.06 |
| S.T.i.ant. | Width | 0.52 | 3.30E-16 | 0.05 |
| S.T.i.ant. | SurfaceArea | 0.41 | 1.60E-10 | 0.06 |
| S.T.i.post. | Length | 0.06 | 2.20E-01 | 0.07 |
| S.T.i.post. | MeanDepth | 0.32 | 4.60E-06 | 0.08 |
| S.T.i.post. | Width | 0.43 | 1.30E-10 | 0.06 |

| | | | | |
|---|---|---|---|---|
| S.T.i.post. | SurfaceArea | 0.12 | 5.50E-02 | 0.08 |
| S.T.pol. | Length | 0.26 | 2.60E-04 | 0.09 |
| S.T.pol. | MeanDepth | 0.13 | 6.70E-02 | 0.08 |
| S.T.pol. | Width | 0.29 | 1.40E-04 | 0.08 |
| S.T.pol. | SurfaceArea | 0.27 | 4.10E-04 | 0.07 |
| S.T.s. | Length | 0.13 | 3.00E-02 | 0.07 |
| S.T.s. | MeanDepth | 0.40 | 4.80E-08 | 0.07 |
| S.T.s. | Width | 0.34 | 4.00E-07 | 0.06 |
| S.T.s. | SurfaceArea | 0.29 | 1.20E-05 | 0.07 |
| S.T.s.ter.asc.ant. | Length | 0.10 | 7.80E-02 | 0.09 |
| S.T.s.ter.asc.ant. | MeanDepth | 0.10 | 2.40E-01 | 0.08 |
| S.T.s.ter.asc.ant. | Width | 0.37 | 1.60E-06 | 0.07 |
| S.T.s.ter.asc.ant. | SurfaceArea | 0.01 | 5.00E-01 | 0.08 |
| S.T.s.ter.asc.post. | Length | 0.09 | 1.10E-01 | 0.08 |
| S.T.s.ter.asc.post. | MeanDepth | 0.19 | 1.40E-02 | 0.07 |
| S.T.s.ter.asc.post. | Width | 0.27 | 2.10E-04 | 0.07 |
| S.T.s.ter.asc.post. | SurfaceArea | 0.17 | 1.10E-02 | 0.07 |
| Globals | Length | 0.27 | 9.10E-05 | 0.07 |
| Globals | MeanDepth | 0.16 | 1.20E-02 | 0.07 |
| Globals | Width | 0.23 | 9.90E-04 | 0.07 |
| Globals | SurfaceArea | 0.31 | 8.70E-06 | 0.07 |

**Table S13:** heritability eatimates for left hemisphere sulcal descriptors (yellow: bonferroni corrected for 123*4 comparisons) in HCP.

| Sulci | Descriptor | h2 | pval | SE |
|---|---|---|---|---|
| F.C.L.a. | Length | 0.07 | 2.29E-01 | 0.09 |
| F.C.L.a. | MeanDepth | 0.21 | 5.47E-03 | 0.09 |
| F.C.L.a. | Width | 0.34 | 5.95E-05 | 0.09 |
| F.C.L.a. | SurfaceArea | 0.28 | 3.84E-04 | 0.09 |
| F.C.L.p. | Length | - | - | - |
| F.C.L.p. | MeanDepth | 0.13 | 3.33E-02 | 0.07 |
| F.C.L.p. | Width | 0.55 | 6.61E-12 | 0.07 |
| F.C.L.p. | SurfaceArea | 0.32 | 4.90E-06 | 0.07 |
| F.C.L.r.ant. | Length | 0.06 | 2.36E-01 | 0.08 |
| F.C.L.r.ant. | MeanDepth | 0.13 | 4.86E-02 | 0.08 |
| F.C.L.r.ant. | Width | 0.11 | 7.06E-02 | 0.08 |
| F.C.L.r.ant. | SurfaceArea | 0.20 | 2.58E-03 | 0.07 |
| F.C.L.r.asc. | Length | - | - | - |
| F.C.L.r.asc. | MeanDepth | 0.09 | 1.33E-01 | 0.08 |
| F.C.L.r.asc. | Width | 0.17 | 2.98E-02 | 0.09 |
| F.C.L.r.asc. | SurfaceArea | 0.26 | 1.37E-03 | 0.09 |
| F.C.L.r.diag. | Length | - | - | - |
| F.C.L.r.diag. | MeanDepth | 0.16 | 1.58E-01 | 0.17 |
| F.C.L.r.diag. | Width | 0.24 | 3.55E-02 | 0.13 |
| F.C.L.r.diag. | SurfaceArea | 0.08 | 3.06E-01 | 0.16 |
| F.C.L.r.retroC.tr. | Length | 0.15 | 3.62E-02 | 0.08 |
| F.C.L.r.retroC.tr. | MeanDepth | 0.14 | 3.77E-02 | 0.08 |
| F.C.L.r.retroC.tr. | Width | 0.08 | 1.78E-01 | 0.09 |
| F.C.L.r.retroC.tr. | SurfaceArea | 0.07 | 1.72E-01 | 0.08 |
| F.C.L.r.sc.ant. | Length | - | - | - |
| F.C.L.r.sc.ant. | MeanDepth | - | - | - |
| F.C.L.r.sc.ant. | Width | - | - | - |
| F.C.L.r.sc.ant. | SurfaceArea | 0.26 | 2.96E-01 | 0.46 |
| F.C.L.r.sc.post. | Length | - | - | - |
| F.C.L.r.sc.post. | MeanDepth | - | - | - |
| F.C.L.r.sc.post. | Width | 0.73 | 1.64E-02 | 0.17 |
| F.C.L.r.sc.post. | SurfaceArea | 0.03 | 3.63E-01 | 0.09 |
| F.C.M.ant. | Length | - | - | - |
| F.C.M.ant. | MeanDepth | 0.12 | 5.20E-02 | 0.07 |
| F.C.M.ant. | Width | 0.17 | 1.26E-02 | 0.08 |
| F.C.M.ant. | SurfaceArea | - | - | - |
| F.C.M.post. | Length | 0.09 | 1.08E-01 | 0.07 |
| F.C.M.post. | MeanDepth | 0.32 | 5.90E-06 | 0.07 |
| F.C.M.post. | Width | 0.59 | 2.49E-16 | 0.06 |
| F.C.M.post. | SurfaceArea | 0.21 | 1.96E-03 | 0.07 |
| F.Cal.ant.-Sc.Cal. | Length | 0.01 | 4.45E-01 | 0.08 |
| F.Cal.ant.-Sc.Cal. | MeanDepth | 0.40 | 6.00E-07 | 0.08 |
| F.Cal.ant.-Sc.Cal. | Width | 0.31 | 3.10E-05 | 0.07 |

| | | | | |
|---|---|---|---|---|
| F.Cal.ant.-Sc.Cal. | SurfaceArea | 0.44 | 3.52E-08 | 0.07 |
| F.Coll. | Length | 0.12 | 4.31E-02 | 0.07 |
| F.Coll. | MeanDepth | 0.34 | 1.90E-06 | 0.07 |
| F.Coll. | Width | 0.40 | 8.20E-10 | 0.06 |
| F.Coll. | SurfaceArea | 0.21 | 2.28E-03 | 0.08 |
| F.I.P. | Length | 0.12 | 5.71E-02 | 0.08 |
| F.I.P. | MeanDepth | 0.31 | 7.10E-06 | 0.07 |
| F.I.P. | Width | 0.56 | 5.41E-15 | 0.06 |
| F.I.P. | SurfaceArea | 0.24 | 6.49E-04 | 0.08 |
| F.I.P.Po.C.inf. | Length | 0.10 | 7.81E-02 | 0.07 |
| F.I.P.Po.C.inf. | MeanDepth | 0.13 | 4.18E-02 | 0.08 |
| F.I.P.Po.C.inf. | Width | 0.46 | 6.20E-10 | 0.07 |
| F.I.P.Po.C.inf. | SurfaceArea | 0.19 | 3.70E-03 | 0.07 |
| F.I.P.r.int.1 | Length | 0.07 | 1.64E-01 | 0.08 |
| F.I.P.r.int.1 | MeanDepth | 0.01 | 4.59E-01 | 0.08 |
| F.I.P.r.int.1 | Width | 0.10 | 7.50E-02 | 0.07 |
| F.I.P.r.int.1 | SurfaceArea | 0.06 | 2.20E-01 | 0.08 |
| F.I.P.r.int.2 | Length | - | - | - |
| F.I.P.r.int.2 | MeanDepth | - | - | - |
| F.I.P.r.int.2 | Width | - | - | - |
| F.I.P.r.int.2 | SurfaceArea | - | - | - |
| F.P.O. | Length | 0.10 | 7.32E-02 | 0.07 |
| F.P.O. | MeanDepth | 0.46 | 5.68E-09 | 0.07 |
| F.P.O. | Width | 0.45 | 8.20E-11 | 0.07 |
| F.P.O. | SurfaceArea | 0.34 | 1.30E-06 | 0.07 |
| INSULA | Length | 0.21 | 2.57E-01 | 0.31 |
| INSULA | MeanDepth | 0.07 | 3.81E-01 | 0.23 |
| INSULA | Width | 0.59 | 5.61E-03 | 0.18 |
| INSULA | SurfaceArea | 0.22 | 5.37E-03 | 0.09 |
| OCCIPITAL | Length | 0.33 | 3.40E-06 | 0.07 |
| OCCIPITAL | MeanDepth | 0.30 | 9.20E-06 | 0.07 |
| OCCIPITAL | Width | 0.37 | 2.00E-07 | 0.07 |
| OCCIPITAL | SurfaceArea | 0.41 | 4.08E-09 | 0.07 |
| S.C. | Length | 0.11 | 9.53E-02 | 0.08 |
| S.C. | MeanDepth | 0.42 | 3.00E-07 | 0.08 |
| S.C. | Width | 0.68 | 1.09E-21 | 0.05 |
| S.C. | SurfaceArea | 0.34 | 4.00E-06 | 0.07 |
| S.C.LPC. | Length | 0.03 | 4.34E-01 | 0.18 |
| S.C.LPC. | MeanDepth | 0.40 | 3.64E-03 | 0.15 |
| S.C.LPC. | Width | 0.30 | 8.44E-03 | 0.13 |
| S.C.LPC. | SurfaceArea | 0.18 | 1.11E-01 | 0.15 |
| S.C.sylvian. | Length | 0.37 | 8.37E-04 | 0.11 |
| S.C.sylvian. | MeanDepth | 0.24 | 2.97E-02 | 0.13 |
| S.C.sylvian. | Width | 0.37 | 1.99E-03 | 0.12 |
| S.C.sylvian. | SurfaceArea | 0.37 | 1.49E-04 | 0.10 |
| S.Call. | Length | 0.17 | 1.76E-02 | 0.09 |

| | | | | |
|---|---|---|---|---|
| S.Call. | MeanDepth | 0.44 | 7.23E-10 | 0.07 |
| S.Call. | Width | 0.26 | 9.63E-05 | 0.07 |
| S.Call. | SurfaceArea | 0.24 | 1.89E-03 | 0.08 |
| S.Cu. | Length | 0.26 | 1.49E-04 | 0.07 |
| S.Cu. | MeanDepth | - | - | - |
| S.Cu. | Width | 0.16 | 2.07E-02 | 0.08 |
| S.Cu. | SurfaceArea | 0.13 | 4.66E-02 | 0.08 |
| S.F.inf. | Length | 0.09 | 9.11E-02 | 0.07 |
| S.F.inf. | MeanDepth | 0.35 | 1.00E-06 | 0.07 |
| S.F.inf. | Width | 0.43 | 3.47E-09 | 0.07 |
| S.F.inf. | SurfaceArea | 0.28 | 4.04E-05 | 0.07 |
| S.F.inf.ant. | Length | - | - | - |
| S.F.inf.ant. | MeanDepth | 0.10 | 7.56E-02 | 0.07 |
| S.F.inf.ant. | Width | 0.08 | 1.47E-01 | 0.08 |
| S.F.inf.ant. | SurfaceArea | 0.02 | 4.08E-01 | 0.07 |
| S.F.int. | Length | 0.37 | 3.00E-07 | 0.07 |
| S.F.int. | MeanDepth | 0.26 | 1.64E-04 | 0.07 |
| S.F.int. | Width | 0.35 | 1.23E-05 | 0.08 |
| S.F.int. | SurfaceArea | 0.36 | 4.00E-07 | 0.07 |
| S.F.inter. | Length | - | - | - |
| S.F.inter. | MeanDepth | 0.24 | 4.95E-04 | 0.08 |
| S.F.inter. | Width | 0.41 | 4.07E-09 | 0.07 |
| S.F.inter. | SurfaceArea | - | - | - |
| S.F.marginal. | Length | 0.18 | 8.72E-03 | 0.08 |
| S.F.marginal. | MeanDepth | 0.04 | 3.13E-01 | 0.07 |
| S.F.marginal. | Width | 0.21 | 2.12E-03 | 0.08 |
| S.F.marginal. | SurfaceArea | 0.16 | 1.82E-02 | 0.08 |
| S.F.median. | Length | 0.17 | 1.60E-02 | 0.08 |
| S.F.median. | MeanDepth | 0.24 | 1.50E-04 | 0.07 |
| S.F.median. | Width | 0.35 | 2.00E-07 | 0.07 |
| S.F.median. | SurfaceArea | 0.25 | 5.06E-04 | 0.08 |
| S.F.orbitaire. | Length | - | - | - |
| S.F.orbitaire. | MeanDepth | - | - | - |
| S.F.orbitaire. | Width | - | - | - |
| S.F.orbitaire. | SurfaceArea | - | - | - |
| S.F.polaire.tr. | Length | 0.25 | 2.31E-03 | 0.09 |
| S.F.polaire.tr. | MeanDepth | 0.19 | 3.30E-03 | 0.07 |
| S.F.polaire.tr. | Width | 0.36 | 2.90E-06 | 0.08 |
| S.F.polaire.tr. | SurfaceArea | 0.22 | 2.88E-03 | 0.08 |
| S.F.sup. | Length | 0.07 | 1.82E-01 | 0.08 |
| S.F.sup. | MeanDepth | 0.30 | 3.80E-05 | 0.07 |
| S.F.sup. | Width | 0.50 | 5.39E-12 | 0.07 |
| S.F.sup. | SurfaceArea | 0.23 | 7.52E-04 | 0.07 |
| S.GSM. | Length | - | - | - |
| S.GSM. | MeanDepth | 0.18 | 3.11E-02 | 0.10 |
| S.GSM. | Width | 0.18 | 4.54E-02 | 0.11 |

| | | | | |
|---|---|---|---|---|
| S.GSM. | SurfaceArea | 0.10 | 1.64E-01 | 0.10 |
| S.Li.ant. | Length | 0.16 | 3.06E-02 | 0.09 |
| S.Li.ant. | MeanDepth | 0.32 | 2.80E-04 | 0.09 |
| S.Li.ant. | Width | 0.30 | 1.76E-04 | 0.08 |
| S.Li.ant. | SurfaceArea | 0.22 | 6.17E-03 | 0.09 |
| S.Li.post. | Length | 0.14 | 3.83E-02 | 0.08 |
| S.Li.post. | MeanDepth | 0.21 | 3.10E-03 | 0.08 |
| S.Li.post. | Width | 0.27 | 2.68E-04 | 0.08 |
| S.Li.post. | SurfaceArea | 0.12 | 6.08E-02 | 0.08 |
| S.O.p. | Length | - | - | - |
| S.O.p. | MeanDepth | 0.13 | 8.50E-02 | 0.10 |
| S.O.p. | Width | 0.28 | 2.53E-03 | 0.10 |
| S.O.p. | SurfaceArea | 0.02 | 4.27E-01 | 0.10 |
| S.O.T.lat.ant. | Length | 0.13 | 4.35E-02 | 0.07 |
| S.O.T.lat.ant. | MeanDepth | 0.21 | 2.02E-03 | 0.07 |
| S.O.T.lat.ant. | Width | 0.18 | 6.18E-03 | 0.07 |
| S.O.T.lat.ant. | SurfaceArea | 0.20 | 2.72E-03 | 0.07 |
| S.O.T.lat.int. | Length | 0.10 | 1.08E-01 | 0.08 |
| S.O.T.lat.int. | MeanDepth | - | - | - |
| S.O.T.lat.int. | Width | - | - | - |
| S.O.T.lat.int. | SurfaceArea | - | - | - |
| S.O.T.lat.med. | Length | 0.09 | 1.51E-01 | 0.08 |
| S.O.T.lat.med. | MeanDepth | 0.07 | 1.88E-01 | 0.08 |
| S.O.T.lat.med. | Width | 0.08 | 1.27E-01 | 0.08 |
| S.O.T.lat.med. | SurfaceArea | 0.08 | 1.62E-01 | 0.08 |
| S.O.T.lat.post. | Length | 0.05 | 2.75E-01 | 0.08 |
| S.O.T.lat.post. | MeanDepth | 0.17 | 1.41E-02 | 0.08 |
| S.O.T.lat.post. | Width | 0.11 | 6.44E-02 | 0.08 |
| S.O.T.lat.post. | SurfaceArea | 0.09 | 1.08E-01 | 0.08 |
| S.Olf. | Length | 0.18 | 1.81E-02 | 0.09 |
| S.Olf. | MeanDepth | 0.47 | 1.59E-10 | 0.07 |
| S.Olf. | Width | 0.28 | 1.55E-05 | 0.07 |
| S.Olf. | SurfaceArea | 0.41 | 1.00E-07 | 0.07 |
| S.Or. | Length | 0.29 | 1.18E-04 | 0.08 |
| S.Or. | MeanDepth | 0.22 | 7.75E-03 | 0.09 |
| S.Or. | Width | 0.19 | 1.13E-02 | 0.08 |
| S.Or. | SurfaceArea | 0.39 | 1.10E-06 | 0.08 |
| S.p.C. | Length | 0.07 | 2.08E-01 | 0.08 |
| S.p.C. | MeanDepth | 0.13 | 5.00E-02 | 0.08 |
| S.p.C. | Width | 0.27 | 1.86E-04 | 0.08 |
| S.p.C. | SurfaceArea | 0.12 | 6.22E-02 | 0.08 |
| S.Pa.int. | Length | 0.16 | 1.02E-02 | 0.07 |
| S.Pa.int. | MeanDepth | 0.27 | 3.03E-04 | 0.08 |
| S.Pa.int. | Width | 0.23 | 3.32E-03 | 0.09 |
| S.Pa.int. | SurfaceArea | 0.11 | 5.45E-02 | 0.07 |
| S.Pa.sup. | Length | 0.00 | 4.79E-01 | 0.08 |

| | | | | |
|---|---|---|---|---|
| S.Pa.sup. | MeanDepth | 0.05 | 2.84E-01 | 0.09 |
| S.Pa.sup. | Width | 0.29 | 5.25E-04 | 0.09 |
| S.Pa.sup. | SurfaceArea | 0.07 | 2.00E-01 | 0.08 |
| S.Pa.t. | Length | - | - | - |
| S.Pa.t. | MeanDepth | 0.14 | 1.35E-01 | 0.13 |
| S.Pa.t. | Width | 0.20 | 6.73E-02 | 0.13 |
| S.Pa.t. | SurfaceArea | 0.15 | 5.39E-02 | 0.09 |
| S.Pe.C.inf. | Length | 0.04 | 3.01E-01 | 0.07 |
| S.Pe.C.inf. | MeanDepth | 0.20 | 6.86E-03 | 0.08 |
| S.Pe.C.inf. | Width | 0.21 | 4.12E-03 | 0.08 |
| S.Pe.C.inf. | SurfaceArea | 0.08 | 1.63E-01 | 0.08 |
| S.Pe.C.inter. | Length | 0.04 | 2.92E-01 | 0.08 |
| S.Pe.C.inter. | MeanDepth | 0.10 | 8.50E-02 | 0.07 |
| S.Pe.C.inter. | Width | 0.15 | 3.39E-02 | 0.08 |
| S.Pe.C.inter. | SurfaceArea | 0.07 | 1.77E-01 | 0.08 |
| S.Pe.C.marginal. | Length | - | - | - |
| S.Pe.C.marginal. | MeanDepth | 0.05 | 3.01E-01 | 0.09 |
| S.Pe.C.marginal. | Width | 0.20 | 9.00E-03 | 0.08 |
| S.Pe.C.marginal. | SurfaceArea | - | - | - |
| S.Pe.C.median. | Length | 0.04 | 3.11E-01 | 0.08 |
| S.Pe.C.median. | MeanDepth | 0.08 | 1.42E-01 | 0.08 |
| S.Pe.C.median. | Width | 0.19 | 7.66E-03 | 0.08 |
| S.Pe.C.median. | SurfaceArea | 0.10 | 1.13E-01 | 0.08 |
| S.Pe.C.sup. | Length | 0.12 | 5.36E-02 | 0.08 |
| S.Pe.C.sup. | MeanDepth | 0.00 | 4.82E-01 | 0.08 |
| S.Pe.C.sup. | Width | 0.21 | 4.71E-03 | 0.08 |
| S.Pe.C.sup. | SurfaceArea | 0.16 | 1.57E-02 | 0.07 |
| S.Po.C.sup. | Length | 0.09 | 1.29E-01 | 0.08 |
| S.Po.C.sup. | MeanDepth | 0.15 | 2.49E-02 | 0.08 |
| S.Po.C.sup. | Width | 0.47 | 5.54E-09 | 0.08 |
| S.Po.C.sup. | SurfaceArea | 0.15 | 3.48E-02 | 0.08 |
| S.R.inf. | Length | 0.14 | 6.85E-02 | 0.09 |
| S.R.inf. | MeanDepth | 0.04 | 3.18E-01 | 0.08 |
| S.R.inf. | Width | 0.14 | 5.54E-02 | 0.09 |
| S.R.inf. | SurfaceArea | 0.12 | 8.29E-02 | 0.09 |
| S.Rh. | Length | 0.25 | 4.06E-04 | 0.08 |
| S.Rh. | MeanDepth | 0.38 | 7.00E-07 | 0.08 |
| S.Rh. | Width | 0.43 | 1.67E-08 | 0.07 |
| S.Rh. | SurfaceArea | 0.33 | 8.60E-06 | 0.08 |
| S.s.P. | Length | 0.07 | 1.91E-01 | 0.08 |
| S.s.P. | MeanDepth | 0.30 | 2.94E-05 | 0.07 |
| S.s.P. | Width | 0.32 | 1.29E-05 | 0.08 |
| S.s.P. | SurfaceArea | 0.13 | 5.12E-02 | 0.08 |
| S.T.i.ant. | Length | 0.09 | 1.11E-01 | 0.07 |
| S.T.i.ant. | MeanDepth | 0.12 | 5.32E-02 | 0.08 |
| S.T.i.ant. | Width | 0.21 | 2.07E-03 | 0.07 |

| | | | | |
|---|---|---|---|---|
| S.T.i.ant. | SurfaceArea | 0.16 | 1.42E-02 | 0.07 |
| S.T.i.post. | Length | - | - | - |
| S.T.i.post. | MeanDepth | 0.10 | 1.09E-01 | 0.09 |
| S.T.i.post. | Width | 0.16 | 1.46E-02 | 0.08 |
| S.T.i.post. | SurfaceArea | 0.01 | 4.50E-01 | 0.07 |
| S.T.pol. | Length | 0.43 | 5.82E-09 | 0.07 |
| S.T.pol. | MeanDepth | 0.21 | 1.91E-03 | 0.07 |
| S.T.pol. | Width | 0.24 | 4.49E-04 | 0.07 |
| S.T.pol. | SurfaceArea | 0.45 | 4.55E-09 | 0.07 |
| S.T.s. | Length | 0.05 | 2.43E-01 | 0.07 |
| S.T.s. | MeanDepth | 0.31 | 7.10E-06 | 0.07 |
| S.T.s. | Width | 0.34 | 2.00E-07 | 0.07 |
| S.T.s. | SurfaceArea | 0.02 | 3.88E-01 | 0.08 |
| S.T.s.ter.asc.ant. | Length | - | - | - |
| S.T.s.ter.asc.ant. | MeanDepth | 0.03 | 3.44E-01 | 0.07 |
| S.T.s.ter.asc.ant. | Width | 0.19 | 3.96E-03 | 0.07 |
| S.T.s.ter.asc.ant. | SurfaceArea | - | - | - |
| S.T.s.ter.asc.post. | Length | 0.04 | 3.07E-01 | 0.07 |
| S.T.s.ter.asc.post. | MeanDepth | 0.01 | 4.32E-01 | 0.08 |
| S.T.s.ter.asc.post. | Width | 0.25 | 6.66E-04 | 0.08 |
| S.T.s.ter.asc.post. | SurfaceArea | 0.12 | 5.14E-02 | 0.08 |
| Globals | Length | 0.09 | 1.45E-01 | 0.08 |
| Globals | MeanDepth | 0.09 | 1.45E-01 | 0.08 |
| Globals | Width | 0.27 | 8.02E-05 | 0.07 |
| Globals | SurfaceArea | 0.29 | 3.43E-04 | 0.08 |

**Table S14:** heritability eatimates for right hemisphere sulcal descriptors (yellow: bonferroni corrected for 123*4 comparisons) in HCP.

| Sulci | Descriptor | h2 | pval | SE |
|---|---|---|---|---|
| F.C.L.a. | Length | 0.16 | 1.88E-02 | 0.08 |
| F.C.L.a. | MeanDepth | 0.00 | 4.72E-01 | 0.07 |
| F.C.L.a. | Width | 0.30 | 4.80E-05 | 0.08 |
| F.C.L.a. | SurfaceArea | 0.35 | 4.60E-06 | 0.08 |
| F.C.L.p. | Length | 0.09 | 1.50E-01 | 0.08 |
| F.C.L.p. | MeanDepth | 0.18 | 1.37E-02 | 0.08 |
| F.C.L.p. | Width | 0.43 | 2.11E-08 | 0.07 |
| F.C.L.p. | SurfaceArea | 0.62 | 1.37E-16 | 0.06 |
| F.C.L.r.ant. | Length | 0.28 | 6.47E-04 | 0.09 |
| F.C.L.r.ant. | MeanDepth | 0.09 | 1.66E-01 | 0.10 |
| F.C.L.r.ant. | Width | 0.04 | 3.28E-01 | 0.10 |
| F.C.L.r.ant. | SurfaceArea | 0.15 | 4.40E-02 | 0.09 |
| F.C.L.r.asc. | Length | 0.02 | 3.93E-01 | 0.08 |
| F.C.L.r.asc. | MeanDepth | 0.05 | 2.70E-01 | 0.08 |
| F.C.L.r.asc. | Width | 0.16 | 2.48E-02 | 0.08 |
| F.C.L.r.asc. | SurfaceArea | 0.20 | 6.24E-03 | 0.08 |
| F.C.L.r.diag. | Length | - | - | - |
| F.C.L.r.diag. | MeanDepth | - | - | - |
| F.C.L.r.diag. | Width | 0.18 | 4.56E-02 | 0.11 |
| F.C.L.r.diag. | SurfaceArea | - | - | - |
| F.C.L.r.retroC.tr. | Length | 0.08 | 1.67E-01 | 0.09 |
| F.C.L.r.retroC.tr. | MeanDepth | 0.20 | 1.58E-02 | 0.10 |
| F.C.L.r.retroC.tr. | Width | 0.26 | 2.89E-03 | 0.09 |
| F.C.L.r.retroC.tr. | SurfaceArea | 0.07 | 2.27E-01 | 0.10 |
| F.C.L.r.sc.ant. | Length | - | - | - |
| F.C.L.r.sc.ant. | MeanDepth | - | - | - |
| F.C.L.r.sc.ant. | Width | - | - | - |
| F.C.L.r.sc.ant. | SurfaceArea | - | - | - |
| F.C.L.r.sc.post. | Length | - | - | - |
| F.C.L.r.sc.post. | MeanDepth | - | - | - |
| F.C.L.r.sc.post. | Width | - | - | - |
| F.C.L.r.sc.post. | SurfaceArea | 0.14 | 5.98E-02 | 0.09 |
| F.C.M.ant. | Length | 0.06 | 1.89E-01 | 0.07 |
| F.C.M.ant. | MeanDepth | 0.21 | 1.45E-03 | 0.07 |
| F.C.M.ant. | Width | 0.30 | 5.20E-06 | 0.07 |
| F.C.M.ant. | SurfaceArea | 0.17 | 7.50E-03 | 0.07 |
| F.C.M.post. | Length | 0.01 | 4.14E-01 | 0.07 |
| F.C.M.post. | MeanDepth | 0.35 | 6.00E-07 | 0.07 |
| F.C.M.post. | Width | 0.40 | 1.76E-08 | 0.07 |
| F.C.M.post. | SurfaceArea | 0.23 | 5.31E-04 | 0.07 |
| F.Cal.ant.-Sc.Cal. | Length | 0.13 | 2.63E-02 | 0.07 |
| F.Cal.ant.-Sc.Cal. | MeanDepth | 0.52 | 6.10E-14 | 0.06 |
| F.Cal.ant.-Sc.Cal. | Width | 0.47 | 4.19E-09 | 0.07 |

| | | | | |
|---|---|---|---|---|
| F.Cal.ant.-Sc.Cal. | SurfaceArea | 0.52 | 1.31E-10 | 0.07 |
| F.Coll. | Length | 0.05 | 2.70E-01 | 0.08 |
| F.Coll. | MeanDepth | 0.32 | 1.00E-06 | 0.07 |
| F.Coll. | Width | 0.46 | 3.14E-11 | 0.06 |
| F.Coll. | SurfaceArea | 0.23 | 2.66E-03 | 0.08 |
| F.I.P. | Length | 0.24 | 5.18E-04 | 0.07 |
| F.I.P. | MeanDepth | 0.38 | 3.52E-09 | 0.06 |
| F.I.P. | Width | 0.62 | 9.39E-16 | 0.06 |
| F.I.P. | SurfaceArea | 0.20 | 4.98E-03 | 0.08 |
| F.I.P.Po.C.inf. | Length | 0.16 | 1.91E-02 | 0.08 |
| F.I.P.Po.C.inf. | MeanDepth | 0.28 | 2.96E-04 | 0.08 |
| F.I.P.Po.C.inf. | Width | 0.40 | 2.00E-07 | 0.08 |
| F.I.P.Po.C.inf. | SurfaceArea | 0.16 | 1.61E-02 | 0.08 |
| F.I.P.r.int.1 | Length | 0.23 | 9.03E-04 | 0.08 |
| F.I.P.r.int.1 | MeanDepth | 0.15 | 3.26E-02 | 0.08 |
| F.I.P.r.int.1 | Width | 0.29 | 4.02E-04 | 0.09 |
| F.I.P.r.int.1 | SurfaceArea | 0.24 | 1.88E-03 | 0.09 |
| F.I.P.r.int.2 | Length | - | - | - |
| F.I.P.r.int.2 | MeanDepth | - | - | - |
| F.I.P.r.int.2 | Width | 0.05 | 3.14E-01 | 0.10 |
| F.I.P.r.int.2 | SurfaceArea | - | - | - |
| F.P.O. | Length | 0.16 | 2.19E-02 | 0.08 |
| F.P.O. | MeanDepth | 0.33 | 7.00E-07 | 0.07 |
| F.P.O. | Width | 0.38 | 3.00E-07 | 0.07 |
| F.P.O. | SurfaceArea | 0.51 | 1.19E-12 | 0.07 |
| INSULA | Length | 0.35 | 9.75E-02 | 0.27 |
| INSULA | MeanDepth | 0.26 | 1.65E-01 | 0.26 |
| INSULA | Width | 0.26 | 1.83E-01 | 0.29 |
| INSULA | SurfaceArea | 0.40 | 4.00E-07 | 0.08 |
| OCCIPITAL | Length | 0.36 | 4.00E-07 | 0.07 |
| OCCIPITAL | MeanDepth | 0.34 | 1.44E-05 | 0.08 |
| OCCIPITAL | Width | 0.49 | 1.44E-11 | 0.07 |
| OCCIPITAL | SurfaceArea | 0.42 | 2.38E-09 | 0.07 |
| S.C. | Length | 0.05 | 2.80E-01 | 0.08 |
| S.C. | MeanDepth | 0.31 | 1.55E-05 | 0.08 |
| S.C. | Width | 0.71 | 1.14E-24 | 0.05 |
| S.C. | SurfaceArea | 0.27 | 6.64E-04 | 0.09 |
| S.C.LPC. | Length | 0.03 | 4.86E-01 | 0.76 |
| S.C.LPC. | MeanDepth | - | - | - |
| S.C.LPC. | Width | - | - | - |
| S.C.LPC. | SurfaceArea | 0.34 | 3.14E-01 | 0.70 |
| S.C.sylvian. | Length | 0.28 | 5.94E-03 | 0.11 |
| S.C.sylvian. | MeanDepth | 0.10 | 1.88E-01 | 0.11 |
| S.C.sylvian. | Width | 0.15 | 7.85E-02 | 0.11 |
| S.C.sylvian. | SurfaceArea | 0.09 | 1.95E-01 | 0.11 |
| S.Call. | Length | 0.33 | 9.00E-07 | 0.07 |

| | | | | |
|---|---|---|---|---|
| S.Call. | MeanDepth | 0.41 | 7.94E-09 | 0.07 |
| S.Call. | Width | 0.26 | 2.69E-04 | 0.08 |
| S.Call. | SurfaceArea | 0.35 | 4.00E-07 | 0.07 |
| S.Cu. | Length | 0.21 | 1.53E-03 | 0.07 |
| S.Cu. | MeanDepth | 0.15 | 2.04E-02 | 0.08 |
| S.Cu. | Width | 0.12 | 5.18E-02 | 0.08 |
| S.Cu. | SurfaceArea | 0.17 | 6.47E-03 | 0.07 |
| S.F.inf. | Length | 0.04 | 2.90E-01 | 0.07 |
| S.F.inf. | MeanDepth | 0.29 | 1.51E-04 | 0.08 |
| S.F.inf. | Width | 0.48 | 8.57E-11 | 0.07 |
| S.F.inf. | SurfaceArea | 0.18 | 9.79E-03 | 0.08 |
| S.F.inf.ant. | Length | 0.20 | 6.31E-03 | 0.08 |
| S.F.inf.ant. | MeanDepth | 0.13 | 3.96E-02 | 0.08 |
| S.F.inf.ant. | Width | 0.19 | 7.96E-03 | 0.08 |
| S.F.inf.ant. | SurfaceArea | 0.21 | 5.06E-03 | 0.08 |
| S.F.int. | Length | 0.26 | 3.94E-04 | 0.08 |
| S.F.int. | MeanDepth | 0.20 | 3.32E-03 | 0.07 |
| S.F.int. | Width | 0.20 | 3.85E-03 | 0.08 |
| S.F.int. | SurfaceArea | 0.29 | 4.71E-05 | 0.08 |
| S.F.inter. | Length | 0.07 | 1.47E-01 | 0.07 |
| S.F.inter. | MeanDepth | 0.13 | 3.59E-02 | 0.07 |
| S.F.inter. | Width | 0.37 | 1.00E-07 | 0.07 |
| S.F.inter. | SurfaceArea | 0.12 | 4.51E-02 | 0.07 |
| S.F.marginal. | Length | 0.26 | 1.27E-04 | 0.07 |
| S.F.marginal. | MeanDepth | - | - | - |
| S.F.marginal. | Width | 0.17 | 6.67E-03 | 0.07 |
| S.F.marginal. | SurfaceArea | 0.18 | 4.90E-03 | 0.07 |
| S.F.median. | Length | 0.36 | 8.00E-07 | 0.07 |
| S.F.median. | MeanDepth | 0.16 | 2.17E-02 | 0.08 |
| S.F.median. | Width | 0.30 | 1.99E-05 | 0.07 |
| S.F.median. | SurfaceArea | 0.35 | 1.00E-06 | 0.07 |
| S.F.orbitaire. | Length | 0.14 | 3.70E-02 | 0.08 |
| S.F.orbitaire. | MeanDepth | 0.14 | 5.79E-02 | 0.09 |
| S.F.orbitaire. | Width | 0.21 | 3.15E-03 | 0.08 |
| S.F.orbitaire. | SurfaceArea | 0.16 | 2.31E-02 | 0.08 |
| S.F.polaire.tr. | Length | 0.14 | 3.37E-02 | 0.08 |
| S.F.polaire.tr. | MeanDepth | 0.04 | 2.93E-01 | 0.07 |
| S.F.polaire.tr. | Width | 0.38 | 4.40E-06 | 0.08 |
| S.F.polaire.tr. | SurfaceArea | 0.12 | 6.45E-02 | 0.08 |
| S.F.sup. | Length | 0.06 | 2.18E-01 | 0.07 |
| S.F.sup. | MeanDepth | 0.32 | 6.26E-05 | 0.08 |
| S.F.sup. | Width | 0.47 | 4.87E-11 | 0.07 |
| S.F.sup. | SurfaceArea | 0.22 | 1.55E-03 | 0.07 |
| S.Li.ant. | Length | 0.15 | 3.19E-02 | 0.08 |
| S.Li.ant. | MeanDepth | 0.27 | 9.96E-04 | 0.09 |
| S.Li.ant. | Width | 0.18 | 1.53E-02 | 0.08 |

| | | | | |
|---|---|---|---|---|
| S.Li.ant. | SurfaceArea | 0.21 | 8.32E-03 | 0.09 |
| S.Li.post. | Length | 0.11 | 5.16E-02 | 0.07 |
| S.Li.post. | MeanDepth | 0.15 | 2.43E-02 | 0.08 |
| S.Li.post. | Width | 0.20 | 3.53E-03 | 0.07 |
| S.Li.post. | SurfaceArea | 0.15 | 1.48E-02 | 0.07 |
| S.O.p. | Length | - | - | - |
| S.O.p. | MeanDepth | 0.12 | 6.68E-02 | 0.08 |
| S.O.p. | Width | 0.33 | 1.58E-05 | 0.08 |
| S.O.p. | SurfaceArea | - | - | - |
| S.O.T.lat.ant. | Length | 0.12 | 5.05E-02 | 0.07 |
| S.O.T.lat.ant. | MeanDepth | 0.44 | 2.04E-08 | 0.07 |
| S.O.T.lat.ant. | Width | 0.33 | 1.93E-05 | 0.08 |
| S.O.T.lat.ant. | SurfaceArea | 0.27 | 1.25E-04 | 0.07 |
| S.O.T.lat.int. | Length | - | - | - |
| S.O.T.lat.int. | MeanDepth | - | - | - |
| S.O.T.lat.int. | Width | - | - | - |
| S.O.T.lat.int. | SurfaceArea | - | - | - |
| S.O.T.lat.med. | Length | 0.09 | 1.14E-01 | 0.08 |
| S.O.T.lat.med. | MeanDepth | 0.01 | 4.74E-01 | 0.08 |
| S.O.T.lat.med. | Width | 0.02 | 4.13E-01 | 0.08 |
| S.O.T.lat.med. | SurfaceArea | 0.05 | 2.75E-01 | 0.08 |
| S.O.T.lat.post. | Length | - | - | - |
| S.O.T.lat.post. | MeanDepth | 0.21 | 1.90E-03 | 0.07 |
| S.O.T.lat.post. | Width | 0.27 | 2.23E-04 | 0.08 |
| S.O.T.lat.post. | SurfaceArea | 0.06 | 2.02E-01 | 0.08 |
| S.Olf. | Length | 0.17 | 1.82E-02 | 0.08 |
| S.Olf. | MeanDepth | 0.55 | 1.02E-13 | 0.06 |
| S.Olf. | Width | 0.20 | 2.04E-03 | 0.07 |
| S.Olf. | SurfaceArea | 0.39 | 5.00E-07 | 0.08 |
| S.Or. | Length | 0.27 | 1.10E-04 | 0.07 |
| S.Or. | MeanDepth | 0.41 | 8.74E-09 | 0.07 |
| S.Or. | Width | 0.34 | 1.90E-06 | 0.07 |
| S.Or. | SurfaceArea | 0.53 | 1.12E-14 | 0.06 |
| S.p.C. | Length | - | - | - |
| S.p.C. | MeanDepth | 0.18 | 2.96E-02 | 0.09 |
| S.p.C. | Width | 0.12 | 1.03E-01 | 0.09 |
| S.p.C. | SurfaceArea | 0.01 | 4.38E-01 | 0.09 |
| S.Pa.int. | Length | 0.12 | 7.28E-02 | 0.08 |
| S.Pa.int. | MeanDepth | 0.23 | 3.56E-03 | 0.09 |
| S.Pa.int. | Width | 0.34 | 4.30E-06 | 0.08 |
| S.Pa.int. | SurfaceArea | 0.21 | 6.09E-03 | 0.08 |
| S.Pa.sup. | Length | 0.11 | 7.95E-02 | 0.08 |
| S.Pa.sup. | MeanDepth | 0.13 | 5.02E-02 | 0.08 |
| S.Pa.sup. | Width | 0.30 | 2.80E-06 | 0.07 |
| S.Pa.sup. | SurfaceArea | 0.22 | 5.59E-03 | 0.09 |
| S.Pa.t. | Length | - | - | - |

| | | | | |
|---|---|---|---|---|
| S.Pa.t. | MeanDepth | 0.25 | 9.59E-03 | 0.11 |
| S.Pa.t. | Width | 0.36 | 2.40E-04 | 0.10 |
| S.Pa.t. | SurfaceArea | 0.09 | 1.34E-01 | 0.08 |
| S.Pe.C.inf. | Length | 0.03 | 3.64E-01 | 0.09 |
| S.Pe.C.inf. | MeanDepth | 0.25 | 2.69E-03 | 0.09 |
| S.Pe.C.inf. | Width | 0.36 | 2.25E-05 | 0.09 |
| S.Pe.C.inf. | SurfaceArea | 0.09 | 1.55E-01 | 0.09 |
| S.Pe.C.inter. | Length | 0.12 | 5.82E-02 | 0.08 |
| S.Pe.C.inter. | MeanDepth | 0.14 | 3.12E-02 | 0.08 |
| S.Pe.C.inter. | Width | 0.32 | 5.80E-06 | 0.07 |
| S.Pe.C.inter. | SurfaceArea | 0.16 | 1.33E-02 | 0.08 |
| S.Pe.C.marginal. | Length | 0.11 | 9.66E-02 | 0.08 |
| S.Pe.C.marginal. | MeanDepth | 0.10 | 8.94E-02 | 0.08 |
| S.Pe.C.marginal. | Width | 0.31 | 4.86E-05 | 0.08 |
| S.Pe.C.marginal. | SurfaceArea | 0.09 | 1.27E-01 | 0.08 |
| S.Pe.C.median. | Length | 0.18 | 1.28E-02 | 0.08 |
| S.Pe.C.median. | MeanDepth | - | - | - |
| S.Pe.C.median. | Width | 0.31 | 1.86E-04 | 0.09 |
| S.Pe.C.median. | SurfaceArea | 0.13 | 6.43E-02 | 0.09 |
| S.Pe.C.sup. | Length | 0.08 | 1.51E-01 | 0.08 |
| S.Pe.C.sup. | MeanDepth | 0.15 | 3.63E-02 | 0.08 |
| S.Pe.C.sup. | Width | 0.13 | 4.18E-02 | 0.08 |
| S.Pe.C.sup. | SurfaceArea | 0.09 | 1.33E-01 | 0.08 |
| S.Po.C.sup. | Length | 0.06 | 2.09E-01 | 0.08 |
| S.Po.C.sup. | MeanDepth | 0.02 | 3.75E-01 | 0.07 |
| S.Po.C.sup. | Width | 0.24 | 3.40E-04 | 0.07 |
| S.Po.C.sup. | SurfaceArea | 0.05 | 2.40E-01 | 0.08 |
| S.R.inf. | Length | - | - | - |
| S.R.inf. | MeanDepth | - | - | - |
| S.R.inf. | Width | 0.04 | 2.94E-01 | 0.08 |
| S.R.inf. | SurfaceArea | - | - | - |
| S.Rh. | Length | 0.17 | 2.00E-02 | 0.08 |
| S.Rh. | MeanDepth | 0.33 | 5.34E-05 | 0.09 |
| S.Rh. | Width | 0.33 | 5.06E-05 | 0.08 |
| S.Rh. | SurfaceArea | 0.26 | 1.08E-03 | 0.09 |
| S.s.P. | Length | 0.08 | 1.44E-01 | 0.08 |
| S.s.P. | MeanDepth | 0.11 | 7.77E-02 | 0.08 |
| S.s.P. | Width | 0.24 | 5.89E-04 | 0.07 |
| S.s.P. | SurfaceArea | 0.17 | 1.11E-02 | 0.08 |
| S.T.i.ant. | Length | 0.14 | 2.44E-02 | 0.07 |
| S.T.i.ant. | MeanDepth | 0.01 | 4.65E-01 | 0.07 |
| S.T.i.ant. | Width | 0.28 | 1.23E-04 | 0.08 |
| S.T.i.ant. | SurfaceArea | 0.24 | 5.00E-04 | 0.07 |
| S.T.i.post. | Length | 0.03 | 3.44E-01 | 0.07 |
| S.T.i.post. | MeanDepth | 0.14 | 3.28E-02 | 0.08 |
| S.T.i.post. | Width | 0.36 | 4.00E-07 | 0.07 |

| | | | | |
|---|---|---|---|---|
| S.T.i.post. | SurfaceArea | 0.08 | 1.37E-01 | 0.07 |
| S.T.pol. | Length | 0.38 | 1.00E-07 | 0.07 |
| S.T.pol. | MeanDepth | 0.28 | 9.68E-05 | 0.07 |
| S.T.pol. | Width | 0.43 | 2.03E-09 | 0.07 |
| S.T.pol. | SurfaceArea | 0.42 | 2.81E-09 | 0.07 |
| S.T.s. | Length | 0.27 | 8.76E-05 | 0.08 |
| S.T.s. | MeanDepth | 0.34 | 9.00E-07 | 0.07 |
| S.T.s. | Width | 0.49 | 3.32E-11 | 0.07 |
| S.T.s. | SurfaceArea | 0.42 | 2.27E-09 | 0.07 |
| S.T.s.ter.asc.ant. | Length | 0.07 | 1.76E-01 | 0.07 |
| S.T.s.ter.asc.ant. | MeanDepth | 0.09 | 1.34E-01 | 0.08 |
| S.T.s.ter.asc.ant. | Width | 0.12 | 5.46E-02 | 0.08 |
| S.T.s.ter.asc.ant. | SurfaceArea | 0.09 | 8.82E-02 | 0.07 |
| S.T.s.ter.asc.post. | Length | - | - | - |
| S.T.s.ter.asc.post. | MeanDepth | 0.17 | 2.09E-02 | 0.09 |
| S.T.s.ter.asc.post. | Width | 0.25 | 2.52E-04 | 0.07 |
| S.T.s.ter.asc.post. | SurfaceArea | - | - | - |
| Globals | Length | 0.21 | 5.94E-03 | 0.08 |
| Globals | MeanDepth | 0.15 | 1.75E-02 | 0.07 |
| Globals | Width | 0.27 | 6.44E-05 | 0.07 |
| Globals | SurfaceArea | 0.30 | 1.14E-04 | 0.08 |

**Table S15:** heritability eatimates for bilaterally average sulcal descriptors (yellow: bonferroni corrected for 61*4 comparisons) in HCP.

| Sulci | Descriptor | h2 | pval | SE |
|---|---|---|---|---|
| F.C.L.a. | Length | 0.13 | 6.53E-02 | 0.09 |
| F.C.L.a. | MeanDepth | 0.28 | 1.57E-04 | 0.08 |
| F.C.L.a. | Width | 0.40 | 6.00E-06 | 0.09 |
| F.C.L.a. | SurfaceArea | 0.36 | 1.74E-05 | 0.08 |
| F.C.L.p. | Length | 0.12 | 6.94E-02 | 0.08 |
| F.C.L.p. | MeanDepth | 0.26 | 1.40E-03 | 0.09 |
| F.C.L.p. | Width | 0.58 | 3.81E-12 | 0.07 |
| F.C.L.p. | SurfaceArea | 0.60 | 6.54E-16 | 0.06 |
| F.C.L.r.ant. | Length | 0.11 | 1.16E-01 | 0.09 |
| F.C.L.r.ant. | MeanDepth | 0.16 | 8.66E-02 | 0.12 |
| F.C.L.r.ant. | Width | 0.02 | 4.38E-01 | 0.11 |
| F.C.L.r.ant. | SurfaceArea | 0.26 | 2.22E-03 | 0.09 |
| F.C.L.r.asc. | Length | 0.07 | 2.02E-01 | 0.08 |
| F.C.L.r.asc. | MeanDepth | 0.22 | 7.27E-03 | 0.09 |
| F.C.L.r.asc. | Width | 0.25 | 3.36E-03 | 0.09 |
| F.C.L.r.asc. | SurfaceArea | 0.26 | 7.55E-04 | 0.08 |
| F.C.L.r.diag. | Length | - | - | - |
| F.C.L.r.diag. | MeanDepth | - | - | - |
| F.C.L.r.diag. | Width | 0.30 | 8.44E-02 | 0.19 |
| F.C.L.r.diag. | SurfaceArea | - | - | - |
| F.C.L.r.retroC.tr. | Length | 0.17 | 2.81E-02 | 0.09 |
| F.C.L.r.retroC.tr. | MeanDepth | 0.32 | 2.15E-03 | 0.11 |
| F.C.L.r.retroC.tr. | Width | 0.19 | 3.97E-02 | 0.11 |
| F.C.L.r.retroC.tr. | SurfaceArea | 0.24 | 1.12E-02 | 0.10 |
| F.C.L.r.sc.ant. | Length | - | - | - |
| F.C.L.r.sc.ant. | MeanDepth | - | - | - |
| F.C.L.r.sc.ant. | Width | - | - | - |
| F.C.L.r.sc.ant. | SurfaceArea | - | - | - |
| F.C.L.r.sc.post. | Length | - | - | - |
| F.C.L.r.sc.post. | MeanDepth | - | - | - |
| F.C.L.r.sc.post. | Width | - | - | - |
| F.C.L.r.sc.post. | SurfaceArea | 0.03 | 3.92E-01 | 0.11 |
| F.C.M.ant. | Length | 0.06 | 2.11E-01 | 0.08 |
| F.C.M.ant. | MeanDepth | 0.26 | 1.70E-04 | 0.07 |
| F.C.M.ant. | Width | 0.32 | 8.30E-06 | 0.07 |
| F.C.M.ant. | SurfaceArea | 0.18 | 7.11E-03 | 0.08 |
| F.C.M.post. | Length | 0.10 | 1.24E-01 | 0.08 |
| F.C.M.post. | MeanDepth | 0.49 | 3.66E-10 | 0.07 |
| F.C.M.post. | Width | 0.57 | 1.37E-15 | 0.06 |
| F.C.M.post. | SurfaceArea | 0.34 | 8.00E-06 | 0.08 |
| F.Cal.ant.-Sc.Cal. | Length | 0.18 | 9.33E-03 | 0.08 |
| F.Cal.ant.-Sc.Cal. | MeanDepth | 0.57 | 2.05E-13 | 0.06 |
| F.Cal.ant.-Sc.Cal. | Width | 0.50 | 8.23E-10 | 0.07 |

| | | | | |
|---|---|---|---|---|
| F.Cal.ant.-Sc.Cal. | SurfaceArea | 0.61 | 6.27E-14 | 0.06 |
| F.Coll. | Length | 0.11 | 8.60E-02 | 0.08 |
| F.Coll. | MeanDepth | 0.40 | 9.23E-09 | 0.07 |
| F.Coll. | Width | 0.54 | 1.03E-15 | 0.06 |
| F.Coll. | SurfaceArea | 0.24 | 1.59E-03 | 0.08 |
| F.I.P. | Length | 0.30 | 1.11E-04 | 0.08 |
| F.I.P. | MeanDepth | 0.45 | 2.78E-11 | 0.06 |
| F.I.P. | Width | 0.65 | 2.03E-16 | 0.06 |
| F.I.P. | SurfaceArea | 0.34 | 7.70E-06 | 0.08 |
| F.I.P.Po.C.inf. | Length | 0.25 | 1.38E-03 | 0.08 |
| F.I.P.Po.C.inf. | MeanDepth | 0.29 | 1.19E-04 | 0.08 |
| F.I.P.Po.C.inf. | Width | 0.50 | 1.01E-10 | 0.07 |
| F.I.P.Po.C.inf. | SurfaceArea | 0.35 | 8.70E-06 | 0.08 |
| F.I.P.r.int.1 | Length | 0.21 | 4.05E-03 | 0.08 |
| F.I.P.r.int.1 | MeanDepth | 0.11 | 1.04E-01 | 0.09 |
| F.I.P.r.int.1 | Width | 0.16 | 3.01E-02 | 0.09 |
| F.I.P.r.int.1 | SurfaceArea | 0.18 | 1.78E-02 | 0.09 |
| F.I.P.r.int.2 | Length | 0.00 | 5.00E-01 | 0.12 |
| F.I.P.r.int.2 | MeanDepth | - | - | - |
| F.I.P.r.int.2 | Width | 0.08 | 2.78E-01 | 0.13 |
| F.I.P.r.int.2 | SurfaceArea | - | - | - |
| F.P.O. | Length | 0.12 | 4.35E-02 | 0.08 |
| F.P.O. | MeanDepth | 0.52 | 4.24E-12 | 0.06 |
| F.P.O. | Width | 0.48 | 1.87E-11 | 0.06 |
| F.P.O. | SurfaceArea | 0.57 | 1.27E-15 | 0.06 |
| INSULA | Length | - | - | - |
| INSULA | MeanDepth | - | - | - |
| INSULA | Width | 0.21 | 3.71E-01 | 0.64 |
| INSULA | SurfaceArea | 0.43 | 3.10E-06 | 0.09 |
| OCCIPITAL | Length | 0.38 | 5.00E-07 | 0.08 |
| OCCIPITAL | MeanDepth | 0.40 | 3.22E-08 | 0.07 |
| OCCIPITAL | Width | 0.54 | 1.17E-13 | 0.07 |
| OCCIPITAL | SurfaceArea | 0.47 | 1.61E-10 | 0.07 |
| S.C. | Length | 0.16 | 2.72E-02 | 0.08 |
| S.C. | MeanDepth | 0.47 | 5.69E-10 | 0.07 |
| S.C. | Width | 0.80 | 5.31E-31 | 0.03 |
| S.C. | SurfaceArea | 0.47 | 2.66E-09 | 0.07 |
| S.C.LPC. | Length | - | - | - |
| S.C.LPC. | MeanDepth | - | - | - |
| S.C.LPC. | Width | - | - | - |
| S.C.LPC. | SurfaceArea | - | - | - |
| S.C.sylvian. | Length | 0.38 | 7.57E-03 | 0.15 |
| S.C.sylvian. | MeanDepth | 0.21 | 9.66E-02 | 0.16 |
| S.C.sylvian. | Width | 0.45 | 5.89E-03 | 0.16 |
| S.C.sylvian. | SurfaceArea | 0.20 | 6.83E-02 | 0.13 |
| S.Call. | Length | 0.49 | 8.64E-09 | 0.08 |

| | | | | |
|---|---|---|---|---|
| S.Call. | MeanDepth | 0.58 | 1.38E-14 | 0.06 |
| S.Call. | Width | 0.34 | 4.80E-06 | 0.08 |
| S.Call. | SurfaceArea | 0.60 | 8.46E-12 | 0.07 |
| S.Cu. | Length | 0.29 | 9.62E-05 | 0.08 |
| S.Cu. | MeanDepth | 0.25 | 1.57E-03 | 0.08 |
| S.Cu. | Width | 0.23 | 3.62E-03 | 0.09 |
| S.Cu. | SurfaceArea | 0.27 | 3.93E-04 | 0.08 |
| S.F.inf. | Length | 0.14 | 3.57E-02 | 0.08 |
| S.F.inf. | MeanDepth | 0.41 | 8.00E-07 | 0.08 |
| S.F.inf. | Width | 0.58 | 6.32E-15 | 0.06 |
| S.F.inf. | SurfaceArea | 0.36 | 1.60E-06 | 0.07 |
| S.F.inf.ant. | Length | 0.17 | 2.03E-02 | 0.08 |
| S.F.inf.ant. | MeanDepth | 0.22 | 1.80E-03 | 0.08 |
| S.F.inf.ant. | Width | 0.14 | 4.30E-02 | 0.08 |
| S.F.inf.ant. | SurfaceArea | 0.23 | 3.03E-03 | 0.09 |
| S.F.int. | Length | 0.47 | 3.34E-11 | 0.07 |
| S.F.int. | MeanDepth | 0.39 | 1.00E-07 | 0.07 |
| S.F.int. | Width | 0.46 | 1.00E-07 | 0.08 |
| S.F.int. | SurfaceArea | 0.50 | 4.28E-12 | 0.06 |
| S.F.inter. | Length | 0.09 | 1.00E-01 | 0.07 |
| S.F.inter. | MeanDepth | 0.28 | 1.32E-04 | 0.08 |
| S.F.inter. | Width | 0.54 | 6.53E-13 | 0.07 |
| S.F.inter. | SurfaceArea | 0.09 | 1.04E-01 | 0.07 |
| S.F.margi-l. | Length | 0.34 | 2.10E-06 | 0.07 |
| S.F.margi-l. | MeanDepth | 0.12 | 5.07E-02 | 0.07 |
| S.F.margi-l. | Width | 0.22 | 8.38E-04 | 0.07 |
| S.F.margi-l. | SurfaceArea | 0.30 | 2.82E-05 | 0.08 |
| S.F.median. | Length | 0.39 | 2.50E-06 | 0.08 |
| S.F.median. | MeanDepth | 0.22 | 3.18E-03 | 0.08 |
| S.F.median. | Width | 0.40 | 1.00E-07 | 0.07 |
| S.F.median. | SurfaceArea | 0.40 | 1.10E-06 | 0.08 |
| S.F.orbitaire. | Length | 0.12 | 8.14E-02 | 0.08 |
| S.F.orbitaire. | MeanDepth | 0.10 | 1.19E-01 | 0.08 |
| S.F.orbitaire. | Width | 0.14 | 4.16E-02 | 0.08 |
| S.F.orbitaire. | SurfaceArea | 0.12 | 5.81E-02 | 0.08 |
| S.F.polaire.tr. | Length | 0.34 | 1.26E-04 | 0.09 |
| S.F.polaire.tr. | MeanDepth | 0.22 | 2.45E-03 | 0.08 |
| S.F.polaire.tr. | Width | 0.47 | 1.06E-08 | 0.07 |
| S.F.polaire.tr. | SurfaceArea | 0.32 | 1.65E-04 | 0.09 |
| S.F.sup. | Length | 0.09 | 5.00E-01 | 0.09 |
| S.F.sup. | MeanDepth | 0.38 | 5.00E-07 | 0.08 |
| S.F.sup. | Width | 0.61 | 5.44E-16 | 0.06 |
| S.F.sup. | SurfaceArea | 0.24 | 2.05E-03 | 0.08 |
| S.Li.ant. | Length | 0.17 | 2.22E-02 | 0.08 |
| S.Li.ant. | MeanDepth | 0.34 | 6.95E-04 | 0.11 |
| S.Li.ant. | Width | 0.24 | 5.24E-03 | 0.09 |

| | | | | |
|---|---|---|---|---|
| S.Li.ant. | SurfaceArea | 0.20 | 1.34E-02 | 0.09 |
| S.Li.post. | Length | 0.11 | 7.03E-02 | 0.08 |
| S.Li.post. | MeanDepth | 0.26 | 1.07E-03 | 0.09 |
| S.Li.post. | Width | 0.42 | 1.00E-07 | 0.07 |
| S.Li.post. | SurfaceArea | 0.13 | 5.86E-02 | 0.08 |
| S.O.p. | Length | - | - | - |
| S.O.p. | MeanDepth | 0.14 | 1.02E-01 | 0.11 |
| S.O.p. | Width | 0.34 | 1.28E-03 | 0.11 |
| S.O.p. | SurfaceArea | 0.10 | 1.53E-01 | 0.10 |
| S.O.T.lat.ant. | Length | 0.23 | 2.06E-03 | 0.08 |
| S.O.T.lat.ant. | MeanDepth | 0.43 | 1.00E-07 | 0.08 |
| S.O.T.lat.ant. | Width | 0.34 | 5.90E-06 | 0.08 |
| S.O.T.lat.ant. | SurfaceArea | 0.37 | 2.70E-06 | 0.08 |
| S.O.T.lat.int. | Length | - | - | - |
| S.O.T.lat.int. | MeanDepth | - | - | - |
| S.O.T.lat.int. | Width | - | - | - |
| S.O.T.lat.int. | SurfaceArea | 0.01 | 5.00E-01 | 0.12 |
| S.O.T.lat.med. | Length | 0.10 | 1.45E-01 | 0.09 |
| S.O.T.lat.med. | MeanDepth | 0.15 | 5.94E-02 | 0.09 |
| S.O.T.lat.med. | Width | 0.06 | 2.25E-01 | 0.08 |
| S.O.T.lat.med. | SurfaceArea | 0.14 | 6.14E-02 | 0.09 |
| S.O.T.lat.post. | Length | - | - | - |
| S.O.T.lat.post. | MeanDepth | 0.22 | 6.91E-03 | 0.09 |
| S.O.T.lat.post. | Width | 0.17 | 2.37E-02 | 0.09 |
| S.O.T.lat.post. | SurfaceArea | 0.02 | 4.02E-01 | 0.09 |
| S.Olf. | Length | 0.22 | 7.59E-03 | 0.09 |
| S.Olf. | MeanDepth | 0.65 | 6.80E-17 | 0.06 |
| S.Olf. | Width | 0.32 | 3.10E-06 | 0.07 |
| S.Olf. | SurfaceArea | 0.52 | 7.63E-11 | 0.07 |
| S.Or. | Length | 0.39 | 8.00E-07 | 0.08 |
| S.Or. | MeanDepth | 0.44 | 1.00E-07 | 0.08 |
| S.Or. | Width | 0.36 | 9.90E-06 | 0.08 |
| S.Or. | SurfaceArea | 0.56 | 9.10E-13 | 0.07 |
| S.p.C. | Length | - | - | - |
| S.p.C. | MeanDepth | 0.16 | 6.91E-02 | 0.11 |
| S.p.C. | Width | 0.29 | 1.21E-03 | 0.09 |
| S.p.C. | SurfaceArea | - | - | - |
| S.Pa.int. | Length | 0.23 | 3.36E-03 | 0.09 |
| S.Pa.int. | MeanDepth | 0.36 | 2.82E-05 | 0.08 |
| S.Pa.int. | Width | 0.40 | 6.00E-07 | 0.08 |
| S.Pa.int. | SurfaceArea | 0.26 | 1.49E-03 | 0.09 |
| S.Pa.sup. | Length | 0.16 | 3.71E-02 | 0.09 |
| S.Pa.sup. | MeanDepth | 0.21 | 7.33E-03 | 0.09 |
| S.Pa.sup. | Width | 0.39 | 3.20E-06 | 0.08 |
| S.Pa.sup. | SurfaceArea | 0.24 | 2.47E-03 | 0.09 |
| S.Pa.t. | Length | 0.10 | 2.36E-01 | 0.19 |

| | | | | |
|---|---|---|---|---|
| S.Pa.t. | MeanDepth | 0.01 | 4.63E-01 | 0.16 |
| S.Pa.t. | Width | 0.27 | 3.59E-02 | 0.14 |
| S.Pa.t. | SurfaceArea | 0.24 | 1.31E-02 | 0.11 |
| S.Pe.C.inf. | Length | 0.04 | 3.29E-01 | 0.09 |
| S.Pe.C.inf. | MeanDepth | 0.31 | 6.43E-04 | 0.09 |
| S.Pe.C.inf. | Width | 0.40 | 3.50E-06 | 0.08 |
| S.Pe.C.inf. | SurfaceArea | 0.05 | 2.87E-01 | 0.09 |
| S.Pe.C.inter. | Length | 0.16 | 2.49E-02 | 0.08 |
| S.Pe.C.inter. | MeanDepth | 0.25 | 7.36E-04 | 0.08 |
| S.Pe.C.inter. | Width | 0.33 | 2.21E-05 | 0.08 |
| S.Pe.C.inter. | SurfaceArea | 0.27 | 3.66E-04 | 0.08 |
| S.Pe.C.margi-l. | Length | 0.04 | 3.43E-01 | 0.10 |
| S.Pe.C.margi-l. | MeanDepth | 0.24 | 7.14E-03 | 0.10 |
| S.Pe.C.margi-l. | Width | 0.37 | 1.45E-05 | 0.08 |
| S.Pe.C.margi-l. | SurfaceArea | 0.08 | 2.01E-01 | 0.10 |
| S.Pe.C.median. | Length | 0.09 | 1.38E-01 | 0.09 |
| S.Pe.C.median. | MeanDepth | 0.04 | 3.47E-01 | 0.10 |
| S.Pe.C.median. | Width | 0.38 | 2.70E-05 | 0.09 |
| S.Pe.C.median. | SurfaceArea | 0.17 | 3.60E-02 | 0.10 |
| S.Pe.C.sup. | Length | 0.22 | 5.00E-03 | 0.09 |
| S.Pe.C.sup. | MeanDepth | 0.15 | 5.05E-02 | 0.09 |
| S.Pe.C.sup. | Width | 0.39 | 9.90E-06 | 0.09 |
| S.Pe.C.sup. | SurfaceArea | 0.24 | 4.78E-03 | 0.09 |
| S.Po.C.sup. | Length | 0.25 | 1.79E-03 | 0.09 |
| S.Po.C.sup. | MeanDepth | 0.20 | 2.27E-03 | 0.07 |
| S.Po.C.sup. | Width | 0.41 | 1.53E-08 | 0.07 |
| S.Po.C.sup. | SurfaceArea | 0.25 | 1.13E-03 | 0.08 |
| S.R.inf. | Length | 0.34 | 9.72E-05 | 0.09 |
| S.R.inf. | MeanDepth | 0.12 | 9.08E-02 | 0.09 |
| S.R.inf. | Width | 0.17 | 4.17E-02 | 0.10 |
| S.R.inf. | SurfaceArea | 0.36 | 4.06E-05 | 0.09 |
| S.Rh. | Length | 0.21 | 5.07E-03 | 0.08 |
| S.Rh. | MeanDepth | 0.34 | 6.95E-05 | 0.09 |
| S.Rh. | Width | 0.37 | 5.00E-06 | 0.08 |
| S.Rh. | SurfaceArea | 0.28 | 4.01E-04 | 0.08 |
| S.s.P. | Length | 0.12 | 5.45E-02 | 0.08 |
| S.s.P. | MeanDepth | 0.39 | 9.00E-07 | 0.08 |
| S.s.P. | Width | 0.37 | 9.00E-07 | 0.07 |
| S.s.P. | SurfaceArea | 0.23 | 2.28E-03 | 0.08 |
| S.T.i.ant. | Length | 0.17 | 1.03E-02 | 0.07 |
| S.T.i.ant. | MeanDepth | 0.18 | 1.24E-02 | 0.08 |
| S.T.i.ant. | Width | 0.41 | 2.00E-07 | 0.08 |
| S.T.i.ant. | SurfaceArea | 0.31 | 2.65E-05 | 0.08 |
| S.T.i.post. | Length | 0.02 | 3.68E-01 | 0.07 |
| S.T.i.post. | MeanDepth | 0.33 | 1.80E-04 | 0.09 |
| S.T.i.post. | Width | 0.48 | 3.26E-08 | 0.08 |

| | | | | |
|---|---|---|---|---|
| S.T.i.post. | SurfaceArea | 0.14 | 4.18E-02 | 0.08 |
| S.T.pol. | Length | 0.45 | 1.68E-09 | 0.07 |
| S.T.pol. | MeanDepth | 0.24 | 8.33E-04 | 0.08 |
| S.T.pol. | Width | 0.38 | 9.00E-07 | 0.08 |
| S.T.pol. | SurfaceArea | 0.51 | 1.07E-11 | 0.07 |
| S.T.s. | Length | 0.16 | 1.36E-02 | 0.08 |
| S.T.s. | MeanDepth | 0.52 | 6.95E-13 | 0.06 |
| S.T.s. | Width | 0.46 | 9.14E-11 | 0.07 |
| S.T.s. | SurfaceArea | 0.29 | 9.75E-05 | 0.08 |
| S.T.s.ter.asc.ant. | Length | 0.14 | 3.69E-02 | 0.08 |
| S.T.s.ter.asc.ant. | MeanDepth | 0.17 | 1.79E-02 | 0.08 |
| S.T.s.ter.asc.ant. | Width | 0.33 | 1.79E-05 | 0.08 |
| S.T.s.ter.asc.ant. | SurfaceArea | 0.10 | 7.81E-02 | 0.07 |
| S.T.s.ter.asc.post. | Length | 0.01 | 4.38E-01 | 0.08 |
| S.T.s.ter.asc.post. | MeanDepth | 0.16 | 2.30E-02 | 0.08 |
| S.T.s.ter.asc.post. | Width | 0.45 | 1.22E-08 | 0.08 |
| S.T.s.ter.asc.post. | SurfaceArea | 0.17 | 1.67E-02 | 0.08 |
| Globals | Length | 0.26 | 1.74E-03 | 0.09 |
| Globals | MeanDepth | 0.19 | 8.24E-03 | 0.08 |
| Globals | Width | 0.25 | 2.70E-04 | 0.07 |
| Globals | SurfaceArea | 0.33 | 1.62E-04 | 0.09 |

**Table S16:** heritability eatimates for left hemisphere sulcal descriptors (yellow: bonferroni corrected for 123*4 comparisons) in GOBS.

| Sulci | Descriptor | h2 | pval | SE |
|---|---|---|---|---|
| F.C.L.a. | Length | 0.09 | 5.18E-02 | 0.06 |
| F.C.L.a. | MeanDepth | 0.13 | 1.55E-02 | 0.07 |
| F.C.L.a. | Width | 0.31 | 3.50E-06 | 0.07 |
| F.C.L.a. | SurfaceArea | 0.17 | 2.20E-03 | 0.07 |
| F.C.L.p. | Length | 0.13 | 1.90E-02 | 0.07 |
| F.C.L.p. | MeanDepth | 0.07 | 9.71E-02 | 0.06 |
| F.C.L.p. | Width | 0.32 | 3.47E-10 | 0.06 |
| F.C.L.p. | SurfaceArea | 0.35 | 2.48E-08 | 0.07 |
| F.C.L.r.ant. | Length | - | - | - |
| F.C.L.r.ant. | MeanDepth | 0.17 | 1.45E-03 | 0.06 |
| F.C.L.r.ant. | Width | 0.07 | 1.24E-01 | 0.06 |
| F.C.L.r.ant. | SurfaceArea | 0.07 | 1.04E-01 | 0.06 |
| F.C.L.r.asc. | Length | - | - | - |
| F.C.L.r.asc. | MeanDepth | 0.09 | 4.73E-02 | 0.06 |
| F.C.L.r.asc. | Width | 0.12 | 1.25E-02 | 0.06 |
| F.C.L.r.asc. | SurfaceArea | 0.18 | 4.69E-04 | 0.06 |
| F.C.L.r.diag. | Length | 0.01 | 4.75E-01 | 0.12 |
| F.C.L.r.diag. | MeanDepth | 0.02 | 4.31E-01 | 0.13 |
| F.C.L.r.diag. | Width | 0.13 | 1.53E-01 | 0.13 |
| F.C.L.r.diag. | SurfaceArea | - | - | - |
| F.C.L.r.retroC.tr. | Length | 0.09 | 6.49E-02 | 0.06 |
| F.C.L.r.retroC.tr. | MeanDepth | 0.16 | 4.18E-03 | 0.07 |
| F.C.L.r.retroC.tr. | Width | 0.13 | 9.28E-03 | 0.06 |
| F.C.L.r.retroC.tr. | SurfaceArea | 0.12 | 2.04E-02 | 0.07 |
| F.C.L.r.sc.ant. | Length | - | - | - |
| F.C.L.r.sc.ant. | MeanDepth | - | - | - |
| F.C.L.r.sc.ant. | Width | 0.74 | 9.08E-02 | 0.56 |
| F.C.L.r.sc.ant. | SurfaceArea | - | - | - |
| F.C.L.r.sc.post. | Length | 0.50 | 1.47E-01 | 0.46 |
| F.C.L.r.sc.post. | MeanDepth | 0.03 | 3.03E-01 | 0.07 |
| F.C.L.r.sc.post. | Width | 0.03 | 3.51E-01 | 0.08 |
| F.C.L.r.sc.post. | SurfaceArea | 0.03 | 3.22E-01 | 0.07 |
| F.C.M.ant. | Length | 0.10 | 3.70E-02 | 0.06 |
| F.C.M.ant. | MeanDepth | 0.33 | 1.13E-08 | 0.07 |
| F.C.M.ant. | Width | 0.29 | 4.00E-07 | 0.07 |
| F.C.M.ant. | SurfaceArea | 0.18 | 9.57E-04 | 0.07 |
| F.C.M.post. | Length | 0.04 | 1.76E-01 | 0.05 |
| F.C.M.post. | MeanDepth | 0.27 | 2.00E-07 | 0.06 |
| F.C.M.post. | Width | 0.19 | 3.05E-04 | 0.06 |
| F.C.M.post. | SurfaceArea | 0.18 | 3.09E-04 | 0.06 |
| F.Cal.ant.-Sc.Cal. | Length | - | - | - |
| F.Cal.ant.-Sc.Cal. | MeanDepth | - | - | - |
| F.Cal.ant.-Sc.Cal. | Width | - | - | - |

| | | | | |
|---|---|---|---|---|
| F.Cal.ant.-Sc.Cal. | SurfaceArea | - | - | - |
| F.Coll. | Length | 0.02 | 3.54E-01 | 0.05 |
| F.Coll. | MeanDepth | 0.38 | 4.68E-12 | 0.07 |
| F.Coll. | Width | 0.25 | 9.50E-06 | 0.07 |
| F.Coll. | SurfaceArea | 0.18 | 1.71E-04 | 0.06 |
| F.I.P. | Length | 0.12 | 1.50E-02 | 0.06 |
| F.I.P. | MeanDepth | 0.21 | 1.62E-04 | 0.07 |
| F.I.P. | Width | 0.37 | 1.19E-10 | 0.07 |
| F.I.P. | SurfaceArea | 0.24 | 8.00E-06 | 0.07 |
| F.I.P.Po.C.inf. | Length | 0.08 | 5.19E-02 | 0.06 |
| F.I.P.Po.C.inf. | MeanDepth | 0.16 | 2.08E-03 | 0.06 |
| F.I.P.Po.C.inf. | Width | 0.23 | 3.08E-05 | 0.07 |
| F.I.P.Po.C.inf. | SurfaceArea | 0.11 | 1.68E-02 | 0.06 |
| F.I.P.r.int.1 | Length | 0.11 | 3.81E-02 | 0.07 |
| F.I.P.r.int.1 | MeanDepth | - | - | - |
| F.I.P.r.int.1 | Width | - | - | - |
| F.I.P.r.int.1 | SurfaceArea | 0.15 | 5.34E-03 | 0.07 |
| F.I.P.r.int.2 | Length | 0.11 | 3.81E-02 | 0.07 |
| F.I.P.r.int.2 | MeanDepth | - | - | - |
| F.I.P.r.int.2 | Width | - | - | - |
| F.I.P.r.int.2 | SurfaceArea | 0.15 | 5.34E-03 | 0.07 |
| F.P.O. | Length | 0.08 | 5.00E-02 | 0.06 |
| F.P.O. | MeanDepth | 0.28 | 6.00E-07 | 0.07 |
| F.P.O. | Width | 0.41 | 3.88E-14 | 0.07 |
| F.P.O. | SurfaceArea | 0.33 | 6.66E-09 | 0.07 |
| INSULA | Length | 0.15 | 3.71E-01 | 0.48 |
| INSULA | MeanDepth | 0.09 | 5.80E-02 | 0.07 |
| INSULA | Width | 0.22 | 7.23E-05 | 0.07 |
| INSULA | SurfaceArea | 0.27 | 3.10E-06 | 0.07 |
| OCCIPITAL | Length | 0.18 | 4.36E-04 | 0.06 |
| OCCIPITAL | MeanDepth | 0.26 | 1.20E-06 | 0.07 |
| OCCIPITAL | Width | 0.35 | 7.70E-10 | 0.07 |
| OCCIPITAL | SurfaceArea | 0.23 | 1.22E-05 | 0.06 |
| S.C. | Length | 0.09 | 3.81E-02 | 0.06 |
| S.C. | MeanDepth | 0.30 | 3.00E-07 | 0.07 |
| S.C. | Width | 0.39 | 3.10E-11 | 0.07 |
| S.C. | SurfaceArea | 0.23 | 2.73E-05 | 0.07 |
| S.C.LPC. | Length | 0.06 | 2.90E-01 | 0.11 |
| S.C.LPC. | MeanDepth | - | - | - |
| S.C.LPC. | Width | 0.08 | 2.63E-01 | 0.13 |
| S.C.LPC. | SurfaceArea | 0.08 | 2.51E-01 | 0.12 |
| S.C.sylvian. | Length | - | - | - |
| S.C.sylvian. | MeanDepth | 0.14 | 2.84E-02 | 0.08 |
| S.C.sylvian. | Width | 0.04 | 3.10E-01 | 0.07 |
| S.C.sylvian. | SurfaceArea | 0.05 | 2.76E-01 | 0.08 |
| S.Call. | Length | 0.39 | 3.44E-10 | 0.07 |

| | | | | |
|---|---|---|---|---|
| S.Call. | MeanDepth | 0.32 | 1.00E-07 | 0.07 |
| S.Call. | Width | 0.27 | 1.90E-06 | 0.07 |
| S.Call. | SurfaceArea | 0.43 | 8.56E-13 | 0.07 |
| S.Cu. | Length | - | - | - |
| S.Cu. | MeanDepth | 0.19 | 4.96E-04 | 0.06 |
| S.Cu. | Width | 0.15 | 8.16E-03 | 0.07 |
| S.Cu. | SurfaceArea | 0.03 | 3.07E-01 | 0.06 |
| S.F.inf. | Length | 0.14 | 8.18E-03 | 0.07 |
| S.F.inf. | MeanDepth | 0.32 | 4.13E-08 | 0.07 |
| S.F.inf. | Width | 0.24 | 1.27E-05 | 0.07 |
| S.F.inf. | SurfaceArea | 0.22 | 8.43E-05 | 0.07 |
| S.F.inf.ant. | Length | 0.10 | 4.14E-02 | 0.06 |
| S.F.inf.ant. | MeanDepth | 0.06 | 1.68E-01 | 0.06 |
| S.F.inf.ant. | Width | 0.10 | 5.84E-02 | 0.07 |
| S.F.inf.ant. | SurfaceArea | 0.11 | 2.72E-02 | 0.06 |
| S.F.int. | Length | 0.14 | 2.85E-03 | 0.06 |
| S.F.int. | MeanDepth | 0.26 | 2.60E-06 | 0.07 |
| S.F.int. | Width | 0.23 | 4.60E-06 | 0.06 |
| S.F.int. | SurfaceArea | 0.22 | 1.20E-05 | 0.06 |
| S.F.inter. | Length | 0.04 | 2.21E-01 | 0.05 |
| S.F.inter. | MeanDepth | 0.09 | 3.99E-02 | 0.06 |
| S.F.inter. | Width | 0.28 | 1.00E-06 | 0.07 |
| S.F.inter. | SurfaceArea | 0.06 | 1.02E-01 | 0.05 |
| S.F.marginal. | Length | 0.01 | 4.28E-01 | 0.05 |
| S.F.marginal. | MeanDepth | 0.04 | 2.48E-01 | 0.06 |
| S.F.marginal. | Width | 0.09 | 4.00E-02 | 0.06 |
| S.F.marginal. | SurfaceArea | 0.07 | 7.93E-02 | 0.05 |
| S.F.median. | Length | 0.12 | 1.05E-02 | 0.06 |
| S.F.median. | MeanDepth | - | - | - |
| S.F.median. | Width | 0.12 | 7.69E-03 | 0.06 |
| S.F.median. | SurfaceArea | 0.11 | 1.80E-02 | 0.06 |
| S.F.orbitaire. | Length | 0.04 | 2.21E-01 | 0.06 |
| S.F.orbitaire. | MeanDepth | 0.02 | 3.48E-01 | 0.06 |
| S.F.orbitaire. | Width | 0.14 | 7.15E-03 | 0.07 |
| S.F.orbitaire. | SurfaceArea | 0.06 | 1.34E-01 | 0.06 |
| S.F.polaire.tr. | Length | 0.13 | 1.15E-02 | 0.06 |
| S.F.polaire.tr. | MeanDepth | 0.00 | 4.66E-01 | 0.05 |
| S.F.polaire.tr. | Width | 0.15 | 4.77E-03 | 0.07 |
| S.F.polaire.tr. | SurfaceArea | 0.08 | 7.31E-02 | 0.06 |
| S.F.sup. | Length | 0.08 | 7.75E-02 | 0.06 |
| S.F.sup. | MeanDepth | 0.24 | 6.80E-06 | 0.06 |
| S.F.sup. | Width | 0.25 | 1.00E-06 | 0.06 |
| S.F.sup. | SurfaceArea | 0.12 | 1.33E-02 | 0.06 |
| S.Li.ant. | Length | 0.20 | 8.97E-04 | 0.07 |
| S.Li.ant. | MeanDepth | 0.11 | 5.28E-02 | 0.08 |
| S.Li.ant. | Width | 0.05 | 2.12E-01 | 0.07 |

| | | | | |
|---|---|---|---|---|
| S.Li.ant. | SurfaceArea | 0.22 | 1.03E-03 | 0.08 |
| S.Li.post. | Length | - | - | - |
| S.Li.post. | MeanDepth | 0.01 | 4.01E-01 | 0.05 |
| S.Li.post. | Width | 0.05 | 1.92E-01 | 0.06 |
| S.Li.post. | SurfaceArea | - | - | - |
| S.O.p. | Length | 0.10 | 6.98E-02 | 0.07 |
| S.O.p. | MeanDepth | 0.06 | 1.76E-01 | 0.07 |
| S.O.p. | Width | 0.06 | 1.95E-01 | 0.07 |
| S.O.p. | SurfaceArea | 0.13 | 2.25E-02 | 0.07 |
| S.O.T.lat.ant. | Length | 0.25 | 1.34E-05 | 0.07 |
| S.O.T.lat.ant. | MeanDepth | 0.22 | 7.45E-05 | 0.07 |
| S.O.T.lat.ant. | Width | 0.18 | 2.41E-04 | 0.06 |
| S.O.T.lat.ant. | SurfaceArea | 0.29 | 5.00E-07 | 0.07 |
| S.O.T.lat.int. | Length | 0.07 | 1.55E-01 | 0.07 |
| S.O.T.lat.int. | MeanDepth | 0.18 | 3.35E-03 | 0.07 |
| S.O.T.lat.int. | Width | 0.17 | 1.30E-02 | 0.08 |
| S.O.T.lat.int. | SurfaceArea | 0.08 | 1.60E-01 | 0.08 |
| S.O.T.lat.med. | Length | 0.03 | 2.97E-01 | 0.05 |
| S.O.T.lat.med. | MeanDepth | 0.06 | 1.05E-01 | 0.05 |
| S.O.T.lat.med. | Width | 0.06 | 1.34E-01 | 0.05 |
| S.O.T.lat.med. | SurfaceArea | 0.01 | 3.91E-01 | 0.05 |
| S.O.T.lat.post. | Length | 0.01 | 4.04E-01 | 0.05 |
| S.O.T.lat.post. | MeanDepth | 0.00 | 4.79E-01 | 0.06 |
| S.O.T.lat.post. | Width | 0.07 | 9.35E-02 | 0.06 |
| S.O.T.lat.post. | SurfaceArea | 0.06 | 1.17E-01 | 0.06 |
| S.Olf. | Length | 0.35 | 3.36E-10 | 0.07 |
| S.Olf. | MeanDepth | 0.42 | 1.54E-14 | 0.06 |
| S.Olf. | Width | 0.28 | 5.00E-06 | 0.07 |
| S.Olf. | SurfaceArea | 0.46 | 2.25E-14 | 0.07 |
| S.Or. | Length | 0.08 | 8.32E-02 | 0.06 |
| S.Or. | MeanDepth | 0.23 | 2.67E-05 | 0.07 |
| S.Or. | Width | 0.21 | 7.53E-04 | 0.07 |
| S.Or. | SurfaceArea | 0.11 | 2.95E-02 | 0.06 |
| S.p.C. | Length | 0.06 | 1.80E-01 | 0.07 |
| S.p.C. | MeanDepth | 0.08 | 9.61E-02 | 0.06 |
| S.p.C. | Width | 0.05 | 1.87E-01 | 0.06 |
| S.p.C. | SurfaceArea | 0.04 | 2.70E-01 | 0.07 |
| S.Pa.int. | Length | 0.11 | 3.58E-02 | 0.07 |
| S.Pa.int. | MeanDepth | 0.18 | 1.48E-03 | 0.07 |
| S.Pa.int. | Width | 0.20 | 2.15E-04 | 0.07 |
| S.Pa.int. | SurfaceArea | 0.11 | 3.10E-02 | 0.07 |
| S.Pa.sup. | Length | 0.04 | 3.03E-01 | 0.07 |
| S.Pa.sup. | MeanDepth | - | - | - |
| S.Pa.sup. | Width | 0.16 | 8.37E-03 | 0.08 |
| S.Pa.sup. | SurfaceArea | 0.11 | 6.69E-02 | 0.08 |
| S.Pa.t. | Length | 0.06 | 2.53E-01 | 0.10 |

| | | | | |
|---|---|---|---|---|
| S.Pa.t. | MeanDepth | 0.02 | 4.13E-01 | 0.08 |
| S.Pa.t. | Width | 0.09 | 1.07E-01 | 0.08 |
| S.Pa.t. | SurfaceArea | - | - | - |
| S.Pe.C.inf. | Length | 0.08 | 4.03E-02 | 0.05 |
| S.Pe.C.inf. | MeanDepth | 0.06 | 1.33E-01 | 0.06 |
| S.Pe.C.inf. | Width | 0.26 | 9.60E-06 | 0.07 |
| S.Pe.C.inf. | SurfaceArea | 0.05 | 1.48E-01 | 0.05 |
| S.Pe.C.inter. | Length | 0.01 | 4.43E-01 | 0.05 |
| S.Pe.C.inter. | MeanDepth | 0.12 | 1.24E-02 | 0.06 |
| S.Pe.C.inter. | Width | 0.18 | 2.94E-04 | 0.06 |
| S.Pe.C.inter. | SurfaceArea | 0.03 | 2.92E-01 | 0.06 |
| S.Pe.C.marginal. | Length | - | - | - |
| S.Pe.C.marginal. | MeanDepth | 0.09 | 9.43E-02 | 0.07 |
| S.Pe.C.marginal. | Width | 0.10 | 6.88E-02 | 0.08 |
| S.Pe.C.marginal. | SurfaceArea | 0.13 | 2.35E-02 | 0.07 |
| S.Pe.C.median. | Length | 0.00 | 4.68E-01 | 0.06 |
| S.Pe.C.median. | MeanDepth | 0.02 | 3.78E-01 | 0.07 |
| S.Pe.C.median. | Width | 0.12 | 1.89E-02 | 0.06 |
| S.Pe.C.median. | SurfaceArea | - | - | - |
| S.Pe.C.sup. | Length | 0.20 | 5.78E-04 | 0.07 |
| S.Pe.C.sup. | MeanDepth | 0.13 | 2.51E-02 | 0.07 |
| S.Pe.C.sup. | Width | 0.13 | 6.19E-03 | 0.06 |
| S.Pe.C.sup. | SurfaceArea | 0.19 | 1.34E-03 | 0.07 |
| S.Po.C.sup. | Length | - | - | - |
| S.Po.C.sup. | MeanDepth | 0.16 | 7.00E-03 | 0.07 |
| S.Po.C.sup. | Width | 0.24 | 1.63E-05 | 0.07 |
| S.Po.C.sup. | SurfaceArea | 0.15 | 7.55E-03 | 0.07 |
| S.R.inf. | Length | - | - | - |
| S.R.inf. | MeanDepth | - | - | - |
| S.R.inf. | Width | - | - | - |
| S.R.inf. | SurfaceArea | - | - | - |
| S.Rh. | Length | 0.30 | 5.80E-06 | 0.08 |
| S.Rh. | MeanDepth | 0.12 | 2.20E-02 | 0.07 |
| S.Rh. | Width | 0.16 | 4.68E-03 | 0.07 |
| S.Rh. | SurfaceArea | 0.20 | 1.11E-03 | 0.07 |
| S.s.P. | Length | 0.10 | 4.39E-02 | 0.06 |
| S.s.P. | MeanDepth | 0.18 | 5.38E-04 | 0.07 |
| S.s.P. | Width | 0.25 | 3.30E-06 | 0.06 |
| S.s.P. | SurfaceArea | 0.17 | 3.20E-03 | 0.07 |
| S.T.i.ant. | Length | 0.05 | 1.66E-01 | 0.06 |
| S.T.i.ant. | MeanDepth | 0.30 | 1.00E-07 | 0.07 |
| S.T.i.ant. | Width | 0.29 | 1.00E-07 | 0.06 |
| S.T.i.ant. | SurfaceArea | 0.13 | 6.01E-03 | 0.06 |
| S.T.i.post. | Length | 0.09 | 6.45E-02 | 0.06 |
| S.T.i.post. | MeanDepth | 0.15 | 1.43E-03 | 0.06 |
| S.T.i.post. | Width | 0.25 | 3.50E-06 | 0.07 |

| | | | | |
|---|---|---|---|---|
| S.T.i.post. | SurfaceArea | 0.21 | 6.52E-04 | 0.08 |
| S.T.pol. | Length | 0.18 | 5.37E-04 | 0.06 |
| S.T.pol. | MeanDepth | 0.18 | 1.14E-03 | 0.07 |
| S.T.pol. | Width | 0.24 | 1.11E-05 | 0.07 |
| S.T.pol. | SurfaceArea | 0.23 | 2.70E-05 | 0.07 |
| S.T.s. | Length | 0.09 | 4.24E-02 | 0.06 |
| S.T.s. | MeanDepth | 0.32 | 1.31E-09 | 0.06 |
| S.T.s. | Width | 0.28 | 4.00E-07 | 0.07 |
| S.T.s. | SurfaceArea | 0.03 | 2.89E-01 | 0.06 |
| S.T.s.ter.asc.ant. | Length | - | - | - |
| S.T.s.ter.asc.ant. | MeanDepth | - | - | - |
| S.T.s.ter.asc.ant. | Width | 0.09 | 4.25E-02 | 0.06 |
| S.T.s.ter.asc.ant. | SurfaceArea | - | - | - |
| S.T.s.ter.asc.post. | Length | - | - | - |
| S.T.s.ter.asc.post. | MeanDepth | 0.10 | 5.72E-02 | 0.07 |
| S.T.s.ter.asc.post. | Width | 0.14 | 7.81E-03 | 0.06 |
| S.T.s.ter.asc.post. | SurfaceArea | 0.04 | 2.72E-01 | 0.06 |
| Globals | Length | 0.16 | 1.77E-03 | 0.06 |
| Globals | MeanDepth | 0.14 | 1.53E-02 | 0.07 |
| Globals | Width | 0.22 | 2.53E-05 | 0.07 |
| Globals | SurfaceArea | 0.24 | 7.80E-06 | 0.07 |

**Table S17:** heritability eatimates for right hemisphere sulcal descriptors (yellow: bonferroni corrected for 123*4 comparisons) in GOBS.

| Sulci | Descriptor | h2 | pval | SE |
|---|---|---|---|---|
| F.C.L.a. | Length | 0.03 | 3.00E-01 | 0.06 |
| F.C.L.a. | MeanDepth | 0.07 | 1.20E-01 | 0.06 |
| F.C.L.a. | SurfaceArea | 0.34 | 1.00E-07 | 0.08 |
| F.C.L.a. | Width | 0.35 | 1.00E-07 | 0.08 |
| F.C.L.p. | Length | - | - | - |
| F.C.L.p. | MeanDepth | - | 4.80E-01 | 0.06 |
| F.C.L.p. | SurfaceArea | 0.21 | 7.10E-04 | 0.08 |
| F.C.L.p. | Width | 0.33 | 1.30E-10 | 0.06 |
| F.C.L.r.ant. | Length | 0.13 | 3.70E-02 | 0.08 |
| F.C.L.r.ant. | MeanDepth | 0.09 | 6.00E-02 | 0.07 |
| F.C.L.r.ant. | SurfaceArea | 0.15 | 7.50E-03 | 0.07 |
| F.C.L.r.ant. | Width | 0.14 | 2.10E-02 | 0.08 |
| F.C.L.r.asc. | Length | 0.17 | 1.90E-03 | 0.07 |
| F.C.L.r.asc. | MeanDepth | 0.09 | 4.30E-02 | 0.06 |
| F.C.L.r.asc. | SurfaceArea | 0.21 | 8.90E-05 | 0.07 |
| F.C.L.r.asc. | Width | 0.16 | 7.30E-04 | 0.06 |
| F.C.L.r.diag. | Length | 0.02 | 4.20E-01 | 0.09 |
| F.C.L.r.diag. | MeanDepth | - | - | - |
| F.C.L.r.diag. | SurfaceArea | - | - | - |
| F.C.L.r.diag. | Width | 0.11 | 1.20E-01 | 0.10 |
| F.C.L.r.retroC.tr. | Length | 0.08 | 7.90E-02 | 0.06 |
| F.C.L.r.retroC.tr. | MeanDepth | 0.14 | 4.50E-03 | 0.06 |
| F.C.L.r.retroC.tr. | SurfaceArea | 0.11 | 2.90E-02 | 0.06 |
| F.C.L.r.retroC.tr. | Width | 0.09 | 6.10E-02 | 0.06 |
| F.C.L.r.sc.ant. | Length | - | - | - |
| F.C.L.r.sc.ant. | MeanDepth | 0.13 | 3.20E-01 | 0.28 |
| F.C.L.r.sc.ant. | SurfaceArea | - | - | - |
| F.C.L.r.sc.ant. | Width | 0.41 | 1.10E-01 | 0.34 |
| F.C.L.r.sc.post. | Length | - | - | - |
| F.C.L.r.sc.post. | MeanDepth | - | 4.70E-01 | 0.06 |
| F.C.L.r.sc.post. | SurfaceArea | 0.05 | 2.20E-01 | 0.07 |
| F.C.L.r.sc.post. | Width | 0.12 | 3.10E-02 | 0.07 |
| F.C.M.ant. | Length | 0.14 | 3.80E-03 | 0.06 |
| F.C.M.ant. | MeanDepth | 0.13 | 4.20E-03 | 0.06 |
| F.C.M.ant. | SurfaceArea | 0.15 | 1.80E-03 | 0.06 |
| F.C.M.ant. | Width | 0.17 | 4.40E-04 | 0.06 |
| F.C.M.post. | Length | - | - | - |
| F.C.M.post. | MeanDepth | 0.25 | 4.60E-06 | 0.07 |
| F.C.M.post. | SurfaceArea | 0.09 | 4.80E-02 | 0.06 |
| F.C.M.post. | Width | 0.23 | 1.00E-05 | 0.06 |
| F.Cal.ant.-Sc.Cal. | Length | - | - | - |
| F.Cal.ant.-Sc.Cal. | MeanDepth | - | - | - |
| F.Cal.ant.-Sc.Cal. | SurfaceArea | - | - | - |

| | | | | |
|---|---|---|---|---|
| F.Cal.ant.-Sc.Cal. | Width | - | - | - |
| F.Coll. | Length | 0.06 | 1.10E-01 | 0.06 |
| F.Coll. | MeanDepth | 0.52 | 1.60E-17 | 0.07 |
| F.Coll. | SurfaceArea | 0.27 | 2.80E-06 | 0.07 |
| F.Coll. | Width | 0.26 | 5.00E-07 | 0.06 |
| F.I.P. | Length | 0.16 | 1.70E-03 | 0.06 |
| F.I.P. | MeanDepth | 0.13 | 6.90E-03 | 0.06 |
| F.I.P. | SurfaceArea | 0.17 | 1.30E-03 | 0.06 |
| F.I.P. | Width | 0.31 | 2.70E-08 | 0.07 |
| F.I.P.Po.C.inf. | Length | 0.07 | 9.10E-02 | 0.06 |
| F.I.P.Po.C.inf. | MeanDepth | 0.08 | 7.50E-02 | 0.06 |
| F.I.P.Po.C.inf. | SurfaceArea | 0.09 | 3.80E-02 | 0.06 |
| F.I.P.Po.C.inf. | Width | 0.23 | 4.40E-05 | 0.07 |
| F.I.P.r.int.1 | Length | - | - | - |
| F.I.P.r.int.1 | MeanDepth | - | - | - |
| F.I.P.r.int.1 | SurfaceArea | - | - | - |
| F.I.P.r.int.1 | Width | - | - | - |
| F.I.P.r.int.2 | Length | - | - | - |
| F.I.P.r.int.2 | MeanDepth | - | - | - |
| F.I.P.r.int.2 | SurfaceArea | - | - | - |
| F.I.P.r.int.2 | Width | - | - | - |
| F.P.O. | Length | 0.10 | 4.60E-02 | 0.06 |
| F.P.O. | MeanDepth | 0.27 | 1.50E-06 | 0.07 |
| F.P.O. | SurfaceArea | 0.24 | 3.90E-05 | 0.07 |
| F.P.O. | Width | 0.24 | 4.60E-06 | 0.06 |
| INSULA | Length | - | - | - |
| INSULA | MeanDepth | 0.18 | 6.70E-04 | 0.07 |
| INSULA | SurfaceArea | 0.27 | 5.20E-06 | 0.07 |
| INSULA | Width | 0.27 | 1.10E-05 | 0.07 |
| OCCIPITAL | Length | 0.27 | 2.60E-06 | 0.07 |
| OCCIPITAL | MeanDepth | 0.16 | 9.70E-04 | 0.06 |
| OCCIPITAL | SurfaceArea | 0.30 | 1.40E-06 | 0.07 |
| OCCIPITAL | Width | 0.35 | 1.60E-10 | 0.07 |
| S.C. | Length | 0.18 | 1.70E-03 | 0.07 |
| S.C. | MeanDepth | 0.25 | 1.50E-06 | 0.06 |
| S.C. | SurfaceArea | 0.30 | 1.10E-06 | 0.07 |
| S.C. | Width | 0.40 | 1.00E-12 | 0.07 |
| S.C.LPC. | Length | - | - | - |
| S.C.LPC. | MeanDepth | - | - | - |
| S.C.LPC. | SurfaceArea | - | - | - |
| S.C.LPC. | Width | - | - | - |
| S.C.sylvian. | Length | 0.12 | 9.20E-02 | 0.09 |
| S.C.sylvian. | MeanDepth | 0.21 | 6.40E-03 | 0.10 |
| S.C.sylvian. | SurfaceArea | 0.14 | 5.50E-02 | 0.09 |
| S.C.sylvian. | Width | 0.22 | 1.90E-03 | 0.09 |
| S.Call. | Length | 0.22 | 9.10E-05 | 0.07 |

| | | | | |
|---|---|---|---|---|
| S.Call. | MeanDepth | 0.40 | 8.70E-09 | 0.08 |
| S.Call. | SurfaceArea | 0.31 | 4.00E-07 | 0.07 |
| S.Call. | Width | 0.23 | 9.80E-05 | 0.07 |
| S.Cu. | Length | - | 4.90E-01 | 0.05 |
| S.Cu. | MeanDepth | 0.08 | 9.30E-02 | 0.06 |
| S.Cu. | SurfaceArea | 0.01 | 4.00E-01 | 0.05 |
| S.Cu. | Width | 0.19 | 4.80E-04 | 0.06 |
| S.F.inf. | Length | 0.07 | 1.40E-01 | 0.07 |
| S.F.inf. | MeanDepth | 0.22 | 2.50E-04 | 0.07 |
| S.F.inf. | SurfaceArea | 0.16 | 3.70E-03 | 0.07 |
| S.F.inf. | Width | 0.21 | 5.80E-05 | 0.06 |
| S.F.inf.ant. | Length | - | - | - |
| S.F.inf.ant. | MeanDepth | 0.10 | 5.00E-02 | 0.06 |
| S.F.inf.ant. | SurfaceArea | 0.07 | 1.20E-01 | 0.06 |
| S.F.inf.ant. | Width | 0.17 | 1.40E-03 | 0.06 |
| S.F.int. | Length | 0.06 | 1.50E-01 | 0.06 |
| S.F.int. | MeanDepth | 0.13 | 1.70E-02 | 0.07 |
| S.F.int. | SurfaceArea | 0.06 | 1.60E-01 | 0.06 |
| S.F.int. | Width | 0.22 | 1.10E-04 | 0.07 |
| S.F.inter. | Length | 0.19 | 1.10E-04 | 0.06 |
| S.F.inter. | MeanDepth | 0.10 | 1.90E-02 | 0.06 |
| S.F.inter. | SurfaceArea | 0.24 | 4.40E-06 | 0.06 |
| S.F.inter. | Width | 0.30 | 1.40E-09 | 0.06 |
| S.F.marginal. | Length | 0.12 | 2.60E-02 | 0.07 |
| S.F.marginal. | MeanDepth | 0.03 | 2.80E-01 | 0.05 |
| S.F.marginal. | SurfaceArea | 0.13 | 1.10E-02 | 0.06 |
| S.F.marginal. | Width | 0.15 | 8.90E-04 | 0.06 |
| S.F.median. | Length | 0.06 | 1.40E-01 | 0.06 |
| S.F.median. | MeanDepth | 0.22 | 2.00E-05 | 0.06 |
| S.F.median. | SurfaceArea | 0.10 | 3.50E-02 | 0.06 |
| S.F.median. | Width | 0.27 | 1.60E-06 | 0.07 |
| S.F.orbitaire. | Length | 0.04 | 2.50E-01 | 0.07 |
| S.F.orbitaire. | MeanDepth | 0.04 | 2.70E-01 | 0.06 |
| S.F.orbitaire. | SurfaceArea | 0.03 | 3.10E-01 | 0.07 |
| S.F.orbitaire. | Width | 0.11 | 4.80E-02 | 0.07 |
| S.F.polaire.tr. | Length | 0.24 | 1.20E-05 | 0.07 |
| S.F.polaire.tr. | MeanDepth | 0.17 | 8.10E-04 | 0.06 |
| S.F.polaire.tr. | SurfaceArea | 0.24 | 1.50E-05 | 0.07 |
| S.F.polaire.tr. | Width | 0.16 | 1.30E-03 | 0.06 |
| S.F.sup. | Length | 0.06 | 1.10E-01 | 0.05 |
| S.F.sup. | MeanDepth | 0.32 | 4.00E-07 | 0.07 |
| S.F.sup. | SurfaceArea | 0.10 | 3.00E-02 | 0.06 |
| S.F.sup. | Width | 0.25 | 2.00E-07 | 0.06 |
| S.GSM. | Length | - | - | - |
| S.GSM. | MeanDepth | - | - | - |
| S.GSM. | SurfaceArea | - | - | - |

| | | | | |
|---|---|---|---|---|
| S.GSM. | Width | - | - | - |
| S.Li.ant. | Length | 0.16 | 5.20E-03 | 0.07 |
| S.Li.ant. | MeanDepth | 0.09 | 6.20E-02 | 0.06 |
| S.Li.ant. | SurfaceArea | 0.11 | 2.00E-02 | 0.06 |
| S.Li.ant. | Width | 0.11 | 3.70E-02 | 0.07 |
| S.Li.post. | Length | 0.05 | 1.50E-01 | 0.05 |
| S.Li.post. | MeanDepth | 0.08 | 1.10E-01 | 0.07 |
| S.Li.post. | SurfaceArea | - | - | - |
| S.Li.post. | Width | 0.07 | 1.20E-01 | 0.06 |
| S.O.p. | Length | 0.11 | 3.40E-02 | 0.07 |
| S.O.p. | MeanDepth | 0.07 | 7.50E-02 | 0.06 |
| S.O.p. | SurfaceArea | 0.08 | 9.40E-02 | 0.06 |
| S.O.p. | Width | 0.16 | 1.40E-03 | 0.06 |
| S.O.T.lat.ant. | Length | 0.28 | 6.50E-06 | 0.08 |
| S.O.T.lat.ant. | MeanDepth | 0.37 | 1.50E-10 | 0.07 |
| S.O.T.lat.ant. | SurfaceArea | 0.40 | 9.30E-11 | 0.08 |
| S.O.T.lat.ant. | Width | 0.37 | 4.60E-09 | 0.07 |
| S.O.T.lat.int. | Length | 0.08 | 1.10E-01 | 0.07 |
| S.O.T.lat.int. | MeanDepth | 0.11 | 5.80E-02 | 0.07 |
| S.O.T.lat.int. | SurfaceArea | 0.02 | 3.90E-01 | 0.06 |
| S.O.T.lat.int. | Width | 0.01 | 4.30E-01 | 0.07 |
| S.O.T.lat.med. | Length | 0.04 | 2.00E-01 | 0.05 |
| S.O.T.lat.med. | MeanDepth | 0.10 | 3.60E-02 | 0.06 |
| S.O.T.lat.med. | SurfaceArea | 0.07 | 7.00E-02 | 0.06 |
| S.O.T.lat.med. | Width | 0.13 | 5.30E-03 | 0.06 |
| S.O.T.lat.post. | Length | 0.09 | 6.70E-02 | 0.06 |
| S.O.T.lat.post. | MeanDepth | 0.09 | 7.20E-02 | 0.06 |
| S.O.T.lat.post. | SurfaceArea | 0.12 | 1.70E-02 | 0.06 |
| S.O.T.lat.post. | Width | 0.08 | 7.80E-02 | 0.06 |
| S.Olf. | Length | 0.24 | 7.40E-06 | 0.07 |
| S.Olf. | MeanDepth | 0.22 | 7.90E-05 | 0.07 |
| S.Olf. | SurfaceArea | 0.31 | 1.00E-07 | 0.07 |
| S.Olf. | Width | 0.14 | 7.70E-03 | 0.06 |
| S.Or. | Length | 0.22 | 5.90E-05 | 0.07 |
| S.Or. | MeanDepth | 0.30 | 2.00E-07 | 0.07 |
| S.Or. | SurfaceArea | 0.27 | 4.70E-06 | 0.07 |
| S.Or. | Width | 0.27 | 8.00E-06 | 0.07 |
| S.p.C. | Length | 0.02 | 3.80E-01 | 0.06 |
| S.p.C. | MeanDepth | - | - | - |
| S.p.C. | SurfaceArea | - | - | - |
| S.p.C. | Width | 0.04 | 2.60E-01 | 0.07 |
| S.Pa.int. | Length | 0.03 | 2.50E-01 | 0.05 |
| S.Pa.int. | MeanDepth | 0.02 | 3.60E-01 | 0.05 |
| S.Pa.int. | SurfaceArea | - | - | - |
| S.Pa.int. | Width | 0.22 | 4.50E-06 | 0.06 |
| S.Pa.sup. | Length | 0.03 | 2.70E-01 | 0.05 |

| | | | | |
|---|---|---|---|---|
| S.Pa.sup. | MeanDepth | 0.10 | 4.60E-02 | 0.06 |
| S.Pa.sup. | SurfaceArea | 0.05 | 1.60E-01 | 0.06 |
| S.Pa.sup. | Width | 0.09 | 5.20E-02 | 0.06 |
| S.Pa.t. | Length | 0.06 | 1.90E-01 | 0.08 |
| S.Pa.t. | MeanDepth | 0.10 | 7.80E-02 | 0.07 |
| S.Pa.t. | SurfaceArea | 0.02 | 4.00E-01 | 0.06 |
| S.Pa.t. | Width | 0.21 | 3.20E-03 | 0.08 |
| S.Pe.C.inf. | Length | 0.07 | 1.30E-01 | 0.06 |
| S.Pe.C.inf. | MeanDepth | 0.27 | 1.30E-05 | 0.07 |
| S.Pe.C.inf. | SurfaceArea | 0.10 | 4.30E-02 | 0.07 |
| S.Pe.C.inf. | Width | 0.19 | 1.40E-03 | 0.07 |
| S.Pe.C.inter. | Length | - | - | - |
| S.Pe.C.inter. | MeanDepth | 0.13 | 4.60E-03 | 0.06 |
| S.Pe.C.inter. | SurfaceArea | 0.11 | 1.50E-02 | 0.06 |
| S.Pe.C.inter. | Width | 0.23 | 1.00E-06 | 0.06 |
| S.Pe.C.marginal. | Length | - | - | - |
| S.Pe.C.marginal. | MeanDepth | - | - | - |
| S.Pe.C.marginal. | SurfaceArea | - | - | - |
| S.Pe.C.marginal. | Width | 0.17 | 5.20E-04 | 0.06 |
| S.Pe.C.median. | Length | 0.04 | 2.50E-01 | 0.06 |
| S.Pe.C.median. | MeanDepth | 0.06 | 1.10E-01 | 0.05 |
| S.Pe.C.median. | SurfaceArea | 0.06 | 1.20E-01 | 0.06 |
| S.Pe.C.median. | Width | 0.15 | 4.00E-03 | 0.06 |
| S.Pe.C.sup. | Length | - | - | - |
| S.Pe.C.sup. | MeanDepth | 0.12 | 1.70E-02 | 0.06 |
| S.Pe.C.sup. | SurfaceArea | - | - | - |
| S.Pe.C.sup. | Width | 0.24 | 5.70E-06 | 0.06 |
| S.Po.C.sup. | Length | 0.18 | 5.70E-04 | 0.07 |
| S.Po.C.sup. | MeanDepth | 0.14 | 5.20E-03 | 0.06 |
| S.Po.C.sup. | SurfaceArea | 0.20 | 2.10E-04 | 0.07 |
| S.Po.C.sup. | Width | 0.19 | 9.40E-04 | 0.07 |
| S.R.inf. | Length | 0.20 | 3.10E-04 | 0.07 |
| S.R.inf. | MeanDepth | 0.11 | 2.40E-02 | 0.06 |
| S.R.inf. | SurfaceArea | 0.19 | 3.30E-04 | 0.07 |
| S.R.inf. | Width | 0.02 | 3.20E-01 | 0.06 |
| S.Rh. | Length | 0.19 | 9.10E-04 | 0.07 |
| S.Rh. | MeanDepth | 0.16 | 7.20E-03 | 0.07 |
| S.Rh. | SurfaceArea | 0.17 | 3.50E-03 | 0.07 |
| S.Rh. | Width | 0.04 | 2.80E-01 | 0.07 |
| S.s.P. | Length | 0.19 | 1.30E-03 | 0.07 |
| S.s.P. | MeanDepth | 0.14 | 6.30E-03 | 0.06 |
| S.s.P. | SurfaceArea | 0.20 | 6.10E-04 | 0.07 |
| S.s.P. | Width | 0.24 | 8.30E-06 | 0.07 |
| S.T.i.ant. | Length | 0.15 | 8.20E-03 | 0.07 |
| S.T.i.ant. | MeanDepth | 0.19 | 6.50E-04 | 0.07 |
| S.T.i.ant. | SurfaceArea | 0.24 | 5.10E-05 | 0.07 |

| | | | | |
|---|---|---|---|---|
| S.T.i.ant. | Width | 0.23 | 1.30E-04 | 0.07 |
| S.T.i.post. | Length | - | - | - |
| S.T.i.post. | MeanDepth | 0.09 | 5.20E-02 | 0.06 |
| S.T.i.post. | SurfaceArea | 0.04 | 2.60E-01 | 0.06 |
| S.T.i.post. | Width | 0.29 | 1.00E-07 | 0.06 |
| S.T.pol. | Length | 0.19 | 2.60E-04 | 0.06 |
| S.T.pol. | MeanDepth | 0.16 | 1.40E-03 | 0.06 |
| S.T.pol. | SurfaceArea | 0.23 | 1.20E-05 | 0.06 |
| S.T.pol. | Width | 0.18 | 4.30E-05 | 0.06 |
| S.T.s. | Length | 0.17 | 8.30E-04 | 0.06 |
| S.T.s. | MeanDepth | 0.36 | 8.30E-10 | 0.07 |
| S.T.s. | SurfaceArea | 0.19 | 4.40E-04 | 0.07 |
| S.T.s. | Width | 0.22 | 4.00E-05 | 0.06 |
| S.T.s.ter.asc.ant. | Length | - | - | - |
| S.T.s.ter.asc.ant. | MeanDepth | - | - | - |
| S.T.s.ter.asc.ant. | SurfaceArea | - | - | - |
| S.T.s.ter.asc.ant. | Width | 0.03 | 2.60E-01 | 0.05 |
| S.T.s.ter.asc.post. | Length | 0.04 | 1.70E-01 | 0.05 |
| S.T.s.ter.asc.post. | MeanDepth | 0.06 | 1.40E-01 | 0.06 |
| S.T.s.ter.asc.post. | SurfaceArea | 0.09 | 4.40E-02 | 0.06 |
| S.T.s.ter.asc.post. | Width | 0.14 | 2.00E-03 | 0.06 |
| Globals | Length | 0.12 | 2.90E-02 | 0.07 |
| Globals | MeanDepth | 0.10 | 3.00E-02 | 0.06 |
| Globals | SurfaceArea | 0.28 | 9.00E-07 | 0.07 |
| Globals | Width | 0.36 | 3.70E-12 | 0.06 |

**Table S18:** heritability eatimates for bilaterally average sulcal descriptors (yellow: bonferroni corrected for 61*4 comparisons) in GOBS.

| Sulci | Descriptor | h2 | pval | SE |
|---|---|---|---|---|
| F.C.L.a._left | Length | 0.06 | 1.61E-01 | 0.06 |
| F.C.L.a._left | MeanDepth | 0.21 | 1.29E-03 | 0.08 |
| F.C.L.a._left | Width | 0.28 | 1.01E-04 | 0.08 |
| F.C.L.a._left | SurfaceArea | 0.26 | 5.03E-05 | 0.08 |
| F.C.L.p._left | Length | 0.14 | 1.34E-02 | 0.07 |
| F.C.L.p._left | MeanDepth | 0.06 | 1.79E-01 | 0.07 |
| F.C.L.p._left | Width | 0.38 | 1.27E-12 | 0.06 |
| F.C.L.p._left | SurfaceArea | 0.39 | 3.01E-09 | 0.08 |
| F.C.L.r.ant._left | Length | 0.11 | 6.88E-02 | 0.08 |
| F.C.L.r.ant._left | MeanDepth | 0.14 | 1.78E-02 | 0.07 |
| F.C.L.r.ant._left | Width | 0.19 | 8.83E-03 | 0.09 |
| F.C.L.r.ant._left | SurfaceArea | 0.25 | 4.68E-04 | 0.09 |
| F.C.L.r.asc._left | Length | 0.18 | 1.44E-03 | 0.07 |
| F.C.L.r.asc._left | MeanDepth | 0.22 | 6.27E-05 | 0.07 |
| F.C.L.r.asc._left | Width | 0.17 | 9.84E-04 | 0.06 |
| F.C.L.r.asc._left | SurfaceArea | 0.35 | 3.57E-09 | 0.07 |
| F.C.L.r.diag._left | Length | - | - | - |
| F.C.L.r.diag._left | MeanDepth | 0.42 | 7.97E-02 | 0.28 |
| F.C.L.r.diag._left | Width | 0.26 | 1.42E-01 | 0.24 |
| F.C.L.r.diag._left | SurfaceArea | 0.01 | 4.77E-01 | 0.24 |
| F.C.L.r.retroC.tr._left | Length | 0.10 | 6.91E-02 | 0.07 |
| F.C.L.r.retroC.tr._left | MeanDepth | 0.19 | 5.78E-04 | 0.07 |
| F.C.L.r.retroC.tr._left | Width | 0.10 | 4.57E-02 | 0.07 |
| F.C.L.r.retroC.tr._left | SurfaceArea | 0.13 | 1.36E-02 | 0.06 |
| F.C.L.r.sc.ant._left | Length | - | - | - |
| F.C.L.r.sc.ant._left | MeanDepth | - | - | - |
| F.C.L.r.sc.ant._left | Width | - | - | - |
| F.C.L.r.sc.ant._left | SurfaceArea | - | - | - |
| F.C.L.r.sc.post._left | Length | - | - | - |
| F.C.L.r.sc.post._left | MeanDepth | 0.05 | 2.93E-01 | 0.09 |
| F.C.L.r.sc.post._left | Width | 0.17 | 2.93E-02 | 0.10 |
| F.C.L.r.sc.post._left | SurfaceArea | 0.02 | 3.99E-01 | 0.09 |
| F.C.M.ant._left | Length | 0.12 | 1.06E-02 | 0.06 |
| F.C.M.ant._left | MeanDepth | 0.29 | 1.20E-06 | 0.07 |
| F.C.M.ant._left | Width | 0.32 | 1.00E-07 | 0.07 |
| F.C.M.ant._left | SurfaceArea | 0.24 | 9.70E-06 | 0.07 |
| F.C.M.post._left | Length | 0.00 | 4.97E-01 | 0.06 |
| F.C.M.post._left | MeanDepth | 0.30 | 2.00E-07 | 0.07 |
| F.C.M.post._left | Width | 0.25 | 1.04E-05 | 0.07 |
| F.C.M.post._left | SurfaceArea | 0.25 | 3.47E-05 | 0.07 |
| F.Cal.ant.-Sc.Cal._left | Length | - | - | - |
| F.Cal.ant.-Sc.Cal._left | MeanDepth | - | - | - |
| F.Cal.ant.-Sc.Cal._left | Width | - | - | - |

| | | | | |
|---|---|---|---|---|
| F.Cal.ant.-Sc.Cal._left | SurfaceArea | - | - | - |
| F.Coll._left | Length | 0.03 | 2.71E-01 | 0.05 |
| F.Coll._left | MeanDepth | 0.55 | 5.48E-20 | 0.07 |
| F.Coll._left | Width | 0.34 | 5.04E-09 | 0.07 |
| F.Coll._left | SurfaceArea | 0.29 | 3.00E-07 | 0.07 |
| F.I.P._left | Length | 0.30 | 7.00E-07 | 0.07 |
| F.I.P._left | MeanDepth | 0.23 | 6.74E-05 | 0.07 |
| F.I.P._left | Width | 0.37 | 2.23E-10 | 0.07 |
| F.I.P._left | SurfaceArea | 0.27 | 2.80E-06 | 0.07 |
| F.I.P.Po.C.inf._left | Length | 0.12 | 2.47E-02 | 0.07 |
| F.I.P.Po.C.inf._left | MeanDepth | 0.19 | 1.19E-03 | 0.07 |
| F.I.P.Po.C.inf._left | Width | 0.25 | 1.85E-05 | 0.07 |
| F.I.P.Po.C.inf._left | SurfaceArea | 0.16 | 3.57E-03 | 0.07 |
| F.I.P.r.int.1_left | Length | - | - | - |
| F.I.P.r.int.1_left | MeanDepth | - | - | - |
| F.I.P.r.int.1_left | Width | - | - | - |
| F.I.P.r.int.1_left | SurfaceArea | - | - | - |
| F.I.P.r.int.2_left | Length | - | - | - |
| F.I.P.r.int.2_left | MeanDepth | - | - | - |
| F.I.P.r.int.2_left | Width | - | - | - |
| F.I.P.r.int.2_left | SurfaceArea | - | - | - |
| F.P.O._left | Length | 0.12 | 2.91E-02 | 0.07 |
| F.P.O._left | MeanDepth | 0.50 | 1.75E-12 | 0.08 |
| F.P.O._left | Width | 0.36 | 6.55E-11 | 0.06 |
| F.P.O._left | SurfaceArea | 0.37 | 3.00E-08 | 0.08 |
| INSULA_left | Length | - | - | - |
| INSULA_left | MeanDepth | 0.18 | 1.41E-03 | 0.07 |
| INSULA_left | Width | 0.25 | 4.55E-05 | 0.07 |
| INSULA_left | SurfaceArea | 0.43 | 7.20E-11 | 0.08 |
| OCCIPITAL_left | Length | 0.29 | 1.00E-06 | 0.07 |
| OCCIPITAL_left | MeanDepth | 0.32 | 1.13E-08 | 0.07 |
| OCCIPITAL_left | Width | 0.46 | 2.12E-13 | 0.07 |
| OCCIPITAL_left | SurfaceArea | 0.34 | 3.11E-08 | 0.07 |
| S.C._left | Length | 0.22 | 1.21E-04 | 0.07 |
| S.C._left | MeanDepth | 0.39 | 1.28E-10 | 0.07 |
| S.C._left | Width | 0.38 | 4.97E-11 | 0.07 |
| S.C._left | SurfaceArea | 0.39 | 5.85E-10 | 0.07 |
| S.C.LPC._left | Length | - | - | - |
| S.C.LPC._left | MeanDepth | 0.23 | 4.66E-01 | 2.57 |
| S.C.LPC._left | Width | - | - | - |
| S.C.LPC._left | SurfaceArea | - | - | - |
| S.C.sylvian._left | Length | 0.02 | 4.47E-01 | 0.12 |
| S.C.sylvian._left | MeanDepth | 0.33 | 2.48E-03 | 0.13 |
| S.C.sylvian._left | Width | 0.27 | 1.16E-02 | 0.13 |
| S.C.sylvian._left | SurfaceArea | 0.22 | 1.76E-02 | 0.12 |
| S.Call._left | Length | 0.47 | 2.16E-12 | 0.08 |

| | | | | |
|---|---|---|---|---|
| S.Call._left | MeanDepth | 0.51 | 4.54E-13 | 0.08 |
| S.Call._left | Width | 0.38 | 7.74E-10 | 0.07 |
| S.Call._left | SurfaceArea | 0.53 | 9.86E-15 | 0.07 |
| S.Cu._left | Length | - | - | - |
| S.Cu._left | MeanDepth | 0.26 | 1.34E-05 | 0.07 |
| S.Cu._left | Width | 0.28 | 4.20E-06 | 0.07 |
| S.Cu._left | SurfaceArea | 0.03 | 2.75E-01 | 0.06 |
| S.F.inf._left | Length | 0.17 | 6.76E-03 | 0.08 |
| S.F.inf._left | MeanDepth | 0.41 | 5.60E-09 | 0.09 |
| S.F.inf._left | Width | 0.27 | 5.20E-06 | 0.07 |
| S.F.inf._left | SurfaceArea | 0.25 | 2.81E-05 | 0.07 |
| S.F.inf.ant._left | Length | 0.10 | 5.05E-02 | 0.06 |
| S.F.inf.ant._left | MeanDepth | 0.08 | 9.68E-02 | 0.07 |
| S.F.inf.ant._left | Width | 0.16 | 8.63E-03 | 0.07 |
| S.F.inf.ant._left | SurfaceArea | 0.13 | 1.09E-02 | 0.07 |
| S.F.int._left | Length | 0.18 | 2.33E-03 | 0.07 |
| S.F.int._left | MeanDepth | 0.29 | 2.30E-06 | 0.07 |
| S.F.int._left | Width | 0.31 | 1.00E-07 | 0.07 |
| S.F.int._left | SurfaceArea | 0.24 | 7.96E-05 | 0.07 |
| S.F.inter._left | Length | 0.18 | 3.16E-04 | 0.06 |
| S.F.inter._left | MeanDepth | 0.13 | 6.74E-03 | 0.06 |
| S.F.inter._left | Width | 0.32 | 2.00E-09 | 0.07 |
| S.F.inter._left | SurfaceArea | 0.23 | 9.70E-06 | 0.06 |
| S.F.marginal._left | Length | 0.12 | 1.77E-02 | 0.07 |
| S.F.marginal._left | MeanDepth | 0.05 | 1.70E-01 | 0.06 |
| S.F.marginal._left | Width | 0.20 | 1.17E-04 | 0.06 |
| S.F.marginal._left | SurfaceArea | 0.18 | 1.77E-03 | 0.07 |
| S.F.median._left | Length | 0.19 | 7.25E-04 | 0.07 |
| S.F.median._left | MeanDepth | 0.18 | 1.08E-03 | 0.07 |
| S.F.median._left | Width | 0.25 | 6.10E-06 | 0.07 |
| S.F.median._left | SurfaceArea | 0.20 | 2.37E-04 | 0.07 |
| S.F.orbitaire._left | Length | 0.13 | 2.91E-02 | 0.08 |
| S.F.orbitaire._left | MeanDepth | 0.08 | 9.93E-02 | 0.07 |
| S.F.orbitaire._left | Width | 0.28 | 8.18E-05 | 0.08 |
| S.F.orbitaire._left | SurfaceArea | 0.15 | 1.44E-02 | 0.08 |
| S.F.polaire.tr._left | Length | 0.24 | 1.60E-05 | 0.07 |
| S.F.polaire.tr._left | MeanDepth | 0.11 | 2.77E-02 | 0.06 |
| S.F.polaire.tr._left | Width | 0.19 | 1.95E-03 | 0.07 |
| S.F.polaire.tr._left | SurfaceArea | 0.23 | 1.31E-04 | 0.07 |
| S.F.sup._left | Length | 0.15 | 2.15E-03 | 0.06 |
| S.F.sup._left | MeanDepth | 0.40 | 1.93E-10 | 0.07 |
| S.F.sup._left | Width | 0.27 | 1.00E-07 | 0.06 |
| S.F.sup._left | SurfaceArea | 0.20 | 3.14E-04 | 0.07 |
| S.Li.ant._left | Length | 0.24 | 4.32E-04 | 0.08 |
| S.Li.ant._left | MeanDepth | 0.18 | 8.11E-03 | 0.09 |
| S.Li.ant._left | Width | 0.03 | 3.32E-01 | 0.07 |

| | | | | |
|---|---|---|---|---|
| S.Li.ant._left | SurfaceArea | 0.28 | 1.47E-04 | 0.09 |
| S.Li.post._left | Length | 0.06 | 1.72E-01 | 0.07 |
| S.Li.post._left | MeanDepth | 0.17 | 6.13E-03 | 0.08 |
| S.Li.post._left | Width | 0.13 | 3.87E-02 | 0.08 |
| S.Li.post._left | SurfaceArea | 0.03 | 3.26E-01 | 0.07 |
| S.O.p._left | Length | 0.06 | 1.98E-01 | 0.07 |
| S.O.p._left | MeanDepth | 0.13 | 2.78E-02 | 0.08 |
| S.O.p._left | Width | 0.26 | 1.11E-04 | 0.08 |
| S.O.p._left | SurfaceArea | 0.11 | 5.32E-02 | 0.08 |
| S.O.T.lat.ant._left | Length | 0.39 | 4.05E-09 | 0.08 |
| S.O.T.lat.ant._left | MeanDepth | 0.38 | 8.30E-10 | 0.07 |
| S.O.T.lat.ant._left | Width | 0.30 | 5.00E-07 | 0.07 |
| S.O.T.lat.ant._left | SurfaceArea | 0.43 | 2.60E-11 | 0.08 |
| S.O.T.lat.int._left | Length | 0.11 | 5.67E-02 | 0.08 |
| S.O.T.lat.int._left | MeanDepth | 0.16 | 2.60E-02 | 0.09 |
| S.O.T.lat.int._left | Width | 0.13 | 6.09E-02 | 0.09 |
| S.O.T.lat.int._left | SurfaceArea | 0.07 | 1.99E-01 | 0.09 |
| S.O.T.lat.med._left | Length | 0.05 | 1.89E-01 | 0.06 |
| S.O.T.lat.med._left | MeanDepth | 0.09 | 4.42E-02 | 0.06 |
| S.O.T.lat.med._left | Width | 0.18 | 1.18E-03 | 0.07 |
| S.O.T.lat.med._left | SurfaceArea | 0.09 | 5.16E-02 | 0.06 |
| S.O.T.lat.post._left | Length | 0.04 | 2.52E-01 | 0.06 |
| S.O.T.lat.post._left | MeanDepth | 0.10 | 6.02E-02 | 0.07 |
| S.O.T.lat.post._left | Width | 0.16 | 3.20E-03 | 0.07 |
| S.O.T.lat.post._left | SurfaceArea | 0.15 | 6.91E-03 | 0.07 |
| S.Olf._left | Length | 0.46 | 2.53E-13 | 0.07 |
| S.Olf._left | MeanDepth | 0.39 | 9.39E-12 | 0.07 |
| S.Olf._left | Width | 0.24 | 7.77E-05 | 0.07 |
| S.Olf._left | SurfaceArea | 0.51 | 1.65E-15 | 0.07 |
| S.Or._left | Length | 0.20 | 4.78E-04 | 0.07 |
| S.Or._left | MeanDepth | 0.32 | 2.00E-07 | 0.08 |
| S.Or._left | Width | 0.32 | 9.00E-07 | 0.08 |
| S.Or._left | SurfaceArea | 0.23 | 1.68E-04 | 0.08 |
| S.p.C._left | Length | 0.13 | 4.19E-02 | 0.08 |
| S.p.C._left | MeanDepth | 0.02 | 3.89E-01 | 0.08 |
| S.p.C._left | Width | 0.01 | 4.60E-01 | 0.09 |
| S.p.C._left | SurfaceArea | 0.08 | 1.48E-01 | 0.08 |
| S.Pa.int._left | Length | 0.20 | 8.63E-04 | 0.07 |
| S.Pa.int._left | MeanDepth | 0.16 | 3.49E-03 | 0.07 |
| S.Pa.int._left | Width | 0.31 | 2.59E-08 | 0.07 |
| S.Pa.int._left | SurfaceArea | 0.14 | 1.27E-02 | 0.07 |
| S.Pa.sup._left | Length | 0.16 | 1.42E-02 | 0.08 |
| S.Pa.sup._left | MeanDepth | 0.12 | 4.77E-02 | 0.08 |
| S.Pa.sup._left | Width | 0.28 | 8.50E-05 | 0.08 |
| S.Pa.sup._left | SurfaceArea | 0.22 | 1.26E-03 | 0.08 |
| S.Pa.t._left | Length | 0.07 | 3.13E-01 | 0.16 |

| | | | | |
|---|---|---|---|---|
| S.Pa.t._left | MeanDepth | 0.04 | 3.41E-01 | 0.09 |
| S.Pa.t._left | Width | 0.11 | 1.15E-01 | 0.10 |
| S.Pa.t._left | SurfaceArea | 0.07 | 2.10E-01 | 0.10 |
| S.Pe.C.inf._left | Length | 0.16 | 4.24E-03 | 0.07 |
| S.Pe.C.inf._left | MeanDepth | 0.23 | 2.84E-04 | 0.08 |
| S.Pe.C.inf._left | Width | 0.34 | 1.50E-06 | 0.08 |
| S.Pe.C.inf._left | SurfaceArea | 0.13 | 1.45E-02 | 0.07 |
| S.Pe.C.inter._left | Length | - | - | - |
| S.Pe.C.inter._left | MeanDepth | 0.24 | 2.47E-05 | 0.07 |
| S.Pe.C.inter._left | Width | 0.24 | 3.60E-06 | 0.06 |
| S.Pe.C.inter._left | SurfaceArea | 0.13 | 8.36E-03 | 0.06 |
| S.Pe.C.marginal._left | Length | - | - | - |
| S.Pe.C.marginal._left | MeanDepth | 0.06 | 2.03E-01 | 0.07 |
| S.Pe.C.marginal._left | Width | 0.29 | 7.21E-05 | 0.09 |
| S.Pe.C.marginal._left | SurfaceArea | - | - | - |
| S.Pe.C.median._left | Length | 0.11 | 6.37E-02 | 0.08 |
| S.Pe.C.median._left | MeanDepth | 0.08 | 1.39E-01 | 0.08 |
| S.Pe.C.median._left | Width | 0.21 | 5.62E-04 | 0.07 |
| S.Pe.C.median._left | SurfaceArea | 0.10 | 6.29E-02 | 0.07 |
| S.Pe.C.sup._left | Length | 0.15 | 9.73E-03 | 0.07 |
| S.Pe.C.sup._left | MeanDepth | 0.17 | 3.14E-03 | 0.07 |
| S.Pe.C.sup._left | Width | 0.20 | 2.76E-04 | 0.07 |
| S.Pe.C.sup._left | SurfaceArea | 0.15 | 9.59E-03 | 0.07 |
| S.Po.C.sup._left | Length | 0.12 | 2.65E-02 | 0.07 |
| S.Po.C.sup._left | MeanDepth | 0.22 | 3.01E-04 | 0.07 |
| S.Po.C.sup._left | Width | 0.32 | 3.00E-07 | 0.08 |
| S.Po.C.sup._left | SurfaceArea | 0.19 | 8.50E-04 | 0.07 |
| S.R.inf._left | Length | 0.18 | 1.79E-02 | 0.09 |
| S.R.inf._left | MeanDepth | 0.16 | 2.73E-02 | 0.09 |
| S.R.inf._left | Width | 0.13 | 3.31E-02 | 0.08 |
| S.R.inf._left | SurfaceArea | 0.18 | 1.86E-02 | 0.10 |
| S.Rh._left | Length | 0.31 | 1.58E-05 | 0.08 |
| S.Rh._left | MeanDepth | 0.20 | 1.81E-03 | 0.08 |
| S.Rh._left | Width | 0.15 | 1.51E-02 | 0.08 |
| S.Rh._left | SurfaceArea | 0.21 | 1.96E-03 | 0.08 |
| S.s.P._left | Length | 0.19 | 1.85E-03 | 0.08 |
| S.s.P._left | MeanDepth | 0.29 | 1.00E-06 | 0.07 |
| S.s.P._left | Width | 0.29 | 4.00E-07 | 0.07 |
| S.s.P._left | SurfaceArea | 0.24 | 1.19E-04 | 0.07 |
| S.T.i.ant._left | Length | 0.22 | 3.71E-04 | 0.07 |
| S.T.i.ant._left | MeanDepth | 0.33 | 2.81E-08 | 0.07 |
| S.T.i.ant._left | Width | 0.41 | 3.94E-10 | 0.08 |
| S.T.i.ant._left | SurfaceArea | 0.28 | 1.40E-06 | 0.07 |
| S.T.i.post._left | Length | 0.02 | 3.38E-01 | 0.06 |
| S.T.i.post._left | MeanDepth | 0.21 | 1.00E-04 | 0.06 |
| S.T.i.post._left | Width | 0.34 | 5.53E-10 | 0.07 |

| | | | | |
|---|---|---|---|---|
| S.T.i.post._left | SurfaceArea | 0.18 | 3.16E-03 | 0.07 |
| S.T.pol._left | Length | 0.29 | 4.00E-07 | 0.07 |
| S.T.pol._left | MeanDepth | 0.24 | 5.74E-05 | 0.07 |
| S.T.pol._left | Width | 0.27 | 2.00E-07 | 0.06 |
| S.T.pol._left | SurfaceArea | 0.34 | 3.43E-09 | 0.07 |
| S.T.s._left | Length | 0.18 | 1.02E-03 | 0.07 |
| S.T.s._left | MeanDepth | 0.43 | 1.69E-13 | 0.07 |
| S.T.s._left | Width | 0.27 | 1.80E-06 | 0.07 |
| S.T.s._left | SurfaceArea | 0.15 | 6.19E-03 | 0.07 |
| S.T.s.ter.asc.ant._left | Length | - | - | - |
| S.T.s.ter.asc.ant._left | MeanDepth | - | - | - |
| S.T.s.ter.asc.ant._left | Width | 0.08 | 8.62E-02 | 0.06 |
| S.T.s.ter.asc.ant._left | SurfaceArea | - | - | - |
| S.T.s.ter.asc.post._left | Length | - | - | - |
| S.T.s.ter.asc.post._left | MeanDepth | 0.15 | 9.03E-03 | 0.07 |
| S.T.s.ter.asc.post._left | Width | 0.19 | 2.74E-04 | 0.06 |
| S.T.s.ter.asc.post._left | SurfaceArea | 0.08 | 7.83E-02 | 0.06 |
| Globals | Length | 0.16 | 3.49E-03 | 0.07 |
| Globals | MeanDepth | 0.22 | 3.18E-04 | 0.07 |
| Globals | Width | 0.39 | 1.19E-12 | 0.07 |
| Globals | SurfaceArea | 0.37 | 5.37E-10 | 0.07 |

**Table S19:** heritability eatimates for bilaterally average sulcal descriptors (yellow: bonferroni corrected for 61*4 comparisons) in UK Biobank.

| Sulci | Descriptor | h2 | pval | SE |
|---|---|---|---|---|
| F.C.L.a | Length | 0.03 | 2.08E-01 | 0.04 |
| F.C.L.a | MeanDepth | 0.04 | 1.18E-01 | 0.04 |
| F.C.L.a | Width | 0.18 | 7.73E-07 | 0.04 |
| F.C.L.a | SurfaceArea | 0.09 | 1.01E-02 | 0.04 |
| F.C.L.p | Length | 0.00 | 4.96E-01 | 0.04 |
| F.C.L.p | MeanDepth | 0.08 | 1.52E-02 | 0.04 |
| F.C.L.p | Width | 0.20 | 5.21E-09 | 0.04 |
| F.C.L.p | SurfaceArea | 0.12 | 3.72E-04 | 0.04 |
| F.C.L.r.ant | Length | 0.08 | 2.17E-02 | 0.04 |
| F.C.L.r.ant | MeanDepth | 0.04 | 1.44E-01 | 0.04 |
| F.C.L.r.ant | Width | 0.05 | 1.08E-01 | 0.04 |
| F.C.L.r.ant | SurfaceArea | 0.07 | 4.09E-02 | 0.04 |
| F.C.L.r.asc | Length | 0.00 | 4.67E-01 | 0.04 |
| F.C.L.r.asc | MeanDepth | 0.10 | 6.56E-03 | 0.04 |
| F.C.L.r.asc | Width | 0.08 | 1.53E-02 | 0.04 |
| F.C.L.r.asc | SurfaceArea | 0.04 | 1.20E-01 | 0.04 |
| F.C.L.r.diag | Length | 0.00 | 5.00E-01 | 0.05 |
| F.C.L.r.diag | MeanDepth | 0.10 | 1.28E-02 | 0.05 |
| F.C.L.r.diag | Width | 0.07 | 5.90E-02 | 0.05 |
| F.C.L.r.diag | SurfaceArea | 0.01 | 4.35E-01 | 0.05 |
| F.C.L.r.retroC.tr | Length | 0.03 | 2.28E-01 | 0.04 |
| F.C.L.r.retroC.tr | MeanDepth | 0.06 | 6.84E-02 | 0.04 |
| F.C.L.r.retroC.tr | Width | 0.06 | 5.67E-02 | 0.04 |
| F.C.L.r.retroC.tr | SurfaceArea | 0.11 | 2.22E-03 | 0.04 |
| F.C.L.r.sc.ant | Length | 0.11 | 9.64E-02 | 0.08 |
| F.C.L.r.sc.ant | MeanDepth | 0.05 | 2.79E-01 | 0.08 |
| F.C.L.r.sc.ant | Width | 0.12 | 6.73E-02 | 0.08 |
| F.C.L.r.sc.ant | SurfaceArea | 0.02 | 3.99E-01 | 0.08 |
| F.C.L.r.sc.post | Length | 0.04 | 1.12E-01 | 0.04 |
| F.C.L.r.sc.post | MeanDepth | 0.01 | 3.87E-01 | 0.04 |
| F.C.L.r.sc.post | Width | 0.05 | 7.91E-02 | 0.04 |
| F.C.L.r.sc.post | SurfaceArea | 0.17 | 5.18E-06 | 0.04 |
| F.C.M.ant | Length | 0.04 | 1.32E-01 | 0.04 |
| F.C.M.ant | MeanDepth | 0.07 | 2.17E-02 | 0.04 |
| F.C.M.ant | Width | 0.06 | 7.44E-02 | 0.04 |
| F.C.M.ant | SurfaceArea | 0.06 | 4.91E-02 | 0.04 |
| F.C.M.post | Length | 0.04 | 1.62E-01 | 0.04 |
| F.C.M.post | MeanDepth | 0.11 | 1.05E-03 | 0.04 |
| F.C.M.post | Width | 0.21 | 6.05E-10 | 0.04 |
| F.C.M.post | SurfaceArea | 0.08 | 1.57E-02 | 0.04 |
| F.Cal.ant.-Sc.Cal | Length | 0.00 | 5.00E-01 | 0.04 |
| F.Cal.ant.-Sc.Cal | MeanDepth | 0.12 | 8.07E-04 | 0.04 |
| F.Cal.ant.-Sc.Cal | Width | 0.13 | 1.81E-04 | 0.04 |

| | | | | |
|---|---|---|---|---|
| F.Cal.ant.-Sc.Cal | SurfaceArea | 0.16 | 1.09E-05 | 0.04 |
| F.Coll | Length | 0.05 | 1.13E-01 | 0.04 |
| F.Coll | MeanDepth | 0.10 | 4.16E-03 | 0.04 |
| F.Coll | Width | 0.17 | 4.83E-06 | 0.04 |
| F.Coll | SurfaceArea | 0.08 | 1.88E-02 | 0.04 |
| F.I.P | Length | 0.00 | 4.47E-01 | 0.04 |
| F.I.P | MeanDepth | 0.18 | 1.35E-06 | 0.04 |
| F.I.P | Width | 0.18 | 1.78E-07 | 0.04 |
| F.I.P | SurfaceArea | 0.03 | 1.96E-01 | 0.04 |
| F.I.P.Po.C.inf | Length | 0.00 | 5.00E-01 | 0.04 |
| F.I.P.Po.C.inf | MeanDepth | 0.01 | 3.79E-01 | 0.04 |
| F.I.P.Po.C.inf | Width | 0.20 | 5.58E-09 | 0.04 |
| F.I.P.Po.C.inf | SurfaceArea | 0.00 | 4.65E-01 | 0.04 |
| F.I.P.r.int. | Length | 0.07 | 2.75E-02 | 0.04 |
| F.I.P.r.int. | MeanDepth | 0.05 | 8.86E-02 | 0.04 |
| F.I.P.r.int. | Width | 0.08 | 1.94E-02 | 0.04 |
| F.I.P.r.int. | SurfaceArea | 0.07 | 3.81E-02 | 0.04 |
| F.I.P.r.int. | Length | 0.06 | 7.39E-02 | 0.04 |
| F.I.P.r.int. | MeanDepth | 0.00 | 5.00E-01 | 0.04 |
| F.I.P.r.int. | Width | 0.10 | 4.76E-03 | 0.04 |
| F.I.P.r.int. | SurfaceArea | 0.02 | 3.20E-01 | 0.04 |
| F.P.O | Length | 0.00 | 5.00E-01 | 0.04 |
| F.P.O | MeanDepth | 0.15 | 2.23E-05 | 0.04 |
| F.P.O | Width | 0.16 | 1.13E-05 | 0.04 |
| F.P.O | SurfaceArea | 0.09 | 5.90E-03 | 0.04 |
| INSULA | Length | 0.00 | 5.00E-01 | 0.04 |
| INSULA | MeanDepth | 0.06 | 4.53E-02 | 0.04 |
| INSULA | Width | 0.14 | 2.59E-05 | 0.04 |
| INSULA | SurfaceArea | 0.16 | 3.82E-06 | 0.04 |
| OCCIPITA | Length | 0.11 | 1.93E-03 | 0.04 |
| OCCIPITA | MeanDepth | 0.12 | 7.21E-04 | 0.04 |
| OCCIPITA | Width | 0.15 | 1.57E-05 | 0.04 |
| OCCIPITA | SurfaceArea | 0.12 | 6.15E-04 | 0.04 |
| S.C | Length | 0.11 | 6.94E-03 | 0.04 |
| S.C | MeanDepth | 0.15 | 1.77E-04 | 0.04 |
| S.C | Width | 0.26 | 3.92E-10 | 0.04 |
| S.C | SurfaceArea | 0.08 | 3.33E-02 | 0.04 |
| S.C.LPC | Length | 0.00 | 5.00E-01 | 0.05 |
| S.C.LPC | MeanDepth | 0.04 | 2.17E-01 | 0.05 |
| S.C.LPC | Width | 0.17 | 7.05E-04 | 0.05 |
| S.C.LPC | SurfaceArea | 0.02 | 3.49E-01 | 0.05 |
| S.C.sylvian | Length | 0.04 | 1.64E-01 | 0.04 |
| S.C.sylvian | MeanDepth | 0.06 | 6.65E-02 | 0.04 |
| S.C.sylvian | Width | 0.05 | 1.20E-01 | 0.04 |
| S.C.sylvian | SurfaceArea | 0.01 | 3.64E-01 | 0.04 |
| S.Call | Length | 0.15 | 4.39E-05 | 0.04 |

| | | | | |
|---|---|---|---|---|
| S.Call | MeanDepth | 0.18 | 8.14E-07 | 0.04 |
| S.Call | Width | 0.09 | 7.59E-03 | 0.04 |
| S.Call | SurfaceArea | 0.18 | 2.04E-06 | 0.04 |
| S.Cu | Length | 0.02 | 3.04E-01 | 0.04 |
| S.Cu | MeanDepth | 0.06 | 5.26E-02 | 0.04 |
| S.Cu | Width | 0.08 | 1.91E-02 | 0.04 |
| S.Cu | SurfaceArea | 0.00 | 5.00E-01 | 0.04 |
| S.F.inf | Length | 0.05 | 1.12E-01 | 0.04 |
| S.F.inf | MeanDepth | 0.20 | 1.63E-08 | 0.04 |
| S.F.inf | Width | 0.17 | 1.93E-06 | 0.04 |
| S.F.inf | SurfaceArea | 0.11 | 1.74E-03 | 0.04 |
| S.F.inf.ant | Length | 0.03 | 2.04E-01 | 0.04 |
| S.F.inf.ant | MeanDepth | 0.02 | 2.62E-01 | 0.04 |
| S.F.inf.ant | Width | 0.14 | 1.29E-04 | 0.04 |
| S.F.inf.ant | SurfaceArea | 0.03 | 1.94E-01 | 0.04 |
| S.F.int | Length | 0.13 | 1.62E-04 | 0.04 |
| S.F.int | MeanDepth | 0.22 | 1.29E-10 | 0.04 |
| S.F.int | Width | 0.16 | 1.98E-06 | 0.04 |
| S.F.int | SurfaceArea | 0.17 | 7.39E-07 | 0.04 |
| S.F.inter | Length | 0.03 | 2.42E-01 | 0.04 |
| S.F.inter | MeanDepth | 0.08 | 1.04E-02 | 0.04 |
| S.F.inter | Width | 0.10 | 2.32E-03 | 0.04 |
| S.F.inter | SurfaceArea | 0.03 | 1.78E-01 | 0.04 |
| S.F.marginal | Length | 0.07 | 3.34E-02 | 0.04 |
| S.F.marginal | MeanDepth | 0.11 | 1.22E-03 | 0.04 |
| S.F.marginal | Width | 0.09 | 6.22E-03 | 0.04 |
| S.F.marginal | SurfaceArea | 0.12 | 6.16E-04 | 0.04 |
| S.F.median | Length | 0.02 | 2.58E-01 | 0.04 |
| S.F.median | MeanDepth | 0.08 | 1.94E-02 | 0.04 |
| S.F.median | Width | 0.08 | 2.17E-02 | 0.04 |
| S.F.median | SurfaceArea | 0.04 | 1.29E-01 | 0.04 |
| S.F.orbitaire | Length | 0.00 | 5.00E-01 | 0.04 |
| S.F.orbitaire | MeanDepth | 0.00 | 5.00E-01 | 0.04 |
| S.F.orbitaire | Width | 0.00 | 5.00E-01 | 0.04 |
| S.F.orbitaire | SurfaceArea | 0.00 | 5.00E-01 | 0.04 |
| S.F.polaire.tr | Length | 0.11 | 2.46E-03 | 0.04 |
| S.F.polaire.tr | MeanDepth | 0.00 | 5.00E-01 | 0.04 |
| S.F.polaire.tr | Width | 0.07 | 1.76E-02 | 0.04 |
| S.F.polaire.tr | SurfaceArea | 0.08 | 1.70E-02 | 0.04 |
| S.F.sup | Length | 0.00 | 5.00E-01 | 0.04 |
| S.F.sup | MeanDepth | 0.05 | 8.60E-02 | 0.04 |
| S.F.sup | Width | 0.14 | 1.75E-05 | 0.04 |
| S.F.sup | SurfaceArea | 0.03 | 1.68E-01 | 0.04 |
| S.Li.ant | Length | 0.07 | 2.77E-02 | 0.04 |
| S.Li.ant | MeanDepth | 0.10 | 2.99E-03 | 0.04 |
| S.Li.ant | Width | 0.09 | 8.34E-03 | 0.04 |

| | | | | |
|---|---|---|---|---|
| S.Li.ant | SurfaceArea | 0.10 | 2.94E-03 | 0.04 |
| S.Li.post | Length | 0.04 | 1.69E-01 | 0.04 |
| S.Li.post | MeanDepth | 0.04 | 1.22E-01 | 0.04 |
| S.Li.post | Width | 0.12 | 8.37E-04 | 0.04 |
| S.Li.post | SurfaceArea | 0.01 | 4.14E-01 | 0.04 |
| S.O.p | Length | 0.00 | 4.90E-01 | 0.04 |
| S.O.p | MeanDepth | 0.08 | 2.39E-02 | 0.04 |
| S.O.p | Width | 0.02 | 3.11E-01 | 0.04 |
| S.O.p | SurfaceArea | 0.01 | 3.60E-01 | 0.04 |
| S.O.T.lat.ant | Length | 0.08 | 1.51E-02 | 0.04 |
| S.O.T.lat.ant | MeanDepth | 0.17 | 2.41E-06 | 0.04 |
| S.O.T.lat.ant | Width | 0.07 | 4.05E-02 | 0.04 |
| S.O.T.lat.ant | SurfaceArea | 0.14 | 1.32E-04 | 0.04 |
| S.O.T.lat.int | Length | 0.02 | 3.26E-01 | 0.04 |
| S.O.T.lat.int | MeanDepth | 0.08 | 1.75E-02 | 0.04 |
| S.O.T.lat.int | Width | 0.07 | 3.01E-02 | 0.04 |
| S.O.T.lat.int | SurfaceArea | 0.01 | 3.68E-01 | 0.04 |
| S.O.T.lat.med | Length | 0.05 | 8.76E-02 | 0.04 |
| S.O.T.lat.med | MeanDepth | 0.10 | 3.70E-03 | 0.04 |
| S.O.T.lat.med | Width | 0.05 | 8.12E-02 | 0.04 |
| S.O.T.lat.med | SurfaceArea | 0.06 | 7.22E-02 | 0.04 |
| S.O.T.lat.post | Length | 0.00 | 5.00E-01 | 0.04 |
| S.O.T.lat.post | MeanDepth | 0.07 | 3.18E-02 | 0.04 |
| S.O.T.lat.post | Width | 0.07 | 3.00E-02 | 0.04 |
| S.O.T.lat.post | SurfaceArea | 0.03 | 2.16E-01 | 0.04 |
| S.Olf | Length | 0.18 | 3.46E-07 | 0.04 |
| S.Olf | MeanDepth | 0.19 | 2.47E-07 | 0.04 |
| S.Olf | Width | 0.09 | 9.26E-03 | 0.04 |
| S.Olf | SurfaceArea | 0.26 | 4.24E-12 | 0.04 |
| S.Or | Length | 0.10 | 3.28E-03 | 0.04 |
| S.Or | MeanDepth | 0.12 | 7.35E-04 | 0.04 |
| S.Or | Width | 0.13 | 4.67E-04 | 0.04 |
| S.Or | SurfaceArea | 0.14 | 1.77E-04 | 0.04 |
| S.p.C | Length | 0.00 | 5.00E-01 | 0.04 |
| S.p.C | MeanDepth | 0.00 | 5.00E-01 | 0.04 |
| S.p.C | Width | 0.10 | 5.28E-03 | 0.04 |
| S.p.C | SurfaceArea | 0.00 | 5.00E-01 | 0.04 |
| S.Pa.int | Length | 0.03 | 1.90E-01 | 0.04 |
| S.Pa.int | MeanDepth | 0.11 | 1.80E-03 | 0.04 |
| S.Pa.int | Width | 0.12 | 2.95E-04 | 0.04 |
| S.Pa.int | SurfaceArea | 0.02 | 3.35E-01 | 0.04 |
| S.Pa.sup | Length | 0.03 | 2.32E-01 | 0.04 |
| S.Pa.sup | MeanDepth | 0.00 | 5.00E-01 | 0.04 |
| S.Pa.sup | Width | 0.09 | 3.26E-03 | 0.04 |
| S.Pa.sup | SurfaceArea | 0.03 | 1.88E-01 | 0.04 |
| S.Pa.t | Length | 0.02 | 2.54E-01 | 0.04 |

| | | | | |
|---|---|---|---|---|
| S.Pa.t | MeanDepth | 0.06 | 6.89E-02 | 0.04 |
| S.Pa.t | Width | 0.07 | 2.78E-02 | 0.04 |
| S.Pa.t | SurfaceArea | 0.04 | 1.39E-01 | 0.04 |
| S.Pe.C.inf | Length | 0.06 | 5.94E-02 | 0.04 |
| S.Pe.C.inf | MeanDepth | 0.08 | 1.71E-02 | 0.04 |
| S.Pe.C.inf | Width | 0.03 | 1.79E-01 | 0.04 |
| S.Pe.C.inf | SurfaceArea | 0.04 | 1.29E-01 | 0.04 |
| S.Pe.C.inter | Length | 0.00 | 5.00E-01 | 0.04 |
| S.Pe.C.inter | MeanDepth | 0.06 | 4.98E-02 | 0.04 |
| S.Pe.C.inter | Width | 0.14 | 7.67E-05 | 0.04 |
| S.Pe.C.inter | SurfaceArea | 0.00 | 4.52E-01 | 0.04 |
| S.Pe.C.marginal | Length | 0.00 | 5.00E-01 | 0.04 |
| S.Pe.C.marginal | MeanDepth | 0.03 | 2.45E-01 | 0.04 |
| S.Pe.C.marginal | Width | 0.09 | 6.51E-03 | 0.04 |
| S.Pe.C.marginal | SurfaceArea | 0.00 | 5.00E-01 | 0.04 |
| S.Pe.C.median | Length | 0.00 | 4.66E-01 | 0.04 |
| S.Pe.C.median | MeanDepth | 0.06 | 7.06E-02 | 0.04 |
| S.Pe.C.median | Width | 0.11 | 2.74E-03 | 0.04 |
| S.Pe.C.median | SurfaceArea | 0.02 | 3.04E-01 | 0.04 |
| S.Pe.C.sup | Length | 0.00 | 4.79E-01 | 0.04 |
| S.Pe.C.sup | MeanDepth | 0.00 | 5.00E-01 | 0.04 |
| S.Pe.C.sup | Width | 0.05 | 7.98E-02 | 0.04 |
| S.Pe.C.sup | SurfaceArea | 0.00 | 5.00E-01 | 0.04 |
| S.Po.C.sup | Length | 0.05 | 9.88E-02 | 0.04 |
| S.Po.C.sup | MeanDepth | 0.00 | 5.00E-01 | 0.04 |
| S.Po.C.sup | Width | 0.20 | 4.14E-09 | 0.04 |
| S.Po.C.sup | SurfaceArea | 0.01 | 3.44E-01 | 0.04 |
| S.R.inf | Length | 0.07 | 2.80E-02 | 0.04 |
| S.R.inf | MeanDepth | 0.07 | 3.55E-02 | 0.04 |
| S.R.inf | Width | 0.06 | 7.06E-02 | 0.04 |
| S.R.inf | SurfaceArea | 0.06 | 4.75E-02 | 0.04 |
| S.Rh | Length | 0.09 | 1.08E-02 | 0.04 |
| S.Rh | MeanDepth | 0.10 | 4.10E-03 | 0.04 |
| S.Rh | Width | 0.13 | 2.19E-04 | 0.04 |
| S.Rh | SurfaceArea | 0.11 | 2.69E-03 | 0.04 |
| S.s.P | Length | 0.05 | 1.06E-01 | 0.04 |
| S.s.P | MeanDepth | 0.07 | 3.02E-02 | 0.04 |
| S.s.P | Width | 0.12 | 3.40E-04 | 0.04 |
| S.s.P | SurfaceArea | 0.06 | 5.30E-02 | 0.04 |
| S.T.i.ant | Length | 0.12 | 9.69E-04 | 0.04 |
| S.T.i.ant | MeanDepth | 0.12 | 9.45E-04 | 0.04 |
| S.T.i.ant | Width | 0.15 | 5.50E-05 | 0.04 |
| S.T.i.ant | SurfaceArea | 0.13 | 4.60E-04 | 0.04 |
| S.T.i.post | Length | 0.05 | 7.46E-02 | 0.04 |
| S.T.i.post | MeanDepth | 0.02 | 2.65E-01 | 0.04 |
| S.T.i.post | Width | 0.10 | 5.19E-03 | 0.04 |

| | | | | |
|---|---|---|---|---|
| S.T.i.post | SurfaceArea | 0.06 | 6.32E-02 | 0.04 |
| S.T.pol | Length | 0.14 | 5.91E-05 | 0.04 |
| S.T.pol | MeanDepth | 0.13 | 4.19E-04 | 0.04 |
| S.T.pol | Width | 0.15 | 3.29E-05 | 0.04 |
| S.T.pol | SurfaceArea | 0.14 | 1.11E-04 | 0.04 |
| S.T.s | Length | 0.00 | 5.00E-01 | 0.04 |
| S.T.s | MeanDepth | 0.18 | 7.22E-07 | 0.04 |
| S.T.s | Width | 0.14 | 6.72E-05 | 0.04 |
| S.T.s | SurfaceArea | 0.03 | 2.21E-01 | 0.04 |
| S.T.s.ter.asc.ant | Length | 0.00 | 5.00E-01 | 0.04 |
| S.T.s.ter.asc.ant | MeanDepth | 0.08 | 1.94E-02 | 0.04 |
| S.T.s.ter.asc.ant | Width | 0.09 | 8.26E-03 | 0.04 |
| S.T.s.ter.asc.ant | SurfaceArea | 0.00 | 5.00E-01 | 0.04 |
| S.T.s.ter.asc.post | Length | 0.06 | 5.02E-02 | 0.04 |
| S.T.s.ter.asc.post | MeanDepth | 0.08 | 1.57E-02 | 0.04 |
| S.T.s.ter.asc.post | Width | 0.06 | 6.30E-02 | 0.04 |
| S.T.s.ter.asc.post | SurfaceArea | 0.06 | 4.26E-02 | 0.04 |

**Table S20:** Meta-analysis of heritability eatimates for bilaterally average sulcal descriptors (yellow: bonferroni corrected for 61*4 comparisons).

| Sulci | Descriptor | h2 | pval | SE |
|---|---|---|---|---|
| F.C.L.a. | Length | 0.11 | 3.40E-03 | 0.01 |
| F.C.L.a. | MeanDepth | 0.21 | 3.60E-06 | 0.01 |
| F.C.L.a. | Width | 0.30 | 3.50E-11 | 0.01 |
| F.C.L.a. | SurfaceArea | 0.26 | 2.60E-08 | 0.01 |
| F.C.L.p. | Length | 0.15 | 1.40E-04 | 0.01 |
| F.C.L.p. | MeanDepth | 0.19 | 1.00E-03 | 0.01 |
| F.C.L.p. | Width | 0.54 | 2.20E-35 | 0.00 |
| F.C.L.p. | SurfaceArea | 0.44 | 1.10E-18 | 0.01 |
| F.C.L.r.ant. | Length | 0.16 | 1.10E-03 | 0.01 |
| F.C.L.r.ant. | MeanDepth | 0.14 | 9.50E-04 | 0.01 |
| F.C.L.r.ant. | Width | 0.17 | 1.50E-02 | 0.01 |
| F.C.L.r.ant. | SurfaceArea | 0.28 | 3.30E-09 | 0.01 |
| F.C.L.r.asc. | Length | 0.11 | 1.20E-02 | 0.01 |
| F.C.L.r.asc. | MeanDepth | 0.20 | 2.20E-06 | 0.01 |
| F.C.L.r.asc. | Width | 0.21 | 1.80E-08 | 0.01 |
| F.C.L.r.asc. | SurfaceArea | 0.30 | 1.70E-10 | 0.01 |
| F.C.L.r.diag. | Length | 0.16 | 6.30E-02 | 0.27 |
| F.C.L.r.diag. | MeanDepth | 0.23 | 2.00E-01 | 0.26 |
| F.C.L.r.diag. | Width | 0.25 | 6.40E-02 | 0.07 |
| F.C.L.r.diag. | SurfaceArea | - | - | - |
| F.C.L.r.retroC.tr. | Length | 0.11 | 2.50E-03 | 0.01 |
| F.C.L.r.retroC.tr. | MeanDepth | 0.28 | 3.40E-09 | 0.01 |
| F.C.L.r.retroC.tr. | Width | 0.20 | 1.00E-04 | 0.01 |
| F.C.L.r.retroC.tr. | SurfaceArea | 0.19 | 2.70E-06 | 0.01 |
| F.C.L.r.sc.ant. | Length | - | - | - |
| F.C.L.r.sc.ant. | MeanDepth | - | - | - |
| F.C.L.r.sc.ant. | Width | - | - | - |
| F.C.L.r.sc.ant. | SurfaceArea | - | - | - |
| F.C.L.r.sc.post. | Length | - | - | - |
| F.C.L.r.sc.post. | MeanDepth | 0.02 | 9.40E-01 | 0.01 |
| F.C.L.r.sc.post. | Width | 0.17 | 8.50E-01 | 0.54 |
| F.C.L.r.sc.post. | SurfaceArea | - | - | - |
| F.C.M.ant. | Length | 0.09 | 5.20E-03 | 0.00 |
| F.C.M.ant. | MeanDepth | 0.35 | 8.40E-13 | 0.01 |
| F.C.M.ant. | Width | 0.34 | 1.00E-16 | 0.01 |
| F.C.M.ant. | SurfaceArea | 0.24 | 6.00E-08 | 0.01 |
| F.C.M.post. | Length | 0.05 | 9.40E-02 | 0.01 |
| F.C.M.post. | MeanDepth | 0.39 | 2.10E-21 | 0.01 |
| F.C.M.post. | Width | 0.49 | 1.90E-16 | 0.00 |
| F.C.M.post. | SurfaceArea | 0.29 | 2.40E-14 | 0.01 |
| F.Cal.ant.-Sc.Cal. | Length | - | - | - |
| F.Cal.ant.-Sc.Cal. | MeanDepth | - | - | - |
| F.Cal.ant.-Sc.Cal. | Width | - | - | - |
| F.Cal.ant.-Sc.Cal. | SurfaceArea | - | - | - |
| F.Coll. | Length | 0.14 | 7.70E-03 | 0.00 |
| F.Coll. | MeanDepth | 0.51 | 1.30E-25 | 0.01 |
| F.Coll. | Width | 0.42 | 1.30E-23 | 0.00 |
| F.Coll. | SurfaceArea | 0.32 | 6.70E-10 | 0.00 |
| F.I.P. | Length | 0.27 | 2.70E-11 | 0.00 |
| F.I.P. | MeanDepth | 0.42 | 5.10E-14 | 0.01 |
| F.I.P. | Width | 0.67 | 1.90E-30 | 0.00 |
| F.I.P. | SurfaceArea | 0.33 | 1.90E-16 | 0.01 |
| F.I.P.Po.C.inf. | Length | 0.20 | 3.20E-06 | 0.01 |
| F.I.P.Po.C.inf. | MeanDepth | 0.27 | 3.80E-10 | 0.01 |
| F.I.P.Po.C.inf. | Width | 0.48 | 1.10E-15 | 0.00 |
| F.I.P.Po.C.inf. | SurfaceArea | 0.24 | 8.40E-09 | 0.01 |

| | | | | |
|---|---|---|---|---|
| F.P.O. | Length | 0.18 | 6.50E-05 | 0.01 |
| F.P.O. | MeanDepth | 0.47 | 2.90E-26 | 0.01 |
| F.P.O. | Width | 0.47 | 9.90E-32 | 0.00 |
| F.P.O. | SurfaceArea | 0.45 | 4.50E-24 | 0.01 |
| INSULA | Length | - | - | - |
| INSULA | MeanDepth | 0.06 | 5.00E-01 | 0.01 |
| INSULA | Width | 0.36 | 1.40E-02 | 0.10 |
| INSULA | SurfaceArea | - | - | - |
| OCCIPITAL | Length | 0.40 | 5.60E-19 | 0.01 |
| OCCIPITAL | MeanDepth | 0.27 | 6.10E-07 | 0.00 |
| OCCIPITAL | Width | 0.52 | 5.90E-39 | 0.00 |
| OCCIPITAL | SurfaceArea | 0.48 | 5.10E-24 | 0.01 |
| S.C. | Length | 0.17 | 4.10E-05 | 0.01 |
| S.C. | MeanDepth | 0.43 | 1.50E-26 | 0.01 |
| S.C. | Width | 0.72 | 2.00E-32 | 0.00 |
| S.C. | SurfaceArea | 0.44 | 5.90E-27 | 0.01 |
| S.C.LPC. | Length | 0.05 | 1.70E-01 | 3.58 |
| S.C.LPC. | MeanDepth | - | - | - |
| S.C.LPC. | Width | - | - | - |
| S.C.LPC. | SurfaceArea | - | - | - |
| S.C.sylvian. | Length | 0.19 | 1.70E-02 | 0.02 |
| S.C.sylvian. | MeanDepth | 0.28 | 3.40E-04 | 0.01 |
| S.C.sylvian. | Width | 0.37 | 1.10E-06 | 0.02 |
| S.C.sylvian. | SurfaceArea | 0.24 | 1.30E-04 | 0.02 |
| S.Call. | Length | 0.49 | 1.10E-25 | 0.01 |
| S.Call. | MeanDepth | 0.50 | 3.90E-35 | 0.01 |
| S.Call. | Width | 0.33 | 5.20E-13 | 0.01 |
| S.Call. | SurfaceArea | 0.56 | 1.00E-34 | 0.01 |
| S.Cu. | Length | 0.18 | 1.30E-12 | 0.01 |
| S.Cu. | MeanDepth | 0.27 | 6.70E-10 | 0.01 |
| S.Cu. | Width | 0.30 | 7.90E-09 | 0.01 |
| S.Cu. | SurfaceArea | - | - | - |
| S.F.inf. | Length | 0.10 | 6.30E-03 | 0.01 |
| S.F.inf. | MeanDepth | 0.45 | 8.70E-20 | 0.01 |
| S.F.inf. | Width | 0.43 | 2.30E-17 | 0.00 |
| S.F.inf. | SurfaceArea | 0.30 | 1.70E-14 | 0.01 |
| S.F.inf.ant. | Length | 0.12 | 9.30E-04 | 0.01 |
| S.F.inf.ant. | MeanDepth | 0.11 | 7.30E-03 | 0.01 |
| S.F.inf.ant. | Width | 0.17 | 2.50E-05 | 0.01 |
| S.F.inf.ant. | SurfaceArea | 0.17 | 4.70E-06 | 0.01 |
| S.F.int. | Length | 0.32 | 2.10E-09 | 0.01 |
| S.F.int. | MeanDepth | 0.37 | 2.30E-18 | 0.01 |
| S.F.int. | Width | 0.40 | 1.30E-21 | 0.01 |
| S.F.int. | SurfaceArea | 0.35 | 1.40E-13 | 0.01 |
| S.F.inter. | Length | 0.17 | 1.70E-04 | 0.00 |
| S.F.inter. | MeanDepth | 0.20 | 6.10E-08 | 0.01 |
| S.F.inter. | Width | 0.44 | 1.30E-27 | 0.00 |
| S.F.inter. | SurfaceArea | 0.20 | 1.90E-04 | 0.00 |
| S.F.marginal. | Length | 0.19 | 1.10E-05 | 0.00 |
| S.F.marginal. | MeanDepth | 0.09 | 7.70E-03 | 0.01 |
| S.F.marginal. | Width | 0.20 | 5.90E-07 | 0.01 |
| S.F.marginal. | SurfaceArea | 0.21 | 2.80E-06 | 0.01 |
| S.F.median. | Length | 0.26 | 5.90E-10 | 0.01 |
| S.F.median. | MeanDepth | 0.18 | 4.50E-06 | 0.01 |
| S.F.median. | Width | 0.34 | 4.60E-17 | 0.01 |
| S.F.median. | SurfaceArea | 0.27 | 4.10E-10 | 0.01 |
| S.F.orbitaire. | Length | 0.10 | 7.40E-03 | 0.01 |
| S.F.orbitaire. | MeanDepth | 0.12 | 2.10E-03 | 0.01 |
| S.F.orbitaire. | Width | 0.20 | 1.60E-05 | 0.01 |

| | | | | |
|---|---|---|---|---|
| S.F.orbitaire. | SurfaceArea | 0.13 | 4.40E-04 | 0.01 |
| S.F.polaire.tr. | Length | 0.30 | 5.00E-13 | 0.00 |
| S.F.polaire.tr. | MeanDepth | 0.17 | 5.00E-06 | 0.01 |
| S.F.polaire.tr. | Width | 0.34 | 1.20E-09 | 0.01 |
| S.F.polaire.tr. | SurfaceArea | 0.29 | 4.50E-12 | 0.01 |
| S.F.sup. | Length | 0.07 | 1.70E-01 | 0.01 |
| S.F.sup. | MeanDepth | 0.41 | 2.20E-20 | 0.01 |
| S.F.sup. | Width | 0.50 | 1.70E-22 | 0.00 |
| S.F.sup. | SurfaceArea | 0.21 | 3.10E-08 | 0.33 |
| S.Li.post. | Length | 0.15 | 2.70E-05 | 0.01 |
| S.Li.post. | MeanDepth | 0.19 | 1.30E-07 | 0.01 |
| S.Li.post. | Width | 0.25 | 6.40E-03 | 0.01 |
| S.Li.post. | SurfaceArea | 0.14 | 5.10E-07 | 0.01 |
| S.O.p. | Length | 0.10 | 2.40E-03 | 0.01 |
| S.O.p. | MeanDepth | 0.17 | 4.90E-06 | 0.01 |
| S.O.p. | Width | 0.28 | 1.80E-05 | 0.01 |
| S.O.p. | SurfaceArea | 0.16 | 9.50E-03 | 0.22 |
| S.O.T.lat.ant. | Length | 0.26 | 6.00E-02 | 0.01 |
| S.O.T.lat.ant. | MeanDepth | 0.32 | 3.80E-04 | 0.01 |
| S.O.T.lat.ant. | Width | 0.31 | 2.80E-09 | 0.01 |
| S.O.T.lat.ant. | SurfaceArea | 0.32 | 1.60E-03 | 0.01 |
| S.O.T.lat.int. | Length | 0.10 | 1.60E-06 | 0.22 |
| S.O.T.lat.int. | MeanDepth | 0.11 | 1.30E-07 | 0.01 |
| S.O.T.lat.int. | Width | 0.14 | 6.20E-15 | 0.22 |
| S.O.T.lat.int. | SurfaceArea | 0.08 | 2.10E-08 | 0.22 |
| S.O.T.lat.med. | Length | 0.05 | 3.60E-01 | 0.01 |
| S.O.T.lat.med. | MeanDepth | 0.09 | 3.70E-02 | 0.01 |
| S.O.T.lat.med. | Width | 0.14 | 2.20E-01 | 0.01 |
| S.O.T.lat.med. | SurfaceArea | 0.10 | 2.30E-01 | 0.33 |
| S.O.T.lat.post. | Length | 0.02 | 9.70E-02 | 0.01 |
| S.O.T.lat.post. | MeanDepth | 0.21 | 1.30E-02 | 0.01 |
| S.O.T.lat.post. | Width | 0.23 | 2.50E-03 | 0.01 |
| S.O.T.lat.post. | SurfaceArea | 0.15 | 3.10E-02 | 0.54 |
| S.Olf. | Length | 0.31 | 3.20E-01 | 0.00 |
| S.Olf. | MeanDepth | 0.46 | 5.00E-05 | 0.01 |
| S.Olf. | Width | 0.28 | 3.20E-06 | 0.01 |
| S.Olf. | SurfaceArea | 0.44 | 1.40E-02 | 0.01 |
| S.Or. | Length | 0.26 | 7.30E-08 | 0.01 |
| S.Or. | MeanDepth | 0.41 | 2.10E-20 | 0.01 |
| S.Or. | Width | 0.37 | 2.60E-13 | 0.01 |
| S.Or. | SurfaceArea | 0.40 | 1.50E-18 | 0.01 |
| S.p.C. | Length | 0.11 | 1.60E-09 | 0.01 |
| S.p.C. | MeanDepth | 0.01 | 4.50E-21 | 0.22 |
| S.p.C. | Width | 0.14 | 2.10E-16 | 0.01 |
| S.p.C. | SurfaceArea | 0.09 | 7.80E-13 | 0.22 |
| S.Pa.int. | Length | 0.23 | 3.20E-02 | 0.01 |
| S.Pa.int. | MeanDepth | 0.26 | 1.50E-01 | 0.01 |
| S.Pa.int. | Width | 0.37 | 2.20E-02 | 0.01 |
| S.Pa.int. | SurfaceArea | 0.25 | 9.20E-02 | 0.01 |
| S.Pa.sup. | Length | 0.12 | 1.70E-08 | 0.01 |
| S.Pa.sup. | MeanDepth | 0.16 | 7.30E-09 | 0.33 |
| S.Pa.sup. | Width | 0.30 | 4.10E-20 | 0.01 |
| S.Pa.sup. | SurfaceArea | 0.17 | 4.80E-07 | 0.01 |
| S.Pa.t. | Length | 0.06 | 2.80E-02 | 0.02 |
| S.Pa.t. | MeanDepth | 0.17 | 1.20E-02 | 0.01 |
| S.Pa.t. | Width | 0.20 | 3.70E-08 | 0.02 |
| S.Pa.t. | SurfaceArea | 0.14 | 2.10E-02 | 0.34 |
| S.Pe.C.inf. | Length | 0.11 | 1.90E-01 | 0.01 |
| S.Pe.C.inf. | MeanDepth | 0.23 | 9.00E-02 | 0.01 |

| | | | | |
|---|---|---|---|---|
| S.Pe.C.inf. | Width | 0.39 | 9.30E-04 | 0.01 |
| S.Pe.C.inf. | SurfaceArea | 0.12 | 8.80E-03 | 0.01 |
| S.Pe.C.inter. | Length | 0.09 | 1.10E-02 | 0.01 |
| S.Pe.C.inter. | MeanDepth | 0.29 | 5.10E-06 | 0.01 |
| S.Pe.C.inter. | Width | 0.32 | 2.10E-17 | 0.00 |
| S.Pe.C.inter. | SurfaceArea | - | - | - |
| S.Pe.C.marginal. | Length | 0.03 | 2.60E-05 | 0.01 |
| S.Pe.C.marginal. | MeanDepth | - | - | - |
| S.Pe.C.marginal. | Width | 0.31 | 2.80E-15 | 0.01 |
| S.Pe.C.marginal. | SurfaceArea | - | - | - |
| S.Pe.C.median. | Length | 0.08 | 7.60E-02 | 0.01 |
| S.Pe.C.median. | MeanDepth | 0.08 | 2.80E-03 | 0.01 |
| S.Pe.C.median. | Width | 0.24 | 1.30E-08 | 0.01 |
| S.Pe.C.median. | SurfaceArea | 0.11 | 6.10E-03 | 0.01 |
| S.Pe.C.sup. | Length | 0.15 | 2.60E-02 | 0.01 |
| S.Pe.C.sup. | MeanDepth | 0.20 | 7.00E-02 | 0.01 |
| S.Pe.C.sup. | Width | 0.31 | 4.70E-06 | 0.01 |
| S.Pe.C.sup. | SurfaceArea | 0.19 | 1.40E-02 | 0.01 |
| S.Po.C.sup. | Length | 0.16 | 2.40E-04 | 0.01 |
| S.Po.C.sup. | MeanDepth | 0.18 | 2.60E-05 | 0.01 |
| S.Po.C.sup. | Width | 0.43 | 3.90E-12 | 0.01 |
| S.Po.C.sup. | SurfaceArea | 0.17 | 1.80E-06 | 0.01 |
| S.R.inf. | Length | 0.27 | 2.60E-04 | 0.01 |
| S.R.inf. | MeanDepth | 0.18 | 7.90E-06 | 0.01 |
| S.R.inf. | Width | 0.15 | 5.20E-21 | 0.01 |
| S.R.inf. | SurfaceArea | 0.29 | 6.40E-05 | 0.01 |
| S.Rh. | Length | 0.20 | 1.00E-06 | 0.01 |
| S.Rh. | MeanDepth | 0.21 | 2.90E-04 | 0.01 |
| S.Rh. | Width | 0.24 | 2.30E-04 | 0.01 |
| S.Rh. | SurfaceArea | 0.19 | 1.10E-06 | 0.01 |
| S.s.P. | Length | 0.21 | 1.80E-03 | 0.01 |
| S.s.P. | MeanDepth | 0.40 | 1.20E-04 | 0.01 |
| S.s.P. | Width | 0.39 | 1.10E-06 | 0.00 |
| S.s.P. | SurfaceArea | 0.28 | 4.50E-04 | 0.01 |
| S.T.i.ant. | Length | 0.27 | 3.00E-05 | 0.01 |
| S.T.i.ant. | MeanDepth | 0.28 | 1.10E-18 | 0.01 |
| S.T.i.ant. | Width | 0.47 | 3.70E-19 | 0.01 |
| S.T.i.ant. | SurfaceArea | 0.35 | 2.30E-09 | 0.01 |
| S.T.i.post. | Length | 0.04 | 2.00E-07 | 0.01 |
| S.T.i.post. | MeanDepth | 0.28 | 3.20E-07 | 0.01 |
| S.T.i.post. | Width | 0.41 | 1.30E-21 | 0.00 |
| S.T.i.post. | SurfaceArea | 0.15 | 3.60E-15 | 0.01 |
| S.T.pol. | Length | 0.34 | 1.30E-01 | 0.01 |
| S.T.pol. | MeanDepth | 0.21 | 3.80E-12 | 0.01 |
| S.T.pol. | Width | 0.31 | 6.10E-24 | 0.01 |
| S.T.pol. | SurfaceArea | 0.37 | 1.70E-04 | 0.01 |
| S.T.s. | Length | 0.16 | 3.00E-12 | 0.01 |
| S.T.s. | MeanDepth | 0.44 | 5.20E-05 | 0.01 |
| S.T.s. | Width | 0.36 | 4.80E-13 | 0.00 |
| S.T.s. | SurfaceArea | 0.24 | 1.20E-11 | 0.01 |
| S.T.s.ter.asc.ant. | Length | - | - | - |
| S.T.s.ter.asc.ant. | MeanDepth | - | - | - |
| S.T.s.ter.asc.ant. | Width | 0.23 | 1.80E-18 | 0.01 |
| S.T.s.ter.asc.ant. | SurfaceArea | - | - | - |
| S.T.s.ter.asc.post. | Length | 0.03 | 2.50E-04 | 0.01 |
| S.T.s.ter.asc.post. | MeanDepth | 0.17 | 2.90E-03 | 0.01 |
| S.T.s.ter.asc.post. | Width | 0.28 | 1.10E-04 | 0.01 |
| S.T.s.ter.asc.post. | SurfaceArea | - | - | - |
| Globals | Length | 0.23 | 2.70E-02 | 0.01 |

| | | | | |
|---|---|---|---|---|
| Globals | MeanDepth | 0.19 | 1.60E-05 | 0.01 |
| Globals | Width | 0.30 | 1.90E-11 | 0.01 |
| Globals | SurfaceArea | 0.34 | 2.00E-04 | 0.01 |

**Table S21:** Mega-analysis of heritability eatimates for bilaterally average sulcal descriptors (yellow: bonferroni corrected for 61*4 comparisons).

| Sulci | Descriptor | h2 | pval | SE |
|---|---|---|---|---|
| F.C.L.a. | Length | 0.06 | 9.40E-02 | 0.05 |
| F.C.L.a. | MeanDepth | 0.23 | 7.30E-06 | 0.06 |
| F.C.L.a. | Width | 0.36 | 5.00E-10 | 0.06 |
| F.C.L.a. | SurfaceArea | 0.26 | 3.60E-07 | 0.06 |
| F.C.L.p. | Length | 0.18 | 2.30E-04 | 0.06 |
| F.C.L.p. | MeanDepth | 0.22 | 2.30E-05 | 0.06 |
| F.C.L.p. | Width | 0.47 | 6.30E-21 | 0.05 |
| F.C.L.p. | SurfaceArea | 0.45 | 2.00E-17 | 0.05 |
| F.C.L.r.ant. | Length | 0.19 | 1.60E-03 | 0.07 |
| F.C.L.r.ant. | MeanDepth | 0.11 | 2.00E-02 | 0.06 |
| F.C.L.r.ant. | Width | 0.20 | 3.00E-04 | 0.06 |
| F.C.L.r.ant. | SurfaceArea | 0.34 | 2.00E-08 | 0.07 |
| F.C.L.r.asc. | Length | 0.14 | 1.50E-03 | 0.05 |
| F.C.L.r.asc. | MeanDepth | 0.19 | 1.90E-05 | 0.05 |
| F.C.L.r.asc. | Width | 0.19 | 2.40E-05 | 0.05 |
| F.C.L.r.asc. | SurfaceArea | 0.30 | 1.80E-10 | 0.05 |
| F.C.L.r.diag. | Length | 0.04 | 3.90E-01 | 0.17 |
| F.C.L.r.diag. | MeanDepth | 0.08 | 3.30E-01 | 0.17 |
| F.C.L.r.diag. | Width | 0.09 | 2.70E-01 | 0.15 |
| F.C.L.r.diag. | SurfaceArea | 0.04 | 4.00E-01 | 0.16 |
| F.C.L.r.retroC.tr. | Length | 0.12 | 1.60E-02 | 0.06 |
| F.C.L.r.retroC.tr. | MeanDepth | 0.29 | 2.80E-08 | 0.06 |
| F.C.L.r.retroC.tr. | Width | 0.17 | 1.20E-03 | 0.06 |
| F.C.L.r.retroC.tr. | SurfaceArea | 0.20 | 6.50E-05 | 0.06 |
| F.C.L.r.sc.ant. | Length | - | - | - |
| F.C.L.r.sc.ant. | MeanDepth | 0.78 | 3.10E-01 | 1.38 |
| F.C.L.r.sc.ant. | Width | - | - | - |
| F.C.L.r.sc.ant. | SurfaceArea | - | - | - |
| F.C.L.r.sc.post. | Length | 1.00 | 4.00E-01 | - |
| F.C.L.r.sc.post. | MeanDepth | 0.10 | 7.80E-02 | 0.07 |
| F.C.L.r.sc.post. | Width | 0.28 | 1.20E-04 | 0.08 |
| F.C.L.r.sc.post. | SurfaceArea | 0.02 | 3.70E-01 | 0.07 |
| F.C.M.ant. | Length | 0.09 | 1.60E-02 | 0.04 |
| F.C.M.ant. | MeanDepth | 0.32 | 1.70E-11 | 0.05 |
| F.C.M.ant. | Width | 0.30 | 1.40E-10 | 0.05 |
| F.C.M.ant. | SurfaceArea | 0.22 | 3.60E-07 | 0.05 |
| F.C.M.post. | Length | 0.02 | 3.40E-01 | 0.05 |
| F.C.M.post. | MeanDepth | 0.37 | 5.10E-15 | 0.05 |
| F.C.M.post. | Width | 0.43 | 2.40E-17 | 0.05 |
| F.C.M.post. | SurfaceArea | 0.25 | 3.00E-08 | 0.05 |
| F.Cal.ant.-Sc.Cal. | Length | - | - | - |
| F.Cal.ant.-Sc.Cal. | MeanDepth | - | - | - |
| F.Cal.ant.-Sc.Cal. | Width | - | - | - |

| | | | | |
|---|---|---|---|---|
| F.Cal.ant.-Sc.Cal. | SurfaceArea | 0.50 | 3.90E-24 | 0.05 |
| F.Coll. | Length | 0.10 | 1.30E-02 | 0.05 |
| F.Coll. | MeanDepth | 0.46 | 3.00E-26 | 0.05 |
| F.Coll. | Width | 0.41 | 3.90E-20 | 0.05 |
| F.Coll. | SurfaceArea | 0.29 | 6.30E-11 | 0.05 |
| F.I.P. | Length | 0.32 | 2.60E-11 | 0.05 |
| F.I.P. | MeanDepth | 0.40 | 6.60E-16 | 0.05 |
| F.I.P. | Width | 0.57 | 2.00E-25 | 0.05 |
| F.I.P. | SurfaceArea | 0.32 | 1.70E-11 | 0.05 |
| F.I.P.Po.C.inf. | Length | 0.23 | 4.40E-06 | 0.06 |
| F.I.P.Po.C.inf. | MeanDepth | 0.24 | 1.00E-06 | 0.06 |
| F.I.P.Po.C.inf. | Width | 0.42 | 2.00E-15 | 0.06 |
| F.I.P.Po.C.inf. | SurfaceArea | 0.24 | 5.40E-07 | 0.05 |
| F.I.P.r.int.1 | Length | 0.11 | 3.10E-02 | 0.06 |
| F.I.P.r.int.1 | MeanDepth | 0.05 | 1.90E-01 | 0.05 |
| F.I.P.r.int.1 | Width | 0.15 | 1.10E-03 | 0.05 |
| F.I.P.r.int.1 | SurfaceArea | 0.09 | 4.10E-02 | 0.06 |
| F.I.P.r.int.2 | Length | 0.08 | 1.50E-01 | 0.08 |
| F.I.P.r.int.2 | MeanDepth | 0.05 | 2.20E-01 | 0.07 |
| F.I.P.r.int.2 | Width | 0.08 | 1.50E-01 | 0.09 |
| F.I.P.r.int.2 | SurfaceArea | 0.07 | 1.40E-01 | 0.07 |
| F.P.O. | Length | 0.16 | 5.20E-04 | 0.05 |
| F.P.O. | MeanDepth | 0.46 | 5.10E-20 | 0.05 |
| F.P.O. | Width | 0.42 | 6.30E-19 | 0.05 |
| F.P.O. | SurfaceArea | 0.47 | 4.90E-21 | 0.05 |
| INSULA | Length | - | - | - |
| INSULA | MeanDepth | 0.19 | 1.00E-04 | 0.06 |
| INSULA | Width | 0.32 | 5.10E-09 | 0.06 |
| INSULA | SurfaceArea | 0.54 | 2.20E-22 | 0.05 |
| OCCIPITAL | Length | 0.36 | 1.00E-12 | 0.05 |
| OCCIPITAL | MeanDepth | 0.29 | 2.90E-11 | 0.05 |
| OCCIPITAL | Width | 0.57 | 1.80E-29 | 0.05 |
| OCCIPITAL | SurfaceArea | 0.41 | 1.20E-15 | 0.05 |
| S.C. | Length | 0.21 | 2.60E-06 | 0.05 |
| S.C. | MeanDepth | 0.40 | 6.00E-18 | 0.05 |
| S.C. | Width | 0.57 | 1.30E-26 | 0.05 |
| S.C. | SurfaceArea | 0.51 | 3.40E-25 | 0.05 |
| S.C.LPC. | Length | - | - | - |
| S.C.LPC. | MeanDepth | - | - | - |
| S.C.LPC. | Width | 0.12 | 4.00E-01 | 0.46 |
| S.C.LPC. | SurfaceArea | 0.92 | 2.00E-01 | 0.15 |
| S.C.sylvian. | Length | 0.19 | 2.70E-02 | 0.10 |
| S.C.sylvian. | MeanDepth | 0.26 | 2.80E-03 | 0.10 |
| S.C.sylvian. | Width | 0.22 | 7.80E-03 | 0.09 |
| S.C.sylvian. | SurfaceArea | 0.20 | 7.50E-03 | 0.09 |
| S.Call. | Length | 0.49 | 8.20E-22 | 0.05 |

| | | | | |
|---|---|---|---|---|
| S.Call. | MeanDepth | 0.57 | 2.50E-25 | 0.05 |
| S.Call. | Width | 0.33 | 8.60E-13 | 0.05 |
| S.Call. | SurfaceArea | 0.58 | 7.80E-28 | 0.05 |
| S.Cu. | Length | 0.06 | 1.30E-01 | 0.05 |
| S.Cu. | MeanDepth | 0.25 | 1.40E-07 | 0.05 |
| S.Cu. | Width | 0.29 | 1.10E-08 | 0.06 |
| S.Cu. | SurfaceArea | 0.09 | 2.80E-02 | 0.05 |
| S.F.inf. | Length | 0.20 | 4.60E-05 | 0.05 |
| S.F.inf. | MeanDepth | 0.43 | 6.20E-17 | 0.05 |
| S.F.inf. | Width | 0.39 | 2.50E-15 | 0.05 |
| S.F.inf. | SurfaceArea | 0.31 | 1.30E-10 | 0.05 |
| S.F.inf.ant. | Length | 0.13 | 2.40E-03 | 0.05 |
| S.F.inf.ant. | MeanDepth | 0.12 | 7.70E-03 | 0.05 |
| S.F.inf.ant. | Width | 0.15 | 1.30E-03 | 0.05 |
| S.F.inf.ant. | SurfaceArea | 0.19 | 2.80E-05 | 0.05 |
| S.F.int. | Length | 0.35 | 2.50E-12 | 0.05 |
| S.F.int. | MeanDepth | 0.36 | 5.90E-13 | 0.05 |
| S.F.int. | Width | 0.33 | 2.60E-11 | 0.05 |
| S.F.int. | SurfaceArea | 0.36 | 1.00E-12 | 0.05 |
| S.F.inter. | Length | 0.19 | 2.90E-06 | 0.05 |
| S.F.inter. | MeanDepth | 0.24 | 1.90E-07 | 0.05 |
| S.F.inter. | Width | 0.41 | 8.20E-18 | 0.05 |
| S.F.inter. | SurfaceArea | 0.25 | 6.40E-09 | 0.05 |
| S.F.marginal. | Length | - | - | - |
| S.F.marginal. | MeanDepth | - | - | - |
| S.F.marginal. | Width | - | - | - |
| S.F.marginal. | SurfaceArea | - | - | - |
| S.F.median. | Length | 0.30 | 7.90E-09 | 0.06 |
| S.F.median. | MeanDepth | 0.18 | 4.50E-05 | 0.05 |
| S.F.median. | Width | 0.35 | 3.90E-12 | 0.06 |
| S.F.median. | SurfaceArea | 0.30 | 1.90E-09 | 0.06 |
| S.F.orbitaire. | Length | 0.15 | 2.30E-03 | 0.06 |
| S.F.orbitaire. | MeanDepth | 0.14 | 3.60E-03 | 0.06 |
| S.F.orbitaire. | Width | 0.21 | 4.50E-05 | 0.06 |
| S.F.orbitaire. | SurfaceArea | 0.17 | 7.00E-04 | 0.06 |
| S.F.polaire.tr. | Length | 0.27 | 8.80E-09 | 0.05 |
| S.F.polaire.tr. | MeanDepth | 0.14 | 1.50E-03 | 0.05 |
| S.F.polaire.tr. | Width | 0.28 | 2.50E-07 | 0.06 |
| S.F.polaire.tr. | SurfaceArea | 0.26 | 2.40E-07 | 0.06 |
| S.F.sup. | Length | 0.11 | 3.70E-03 | 0.05 |
| S.F.sup. | MeanDepth | 0.44 | 2.60E-19 | 0.05 |
| S.F.sup. | Width | 0.41 | 2.20E-18 | 0.05 |
| S.F.sup. | SurfaceArea | 0.28 | 3.80E-09 | 0.05 |
| S.Li.ant. | Length | 0.21 | 4.60E-05 | 0.06 |
| S.Li.ant. | MeanDepth | 0.24 | 1.10E-04 | 0.07 |
| S.Li.ant. | Width | 0.14 | 1.10E-02 | 0.07 |

| | | | | |
|---|---|---|---|---|
| S.Li.ant. | SurfaceArea | 0.24 | 1.50E-05 | 0.06 |
| S.Li.post. | Length | 0.08 | 6.30E-02 | 0.05 |
| S.Li.post. | MeanDepth | 0.16 | 1.10E-03 | 0.06 |
| S.Li.post. | Width | 0.16 | 2.10E-03 | 0.06 |
| S.Li.post. | SurfaceArea | 0.06 | 1.40E-01 | 0.06 |
| S.O.p. | Length | 0.10 | 3.40E-02 | 0.06 |
| S.O.p. | MeanDepth | 0.17 | 2.20E-03 | 0.06 |
| S.O.p. | Width | 0.28 | 3.00E-07 | 0.06 |
| S.O.p. | SurfaceArea | 0.17 | 1.50E-03 | 0.06 |
| S.O.T.lat.ant. | Length | 0.29 | 1.30E-09 | 0.05 |
| S.O.T.lat.ant. | MeanDepth | 0.33 | 8.60E-13 | 0.05 |
| S.O.T.lat.ant. | Width | 0.31 | 2.80E-11 | 0.05 |
| S.O.T.lat.ant. | SurfaceArea | 0.34 | 1.40E-13 | 0.05 |
| S.O.T.lat.int. | Length | 0.10 | 4.10E-02 | 0.06 |
| S.O.T.lat.int. | MeanDepth | 0.10 | 5.00E-02 | 0.07 |
| S.O.T.lat.int. | Width | 0.07 | 1.20E-01 | 0.06 |
| S.O.T.lat.int. | SurfaceArea | 0.04 | 2.60E-01 | 0.07 |
| S.O.T.lat.med. | Length | 0.04 | 1.80E-01 | 0.05 |
| S.O.T.lat.med. | MeanDepth | 0.12 | 5.80E-03 | 0.05 |
| S.O.T.lat.med. | Width | 0.15 | 1.00E-03 | 0.05 |
| S.O.T.lat.med. | SurfaceArea | 0.09 | 2.90E-02 | 0.05 |
| S.O.T.lat.post. | Length | 0.02 | 3.40E-01 | 0.05 |
| S.O.T.lat.post. | MeanDepth | 0.18 | 1.90E-04 | 0.06 |
| S.O.T.lat.post. | Width | 0.18 | 1.10E-04 | 0.05 |
| S.O.T.lat.post. | SurfaceArea | 0.13 | 3.50E-03 | 0.05 |
| S.Olf. | Length | 0.36 | 1.40E-15 | 0.05 |
| S.Olf. | MeanDepth | 0.49 | 4.20E-23 | 0.05 |
| S.Olf. | Width | 0.24 | 1.40E-07 | 0.05 |
| S.Olf. | SurfaceArea | 0.47 | 9.30E-23 | 0.05 |
| S.Or. | Length | 0.27 | 2.30E-08 | 0.05 |
| S.Or. | MeanDepth | 0.35 | 1.20E-12 | 0.05 |
| S.Or. | Width | 0.35 | 4.10E-12 | 0.06 |
| S.Or. | SurfaceArea | 0.41 | 4.00E-13 | 0.06 |
| S.p.C. | Length | 0.07 | 1.20E-01 | 0.06 |
| S.p.C. | MeanDepth | 0.07 | 1.50E-01 | 0.07 |
| S.p.C. | Width | 0.11 | 4.50E-02 | 0.07 |
| S.p.C. | SurfaceArea | 0.04 | 2.50E-01 | 0.06 |
| S.Pa.int. | Length | 0.22 | 5.90E-06 | 0.06 |
| S.Pa.int. | MeanDepth | 0.23 | 2.70E-06 | 0.05 |
| S.Pa.int. | Width | 0.34 | 2.70E-13 | 0.05 |
| S.Pa.int. | SurfaceArea | 0.20 | 3.30E-05 | 0.05 |
| S.Pa.sup. | Length | 0.14 | 6.30E-03 | 0.06 |
| S.Pa.sup. | MeanDepth | 0.17 | 1.40E-03 | 0.06 |
| S.Pa.sup. | Width | 0.36 | 1.70E-09 | 0.06 |
| S.Pa.sup. | SurfaceArea | 0.20 | 1.60E-04 | 0.06 |
| S.Pa.t. | Length | 0.14 | 1.30E-01 | 0.13 |

| | | | | |
|---|---|---|---|---|
| S.Pa.t. | MeanDepth | 0.13 | 3.70E-02 | 0.08 |
| S.Pa.t. | Width | 0.16 | 1.90E-02 | 0.08 |
| S.Pa.t. | SurfaceArea | 0.20 | 3.70E-03 | 0.08 |
| S.Pe.C.inf. | Length | 0.14 | 2.60E-03 | 0.05 |
| S.Pe.C.inf. | MeanDepth | 0.28 | 1.70E-07 | 0.06 |
| S.Pe.C.inf. | Width | 0.41 | 5.90E-14 | 0.05 |
| S.Pe.C.inf. | SurfaceArea | 0.13 | 3.90E-03 | 0.05 |
| S.Pe.C.inter. | Length | 0.03 | 2.40E-01 | 0.05 |
| S.Pe.C.inter. | MeanDepth | 0.32 | 1.10E-10 | 0.06 |
| S.Pe.C.inter. | Width | 0.31 | 9.10E-11 | 0.05 |
| S.Pe.C.inter. | SurfaceArea | 0.16 | 2.40E-04 | 0.05 |
| S.Pe.C.marginal. | Length | - | - | - |
| S.Pe.C.marginal. | MeanDepth | - | - | - |
| S.Pe.C.marginal. | Width | - | - | - |
| S.Pe.C.marginal. | SurfaceArea | - | - | - |
| S.Pe.C.median. | Length | 0.12 | 1.90E-02 | 0.06 |
| S.Pe.C.median. | MeanDepth | 0.09 | 6.30E-02 | 0.06 |
| S.Pe.C.median. | Width | 0.27 | 6.80E-07 | 0.06 |
| S.Pe.C.median. | SurfaceArea | 0.13 | 1.20E-02 | 0.06 |
| S.Pe.C.sup. | Length | 0.17 | 6.90E-04 | 0.06 |
| S.Pe.C.sup. | MeanDepth | 0.20 | 5.20E-05 | 0.06 |
| S.Pe.C.sup. | Width | 0.28 | 7.50E-09 | 0.05 |
| S.Pe.C.sup. | SurfaceArea | 0.18 | 2.90E-04 | 0.06 |
| S.Po.C.sup. | Length | 0.16 | 6.50E-04 | 0.05 |
| S.Po.C.sup. | MeanDepth | 0.17 | 1.60E-04 | 0.05 |
| S.Po.C.sup. | Width | 0.34 | 3.70E-11 | 0.06 |
| S.Po.C.sup. | SurfaceArea | 0.18 | 7.70E-05 | 0.05 |
| S.R.inf. | Length | 0.24 | 5.70E-05 | 0.07 |
| S.R.inf. | MeanDepth | 0.15 | 6.10E-03 | 0.06 |
| S.R.inf. | Width | 0.12 | 1.60E-02 | 0.06 |
| S.R.inf. | SurfaceArea | 0.26 | 3.10E-05 | 0.07 |
| S.Rh. | Length | 0.22 | 8.50E-06 | 0.06 |
| S.Rh. | MeanDepth | 0.24 | 6.70E-06 | 0.06 |
| S.Rh. | Width | 0.26 | 1.80E-06 | 0.06 |
| S.Rh. | SurfaceArea | 0.21 | 3.50E-05 | 0.06 |
| S.s.P. | Length | 0.22 | 2.20E-05 | 0.06 |
| S.s.P. | MeanDepth | 0.39 | 2.00E-14 | 0.05 |
| S.s.P. | Width | 0.35 | 1.60E-12 | 0.05 |
| S.s.P. | SurfaceArea | 0.28 | 7.00E-08 | 0.06 |
| S.T.i.ant. | Length | 0.29 | 2.20E-09 | 0.05 |
| S.T.i.ant. | MeanDepth | 0.28 | 8.40E-10 | 0.05 |
| S.T.i.ant. | Width | 0.41 | 4.80E-16 | 0.05 |
| S.T.i.ant. | SurfaceArea | 0.38 | 1.40E-15 | 0.05 |
| S.T.i.post. | Length | 0.03 | 2.20E-01 | 0.05 |
| S.T.i.post. | MeanDepth | 0.27 | 1.90E-08 | 0.05 |
| S.T.i.post. | Width | 0.33 | 1.50E-13 | 0.05 |

| | | | | |
|---|---|---|---|---|
| S.T.i.post. | SurfaceArea | 0.15 | 8.10E-04 | 0.05 |
| S.T.pol. | Length | 0.28 | 2.70E-09 | 0.05 |
| S.T.pol. | MeanDepth | 0.24 | 7.80E-07 | 0.05 |
| S.T.pol. | Width | 0.32 | 2.50E-13 | 0.05 |
| S.T.pol. | SurfaceArea | 0.33 | 3.20E-12 | 0.05 |
| S.T.s. | Length | 0.21 | 3.80E-06 | 0.05 |
| S.T.s. | MeanDepth | 0.46 | 3.40E-24 | 0.05 |
| S.T.s. | Width | 0.31 | 7.50E-12 | 0.05 |
| S.T.s. | SurfaceArea | 0.32 | 8.80E-11 | 0.05 |
| S.T.s.ter.asc.ant. | Length | 0.04 | 2.30E-01 | 0.05 |
| S.T.s.ter.asc.ant. | MeanDepth | - | - | - |
| S.T.s.ter.asc.ant. | Width | 0.18 | 1.90E-04 | 0.05 |
| S.T.s.ter.asc.ant. | SurfaceArea | - | - | - |
| S.T.s.ter.asc.post. | Length | 0.04 | 1.70E-01 | 0.05 |
| S.T.s.ter.asc.post. | MeanDepth | 0.18 | 2.20E-04 | 0.05 |
| S.T.s.ter.asc.post. | Width | 0.21 | 3.40E-06 | 0.05 |
| S.T.s.ter.asc.post. | SurfaceArea | 0.19 | 8.90E-05 | 0.05 |
| Globals | Length | 0.28 | 1.10E-08 | 0.06 |
| Globals | MeanDepth | 0.17 | 1.30E-04 | 0.05 |
| Globals | Width | 0.32 | 2.50E-14 | 0.05 |
| Globals | SurfaceArea | 0.41 | 1.00E-17 | 0.05 |

**Table S22:** Bivariate analysis: left-right genetic correlation in QTIM (yellow: Bonferroni corrected; grey: $p < 0.05$).

| Sulci | Descriptor | rhoG | rhoG_SE | rhoP=0 | rhoP=1 | rhoE | rhoE.se | rhoE.p |
|---|---|---|---|---|---|---|---|---|
| F.C.L.a. | Length | 1.00 | - | 8.75E-02 | - | -0.08 | 0.06 | 1.94E-01 |
| F.C.L.a. | MeanDepth | -0.04 | 0.52 | 9.31E-01 | - | 0.07 | 0.08 | 3.56E-01 |
| F.C.L.a. | Width | 0.60 | 0.39 | 1.57E-01 | 2.12E-01 | 0.20 | 0.07 | 5.14E-03 |
| F.C.L.a. | SurfaceArea | 1.00 | - | 5.54E-02 | - | 0.05 | 0.06 | 3.69E-01 |
| F.C.L.p. | Length | 0.25 | 0.92 | 7.52E-01 | 4.23E-01 | 0.14 | 0.07 | 6.22E-02 |
| F.C.L.p. | MeanDepth | 1.00 | - | 1.10E-02 | - | -0.05 | 0.06 | 3.87E-01 |
| F.C.L.p. | Width | 0.89 | 0.07 | 4.02E-12 | 7.44E-02 | 0.23 | 0.08 | 3.18E-03 |
| F.C.L.p. | SurfaceArea | 0.95 | 0.18 | 8.73E-06 | 3.93E-01 | 0.05 | 0.08 | 5.45E-01 |
| F.C.L.r.ant. | Length | 0.88 | 0.49 | 3.34E-02 | 4.11E-01 | -0.01 | 0.09 | 9.44E-01 |
| F.C.L.r.ant. | MeanDepth | 0.40 | 0.54 | 4.50E-01 | 2.23E-01 | 0.12 | 0.10 | 2.21E-01 |
| F.C.L.r.ant. | Width | 0.93 | 0.67 | 5.40E-02 | 4.61E-01 | 0.18 | 0.09 | 3.88E-02 |
| F.C.L.r.ant. | SurfaceArea | 0.78 | 0.29 | 5.12E-03 | 2.45E-01 | 0.03 | 0.09 | 7.83E-01 |
| F.C.L.r.asc. | Length | -0.06 | 1.11 | 9.56E-01 | - | 0.04 | 0.07 | 6.21E-01 |
| F.C.L.r.asc. | MeanDepth | 1.00 | - | 6.44E-02 | - | 0.12 | 0.06 | 5.43E-02 |
| F.C.L.r.asc. | Width | 1.00 | - | 3.29E-03 | - | 0.10 | 0.06 | 1.33E-01 |
| F.C.L.r.asc. | SurfaceArea | 0.33 | 0.35 | 3.55E-01 | 1.52E-01 | 0.12 | 0.08 | 1.18E-01 |
| F.C.L.r.diag. | Length | 0.00 | 1.25 | 9.98E-01 | 0.3494618 | 0.14 | 0.38 | 7.26E-01 |
| F.C.L.r.diag. | MeanDepth | 1.00 | - | 6.00E-01 | - | 0.12 | 0.16 | 4.73E-01 |
| F.C.L.r.diag. | Width | -0.16 | 0.91 | 8.35E-01 | - | 0.63 | 0.19 | 1.95E-02 |
| F.C.L.r.diag. | SurfaceArea | 1.00 | - | 6.65E-01 | - | -0.01 | 0.17 | 9.59E-01 |
| F.C.L.r.retroC.tr. | Length | -0.90 | - | 9.32E-01 | - | 0.03 | - | 4.93E-01 |
| F.C.L.r.retroC.tr. | MeanDepth | 0.88 | 0.53 | 5.73E-02 | 4.16E-01 | 0.02 | 0.10 | 8.69E-01 |
| F.C.L.r.retroC.tr. | Width | 0.50 | 0.41 | 1.77E-01 | 2.45E-01 | 0.14 | 0.11 | 1.96E-01 |
| F.C.L.r.retroC.tr. | SurfaceArea | 0.08 | 0.37 | 8.40E-01 | 6.48E-02 | 0.12 | 0.09 | 1.92E-01 |
| F.C.L.r.sc.ant. | Length | - | - | - | - | - | - | - |
| F.C.L.r.sc.ant. | MeanDepth | - | 0.53 | - | - | - | - | - |
| F.C.L.r.sc.ant. | Width | - | 0.41 | - | - | - | - | - |
| F.C.L.r.sc.ant. | SurfaceArea | - | - | - | - | - | - | - |
| F.C.L.r.sc.post. | Length | - | - | - | - | - | - | - |
| F.C.L.r.sc.post. | MeanDepth | - | 0.53 | - | - | - | - | - |
| F.C.L.r.sc.post. | Width | - | 0.41 | - | - | - | - | - |
| F.C.L.r.sc.post. | SurfaceArea | - | - | - | - | - | - | - |
| F.C.M.ant. | Length | -0.02 | - | 1.00E+00 | - | -0.08 | - | 3.37E-01 |
| F.C.M.ant. | MeanDepth | 0.44 | 0.21 | 1.42E-02 | 2.74E-02 | -0.22 | 0.08 | 5.42E-03 |
| F.C.M.ant. | Width | 1.00 | - | 4.89E-05 | - | -0.11 | 0.06 | 1.22E-01 |
| F.C.M.ant. | SurfaceArea | 1.00 | - | 5.83E-03 | - | -0.17 | 0.06 | 4.10E-03 |
| F.C.M.post. | Length | 1.00 | - | 6.30E-01 | - | -0.03 | 0.06 | 6.09E-01 |
| F.C.M.post. | MeanDepth | 0.84 | 0.16 | 5.66E-05 | 1.86E-01 | 0.17 | 0.07 | 2.27E-02 |
| F.C.M.post. | Width | 0.91 | 0.11 | 1.29E-10 | 2.10E-01 | 0.09 | 0.08 | 2.52E-01 |
| F.C.M.post. | SurfaceArea | 0.68 | 0.31 | 1.87E-02 | 2.04E-01 | 0.00 | 0.08 | 9.92E-01 |
| F.Cal.ant.-Sc.Cal. | Length | 1.00 | - | 1.42E-01 | - | 0.01 | 0.07 | 8.24E-01 |
| F.Cal.ant.-Sc.Cal. | MeanDepth | 0.87 | 0.14 | 2.10E-09 | 0.1842361 | 0.03 | 0.08 | 7.26E-01 |
| F.Cal.ant.-Sc.Cal. | Width | 1.00 | - | 8.77E-07 | - | -0.08 | 0.06 | 1.75E-01 |
| F.Cal.ant.-Sc.Cal. | SurfaceArea | 0.78 | 0.10 | 1.92E-09 | 0.0253325 | 0.17 | 0.08 | 3.32E-02 |
| F.Coll. | Length | 0.65 | 0.44 | 1.29E-01 | 2.50E-01 | 0.01 | 0.07 | 8.45E-01 |
| F.Coll. | MeanDepth | 0.85 | 0.09 | 1.63E-10 | 5.79E-02 | 0.12 | 0.08 | 1.19E-01 |
| F.Coll. | Width | 0.72 | 0.17 | 3.49E-04 | 7.59E-02 | 0.13 | 0.07 | 7.64E-02 |
| F.Coll. | SurfaceArea | 1.00 | - | 1.17E-06 | - | 0.06 | 0.06 | 3.87E-01 |
| F.I.P. | Length | 1.00 | - | 1.17E-02 | - | 0.00 | 0.06 | 9.46E-01 |
| F.I.P. | MeanDepth | 0.94 | 0.15 | 1.18E-09 | 3.37E-01 | -0.01 | 0.08 | 9.26E-01 |
| F.I.P. | Width | 0.96 | 0.03 | 3.78E-24 | 1.05E-01 | 0.22 | 0.08 | 5.52E-03 |
| F.I.P. | SurfaceArea | 1.00 | - | 4.92E-05 | - | 0.11 | 0.06 | 7.85E-02 |
| F.I.P.Po.C.inf. | Length | 1.00 | - | 9.02E-03 | - | 0.01 | 0.06 | 9.10E-01 |
| F.I.P.Po.C.inf. | MeanDepth | 1.00 | - | 2.34E-03 | - | 0.01 | 0.06 | 8.95E-01 |
| F.I.P.Po.C.inf. | Width | 1.00 | - | 7.06E-13 | - | 0.14 | 0.07 | 3.96E-02 |
| F.I.P.Po.C.inf. | SurfaceArea | 0.74 | 0.39 | 4.14E-02 | 2.81E-01 | 0.07 | 0.07 | 3.32E-01 |
| F.I.P.r.int.1 | Length | 0.24 | 0.37 | 4.97E-01 | 0.0919835 | -0.11 | 0.09 | 2.17E-01 |
| F.I.P.r.int.1 | MeanDepth | 1.00 | - | 3.43E-01 | - | -0.13 | 0.06 | 3.33E-02 |
| F.I.P.r.int.1 | Width | 1.00 | - | 2.40E-01 | - | -0.05 | 0.07 | 5.05E-01 |
| F.I.P.r.int.1 | SurfaceArea | 0.38 | 0.89 | 6.17E-01 | 0.323528 | -0.11 | 0.08 | 1.70E-01 |
| F.I.P.r.int.2 | Length | -1.00 | - | 8.72E-01 | - | 0.09 | 0.09 | 1.14E-01 |
| F.I.P.r.int.2 | MeanDepth | -0.41 | 1.10 | 6.65E-01 | - | 0.11 | 0.13 | 3.64E-01 |

| | | | | | | | | |
|---|---|---|---|---|---|---|---|---|
| F.I.P.r.int.2 | Width | 0.40 | 0.68 | 5.74E-01 | 0.2600942 | -0.05 | 0.14 | 7.35E-01 |
| F.I.P.r.int.2 | SurfaceArea | 0.15 | - | 1.00E+00 | 1 | 0.05 | - | 3.23E-01 |
| F.P.O. | Length | 0.41 | 0.37 | 2.53E-01 | 1.73E-01 | 0.01 | 0.07 | 9.04E-01 |
| F.P.O. | MeanDepth | 1.00 | - | 1.63E-06 | - | -0.02 | 0.07 | 7.86E-01 |
| F.P.O. | Width | 0.77 | 0.13 | 4.17E-08 | 6.01E-02 | -0.03 | 0.08 | 7.47E-01 |
| F.P.O. | SurfaceArea | 0.97 | 0.20 | 4.54E-07 | 4.42E-01 | 0.01 | 0.07 | 9.26E-01 |
| INSULA | Length | -0.07 | - | 1.00E+00 | - | -0.08 | - | 4.29E-01 |
| INSULA | MeanDepth | 1.00 | - | 5.76E-01 | - | 0.44 | 0.25 | 1.52E-01 |
| INSULA | Width | 0.41 | 0.46 | 4.42E-01 | 9.81E-02 | 0.84 | 0.22 | 9.44E-02 |
| INSULA | SurfaceArea | 0.58 | 0.31 | 9.51E-02 | 1.02E-01 | -0.17 | 0.35 | 6.12E-01 |
| OCCIPITAL | Length | 1.00 | - | 1.81E-15 | - | 0.13 | 0.07 | 5.90E-02 |
| OCCIPITAL | MeanDepth | 0.23 | 0.37 | 5.65E-01 | 4.15E-02 | 0.16 | 0.07 | 2.71E-02 |
| OCCIPITAL | Width | 0.75 | 0.11 | 1.54E-07 | 2.13E-02 | 0.19 | 0.08 | 1.37E-02 |
| OCCIPITAL | SurfaceArea | 0.94 | 0.12 | 2.00E-10 | 3.25E-01 | 0.11 | 0.08 | 1.61E-01 |
| S.C. | Length | 1.00 | - | 3.61E-01 | - | 0.04 | 0.07 | 5.12E-01 |
| S.C. | MeanDepth | 0.95 | 0.20 | 9.83E-05 | 4.12E-01 | 0.16 | 0.08 | 2.58E-02 |
| S.C. | Width | 0.98 | 0.03 | 9.30E-23 | 2.46E-01 | 0.43 | 0.07 | 1.85E-11 |
| S.C. | SurfaceArea | 1.00 | - | 2.18E-06 | - | 0.13 | 0.06 | 3.53E-02 |
| S.C.LPC. | Length | - | - | - | - | - | 0.07 | - |
| S.C.LPC. | MeanDepth | -0.13 | - | 1.00E+00 | - | -0.11 | - | 3.75E-01 |
| S.C.LPC. | Width | -0.08 | - | 9.95E-01 | - | 0.00 | - | 9.77E-01 |
| S.C.LPC. | SurfaceArea | 0.91 | - | 1.00E+00 | 1 | -0.90 | - | 2.55E-02 |
| S.C.sylvian. | Length | 1.00 | - | 1.26E-01 | - | -0.02 | 0.10 | 8.43E-01 |
| S.C.sylvian. | MeanDepth | 0.06 | 0.43 | 8.91E-01 | 9.85E-02 | 0.29 | 0.14 | 4.39E-02 |
| S.C.sylvian. | Width | 0.59 | 0.52 | 2.07E-01 | 2.96E-01 | 0.08 | 0.14 | 5.57E-01 |
| S.C.sylvian. | SurfaceArea | 1.00 | - | 5.42E-02 | - | -0.06 | 0.11 | 5.89E-01 |
| S.Call. | Length | 0.76 | 0.13 | 1.48E-08 | 4.96E-02 | -0.04 | 0.08 | 5.58E-01 |
| S.Call. | MeanDepth | 0.52 | 0.20 | 7.81E-03 | 5.30E-02 | 0.10 | 0.08 | 1.83E-01 |
| S.Call. | Width | 0.58 | 0.31 | 7.89E-02 | 1.80E-01 | 0.17 | 0.07 | 1.78E-02 |
| S.Call. | SurfaceArea | 0.70 | 0.10 | 1.21E-09 | 5.58E-03 | -0.03 | 0.08 | 6.79E-01 |
| S.Cu. | Length | 1.00 | - | 1.87E-04 | - | -0.11 | 0.06 | 7.06E-02 |
| S.Cu. | MeanDepth | 0.41 | 0.29 | 1.46E-01 | 5.66E-02 | -0.02 | 0.08 | 7.66E-01 |
| S.Cu. | Width | 1.00 | - | 9.04E-03 | - | 0.02 | 0.07 | 7.58E-01 |
| S.Cu. | SurfaceArea | 1.00 | - | 5.95E-02 | - | -0.07 | 0.06 | 2.77E-01 |
| S.F.inf. | Length | -1.00 | - | 3.38E-01 | - | 0.11 | 0.06 | 5.11E-02 |
| S.F.inf. | MeanDepth | 0.82 | 0.11 | 1.51E-07 | 6.61E-02 | 0.11 | 0.08 | 1.97E-01 |
| S.F.inf. | Width | 0.86 | 0.14 | 3.36E-06 | 1.59E-01 | 0.22 | 0.07 | 1.62E-03 |
| S.F.inf. | SurfaceArea | 0.81 | 0.32 | 9.60E-03 | 3.06E-01 | 0.13 | 0.07 | 7.05E-02 |
| S.F.inf.ant. | Length | 1.00 | - | 1.69E-01 | - | 0.02 | 0.06 | 7.73E-01 |
| S.F.inf.ant. | MeanDepth | - | 0.11 | - | - | - | - | - |
| S.F.inf.ant. | Width | 0.58 | 0.31 | 8.08E-02 | 1.27E-01 | 0.16 | 0.07 | 2.83E-02 |
| S.F.inf.ant. | SurfaceArea | 0.87 | 0.63 | 9.54E-02 | 4.22E-01 | -0.02 | 0.07 | 7.97E-01 |
| S.F.int. | Length | 0.78 | 0.28 | 3.62E-03 | 2.51E-01 | 0.08 | 0.08 | 2.80E-01 |
| S.F.int. | MeanDepth | 0.51 | 0.25 | 1.83E-02 | 5.96E-02 | -0.14 | 0.08 | 7.29E-02 |
| S.F.int. | Width | 0.75 | 0.24 | 3.56E-04 | 1.81E-01 | -0.02 | 0.08 | 8.40E-01 |
| S.F.int. | SurfaceArea | 0.60 | 0.25 | 1.75E-02 | 9.43E-02 | 0.07 | 0.08 | 3.74E-01 |
| S.F.inter. | Length | 0.71 | 0.54 | 1.46E-01 | 3.24E-01 | 0.08 | 0.07 | 2.26E-01 |
| S.F.inter. | MeanDepth | 1.00 | - | 1.39E-03 | - | 0.14 | 0.06 | 1.41E-02 |
| S.F.inter. | Width | 0.69 | 0.11 | 8.30E-07 | 4.16E-03 | 0.23 | 0.07 | 2.54E-03 |
| S.F.inter. | SurfaceArea | 1.00 | - | 1.06E-02 | - | 0.11 | 0.06 | 4.83E-02 |
| S.F.marginal. | Length | -0.31 | 0.32 | 2.73E-01 | - | 0.28 | 0.07 | 2.02E-04 |
| S.F.marginal. | MeanDepth | 1.00 | - | 8.89E-01 | - | 0.12 | 0.05 | 2.65E-02 |
| S.F.marginal. | Width | 1.00 | - | 5.49E-01 | - | 0.15 | 0.06 | 1.21E-02 |
| S.F.marginal. | SurfaceArea | 0.02 | 0.50 | 9.69E-01 | 1.22E-01 | 0.24 | 0.07 | 7.58E-04 |
| S.F.median. | Length | 0.89 | 0.40 | 3.14E-02 | 3.99E-01 | 0.07 | 0.08 | 3.23E-01 |
| S.F.median. | MeanDepth | 0.42 | 0.37 | 2.85E-01 | 1.20E-01 | 0.12 | 0.07 | 1.06E-01 |
| S.F.median. | Width | 1.00 | - | 1.68E-05 | - | 0.13 | 0.06 | 3.63E-02 |
| S.F.median. | SurfaceArea | 0.75 | 0.34 | 6.44E-02 | 2.53E-01 | 0.14 | 0.07 | 5.35E-02 |
| S.F.orbitaire. | Length | 0.90 | 5.48 | 8.43E-01 | 1.00E+00 | 0.07 | 0.07 | 2.57E-01 |
| S.F.orbitaire. | MeanDepth | 1.00 | - | 2.48E-01 | - | 0.05 | 0.07 | 4.40E-01 |
| S.F.orbitaire. | Width | 0.16 | 0.42 | 7.12E-01 | 9.44E-02 | 0.15 | 0.07 | 5.16E-02 |
| S.F.orbitaire. | SurfaceArea | 0.75 | 1.19 | 4.54E-01 | 4.32E-01 | 0.09 | 0.07 | 1.91E-01 |
| S.F.polaire.tr. | Length | 1.00 | - | 7.63E-05 | - | 0.03 | 0.07 | 6.71E-01 |
| S.F.polaire.tr. | MeanDepth | 0.51 | 0.27 | 6.94E-02 | 7.73E-02 | 0.11 | 0.07 | 1.33E-01 |

| | | | | | | | | |
|---|---|---|---|---|---|---|---|---|
| S.F.polaire.tr. | Width | 0.52 | 0.16 | 3.82E-03 | 4.33E-03 | 0.18 | 0.07 | 1.36E-02 |
| S.F.polaire.tr. | SurfaceArea | 0.95 | 0.27 | 3.04E-04 | 4.35E-01 | 0.09 | 0.08 | 1.94E-01 |
| S.F.sup. | Length | 0.12 | 1037.39 | 1.00E+00 | 1.00E+00 | 0.10 | 0.04 | 3.03E-02 |
| S.F.sup. | MeanDepth | 0.79 | 0.12 | 2.33E-05 | 4.85E-02 | 0.23 | 0.07 | 2.39E-03 |
| S.F.sup. | Width | 0.88 | 0.09 | 1.41E-10 | 8.97E-02 | 0.17 | 0.08 | 2.57E-02 |
| S.F.sup. | SurfaceArea | 1.00 | - | 9.03E-02 | - | 0.12 | 0.06 | 4.93E-02 |
| S.Li.ant. | Length | 1.00 | - | 2.58E-01 | - | 0.05 | 0.07 | 4.81E-01 |
| S.Li.ant. | MeanDepth | 1.00 | - | 8.25E-04 | - | 0.06 | 0.09 | 5.44E-01 |
| S.Li.ant. | Width | 0.90 | 0.23 | 3.08E-04 | 3.45E-01 | 0.10 | 0.09 | 3.04E-01 |
| S.Li.ant. | SurfaceArea | 1.00 | - | 5.64E-02 | - | 0.08 | 0.09 | 3.33E-01 |
| S.Li.post. | Length | 0.74 | 0.40 | 3.81E-02 | 2.79E-01 | 0.00 | 0.07 | 9.98E-01 |
| S.Li.post. | MeanDepth | 1.00 | - | 5.47E-02 | - | 0.08 | 0.06 | 2.18E-01 |
| S.Li.post. | Width | 1.00 | - | 4.19E-01 | - | 0.15 | 0.06 | 1.60E-02 |
| S.Li.post. | SurfaceArea | 1.00 | - | 8.34E-03 | - | -0.04 | 0.06 | 5.14E-01 |
| S.O.p. | Length | 0.77 | 0.67 | 1.80E-01 | 3.82E-01 | -0.03 | 0.08 | 7.08E-01 |
| S.O.p. | MeanDepth | 1.00 | - | 2.84E-02 | - | 0.01 | 0.07 | 9.24E-01 |
| S.O.p. | Width | 1.00 | - | 5.45E-03 | - | 0.09 | 0.07 | 1.89E-01 |
| S.O.p. | SurfaceArea | 0.78 | 0.54 | 1.24E-01 | 3.55E-01 | -0.02 | 0.09 | 8.05E-01 |
| S.O.T.lat.ant. | Length | 0.94 | 0.45 | 4.53E-02 | 4.47E-01 | 0.17 | 0.07 | 5.00E-03 |
| S.O.T.lat.ant. | MeanDepth | 0.83 | 0.31 | 9.85E-02 | 3.13E-01 | 0.45 | 0.06 | 1.41E-14 |
| S.O.T.lat.ant. | Width | 0.28 | 0.17 | 1.59E-01 | 2.76E-04 | 0.38 | 0.06 | 1.37E-07 |
| S.O.T.lat.ant. | SurfaceArea | 0.69 | 0.23 | 4.53E-02 | 1.25E-01 | 0.40 | 0.06 | 4.98E-10 |
| S.O.T.lat.int. | Length | 1.00 | - | 1.22E-01 | - | -0.06 | 0.07 | 3.87E-01 |
| S.O.T.lat.int. | MeanDepth | -0.01 | - | 1.00E+00 | - | 0.03 | - | 5.26E-01 |
| S.O.T.lat.int. | Width | 1.00 | - | 7.70E-01 | - | 0.10 | 0.08 | 1.98E-01 |
| S.O.T.lat.int. | SurfaceArea | 1.00 | - | 3.91E-01 | - | -0.03 | 0.07 | 6.57E-01 |
| S.O.T.lat.med. | Length | 1.00 | - | 8.20E-01 | - | 0.03 | 0.06 | 6.30E-01 |
| S.O.T.lat.med. | MeanDepth | -0.87 | 1.30 | 3.47E-01 | - | 0.15 | 0.08 | 6.89E-02 |
| S.O.T.lat.med. | Width | 0.42 | 0.35 | 2.63E-01 | 8.51E-02 | 0.09 | 0.08 | 2.92E-01 |
| S.O.T.lat.med. | SurfaceArea | 0.20 | 2.04 | 9.19E-01 | 4.15E-01 | 0.04 | 0.08 | 5.91E-01 |
| S.O.T.lat.post. | Length | 1.00 | - | 4.33E-01 | - | 0.05 | 0.06 | 3.36E-01 |
| S.O.T.lat.post. | MeanDepth | 0.84 | 0.29 | 1.84E-03 | 2.99E-01 | 0.00 | 0.08 | 9.65E-01 |
| S.O.T.lat.post. | Width | 0.42 | 0.34 | 1.87E-01 | 2.16E-01 | 0.23 | 0.08 | 3.86E-03 |
| S.O.T.lat.post. | SurfaceArea | 1.00 | - | 1.38E-02 | - | 0.02 | 0.06 | 6.79E-01 |
| S.Olf. | Length | 0.63 | 0.20 | 7.85E-03 | 4.40E-02 | 0.25 | 0.07 | 5.54E-04 |
| S.Olf. | MeanDepth | 0.77 | 0.15 | 4.07E-05 | 7.74E-02 | 0.16 | 0.08 | 3.68E-02 |
| S.Olf. | Width | 0.42 | 0.21 | 7.04E-02 | 1.80E-02 | 0.19 | 0.07 | 8.94E-03 |
| S.Olf. | SurfaceArea | 0.76 | 0.15 | 3.92E-04 | 6.94E-02 | 0.29 | 0.07 | 4.25E-05 |
| S.Or. | Length | 1.00 | - | 4.69E-02 | - | 0.19 | 0.06 | 2.05E-03 |
| S.Or. | MeanDepth | 1.00 | - | 1.70E-05 | - | 0.09 | 0.07 | 1.56E-01 |
| S.Or. | Width | 1.00 | - | 8.96E-05 | - | 0.25 | 0.07 | 1.84E-04 |
| S.Or. | SurfaceArea | 0.96 | 0.27 | 1.26E-03 | 4.38E-01 | 0.25 | 0.08 | 4.74E-04 |
| S.p.C. | Length | 0.82 | 1.64 | 4.79E-01 | 4.62E-01 | -0.07 | 0.08 | 2.86E-01 |
| S.p.C. | MeanDepth | -1.00 | - | 5.32E-01 | - | 0.09 | 0.06 | 1.43E-01 |
| S.p.C. | Width | -0.16 | 1.65 | 9.21E-01 | - | 0.05 | 0.08 | 5.47E-01 |
| S.p.C. | SurfaceArea | 0.58 | 1.34 | 6.21E-01 | 4.00E-01 | -0.03 | 0.08 | 7.24E-01 |
| S.Pa.int. | Length | 1.00 | - | 7.35E-02 | - | -0.02 | 0.06 | 7.38E-01 |
| S.Pa.int. | MeanDepth | 1.00 | - | 9.26E-04 | - | 0.01 | 0.06 | 8.07E-01 |
| S.Pa.int. | Width | 1.00 | - | 2.41E-07 | - | 0.00 | 0.07 | 9.94E-01 |
| S.Pa.int. | SurfaceArea | 1.00 | - | 1.98E-03 | - | -0.06 | 0.06 | 2.98E-01 |
| S.Pa.sup. | Length | 1.00 | - | 5.65E-01 | - | 0.06 | 0.07 | 3.78E-01 |
| S.Pa.sup. | MeanDepth | -0.89 | 1.48 | 3.45E-01 | - | 0.13 | 0.09 | 1.47E-01 |
| S.Pa.sup. | Width | 1.00 | - | 8.45E-02 | - | 0.20 | 0.07 | 6.32E-03 |
| S.Pa.sup. | SurfaceArea | 0.11 | - | 1.00E+00 | 4.87E-01 | 0.09 | - | 2.00E-01 |
| S.Pa.t. | Length | 1.00 | - | 8.52E-01 | - | 0.04 | 0.09 | 6.88E-01 |
| S.Pa.t. | MeanDepth | -1.00 | - | 2.21E-01 | - | 0.53 | 0.11 | 6.55E-05 |
| S.Pa.t. | Width | 0.08 | - | 1.00E+00 | 5.00E-01 | 0.25 | - | 7.39E-02 |
| S.Pa.t. | SurfaceArea | 0.20 | 2.44 | 9.25E-01 | 4.69E-01 | 0.14 | 0.16 | 3.67E-01 |
| S.Pe.C.inf. | Length | 1.00 | - | 2.60E-01 | - | 0.10 | 0.07 | 1.14E-01 |
| S.Pe.C.inf. | MeanDepth | 0.86 | 0.85 | 2.65E-01 | 4.43E-01 | 0.11 | 0.09 | 1.62E-01 |
| S.Pe.C.inf. | Width | 1.00 | - | 2.84E-07 | - | 0.07 | 0.07 | 3.20E-01 |
| S.Pe.C.inf. | SurfaceArea | 0.48 | 0.51 | 3.57E-01 | 2.44E-01 | 0.11 | 0.08 | 1.81E-01 |
| S.Pe.C.inter. | Length | 0.13 | 0.51 | 7.86E-01 | 0.1185144 | 0.02 | 0.07 | 8.06E-01 |
| S.Pe.C.inter. | MeanDepth | 1.00 | - | 1.36E-05 | - | -0.09 | 0.07 | 1.94E-01 |

| | | | | | | | | |
|---|---|---|---|---|---|---|---|---|
| S.Pe.C.inter. | Width | 1.00 | - | 6.35E-07 | - | 0.09 | 0.06 | 1.57E-01 |
| S.Pe.C.inter. | SurfaceArea | -0.07 | 0.29 | 8.06E-01 | - | 0.14 | 0.07 | 5.85E-02 |
| S.Pe.C.marginal. | Length | -1.00 | - | 4.94E-01 | - | 0.02 | 0.09 | 8.26E-01 |
| S.Pe.C.marginal. | MeanDepth | 1.00 | - | 2.14E-01 | - | 0.02 | 0.08 | 8.37E-01 |
| S.Pe.C.marginal. | Width | 1.00 | - | 5.81E-02 | - | 0.20 | 0.08 | 1.79E-02 |
| S.Pe.C.marginal. | SurfaceArea | -0.76 | 1.98 | 4.89E-01 | - | 0.07 | 0.10 | 5.05E-01 |
| S.Pe.C.median. | Length | 1.00 | - | 4.88E-01 | - | 0.11 | 0.07 | 1.35E-01 |
| S.Pe.C.median. | MeanDepth | 1.00 | - | 2.69E-01 | - | 0.15 | 0.07 | 3.23E-02 |
| S.Pe.C.median. | Width | 1.00 | - | 6.41E-02 | - | 0.15 | 0.08 | 5.19E-02 |
| S.Pe.C.median. | SurfaceArea | 1.00 | - | 4.57E-01 | - | 0.12 | 0.07 | 7.75E-02 |
| S.Pe.C.sup. | Length | 1.00 | - | 2.22E-01 | - | -0.01 | 0.07 | 9.34E-01 |
| S.Pe.C.sup. | MeanDepth | 1.00 | - | 1.52E-02 | - | -0.06 | 0.07 | 4.44E-01 |
| S.Pe.C.sup. | Width | 1.00 | - | 7.97E-05 | - | 0.23 | 0.07 | 6.10E-04 |
| S.Pe.C.sup. | SurfaceArea | 1.00 | - | 2.97E-02 | - | -0.06 | 0.07 | 3.45E-01 |
| S.Po.C.sup. | Length | 1.00 | - | 2.63E-01 | - | 0.05 | 0.06 | 3.52E-01 |
| S.Po.C.sup. | MeanDepth | 1.00 | - | 2.87E-02 | - | 0.00 | 0.06 | 9.95E-01 |
| S.Po.C.sup. | Width | 1.00 | - | 1.11E-10 | - | 0.07 | 0.07 | 3.63E-01 |
| S.Po.C.sup. | SurfaceArea | 1.00 | - | 8.24E-02 | - | 0.04 | 0.06 | 5.04E-01 |
| S.R.inf. | Length | 1.00 | - | 7.70E-04 | - | -0.10 | 0.07 | 1.51E-01 |
| S.R.inf. | MeanDepth | 1.00 | - | 6.45E-02 | - | -0.08 | 0.07 | 2.49E-01 |
| S.R.inf. | Width | 1.00 | - | 1.36E-01 | - | -0.03 | 0.06 | 6.73E-01 |
| S.R.inf. | SurfaceArea | 1.00 | - | 3.12E-04 | - | -0.14 | 0.07 | 4.20E-02 |
| S.Rh. | Length | 0.54 | 0.68 | 4.18E-01 | 2.92E-01 | 0.16 | 0.08 | 2.61E-02 |
| S.Rh. | MeanDepth | 0.72 | 0.59 | 3.09E-01 | 3.36E-01 | 0.25 | 0.07 | 2.23E-04 |
| S.Rh. | Width | 1.00 | - | 3.20E-02 | - | 0.22 | 0.06 | 7.73E-04 |
| S.Rh. | SurfaceArea | -0.03 | 0.65 | 9.65E-01 | - | 0.29 | 0.07 | 5.57E-05 |
| S.s.P. | Length | 0.66 | 0.26 | 1.06E-02 | 1.29E-01 | -0.02 | 0.08 | 8.26E-01 |
| S.s.P. | MeanDepth | 1.00 | - | 2.41E-08 | - | 0.04 | 0.06 | 5.01E-01 |
| S.s.P. | Width | 1.00 | - | 3.97E-06 | - | 0.15 | 0.06 | 2.25E-02 |
| S.s.P. | SurfaceArea | 0.83 | 0.20 | 5.60E-04 | 2.09E-01 | 0.07 | 0.08 | 3.76E-01 |
| S.T.i.ant. | Length | 1.00 | - | 3.36E-05 | - | -0.03 | 0.06 | 5.81E-01 |
| S.T.i.ant. | MeanDepth | 0.86 | 0.29 | 4.08E-04 | 3.25E-01 | -0.06 | 0.07 | 3.82E-01 |
| S.T.i.ant. | Width | 0.92 | 0.15 | 1.50E-10 | 2.93E-01 | -0.04 | 0.07 | 5.92E-01 |
| S.T.i.ant. | SurfaceArea | 0.91 | 0.20 | 1.23E-06 | 3.33E-01 | -0.04 | 0.07 | 5.76E-01 |
| S.T.i.post. | Length | 1.00 | - | 3.89E-01 | - | 0.02 | 0.06 | 7.89E-01 |
| S.T.i.post. | MeanDepth | 0.67 | 0.31 | 1.33E-02 | 1.81E-01 | 0.01 | 0.08 | 9.37E-01 |
| S.T.i.post. | Width | 0.64 | 0.14 | 1.84E-04 | 1.60E-02 | 0.17 | 0.07 | 1.96E-02 |
| S.T.i.post. | SurfaceArea | 0.99 | 0.87 | 1.58E-01 | 4.95E-01 | 0.02 | 0.08 | 7.21E-01 |
| S.T.pol. | Length | 0.73 | 0.51 | 1.24E-01 | 3.26E-01 | 0.05 | 0.08 | 4.73E-01 |
| S.T.pol. | MeanDepth | 0.90 | 6.89 | 8.07E-01 | 1.00E+00 | 0.26 | 0.07 | 1.53E-04 |
| S.T.pol. | Width | 1.00 | - | 2.01E-03 | - | 0.22 | 0.06 | 7.69E-04 |
| S.T.pol. | SurfaceArea | 0.61 | 0.36 | 1.26E-01 | 1.80E-01 | 0.14 | 0.08 | 7.45E-02 |
| S.T.s. | Length | 1.00 | - | 8.34E-01 | - | 0.00 | 0.06 | 9.60E-01 |
| S.T.s. | MeanDepth | 0.67 | 0.19 | 2.00E-03 | 6.73E-02 | 0.09 | 0.08 | 2.25E-01 |
| S.T.s. | Width | 0.68 | 0.14 | 2.90E-04 | 2.66E-02 | 0.23 | 0.07 | 1.18E-03 |
| S.T.s. | SurfaceArea | 0.34 | 0.26 | 1.86E-01 | 2.61E-02 | 0.06 | 0.07 | 4.59E-01 |
| S.T.s.ter.asc.ant. | Length | 1.00 | - | 3.22E-01 | - | -0.02 | 0.06 | 7.01E-01 |
| S.T.s.ter.asc.ant. | MeanDepth | 0.74 | 1.92 | 6.51E-01 | 0.4540901 | 0.01 | 0.08 | 8.99E-01 |
| S.T.s.ter.asc.ant. | Width | 0.88 | 0.31 | 1.18E-03 | 3.63E-01 | 0.03 | 0.09 | 7.35E-01 |
| S.T.s.ter.asc.ant. | SurfaceArea | 0.02 | - | 1.00E+00 | 1 | 0.02 | - | 5.41E-01 |
| S.T.s.ter.asc.post. | Length | 1.00 | - | 7.97E-01 | - | 0.05 | 0.07 | 4.01E-01 |
| S.T.s.ter.asc.post. | MeanDepth | 0.64 | 0.30 | 3.89E-02 | 1.48E-01 | -0.02 | 0.08 | 7.51E-01 |
| S.T.s.ter.asc.post. | Width | 0.60 | 0.40 | 1.18E-01 | 2.65E-01 | 0.19 | 0.07 | 1.13E-02 |
| S.T.s.ter.asc.post. | SurfaceArea | 0.72 | 0.90 | 3.34E-01 | 3.98E-01 | 0.01 | 0.07 | 9.21E-01 |
| Globals | Length | 1.00 | - | 2.18E-09 | - | 0.84 | 0.02 | 3.75E-112 |
| Globals | MeanDepth | 1.00 | - | 9.69E-09 | - | 0.84 | 0.02 | 3.41E-123 |
| Globals | Width | 0.93 | 0.04 | 1.78E-09 | 6.27E-02 | 0.68 | 0.04 | 1.55E-48 |
| Globals | SurfaceArea | 1.00 | - | 1.06E-35 | - | 0.89 | 0.01 | 5.71E-166 |

**Table S23:** Bivariate analysis: left-right genetic correlation in HCP (yellow: Bonferroni corrected; grey: $p < 0.05$).

| Sulci | Descriptor | rhoG | rhoG_SE | rhoP=0 | rhoP=1 | rhoE | rhoE.se | rhoE.p |
|---|---|---|---|---|---|---|---|---|
| F.C.L.a. | Length | 1.00 | - | 2.50E-01 | - | 0.07 | 0.07 | 2.89E-01 |
| F.C.L.a. | MeanDepth | 1.00 | - | 1.12E-03 | - | -0.03 | 0.06 | 6.78E-01 |
| F.C.L.a. | Width | 0.99 | 0.16 | 4.06E-05 | 4.75E-01 | 0.25 | 0.08 | 1.08E-03 |
| F.C.L.a. | SurfaceArea | 1.00 | - | 3.96E-05 | - | 0.08 | 0.07 | 2.47E-01 |
| F.C.L.p. | Length | 1.00 | - | 1.07E-01 | - | 0.00 | 0.06 | 9.53E-01 |
| F.C.L.p. | MeanDepth | 0.94 | 0.33 | 9.12E-03 | 4.31E-01 | 0.07 | 0.08 | 3.64E-01 |
| F.C.L.p. | Width | 0.88 | 0.07 | 4.67E-10 | 4.62E-02 | 0.26 | 0.09 | 3.57E-03 |
| F.C.L.p. | SurfaceArea | 0.95 | 0.14 | 2.49E-11 | 3.52E-01 | -0.06 | 0.09 | 5.21E-01 |
| F.C.L.r.ant. | Length | 1.00 | - | 1.64E-01 | - | 0.01 | 0.07 | 9.26E-01 |
| F.C.L.r.ant. | MeanDepth | 0.42 | 0.61 | 5.12E-01 | 2.67E-01 | 0.09 | 0.09 | 3.29E-01 |
| F.C.L.r.ant. | Width | -0.27 | 1.99 | 8.71E-01 | - | 0.27 | 0.08 | 7.71E-04 |
| F.C.L.r.ant. | SurfaceArea | 0.80 | 0.55 | 1.20E-01 | 3.81E-01 | 0.09 | 0.09 | 3.16E-01 |
| F.C.L.r.asc. | Length | 1.00 | - | 6.04E-01 | - | 0.08 | 0.06 | 1.82E-01 |
| F.C.L.r.asc. | MeanDepth | 1.00 | - | 9.36E-03 | - | 0.06 | 0.07 | 3.86E-01 |
| F.C.L.r.asc. | Width | 1.00 | - | 6.37E-03 | - | 0.06 | 0.07 | 4.06E-01 |
| F.C.L.r.asc. | SurfaceArea | 0.64 | 0.33 | 5.14E-02 | 1.68E-01 | 0.06 | 0.08 | 4.44E-01 |
| F.C.L.r.diag. | Length | 0.03 | - | 1.00E+00 | 1 | 0.19 | 0.08 | 1.88E-01 |
| F.C.L.r.diag. | MeanDepth | 1.00 | - | 4.81E-01 | - | 0.11 | 0.15 | 4.82E-01 |
| F.C.L.r.diag. | Width | 1.00 | - | 7.14E-02 | - | -0.15 | 0.15 | 3.15E-01 |
| F.C.L.r.diag. | SurfaceArea | 0.10 | - | 1.00E+00 | 1.00E+00 | 0.14 | - | 1.72E-01 |
| F.C.L.r.retroC.tr. | Length | 0.98 | 1.19 | 2.24E-01 | 4.93E-01 | -0.08 | 0.08 | 3.12E-01 |
| F.C.L.r.retroC.tr. | MeanDepth | 0.54 | 0.25 | 4.55E-02 | 5.83E-02 | 0.01 | 0.11 | 9.46E-01 |
| F.C.L.r.retroC.tr. | Width | 0.29 | 0.50 | 5.52E-01 | 2.49E-01 | 0.06 | 0.10 | 5.01E-01 |
| F.C.L.r.retroC.tr. | SurfaceArea | 1.00 | - | 7.62E-02 | - | -0.02 | 0.08 | 7.52E-01 |
| F.C.L.r.sc.ant. | Length | - | 1.19 | - | - | - | 0.08 | - |
| F.C.L.r.sc.ant. | MeanDepth | - | 0.25 | - | - | - | - | - |
| F.C.L.r.sc.ant. | Width | - | 0.50 | - | - | - | - | - |
| F.C.L.r.sc.ant. | SurfaceArea | - | - | - | - | - | - | - |
| F.C.L.r.sc.post. | Length | - | 1.19 | - | - | - | 0.08 | - |
| F.C.L.r.sc.post. | MeanDepth | 0.76 | - | 4.43E-01 | 1.00E+00 | 0.05 | - | 1.00E+00 |
| F.C.L.r.sc.post. | Width | - | 0.50 | - | - | - | - | - |
| F.C.L.r.sc.post. | SurfaceArea | - | - | - | - | - | - | - |
| F.C.M.ant. | Length | 1.00 | - | 5.30E-01 | - | 0.04 | 0.06 | 4.88E-01 |
| F.C.M.ant. | MeanDepth | 0.58 | 0.48 | 1.35E-01 | 2.65E-01 | -0.09 | 0.07 | 1.91E-01 |
| F.C.M.ant. | Width | 0.77 | 0.23 | 1.37E-03 | 2.04E-01 | 0.09 | 0.07 | 2.48E-01 |
| F.C.M.ant. | SurfaceArea | 1.00 | - | 1.42E-01 | - | -0.04 | 0.06 | 5.04E-01 |
| F.C.M.post. | Length | 1.00 | - | 2.35E-01 | - | 0.02 | 0.06 | 7.46E-01 |
| F.C.M.post. | MeanDepth | 0.95 | 0.12 | 1.87E-08 | 3.53E-01 | 0.05 | 0.08 | 5.66E-01 |
| F.C.M.post. | Width | 0.94 | 0.07 | 8.02E-14 | 2.11E-01 | 0.30 | 0.08 | 7.86E-05 |
| F.C.M.post. | SurfaceArea | 0.88 | 0.22 | 3.10E-04 | 3.01E-01 | 0.04 | 0.07 | 6.20E-01 |
| F.Cal.ant.-Sc.Cal. | Length | 1.00 | - | 6.26E-02 | - | 0.07 | 0.06 | 2.89E-01 |
| F.Cal.ant.-Sc.Cal. | MeanDepth | 0.71 | 0.10 | 9.52E-08 | 0.0041053 | 0.16 | 0.08 | 7.11E-02 |
| F.Cal.ant.-Sc.Cal. | Width | 0.94 | 0.12 | 5.39E-08 | 0.3067672 | 0.18 | 0.08 | 3.12E-02 |
| F.Cal.ant.-Sc.Cal. | SurfaceArea | 1.00 | - | 5.91E-13 | - | 0.09 | 0.08 | 2.74E-01 |
| F.Coll. | Length | 0.25 | 0.66 | 7.16E-01 | 2.43E-01 | 0.13 | 0.07 | 6.09E-02 |
| F.Coll. | MeanDepth | 0.80 | 0.15 | 9.28E-06 | 1.06E-01 | 0.18 | 0.07 | 1.59E-02 |
| F.Coll. | Width | 0.78 | 0.11 | 3.17E-09 | 3.02E-02 | 0.06 | 0.08 | 4.14E-01 |
| F.Coll. | SurfaceArea | 0.55 | 0.23 | 5.81E-02 | 3.47E-02 | 0.26 | 0.07 | 6.72E-04 |
| F.I.P. | Length | 1.00 | - | 3.40E-04 | - | 0.09 | 0.06 | 1.81E-01 |
| F.I.P. | MeanDepth | 0.75 | 0.16 | 5.63E-06 | 7.91E-02 | 0.06 | 0.08 | 4.37E-01 |
| F.I.P. | Width | 0.97 | 0.03 | 9.38E-16 | 1.77E-01 | 0.48 | 0.07 | 6.47E-11 |
| F.I.P. | SurfaceArea | 1.00 | - | 2.05E-05 | - | 0.15 | 0.06 | 2.18E-02 |
| F.I.P.Po.C.inf. | Length | 0.83 | 0.52 | 5.46E-02 | 3.81E-01 | -0.14 | 0.07 | 5.35E-02 |
| F.I.P.Po.C.inf. | MeanDepth | 0.96 | 0.40 | 6.35E-03 | 4.65E-01 | 0.09 | 0.07 | 1.58E-01 |
| F.I.P.Po.C.inf. | Width | 0.89 | 0.08 | 7.95E-09 | 7.52E-02 | 0.37 | 0.07 | 2.86E-06 |
| F.I.P.Po.C.inf. | SurfaceArea | 0.96 | 0.34 | 1.39E-03 | 4.54E-01 | -0.11 | 0.08 | 1.41E-01 |
| F.I.P.r.int.1 | Length | 0.61 | 0.58 | 1.96E-01 | 0.3137874 | -0.04 | 0.07 | 6.22E-01 |
| F.I.P.r.int.1 | MeanDepth | 1.00 | - | 5.62E-01 | - | 0.03 | 0.07 | 6.17E-01 |
| F.I.P.r.int.1 | Width | 0.42 | 0.42 | 3.08E-01 | 0.1573733 | 0.07 | 0.08 | 3.82E-01 |
| F.I.P.r.int.1 | SurfaceArea | 0.60 | 0.95 | 3.85E-01 | 0.3969436 | -0.01 | 0.08 | 9.30E-01 |
| F.I.P.r.int.2 | Length | 0.02 | - | 1.00E+00 | 1 | 0.04 | - | 1.00E+00 |
| F.I.P.r.int.2 | MeanDepth | 0.18 | - | 1.00E+00 | 1 | 0.06 | - | 2.55E-01 |

| | | | | | | | | |
|---|---|---|---|---|---|---|---|---|
| F.I.P.r.int.2 | Width | 0.27 | 0.93 | 7.75E-01 | 0.3802749 | 0.09 | 0.11 | 4.11E-01 |
| F.I.P.r.int.2 | SurfaceArea | 0.09 | - | 1.00E+00 | 1 | 0.10 | - | 1.00E+00 |
| F.P.O. | Length | 0.27 | 0.60 | 6.34E-01 | 2.18E-01 | 0.05 | 0.07 | 4.76E-01 |
| F.P.O. | MeanDepth | 0.85 | 0.13 | 2.43E-08 | 1.44E-01 | 0.06 | 0.08 | 5.00E-01 |
| F.P.O. | Width | 0.86 | 0.10 | 1.01E-08 | 8.71E-02 | 0.28 | 0.07 | 1.98E-04 |
| F.P.O. | SurfaceArea | 1.00 | - | 4.28E-12 | - | -0.05 | 0.07 | 4.61E-01 |
| INSULA | Length | 0.04 | - | 1.00E+00 | 1 | 0.03 | - | 1.00E+00 |
| INSULA | MeanDepth | 1.00 | - | 5.82E-01 | - | 0.86 | 0.10 | 6.79E-05 |
| INSULA | Width | 1.00 | - | 6.26E-01 | - | 0.47 | 0.20 | 8.09E-02 |
| INSULA | SurfaceArea | 0.78 | - | 4.81E-04 | 3.05E-01 | -0.90 | - | 9.20E-01 |
| OCCIPITAL | Length | 0.76 | 0.13 | 1.25E-04 | 4.58E-02 | 0.33 | 0.07 | 1.61E-05 |
| OCCIPITAL | MeanDepth | 0.56 | 0.19 | 3.72E-03 | 2.47E-02 | 0.02 | 0.08 | 8.24E-01 |
| OCCIPITAL | Width | 1.00 | - | 9.79E-13 | - | 0.16 | 0.07 | 3.43E-02 |
| OCCIPITAL | SurfaceArea | 0.80 | 0.10 | 2.74E-07 | 2.17E-02 | 0.30 | 0.07 | 1.61E-04 |
| S.C. | Length | 1.00 | - | 9.50E-02 | - | 0.14 | 0.06 | 2.12E-02 |
| S.C. | MeanDepth | 1.00 | - | 9.29E-08 | - | 0.12 | 0.07 | 1.04E-01 |
| S.C. | Width | 1.00 | - | 2.33E-58 | - | 0.26 | 0.07 | 4.53E-04 |
| S.C. | SurfaceArea | 1.00 | - | 3.43E-08 | - | 0.05 | 0.07 | 5.33E-01 |
| S.C.LPC. | Length | -1.00 | - | 8.31E-01 | - | 0.44 | 1.95 | 1.21E-01 |
| S.C.LPC. | MeanDepth | - | - | - | - | - | - | - |
| S.C.LPC. | Width | 0.19 | 19061.74 | 1.00E+00 | 0.4992021 | 0.22 | 0.22 | 7.90E-01 |
| S.C.LPC. | SurfaceArea | -1.00 | - | 8.39E-01 | - | 0.17 | 0.35 | 4.31E-01 |
| S.C.sylvian. | Length | 0.57 | 0.52 | 2.90E-01 | 2.33E-01 | 0.05 | 0.14 | 7.37E-01 |
| S.C.sylvian. | MeanDepth | 1.00 | - | 1.74E-01 | - | 0.01 | 0.11 | 8.99E-01 |
| S.C.sylvian. | Width | 1.00 | - | 1.20E-02 | - | -0.18 | 0.14 | 1.78E-01 |
| S.C.sylvian. | SurfaceArea | 1.00 | - | 1.19E-01 | - | -0.02 | 0.11 | 8.85E-01 |
| S.Call. | Length | 1.00 | - | 3.83E-06 | - | -0.07 | 0.08 | 4.52E-01 |
| S.Call. | MeanDepth | 0.64 | 0.12 | 7.27E-07 | 2.23E-03 | -0.08 | 0.08 | 3.67E-01 |
| S.Call. | Width | 0.83 | 0.17 | 2.34E-04 | 1.71E-01 | 0.15 | 0.07 | 4.99E-02 |
| S.Call. | SurfaceArea | 0.88 | 0.13 | 7.25E-08 | 1.92E-01 | -0.09 | 0.10 | 3.73E-01 |
| S.Cu. | Length | 0.40 | 0.25 | 1.19E-01 | 0.0234255 | 0.02 | 0.07 | 8.16E-01 |
| S.Cu. | MeanDepth | 1.00 | - | 2.87E-03 | - | 0.02 | 0.06 | 8.12E-01 |
| S.Cu. | Width | 1.00 | - | 1.08E-02 | - | 0.06 | 0.07 | 3.43E-01 |
| S.Cu. | SurfaceArea | 0.90 | 0.49 | 2.12E-02 | 4.26E-01 | -0.07 | 0.07 | 3.36E-01 |
| S.F.inf. | Length | 1.00 | - | 1.08E-01 | - | -0.04 | 0.06 | 4.91E-01 |
| S.F.inf. | MeanDepth | 0.85 | 0.16 | 8.66E-05 | 1.72E-01 | 0.19 | 0.08 | 1.45E-02 |
| S.F.inf. | Width | 0.89 | 0.08 | 8.64E-12 | 9.17E-02 | 0.20 | 0.08 | 1.61E-02 |
| S.F.inf. | SurfaceArea | 1.00 | - | 2.95E-05 | - | 0.01 | 0.07 | 8.23E-01 |
| S.F.inf.ant. | Length | 1.00 | - | 1.12E-01 | - | 0.02 | 0.06 | 7.18E-01 |
| S.F.inf.ant. | MeanDepth | 0.81 | 0.46 | 5.41E-02 | 3.51E-01 | 0.03 | 0.07 | 6.32E-01 |
| S.F.inf.ant. | Width | 0.84 | 0.71 | 1.61E-01 | 4.18E-01 | 0.04 | 0.07 | 5.49E-01 |
| S.F.inf.ant. | SurfaceArea | 1.00 | - | 6.48E-02 | - | -0.01 | 0.07 | 9.33E-01 |
| S.F.int. | Length | 0.91 | 0.22 | 1.99E-06 | 3.40E-01 | -0.09 | 0.08 | 2.46E-01 |
| S.F.int. | MeanDepth | 0.89 | 0.31 | 7.33E-04 | 3.63E-01 | -0.13 | 0.08 | 8.84E-02 |
| S.F.int. | Width | 1.00 | - | 2.75E-06 | - | -0.02 | 0.08 | 8.43E-01 |
| S.F.int. | SurfaceArea | 0.89 | 0.19 | 4.54E-07 | 2.92E-01 | -0.12 | 0.08 | 1.25E-01 |
| S.F.inter. | Length | 1.00 | - | 1.44E-01 | - | 0.11 | 0.05 | 3.85E-02 |
| S.F.inter. | MeanDepth | 1.00 | - | 1.30E-03 | - | 0.16 | 0.06 | 1.05E-02 |
| S.F.inter. | Width | 1.00 | - | 2.07E-18 | - | 0.18 | 0.07 | 1.43E-02 |
| S.F.inter. | SurfaceArea | 1.00 | - | 1.69E-01 | - | 0.18 | 0.05 | 1.21E-03 |
| S.F.marginal. | Length | 1.00 | - | 4.58E-04 | - | -0.02 | 0.06 | 7.56E-01 |
| S.F.marginal. | MeanDepth | 0.10 | - | 1.00E+00 | 1.00E+00 | 0.08 | - | 1.00E+00 |
| S.F.marginal. | Width | 1.00 | - | 9.42E-03 | - | 0.09 | 0.06 | 1.35E-01 |
| S.F.marginal. | SurfaceArea | 1.00 | - | 3.64E-04 | - | -0.03 | 0.06 | 6.19E-01 |
| S.F.median. | Length | 1.00 | - | 3.24E-04 | - | -0.07 | 0.07 | 3.76E-01 |
| S.F.median. | MeanDepth | 0.45 | 0.28 | 1.52E-01 | 5.76E-02 | 0.18 | 0.07 | 1.85E-02 |
| S.F.median. | Width | 0.70 | 0.15 | 7.41E-05 | 3.11E-02 | 0.09 | 0.08 | 2.49E-01 |
| S.F.median. | SurfaceArea | 0.87 | 0.22 | 2.50E-04 | 2.87E-01 | 0.00 | 0.08 | 9.63E-01 |
| S.F.orbitaire. | Length | 1.00 | - | 9.75E-02 | - | 0.06 | 0.06 | 3.63E-01 |
| S.F.orbitaire. | MeanDepth | 0.90 | 2.20 | 4.90E-01 | 4.88E-01 | 0.02 | 0.07 | 7.38E-01 |
| S.F.orbitaire. | Width | 1.00 | - | 5.06E-02 | - | 0.08 | 0.06 | 2.21E-01 |
| S.F.orbitaire. | SurfaceArea | 1.00 | - | 7.84E-02 | - | 0.04 | 0.06 | 4.84E-01 |
| S.F.polaire.tr. | Length | 1.00 | - | 4.69E-04 | - | 0.04 | 0.08 | 6.47E-01 |
| S.F.polaire.tr. | MeanDepth | 1.00 | - | 9.76E-03 | - | 0.07 | 0.06 | 2.40E-01 |

| | | | | | | | | |
|---|---|---|---|---|---|---|---|---|
| S.F.polaire.tr. | Width | 0.93 | 0.17 | 9.30E-07 | 3.40E-01 | 0.03 | 0.09 | 7.13E-01 |
| S.F.polaire.tr. | SurfaceArea | 1.00 | - | 4.77E-04 | - | 0.05 | 0.07 | 4.81E-01 |
| S.F.sup. | Length | -1.00 | - | 3.60E-01 | - | 0.07 | 0.05 | 2.25E-01 |
| S.F.sup. | MeanDepth | 1.00 | - | 2.06E-06 | - | 0.13 | 0.06 | 4.63E-02 |
| S.F.sup. | Width | 1.00 | - | 1.66E-15 | - | 0.22 | 0.07 | 3.80E-03 |
| S.F.sup. | SurfaceArea | 0.34 | 0.25 | 2.20E-01 | 7.99E-03 | 0.13 | 0.07 | 7.85E-02 |
| S.Li.ant. | Length | 1.00 | - | 1.27E-01 | - | -0.01 | 0.06 | 8.43E-01 |
| S.Li.ant. | MeanDepth | 0.43 | 0.23 | 1.12E-01 | 1.49E-02 | 0.14 | 0.10 | 1.68E-01 |
| S.Li.ant. | Width | 0.66 | 0.32 | 6.07E-02 | 1.99E-01 | 0.19 | 0.08 | 2.47E-02 |
| S.Li.ant. | SurfaceArea | 0.52 | 0.42 | 2.11E-01 | 1.80E-01 | 0.10 | 0.09 | 2.58E-01 |
| S.Li.post. | Length | 0.11 | 0.54 | 8.45E-01 | 2.05E-01 | 0.06 | 0.07 | 3.94E-01 |
| S.Li.post. | MeanDepth | 0.44 | 0.30 | 1.41E-01 | 6.08E-02 | 0.01 | 0.08 | 8.89E-01 |
| S.Li.post. | Width | 0.86 | 0.26 | 3.38E-04 | 3.07E-01 | -0.04 | 0.08 | 6.55E-01 |
| S.Li.post. | SurfaceArea | 0.13 | 0.44 | 7.76E-01 | 5.29E-02 | 0.06 | 0.07 | 3.78E-01 |
| S.O.p. | Length | 1.00 | - | 8.08E-01 | - | 0.05 | 0.05 | 3.87E-01 |
| S.O.p. | MeanDepth | 1.00 | - | 2.05E-01 | - | 0.08 | 0.07 | 3.07E-01 |
| S.O.p. | Width | 1.00 | - | 4.66E-02 | - | 0.10 | 0.08 | 2.32E-01 |
| S.O.p. | SurfaceArea | 1.00 | - | 1.63E-01 | - | 0.03 | 0.07 | 6.71E-01 |
| S.O.T.lat.ant. | Length | 0.81 | 0.35 | 3.13E-02 | 3.07E-01 | 0.10 | 0.07 | 1.38E-01 |
| S.O.T.lat.ant. | MeanDepth | 1.00 | - | 2.34E-07 | - | 0.22 | 0.07 | 2.67E-03 |
| S.O.T.lat.ant. | Width | 0.99 | 0.19 | 4.28E-05 | 4.83E-01 | 0.25 | 0.07 | 1.97E-04 |
| S.O.T.lat.ant. | SurfaceArea | 0.94 | 0.15 | 5.59E-05 | 3.59E-01 | 0.26 | 0.07 | 1.58E-04 |
| S.O.T.lat.int. | Length | -0.04 | - | 9.94E-01 | - | 0.13 | - | 3.06E-03 |
| S.O.T.lat.int. | MeanDepth | 0.04 | - | 1.00E+00 | 1.00E+00 | 0.09 | - | 2.47E-01 |
| S.O.T.lat.int. | Width | -1.00 | - | 2.20E-01 | - | 0.24 | 0.06 | 1.63E-05 |
| S.O.T.lat.int. | SurfaceArea | 0.10 | - | 1.00E+00 | 1.00E+00 | 0.11 | - | 7.62E-02 |
| S.O.T.lat.med. | Length | -0.34 | 0.91 | 6.75E-01 | - | 0.10 | 0.08 | 1.79E-01 |
| S.O.T.lat.med. | MeanDepth | 0.57 | 1.20 | 5.64E-01 | 4.07E-01 | 0.01 | 0.08 | 8.71E-01 |
| S.O.T.lat.med. | Width | -0.12 | - | 1.00E+00 | - | 0.07 | - | 1.00E-01 |
| S.O.T.lat.med. | SurfaceArea | 0.75 | 1.00 | 3.37E-01 | 4.15E-01 | -0.04 | 0.08 | 6.27E-01 |
| S.O.T.lat.post. | Length | 0.16 | 24415.84 | 1.00E+00 | 4.84E-01 | 0.07 | 0.04 | 1.06E-01 |
| S.O.T.lat.post. | MeanDepth | 0.76 | 0.39 | 7.21E-02 | 2.89E-01 | 0.09 | 0.08 | 2.52E-01 |
| S.O.T.lat.post. | Width | 0.54 | 0.41 | 2.29E-01 | 2.26E-01 | 0.16 | 0.07 | 3.90E-02 |
| S.O.T.lat.post. | SurfaceArea | 0.32 | 53190503.62 | 1.00E+00 | 1.00E+00 | 0.12 | 0.08 | 5.69E-02 |
| S.Olf. | Length | 0.81 | 0.30 | 3.46E-02 | 2.78E-01 | 0.23 | 0.07 | 1.58E-03 |
| S.Olf. | MeanDepth | 0.99 | 0.07 | 6.59E-15 | 4.47E-01 | 0.16 | 0.09 | 5.38E-02 |
| S.Olf. | Width | 0.88 | 0.18 | 1.95E-04 | 2.56E-01 | 0.28 | 0.07 | 5.65E-05 |
| S.Olf. | SurfaceArea | 1.00 | - | 4.21E-10 | - | 0.31 | 0.07 | 3.31E-05 |
| S.Or. | Length | 0.94 | 0.18 | 4.13E-05 | 3.66E-01 | 0.11 | 0.08 | 1.47E-01 |
| S.Or. | MeanDepth | 0.90 | 0.18 | 1.14E-04 | 3.04E-01 | 0.28 | 0.08 | 5.80E-04 |
| S.Or. | Width | 1.00 | - | 1.82E-05 | - | 0.26 | 0.07 | 1.88E-04 |
| S.Or. | SurfaceArea | 0.82 | 0.09 | 1.28E-08 | 2.61E-02 | 0.22 | 0.08 | 8.82E-03 |
| S.p.C. | Length | - | 0.18 | - | | - | 0.08 | - |
| S.p.C. | MeanDepth | 0.67 | 1.06 | 4.22E-01 | 4.11E-01 | 0.01 | 0.09 | 8.90E-01 |
| S.p.C. | Width | 1.00 | - | 2.74E-02 | - | -0.04 | 0.08 | 6.25E-01 |
| S.p.C. | SurfaceArea | -1.00 | - | 4.06E-01 | - | 0.13 | 0.07 | 7.13E-02 |
| S.Pa.int. | Length | 1.00 | - | 1.11E-02 | - | 0.02 | 0.07 | 7.62E-01 |
| S.Pa.int. | MeanDepth | 0.95 | 0.35 | 3.16E-03 | 4.41E-01 | 0.02 | 0.08 | 8.53E-01 |
| S.Pa.int. | Width | 0.90 | 0.23 | 2.82E-04 | 3.44E-01 | 0.14 | 0.08 | 8.51E-02 |
| S.Pa.int. | SurfaceArea | 1.00 | - | 5.43E-03 | - | 0.01 | 0.07 | 8.77E-01 |
| S.Pa.sup. | Length | 1.00 | - | 1.21E-01 | - | -0.02 | 0.07 | 7.61E-01 |
| S.Pa.sup. | MeanDepth | 1.00 | - | 6.11E-02 | - | 0.00 | 0.07 | 9.84E-01 |
| S.Pa.sup. | Width | 1.00 | - | 2.33E-05 | - | 0.10 | 0.08 | 2.33E-01 |
| S.Pa.sup. | SurfaceArea | 1.00 | - | 1.02E-02 | - | -0.08 | 0.07 | 2.69E-01 |
| S.Pa.t. | Length | 1.00 | - | 3.47E-01 | - | -0.02 | 0.09 | 8.40E-01 |
| S.Pa.t. | MeanDepth | 1.00 | - | 2.27E-01 | - | 0.25 | 0.09 | 1.18E-02 |
| S.Pa.t. | Width | 1.00 | - | 3.89E-01 | - | 0.35 | 0.10 | 6.59E-04 |
| S.Pa.t. | SurfaceArea | 1.00 | - | 1.66E-01 | - | 0.06 | 0.10 | 5.41E-01 |
| S.Pe.C.inf. | Length | 0.79 | 1.26 | 4.95E-01 | 4.41E-01 | 0.05 | 0.07 | 5.14E-01 |
| S.Pe.C.inf. | MeanDepth | 0.69 | 0.49 | 1.39E-01 | 2.90E-01 | 0.05 | 0.09 | 5.84E-01 |
| S.Pe.C.inf. | Width | 1.00 | - | 5.74E-04 | - | 0.14 | 0.07 | 5.71E-02 |
| S.Pe.C.inf. | SurfaceArea | 1.00 | - | 2.86E-01 | - | 0.04 | 0.07 | 5.08E-01 |
| S.Pe.C.inter. | Length | 1.00 | - | 8.81E-02 | - | -0.03 | 0.06 | 6.57E-01 |
| S.Pe.C.inter. | MeanDepth | 1.00 | - | 3.18E-03 | - | -0.01 | 0.06 | 8.20E-01 |

| | | | | | | | | |
|---|---|---|---|---|---|---|---|---|
| S.Pe.C.inter. | Width | 1.00 | - | 1.25E-04 | - | 0.20 | 0.06 | 2.91E-03 |
| S.Pe.C.inter. | SurfaceArea | 1.00 | - | 3.47E-03 | - | -0.03 | 0.06 | 6.49E-01 |
| S.Pe.C.marginal. | Length | 0.90 | 13.06 | 9.11E-01 | 1 | 0.06 | 0.11 | 3.94E-01 |
| S.Pe.C.marginal. | MeanDepth | 1.00 | - | 5.96E-02 | - | -0.02 | 0.08 | 7.96E-01 |
| S.Pe.C.marginal. | Width | 0.79 | 0.29 | 7.11E-03 | 2.50E-01 | 0.17 | 0.09 | 5.36E-02 |
| S.Pe.C.marginal. | SurfaceArea | 1.00 | - | 4.61E-01 | - | 0.05 | 0.08 | 5.52E-01 |
| S.Pe.C.median. | Length | 0.61 | 1.24 | 5.19E-01 | 4.17E-01 | 0.09 | 0.08 | 2.42E-01 |
| S.Pe.C.median. | MeanDepth | 0.10 | - | 1.00E+00 | 4.95E-01 | 0.08 | - | 2.54E-01 |
| S.Pe.C.median. | Width | 1.00 | - | 8.27E-05 | - | 0.05 | 0.08 | 5.30E-01 |
| S.Pe.C.median. | SurfaceArea | 0.60 | 0.72 | 3.80E-01 | 3.34E-01 | 0.06 | 0.09 | 5.18E-01 |
| S.Pe.C.sup. | Length | 1.00 | - | 6.06E-02 | - | -0.12 | 0.07 | 1.20E-01 |
| S.Pe.C.sup. | MeanDepth | 1.00 | - | 3.75E-01 | - | 0.03 | 0.07 | 6.72E-01 |
| S.Pe.C.sup. | Width | 1.00 | - | 1.38E-02 | - | 0.22 | 0.07 | 2.57E-03 |
| S.Pe.C.sup. | SurfaceArea | 0.49 | 0.42 | 2.17E-01 | 2.07E-01 | -0.06 | 0.08 | 4.32E-01 |
| S.Po.C.sup. | Length | 1.00 | - | 3.94E-03 | - | -0.05 | 0.07 | 4.34E-01 |
| S.Po.C.sup. | MeanDepth | 1.00 | - | 3.56E-03 | - | -0.05 | 0.06 | 4.11E-01 |
| S.Po.C.sup. | Width | 1.00 | - | 6.73E-08 | - | 0.18 | 0.07 | 9.65E-03 |
| S.Po.C.sup. | SurfaceArea | 1.00 | - | 7.80E-04 | - | -0.02 | 0.06 | 7.09E-01 |
| S.R.inf. | Length | 1.00 | - | 7.66E-03 | - | -0.20 | 0.07 | 7.92E-03 |
| S.R.inf. | MeanDepth | 1.00 | - | 2.21E-01 | - | -0.06 | 0.07 | 3.84E-01 |
| S.R.inf. | Width | 0.21 | 0.57 | 7.10E-01 | 1.73E-01 | -0.01 | 0.09 | 9.08E-01 |
| S.R.inf. | SurfaceArea | 1.00 | - | 2.53E-03 | - | -0.20 | 0.07 | 5.97E-03 |
| S.Rh. | Length | 0.70 | 0.49 | 1.47E-01 | 3.14E-01 | 0.24 | 0.07 | 1.10E-03 |
| S.Rh. | MeanDepth | 0.67 | 0.23 | 1.46E-02 | 9.94E-02 | 0.23 | 0.09 | 1.01E-02 |
| S.Rh. | Width | 0.42 | 0.20 | 5.18E-02 | 9.45E-03 | 0.29 | 0.09 | 1.53E-03 |
| S.Rh. | SurfaceArea | 0.57 | 0.27 | 7.26E-02 | 1.02E-01 | 0.31 | 0.08 | 1.51E-04 |
| S.s.P. | Length | 1.00 | - | 1.15E-01 | - | 0.13 | 0.06 | 3.18E-02 |
| S.s.P. | MeanDepth | 1.00 | - | 5.83E-06 | - | 0.03 | 0.07 | 6.72E-01 |
| S.s.P. | Width | 0.73 | 0.20 | 7.43E-04 | 1.06E-01 | 0.12 | 0.08 | 1.23E-01 |
| S.s.P. | SurfaceArea | 1.00 | - | 6.51E-03 | - | 0.13 | 0.06 | 4.93E-02 |
| S.T.i.ant. | Length | 0.83 | 0.64 | 1.18E-01 | 4.04E-01 | 0.06 | 0.07 | 3.47E-01 |
| S.T.i.ant. | MeanDepth | 1.00 | - | 2.41E-02 | - | 0.09 | 0.06 | 1.36E-01 |
| S.T.i.ant. | Width | 1.00 | - | 9.49E-07 | - | 0.04 | 0.07 | 5.35E-01 |
| S.T.i.ant. | SurfaceArea | 1.00 | 0.37 | 2.00E-03 | 4.97E-01 | 0.03 | 0.07 | 6.01E-01 |
| S.T.i.post. | Length | 1.00 | - | 5.49E-01 | - | 0.02 | 0.05 | 7.60E-01 |
| S.T.i.post. | MeanDepth | 1.00 | - | 8.03E-04 | - | -0.07 | 0.07 | 3.37E-01 |
| S.T.i.post. | Width | 1.00 | - | 2.71E-07 | - | -0.01 | 0.08 | 8.62E-01 |
| S.T.i.post. | SurfaceArea | 1.00 | - | 9.03E-02 | - | 0.02 | 0.06 | 6.74E-01 |
| S.T.pol. | Length | 0.55 | 0.14 | 5.59E-04 | 1.74E-03 | 0.14 | 0.08 | 8.73E-02 |
| S.T.pol. | MeanDepth | 0.25 | 0.24 | 3.33E-01 | 4.01E-03 | 0.15 | 0.07 | 4.46E-02 |
| S.T.pol. | Width | 0.58 | 0.15 | 1.83E-03 | 1.18E-02 | 0.19 | 0.08 | 1.65E-02 |
| S.T.pol. | SurfaceArea | 0.66 | 0.14 | 1.29E-05 | 1.15E-02 | 0.06 | 0.08 | 5.13E-01 |
| S.T.s. | Length | 0.34 | 0.46 | 4.44E-01 | 2.25E-01 | 0.07 | 0.07 | 3.30E-01 |
| S.T.s. | MeanDepth | 1.00 | - | 1.25E-09 | - | -0.06 | 0.07 | 3.91E-01 |
| S.T.s. | Width | 0.90 | 0.08 | 2.44E-09 | 1.18E-01 | 0.32 | 0.07 | 2.48E-05 |
| S.T.s. | SurfaceArea | 1.00 | - | 9.43E-03 | - | 0.11 | 0.07 | 1.26E-01 |
| S.T.s.ter.asc.ant. | Length | 1.00 | - | 1.38E-01 | - | -0.04 | 0.06 | 5.34E-01 |
| S.T.s.ter.asc.ant. | MeanDepth | 1.00 | - | 6.17E-02 | - | -0.01 | 0.06 | 8.87E-01 |
| S.T.s.ter.asc.ant. | Width | 1.00 | - | 6.96E-04 | - | -0.03 | 0.07 | 7.00E-01 |
| S.T.s.ter.asc.ant. | SurfaceArea | 1.00 | - | 2.03E-01 | - | 0.01 | 0.06 | 8.02E-01 |
| S.T.s.ter.asc.post. | Length | -0.08 | - | 9.96E-01 | - | -0.03 | - | 6.22E-01 |
| S.T.s.ter.asc.post. | MeanDepth | 1.00 | - | 1.14E-01 | - | 0.00 | 0.06 | 9.46E-01 |
| S.T.s.ter.asc.post. | Width | 1.00 | - | 1.57E-06 | - | 0.04 | 0.07 | 5.85E-01 |
| S.T.s.ter.asc.post. | SurfaceArea | 1.00 | - | 2.37E-01 | - | -0.05 | 0.07 | 5.09E-01 |
| Globals | Length | 0.99 | 0.01 | 4.06E-12 | 1.71E-01 | 0.84 | 0.02 | 2.19E-81 |
| Globals | MeanDepth | 1.00 | - | 7.23E-17 | - | 0.86 | 0.02 | 7.61E-97 |
| Globals | Width | 1.00 | - | 1.15E-10 | - | 0.73 | 0.03 | 2.19E-59 |
| Globals | SurfaceArea | 0.99 | 0.01 | 1.62E-16 | 1.60E-01 | 0.90 | 0.02 | 4.92E-127 |

**Table S24:** Bivariate analysis: left-right genetic correlation in GOBS (yellow: Bonferroni corrected; grey: p < 0.05).

| Sulci | Descriptor | rhoG | rhoG_SE | rhoP=0 | rhoP=1 | rhoE | rhoE.se | rhoE.p |
|---|---|---|---|---|---|---|---|---|
| F.C.L.a. | Length | 0.40 | 0.87 | 6.23E-01 | 3.38E-01 | 0.07 | 0.05 | 1.80E-01 |
| F.C.L.a. | MeanDepth | 1.00 | - | 1.27E-02 | - | 0.06 | 0.05 | - |
| F.C.L.a. | Width | 0.83 | 0.07 | 1.21E-07 | 4.39E-03 | 0.53 | 0.05 | 2.03E-10 |
| F.C.L.a. | SurfaceArea | 0.52 | 0.19 | 1.44E-02 | 9.59E-03 | 0.23 | 0.07 | 9.38E-04 |
| F.C.L.p. | Length | 1.00 | - | 1.72E-02 | - | 0.03 | 0.05 | 4.61E-01 |
| F.C.L.p. | MeanDepth | 0.90 | 3.72 | 6.30E-01 | 1.00E+00 | 0.06 | 0.05 | - |
| F.C.L.p. | Width | 1.00 | - | 2.68E-12 | - | 0.61 | 0.03 | 8.51E-19 |
| F.C.L.p. | SurfaceArea | 0.98 | 0.14 | 4.38E-07 | 4.57E-01 | 0.16 | 0.07 | 2.55E-02 |
| F.C.L.r.ant. | Length | 1.00 | - | 2.31E-02 | - | 0.09 | 0.05 | 1.23E-01 |
| F.C.L.r.ant. | MeanDepth | 0.41 | 0.40 | 2.80E-01 | 1.35E-01 | 0.02 | 0.06 | - |
| F.C.L.r.ant. | Width | 1.00 | - | 5.97E-02 | - | 0.22 | 0.05 | 1.21E-04 |
| F.C.L.r.ant. | SurfaceArea | 1.00 | - | 1.10E-04 | - | 0.04 | 0.06 | 4.74E-01 |
| F.C.L.r.asc. | Length | 1.00 | - | 2.13E-03 | - | -0.04 | 0.05 | 4.41E-01 |
| F.C.L.r.asc. | MeanDepth | 1.00 | - | 5.16E-04 | - | 0.07 | 0.05 | - |
| F.C.L.r.asc. | Width | 0.70 | 0.26 | 2.36E-02 | 1.40E-01 | 0.36 | 0.05 | 5.35E-11 |
| F.C.L.r.asc. | SurfaceArea | 1.00 | - | 1.06E-06 | - | -0.03 | 0.06 | 6.35E-01 |
| F.C.L.r.diag. | Length | 1.00 | - | 7.63E-01 | - | 0.10 | 0.08 | 2.10E-01 |
| F.C.L.r.diag. | MeanDepth | 0.16 | - | 1.00E+00 | 1.00E+00 | 0.14 | - | - |
| F.C.L.r.diag. | Width | 1.00 | 0.80 | 1.53E-01 | 5.00E-01 | 0.08 | 0.11 | 4.13E-01 |
| F.C.L.r.diag. | SurfaceArea | 0.22 | - | 1.00E+00 | 1.00E+00 | 0.12 | - | 6.24E-02 |
| F.C.L.r.retroC.tr. | Length | 0.25 | 0.54 | 6.36E-01 | 1.34E-01 | -0.03 | 0.06 | 6.61E-01 |
| F.C.L.r.retroC.tr. | MeanDepth | 1.00 | - | 2.95E-03 | - | -0.02 | 0.05 | - |
| F.C.L.r.retroC.tr. | Width | 1.00 | - | 1.17E-03 | - | 0.06 | 0.05 | 2.14E-01 |
| F.C.L.r.retroC.tr. | SurfaceArea | 0.80 | 0.53 | 7.26E-02 | 3.63E-01 | -0.03 | 0.06 | 6.34E-01 |
| F.C.L.r.sc.ant. | Length | - | - | - | - | - | - | - |
| F.C.L.r.sc.ant. | MeanDepth | -0.08 | 0.86 | 9.34E-01 | - | 0.90 | 5.56 | - |
| F.C.L.r.sc.ant. | Width | -0.47 | 0.97 | 6.00E-01 | - | 0.32 | 1.05 | 7.69E-01 |
| F.C.L.r.sc.ant. | SurfaceArea | 0.21 | - | 1.00E+00 | 1 | 0.07 | - | 7.07E-01 |
| F.C.L.r.sc.post. | Length | 1.00 | - | 3.54E-02 | - | -1.00 | - | 3.36E-01 |
| F.C.L.r.sc.post. | MeanDepth | 1.00 | - | 1.49E-01 | - | -0.01 | 0.05 | - |
| F.C.L.r.sc.post. | Width | 1.00 | - | 6.64E-03 | - | 0.19 | 0.05 | 1.93E-03 |
| F.C.L.r.sc.post. | SurfaceArea | 0.07 | 1.29 | 9.56E-01 | 3.51E-01 | 0.06 | 0.06 | 3.46E-01 |
| F.C.M.ant. | Length | 0.48 | 0.41 | 2.06E-01 | 1.49E-01 | -0.03 | 0.06 | 6.16E-01 |
| F.C.M.ant. | MeanDepth | 0.53 | 0.22 | 1.61E-02 | 2.88E-02 | -0.16 | 0.06 | - |
| F.C.M.ant. | Width | 0.85 | 0.14 | 1.22E-05 | 1.46E-01 | 0.18 | 0.06 | 4.41E-03 |
| F.C.M.ant. | SurfaceArea | 0.83 | 0.29 | 2.47E-03 | 2.88E-01 | -0.11 | 0.06 | 5.30E-02 |
| F.C.M.post. | Length | 1.00 | - | 7.05E-01 | - | 0.06 | 0.04 | 1.39E-01 |
| F.C.M.post. | MeanDepth | 0.94 | 0.15 | 3.25E-07 | 3.40E-01 | 0.16 | 0.06 | - |
| F.C.M.post. | Width | 0.87 | 0.10 | 4.51E-05 | 1.01E-01 | 0.56 | 0.04 | 1.23E-17 |
| F.C.M.post. | SurfaceArea | 1.00 | - | 5.48E-05 | - | 0.09 | 0.05 | 9.99E-02 |
| F.Cal.ant.-Sc.Cal. | Length | - | - | - | - | - | - | - |
| F.Cal.ant.-Sc.Cal. | MeanDepth | - | - | - | - | - | - | - |
| F.Cal.ant.-Sc.Cal. | Width | - | - | - | - | - | - | - |
| F.Cal.ant.-Sc.Cal. | SurfaceArea | - | - | - | - | - | - | - |
| F.Coll. | Length | 1.00 | - | 1.84E-01 | - | 0.08 | 0.04 | 7.46E-02 |
| F.Coll. | MeanDepth | 0.84 | 0.09 | 3.08E-12 | 2.35E-02 | 0.10 | 0.08 | - |
| F.Coll. | Width | 0.88 | 0.12 | 1.90E-06 | 1.56E-01 | 0.33 | 0.05 | 1.59E-07 |
| F.Coll. | SurfaceArea | 1.00 | - | 9.58E-08 | - | 0.10 | 0.05 | 9.20E-02 |
| F.I.P. | Length | 1.00 | - | 1.28E-04 | - | 0.01 | 0.05 | 8.27E-01 |
| F.I.P. | MeanDepth | 0.88 | 0.16 | 3.35E-04 | 2.31E-01 | 0.25 | 0.05 | - |
| F.I.P. | Width | 1.00 | 0.04 | 3.77E-11 | 4.62E-01 | 0.65 | 0.04 | 1.01E-15 |
| F.I.P. | SurfaceArea | 0.68 | 0.19 | 2.13E-03 | 4.89E-02 | 0.12 | 0.06 | 5.24E-02 |
| F.I.P.Po.C.inf. | Length | 0.51 | 0.52 | 3.17E-01 | 2.11E-01 | 0.00 | 0.05 | 9.60E-01 |
| F.I.P.Po.C.inf. | MeanDepth | 0.89 | 0.29 | 5.94E-03 | 3.55E-01 | 0.06 | 0.06 | - |
| F.I.P.Po.C.inf. | Width | 0.93 | 0.09 | 1.11E-05 | 2.06E-01 | 0.55 | 0.04 | 2.13E-16 |
| F.I.P.Po.C.inf. | SurfaceArea | 0.63 | 0.43 | 1.27E-01 | 2.19E-01 | 0.01 | 0.05 | 8.22E-01 |
| F.I.P.r.int.1 | Length | -0.40 | 1.82 | 7.60E-01 | - | 0.13 | 0.06 | 2.24E-02 |
| F.I.P.r.int.1 | MeanDepth | -0.54 | 0.38 | 1.27E-01 | - | 0.18 | 0.06 | - |
| F.I.P.r.int.1 | Width | 0.39 | 0.29 | 2.26E-01 | 0.0240789 | 0.22 | 0.05 | 2.38E-04 |
| F.I.P.r.int.1 | SurfaceArea | -0.34 | 0.85 | 6.46E-01 | - | 0.13 | 0.06 | 2.27E-02 |
| F.I.P.r.int.2 | Length | 1.00 | - | 4.61E-01 | - | -0.02 | 0.06 | 7.98E-01 |
| F.I.P.r.int.2 | MeanDepth | 0.24 | 1.11 | 8.26E-01 | 0.3901107 | 0.04 | 0.07 | - |

| | | | | | | | | |
|---|---|---|---|---|---|---|---|---|
| F.I.P.r.int.2 | Width | 0.06 | 0.66 | 9.24E-01 | 0.1923586 | 0.09 | 0.07 | 2.38E-01 |
| F.I.P.r.int.2 | SurfaceArea | 1.00 | - | 2.03E-01 | - | 0.00 | 0.06 | 9.93E-01 |
| F.P.O. | Length | 0.49 | 0.42 | 2.84E-01 | 1.32E-01 | 0.14 | 0.05 | 8.17E-03 |
| F.P.O. | MeanDepth | 0.98 | 0.08 | 1.77E-11 | 4.27E-01 | -0.03 | 0.08 | - |
| F.P.O. | Width | 1.00 | - | 7.40E-13 | - | 0.41 | 0.05 | 1.54E-09 |
| F.P.O. | SurfaceArea | 0.74 | 0.12 | 9.97E-06 | 1.31E-02 | 0.14 | 0.07 | 5.72E-02 |
| INSULA | Length | -1.00 | - | 4.63E-01 | - | 0.08 | 0.29 | 7.86E-01 |
| INSULA | MeanDepth | 0.73 | 0.38 | 4.62E-02 | 2.63E-01 | 0.01 | 0.06 | - |
| INSULA | Width | 0.93 | 0.07 | 8.16E-06 | 1.20E-01 | 0.74 | 0.03 | 6.66E-26 |
| INSULA | SurfaceArea | 0.98 | 0.15 | 1.81E-07 | 4.42E-01 | 0.08 | 0.07 | 2.76E-01 |
| OCCIPITAL | Length | 0.85 | 0.17 | 3.56E-05 | 1.80E-01 | 0.12 | 0.06 | 6.25E-02 |
| OCCIPITAL | MeanDepth | 0.86 | 0.23 | 2.93E-04 | 2.74E-01 | 0.07 | 0.06 | - |
| OCCIPITAL | Width | 0.87 | 0.07 | 7.96E-10 | 1.28E-02 | 0.42 | 0.06 | 1.45E-07 |
| OCCIPITAL | SurfaceArea | 0.75 | 0.14 | 3.89E-05 | 3.62E-02 | 0.13 | 0.07 | 6.53E-02 |
| S.C. | Length | 1.00 | - | 6.16E-03 | - | 0.05 | 0.05 | 3.57E-01 |
| S.C. | MeanDepth | 0.83 | 0.15 | 7.20E-06 | 1.19E-01 | 0.14 | 0.06 | - |
| S.C. | Width | 1.00 | - | 3.14E-16 | - | 0.74 | 0.03 | 4.11E-19 |
| S.C. | SurfaceArea | 1.00 | - | 7.97E-08 | - | 0.10 | 0.06 | 1.18E-01 |
| S.C.LPC. | Length | -1.00 | - | 1.53E-01 | - | 0.25 | 0.31 | 3.19E-01 |
| S.C.LPC. | MeanDepth | 0.13 | 442538603000.0( | 1.00E+00 | 1.00E+00 | 0.05 | 0.16 | - |
| S.C.LPC. | Width | -0.90 | - | 8.09E-01 | - | 0.20 | - | 2.66E-01 |
| S.C.LPC. | SurfaceArea | -1.00 | - | 3.57E-01 | - | 0.12 | 0.22 | 5.65E-01 |
| S.C.sylvian. | Length | 1.00 | - | 2.35E-01 | - | 0.11 | 0.06 | 1.13E-01 |
| S.C.sylvian. | MeanDepth | 1.00 | - | 1.01E-03 | - | -0.09 | 0.08 | - |
| S.C.sylvian. | Width | 1.00 | - | 4.21E-02 | - | 0.06 | 0.07 | 3.98E-01 |
| S.C.sylvian. | SurfaceArea | 1.00 | - | 5.82E-02 | - | 0.04 | 0.07 | 6.02E-01 |
| S.Call. | Length | 0.84 | 0.14 | 1.46E-06 | 1.27E-01 | 0.06 | 0.07 | 4.43E-01 |
| S.Call. | MeanDepth | 0.81 | 0.12 | 1.56E-07 | 5.47E-02 | -0.02 | 0.08 | - |
| S.Call. | Width | 1.00 | - | 3.56E-07 | - | 0.06 | 0.06 | 3.10E-01 |
| S.Call. | SurfaceArea | 0.87 | 0.10 | 4.94E-10 | 8.57E-02 | 0.07 | 0.08 | 3.93E-01 |
| S.Cu. | Length | -0.90 | - | 8.54E-01 | - | 0.06 | - | 5.77E-02 |
| S.Cu. | MeanDepth | 1.00 | - | 2.37E-04 | - | -0.08 | 0.05 | - |
| S.Cu. | Width | 1.00 | - | 2.76E-07 | - | 0.04 | 0.06 | 4.43E-01 |
| S.Cu. | SurfaceArea | 0.78 | 2.48 | 6.93E-01 | 4.69E-01 | 0.03 | 0.05 | 4.54E-01 |
| S.F.inf. | Length | 1.00 | - | 2.63E-02 | - | 0.00 | 0.05 | 9.35E-01 |
| S.F.inf. | MeanDepth | 1.00 | - | 2.31E-10 | - | 0.13 | 0.07 | - |
| S.F.inf. | Width | 0.81 | 0.12 | 7.80E-05 | 4.34E-02 | 0.36 | 0.05 | 3.60E-08 |
| S.F.inf. | SurfaceArea | 0.78 | 0.24 | 3.47E-03 | 1.87E-01 | 0.12 | 0.06 | 5.92E-02 |
| S.F.inf.ant. | Length | 1.00 | - | 1.66E-01 | - | 0.01 | 0.05 | 8.48E-01 |
| S.F.inf.ant. | MeanDepth | 0.59 | 0.59 | 3.03E-01 | 2.83E-01 | 0.02 | 0.06 | - |
| S.F.inf.ant. | Width | 0.69 | 0.34 | 5.20E-02 | 2.21E-01 | 0.12 | 0.06 | 4.17E-02 |
| S.F.inf.ant. | SurfaceArea | 0.86 | 0.64 | 1.07E-01 | 4.20E-01 | 0.02 | 0.06 | 7.64E-01 |
| S.F.int. | Length | 0.99 | 0.40 | 1.19E-02 | 4.85E-01 | 0.07 | 0.05 | 1.52E-01 |
| S.F.int. | MeanDepth | 0.78 | 0.27 | 3.08E-03 | 2.27E-01 | -0.06 | 0.06 | - |
| S.F.int. | Width | 0.99 | 0.13 | 3.52E-07 | 4.61E-01 | 0.17 | 0.06 | 3.28E-03 |
| S.F.int. | SurfaceArea | 1.00 | - | 1.13E-03 | - | 0.06 | 0.05 | 2.85E-01 |
| S.F.inter. | Length | 1.00 | - | 2.40E-02 | - | 0.06 | 0.05 | 1.98E-01 |
| S.F.inter. | MeanDepth | 0.68 | 0.38 | 9.85E-02 | 2.22E-01 | 0.23 | 0.05 | - |
| S.F.inter. | Width | 0.85 | 0.10 | 1.76E-07 | 5.47E-02 | 0.41 | 0.05 | 2.51E-09 |
| S.F.inter. | SurfaceArea | 1.00 | - | 7.54E-04 | - | 0.11 | 0.05 | 4.15E-02 |
| S.F.marginal. | Length | 1.00 | - | 8.49E-02 | - | -0.01 | 0.05 | 9.07E-01 |
| S.F.marginal. | MeanDepth | 0.83 | 1.21 | 4.69E-01 | 4.47E-01 | 0.11 | 0.05 | - |
| S.F.marginal. | Width | 1.00 | - | 1.90E-04 | - | 0.22 | 0.05 | 2.64E-05 |
| S.F.marginal. | SurfaceArea | 0.89 | 0.39 | 2.59E-02 | 3.89E-01 | 0.05 | 0.05 | 3.15E-01 |
| S.F.median. | Length | 1.00 | - | 1.90E-04 | - | 0.15 | 0.05 | 3.28E-03 |
| S.F.median. | MeanDepth | 1.00 | - | 2.14E-02 | - | 0.06 | 0.05 | - |
| S.F.median. | Width | 1.00 | - | 1.06E-05 | - | 0.24 | 0.05 | 3.63E-05 |
| S.F.median. | SurfaceArea | 1.00 | - | 1.93E-04 | - | 0.15 | 0.05 | 3.97E-03 |
| S.F.orbitaire. | Length | 0.26 | 1.04 | 8.08E-01 | 2.85E-01 | 0.03 | 0.06 | 5.98E-01 |
| S.F.orbitaire. | MeanDepth | 0.32 | 1.60 | 8.33E-01 | 3.80E-01 | 0.00 | 0.05 | - |
| S.F.orbitaire. | Width | 1.00 | - | 1.09E-03 | - | -0.05 | 0.06 | 3.65E-01 |
| S.F.orbitaire. | SurfaceArea | 0.41 | 1.00 | 6.64E-01 | 3.46E-01 | 0.00 | 0.06 | 9.45E-01 |
| S.F.polaire.tr. | Length | 0.95 | 0.34 | 1.05E-03 | 4.48E-01 | 0.06 | 0.06 | 2.90E-01 |
| S.F.polaire.tr. | MeanDepth | 1.00 | - | 1.00E-01 | - | 0.07 | 0.05 | - |

| | | | | | | | | |
|---|---|---|---|---|---|---|---|---|
| S.F.polaire.tr. | Width | 1.00 | - | 4.86E-05 | - | 0.20 | 0.05 | 3.56E-04 |
| S.F.polaire.tr. | SurfaceArea | 0.99 | 0.45 | 4.32E-03 | 4.89E-01 | 0.08 | 0.06 | 1.54E-01 |
| S.F.sup. | Length | 1.00 | - | 2.58E-03 | - | -0.05 | 0.05 | 3.03E-01 |
| S.F.sup. | MeanDepth | 0.99 | 0.11 | 2.04E-09 | 4.59E-01 | 0.09 | 0.07 | - |
| S.F.sup. | Width | 0.85 | 0.08 | 3.56E-06 | 1.92E-02 | 0.63 | 0.03 | 1.40E-23 |
| S.F.sup. | SurfaceArea | 1.00 | - | 2.65E-04 | - | 0.04 | 0.05 | 3.73E-01 |
| S.Li.ant. | Length | 0.27 | 0.29 | 3.57E-01 | 1.15E-02 | 0.04 | 0.07 | 5.21E-01 |
| S.Li.ant. | MeanDepth | 0.30 | 0.50 | 5.69E-01 | 1.10E-01 | 0.14 | 0.06 | - |
| S.Li.ant. | Width | 0.65 | 0.58 | 3.06E-01 | 3.21E-01 | 0.18 | 0.06 | 3.09E-03 |
| S.Li.ant. | SurfaceArea | 0.08 | 0.33 | 8.06E-01 | 1.96E-02 | 0.11 | 0.07 | 1.19E-01 |
| S.Li.post. | Length | 0.90 | 1.84 | 5.29E-01 | 1.00E+00 | 0.01 | 0.05 | 8.56E-01 |
| S.Li.post. | MeanDepth | 1.00 | - | 2.62E-02 | - | 0.03 | 0.05 | - |
| S.Li.post. | Width | 1.00 | - | 3.63E-02 | - | 0.13 | 0.05 | 1.66E-02 |
| S.Li.post. | SurfaceArea | 0.39 | - | 1.00E+00 | 1.00E+00 | 0.05 | - | 1.44E-01 |
| S.O.p. | Length | -0.04 | 0.46 | 9.34E-01 | - | 0.05 | 0.06 | 4.01E-01 |
| S.O.p. | MeanDepth | 0.53 | 0.66 | 4.37E-01 | 2.83E-01 | 0.09 | 0.06 | - |
| S.O.p. | Width | 1.00 | - | 2.24E-04 | - | 0.04 | 0.06 | 5.27E-01 |
| S.O.p. | SurfaceArea | 0.43 | 0.49 | 3.73E-01 | 1.72E-01 | 0.04 | 0.06 | 5.29E-01 |
| S.O.T.lat.ant. | Length | 1.00 | - | 9.64E-09 | - | 0.02 | 0.07 | 7.98E-01 |
| S.O.T.lat.ant. | MeanDepth | 0.93 | 0.11 | 5.62E-08 | 2.75E-01 | 0.28 | 0.06 | - |
| S.O.T.lat.ant. | Width | 0.66 | 0.14 | 5.90E-04 | 7.44E-03 | 0.42 | 0.06 | 2.72E-09 |
| S.O.T.lat.ant. | SurfaceArea | 0.99 | 0.09 | 3.77E-11 | 4.41E-01 | 0.20 | 0.07 | 1.19E-02 |
| S.O.T.lat.int. | Length | 1.00 | - | 1.28E-01 | - | 0.06 | 0.05 | 2.62E-01 |
| S.O.T.lat.int. | MeanDepth | 0.02 | 0.38 | 9.53E-01 | 5.30E-02 | 0.16 | 0.07 | - |
| S.O.T.lat.int. | Width | 1.00 | - | 2.31E-01 | - | 0.13 | 0.06 | 3.58E-02 |
| S.O.T.lat.int. | SurfaceArea | 0.69 | 1.57 | 5.82E-01 | 4.39E-01 | 0.09 | 0.06 | 1.15E-01 |
| S.O.T.lat.med. | Length | 0.90 | 1.32 | 3.70E-01 | 1.00E+00 | 0.00 | 0.05 | 1.00E+00 |
| S.O.T.lat.med. | MeanDepth | 0.80 | 0.61 | 1.26E-01 | 3.81E-01 | -0.02 | 0.05 | - |
| S.O.T.lat.med. | Width | 1.00 | - | 1.46E-03 | - | 0.00 | 0.05 | 9.41E-01 |
| S.O.T.lat.med. | SurfaceArea | 1.00 | - | 6.78E-02 | - | 0.02 | 0.04 | 6.27E-01 |
| S.O.T.lat.post. | Length | 0.84 | 1.74 | 4.56E-01 | 4.69E-01 | 0.02 | 0.05 | 7.29E-01 |
| S.O.T.lat.post. | MeanDepth | 1.00 | - | 1.11E-01 | - | 0.13 | 0.05 | - |
| S.O.T.lat.post. | Width | 1.00 | - | 1.42E-02 | - | 0.20 | 0.04 | 6.87E-05 |
| S.O.T.lat.post. | SurfaceArea | 0.82 | 0.48 | 6.74E-02 | 3.67E-01 | 0.02 | 0.05 | 6.76E-01 |
| S.Olf. | Length | 0.99 | 0.09 | 3.99E-11 | 4.54E-01 | 0.10 | 0.07 | 1.74E-01 |
| S.Olf. | MeanDepth | 0.90 | 0.08 | 7.65E-09 | 1.27E-01 | 0.42 | 0.05 | - |
| S.Olf. | Width | 0.97 | 0.12 | 1.18E-05 | 4.02E-01 | 0.37 | 0.05 | 6.47E-09 |
| S.Olf. | SurfaceArea | 1.00 | - | 2.13E-14 | - | 0.27 | 0.07 | 1.31E-03 |
| S.Or. | Length | 1.00 | - | 2.07E-03 | - | 0.12 | 0.05 | 2.55E-02 |
| S.Or. | MeanDepth | 0.68 | 0.13 | 1.52E-04 | 6.31E-03 | 0.28 | 0.06 | - |
| S.Or. | Width | 1.00 | - | 1.56E-06 | - | 0.26 | 0.06 | 1.35E-04 |
| S.Or. | SurfaceArea | 1.00 | - | 3.88E-05 | - | 0.26 | 0.05 | 6.15E-05 |
| S.p.C. | Length | 1.00 | - | 1.40E-01 | - | -0.05 | 0.05 | 3.36E-01 |
| S.p.C. | MeanDepth | 1.00 | - | 4.73E-01 | - | 0.00 | 0.05 | - |
| S.p.C. | Width | 0.47 | 0.86 | 6.05E-01 | 3.00E-01 | 0.09 | 0.06 | 1.21E-01 |
| S.p.C. | SurfaceArea | 1.00 | - | 2.30E-01 | - | -0.01 | 0.05 | 9.10E-01 |
| S.Pa.int. | Length | 1.00 | - | 6.60E-03 | - | 0.04 | 0.05 | 4.15E-01 |
| S.Pa.int. | MeanDepth | 1.00 | - | 1.48E-02 | - | 0.01 | 0.05 | - |
| S.Pa.int. | Width | 0.94 | 0.16 | 6.52E-06 | 3.61E-01 | 0.15 | 0.06 | 1.00E-02 |
| S.Pa.int. | SurfaceArea | 1.00 | - | 1.46E-02 | - | 0.06 | 0.05 | 2.21E-01 |
| S.Pa.sup. | Length | 1.00 | - | 1.09E-02 | - | -0.06 | 0.05 | 2.17E-01 |
| S.Pa.sup. | MeanDepth | 1.00 | - | 8.79E-02 | - | -0.01 | 0.05 | - |
| S.Pa.sup. | Width | 1.00 | - | 2.44E-03 | - | 0.11 | 0.05 | 6.54E-02 |
| S.Pa.sup. | SurfaceArea | 1.00 | - | 1.39E-03 | - | -0.05 | 0.05 | 3.10E-01 |
| S.Pa.t. | Length | 1.00 | - | 1.42E-01 | - | 0.00 | 0.07 | 9.49E-01 |
| S.Pa.t. | MeanDepth | 0.19 | - | 1.00E+00 | 1.00E+00 | 0.12 | - | - |
| S.Pa.t. | Width | 0.48 | 0.41 | 2.40E-01 | 1.71E-01 | 0.11 | 0.07 | 1.27E-01 |
| S.Pa.t. | SurfaceArea | -0.90 | 11.00 | 8.75E-01 | - | 0.13 | 0.08 | 8.72E-04 |
| S.Pe.C.inf. | Length | 1.00 | - | 9.50E-03 | - | -0.03 | 0.05 | 5.91E-01 |
| S.Pe.C.inf. | MeanDepth | 1.00 | - | 2.09E-04 | - | 0.09 | 0.06 | - |
| S.Pe.C.inf. | Width | 1.00 | - | 6.26E-08 | - | 0.32 | 0.05 | 1.77E-06 |
| S.Pe.C.inf. | SurfaceArea | 1.00 | - | 3.09E-03 | - | 0.03 | 0.05 | 6.13E-01 |
| S.Pe.C.inter. | Length | 1.00 | - | 7.07E-01 | - | -0.01 | 0.04 | 8.58E-01 |
| S.Pe.C.inter. | MeanDepth | 0.99 | 0.33 | 2.34E-03 | 4.84E-01 | 0.03 | 0.05 | - |

| | | | | | | | | |
|---|---|---|---|---|---|---|---|---|
| S.Pe.C.inter. | Width | 0.97 | 0.13 | 1.02E-06 | 4.20E-01 | 0.32 | 0.05 | 8.97E-08 |
| S.Pe.C.inter. | SurfaceArea | 1.00 | - | 3.90E-02 | - | 0.01 | 0.05 | 8.92E-01 |
| S.Pe.C.marginal. | Length | -1.00 | - | 3.83E-01 | - | 0.07 | 0.05 | 1.65E-01 |
| S.Pe.C.marginal. | MeanDepth | 0.90 | 2.47 | 5.92E-01 | 1.00E+00 | -0.01 | 0.07 | - |
| S.Pe.C.marginal. | Width | 1.00 | - | 5.35E-04 | - | 0.07 | 0.06 | 2.37E-01 |
| S.Pe.C.marginal. | SurfaceArea | -1.00 | - | 1.53E-01 | - | 0.09 | 0.05 | 8.75E-02 |
| S.Pe.C.median. | Length | 1.00 | - | 2.83E-01 | - | 0.05 | 0.05 | 2.52E-01 |
| S.Pe.C.median. | MeanDepth | 1.00 | - | 2.42E-01 | - | 0.08 | 0.05 | - |
| S.Pe.C.median. | Width | 1.00 | - | 4.91E-04 | - | 0.09 | 0.05 | 1.29E-01 |
| S.Pe.C.median. | SurfaceArea | 1.00 | - | 1.93E-01 | - | 0.08 | 0.05 | 1.00E-01 |
| S.Pe.C.sup. | Length | 1.00 | - | 1.01E-02 | - | 0.01 | 0.05 | 8.02E-01 |
| S.Pe.C.sup. | MeanDepth | 0.89 | 0.43 | 2.40E-02 | 4.02E-01 | 0.01 | 0.06 | - |
| S.Pe.C.sup. | Width | 1.00 | - | 1.74E-05 | - | 0.27 | 0.05 | 1.19E-06 |
| S.Pe.C.sup. | SurfaceArea | 1.00 | - | 3.26E-02 | - | 0.01 | 0.05 | 8.69E-01 |
| S.Po.C.sup. | Length | 0.90 | 12.82 | 8.20E-01 | 1.00E+00 | 0.08 | 0.05 | 1.57E-01 |
| S.Po.C.sup. | MeanDepth | 0.59 | 0.30 | 5.45E-02 | 1.03E-01 | -0.01 | 0.06 | - |
| S.Po.C.sup. | Width | 1.00 | - | 3.07E-07 | - | 0.18 | 0.06 | 5.03E-03 |
| S.Po.C.sup. | SurfaceArea | 0.12 | 0.28 | 6.73E-01 | 8.22E-03 | 0.10 | 0.06 | 1.01E-01 |
| S.R.inf. | Length | 1.00 | - | 2.00E-01 | - | -0.11 | 0.06 | 7.33E-02 |
| S.R.inf. | MeanDepth | 1.00 | - | 1.57E-01 | - | -0.11 | 0.05 | - |
| S.R.inf. | Width | 1.00 | - | 3.68E-02 | - | 0.02 | 0.05 | 7.36E-01 |
| S.R.inf. | SurfaceArea | 1.00 | - | 1.17E-01 | - | -0.11 | 0.06 | 4.84E-02 |
| S.Rh. | Length | 0.74 | 0.21 | 1.13E-03 | 1.13E-01 | 0.11 | 0.07 | 1.26E-01 |
| S.Rh. | MeanDepth | 0.86 | 0.26 | 1.13E-02 | 2.95E-01 | 0.24 | 0.06 | - |
| S.Rh. | Width | 1.00 | - | 2.62E-02 | - | 0.37 | 0.04 | 5.27E-10 |
| S.Rh. | SurfaceArea | 0.73 | 0.25 | 7.63E-03 | 1.54E-01 | 0.17 | 0.06 | 1.35E-02 |
| S.s.P. | Length | 0.67 | 0.32 | 4.78E-02 | 1.78E-01 | 0.08 | 0.06 | 2.00E-01 |
| S.s.P. | MeanDepth | 1.00 | - | 5.16E-07 | - | 0.04 | 0.05 | - |
| S.s.P. | Width | 1.00 | - | 1.54E-08 | - | 0.26 | 0.05 | 1.99E-05 |
| S.s.P. | SurfaceArea | 0.96 | 0.22 | 1.67E-04 | 4.24E-01 | 0.08 | 0.06 | 1.89E-01 |
| S.T.i.ant. | Length | 1.00 | - | 6.67E-04 | - | 0.09 | 0.05 | 1.03E-01 |
| S.T.i.ant. | MeanDepth | 0.96 | 0.20 | 3.14E-06 | 4.18E-01 | 0.01 | 0.06 | - |
| S.T.i.ant. | Width | 0.97 | 0.12 | 2.38E-07 | 4.06E-01 | 0.26 | 0.06 | 1.12E-04 |
| S.T.i.ant. | SurfaceArea | 1.00 | - | 8.83E-07 | - | 0.11 | 0.05 | 5.43E-02 |
| S.T.i.post. | Length | -1.00 | - | 6.78E-01 | - | 0.01 | 0.05 | 8.89E-01 |
| S.T.i.post. | MeanDepth | 0.78 | 0.39 | 2.91E-02 | 3.10E-01 | 0.03 | 0.05 | - |
| S.T.i.post. | Width | 1.00 | 0.14 | 3.89E-08 | 4.88E-01 | 0.15 | 0.06 | 1.01E-02 |
| S.T.i.post. | SurfaceArea | 0.55 | 0.69 | 3.25E-01 | 3.42E-01 | -0.01 | 0.06 | 8.89E-01 |
| S.T.pol. | Length | 0.82 | 0.24 | 9.90E-04 | 2.32E-01 | 0.03 | 0.06 | 6.68E-01 |
| S.T.pol. | MeanDepth | 0.77 | 0.23 | 3.72E-03 | 1.67E-01 | 0.15 | 0.06 | - |
| S.T.pol. | Width | 0.85 | 0.16 | 8.91E-05 | 1.82E-01 | 0.33 | 0.05 | 2.10E-08 |
| S.T.pol. | SurfaceArea | 0.79 | 0.19 | 1.44E-04 | 1.36E-01 | 0.08 | 0.06 | 2.14E-01 |
| S.T.s. | Length | 0.44 | 0.32 | 1.90E-01 | 7.91E-02 | 0.02 | 0.06 | 7.16E-01 |
| S.T.s. | MeanDepth | 0.81 | 0.10 | 1.83E-08 | 3.07E-02 | 0.12 | 0.07 | - |
| S.T.s. | Width | 0.92 | 0.06 | 2.90E-06 | 7.97E-02 | 0.63 | 0.03 | 1.74E-19 |
| S.T.s. | SurfaceArea | 0.87 | 0.68 | 9.31E-02 | 4.34E-01 | 0.11 | 0.05 | 5.28E-02 |
| S.T.s.ter.asc.ant. | Length | 0.09 | - | 1.00E+00 | 1 | 0.01 | - | 7.81E-01 |
| S.T.s.ter.asc.ant. | MeanDepth | 0.03 | - | 1.00E+00 | 0.5 | 0.07 | - | - |
| S.T.s.ter.asc.ant. | Width | 1.00 | - | 1.16E-01 | - | 0.22 | 0.04 | 3.52E-06 |
| S.T.s.ter.asc.ant. | SurfaceArea | 0.11 | - | 1.00E+00 | 1 | 0.03 | - | 3.00E-01 |
| S.T.s.ter.asc.post. | Length | -1.00 | - | 1.22E-01 | - | 0.08 | 0.05 | 6.94E-02 |
| S.T.s.ter.asc.post. | MeanDepth | 0.88 | 0.59 | 1.07E-01 | 4.26E-01 | 0.03 | 0.06 | - |
| S.T.s.ter.asc.post. | Width | 0.64 | 0.25 | 4.26E-02 | 8.30E-02 | 0.31 | 0.05 | 1.06E-08 |
| S.T.s.ter.asc.post. | SurfaceArea | 0.21 | 0.77 | 7.75E-01 | 2.89E-01 | 0.03 | 0.05 | 5.43E-01 |
| Globals | Length | 0.55 | 0.32 | 1.16E-01 | 1.11E-01 | 0.20 | 0.06 | 1.06E-03 |
| Globals | MeanDepth | 0.93 | 0.40 | 1.68E-02 | 4.31E-01 | 0.02 | 0.06 | - |
| Globals | Width | 0.97 | 0.15 | 6.06E-08 | 4.14E-01 | 0.22 | 0.06 | 5.44E-04 |
| Globals | SurfaceArea | 0.72 | 0.16 | 1.31E-04 | 3.93E-02 | 0.18 | 0.06 | 7.31E-03 |

**Table S25:** Meta-analysis of left-right genetic correlation
(yellow: Bonferroni corrected; grey: $p < 0.05$).

| Sulci | Descriptor | rhoG | rhoP=0 | rhoP=1 |
|---|---|---|---|---|
| F.C.L.a. | Length | 0.76 | 1.48E-01 | 6.44E-03 |
| F.C.L.a. | MeanDepth | 0.65 | 1.41E-01 | - |
| F.C.L.a. | Width | 0.80 | 6.41E-04 | 5.49E-02 |
| F.C.L.a. | SurfaceArea | 0.81 | 5.67E-05 | 1.47E-07 |
| F.C.L.p. | Length | 0.75 | 1.12E-01 | 1.26E-02 |
| F.C.L.p. | MeanDepth | 0.94 | 4.57E-02 | 4.49E-01 |
| F.C.L.p. | Width | 0.93 | 1.77E-29 | 2.92E-04 |
| F.C.L.p. | SurfaceArea | 0.96 | 1.29E-16 | 2.85E-01 |
| F.C.L.r.ant. | Length | 0.96 | 1.78E-03 | 1.15E-02 |
| F.C.L.r.ant. | MeanDepth | 0.41 | 3.11E-01 | 4.05E-02 |
| F.C.L.r.ant. | Width | 0.63 | 1.59E-01 | 1.64E-02 |
| F.C.L.r.ant. | SurfaceArea | 0.87 | 3.28E-04 | 4.09E-02 |
| F.C.L.r.asc. | Length | 0.65 | 5.47E-01 | - |
| F.C.L.r.asc. | MeanDepth | 1.00 | 6.82E-05 | - |
| F.C.L.r.asc. | Width | 0.88 | 6.15E-06 | 4.60E-04 |
| F.C.L.r.asc. | SurfaceArea | 0.68 | 1.12E-02 | 5.45E-03 |
| F.C.L.r.diag. | Length | 0.41 | 9.98E-01 | 3.88E-01 |
| F.C.L.r.diag. | MeanDepth | 0.67 | 8.71E-01 | 1.67E-01 |
| F.C.L.r.diag. | Width | 0.61 | 1.98E-01 | 2.08E-02 |
| F.C.L.r.diag. | SurfaceArea | 0.45 | 9.94E-01 | 8.33E-01 |
| F.C.L.r.retroC.tr. | Length | 0.06 | 7.11E-01 | 4.09E-02 |
| F.C.L.r.retroC.tr. | MeanDepth | 0.84 | 1.97E-04 | 1.78E-02 |
| F.C.L.r.retroC.tr. | Width | 0.64 | 6.49E-02 | 2.01E-02 |
| F.C.L.r.retroC.tr. | SurfaceArea | 0.61 | 1.61E-01 | 1.31E-02 |
| F.C.L.r.sc.ant. | Length | 0.00 | 0.00E+00 | - |
| F.C.L.r.sc.ant. | MeanDepth | -0.03 | 1.36E-01 | - |
| F.C.L.r.sc.ant. | Width | -0.19 | 3.61E-02 | - |
| F.C.L.r.sc.ant. | SurfaceArea | 0.08 | 1.67E-01 | 1.67E-01 |
| F.C.L.r.sc.post. | Length | 0.40 | 7.39E-06 | - |
| F.C.L.r.sc.post. | MeanDepth | 0.60 | 3.46E-02 | 1.67E-01 |
| F.C.L.r.sc.post. | Width | 0.40 | 4.88E-08 | - |
| F.C.L.r.sc.post. | SurfaceArea | 0.03 | 1.46E-01 | 7.22E-03 |
| F.C.M.ant. | Length | 0.45 | 6.72E-01 | 5.54E-04 |
| F.C.M.ant. | MeanDepth | 0.51 | 7.56E-04 | 5.55E-03 |
| F.C.M.ant. | Width | 0.88 | 4.90E-10 | 7.12E-03 |
| F.C.M.ant. | SurfaceArea | 0.93 | 5.67E-04 | 4.00E-03 |
| F.C.M.post. | Length | 1.00 | 5.52E-01 | - |
| F.C.M.post. | MeanDepth | 0.91 | 3.08E-14 | 1.13E-01 |
| F.C.M.post. | Width | 0.90 | 1.53E-14 | 2.37E-02 |
| F.C.M.post. | SurfaceArea | 0.86 | 1.16E-06 | 2.15E-02 |
| F.Cal.ant.-Sc.Cal. | Length | 0.60 | 1.42E-03 | - |
| F.Cal.ant.-Sc.Cal. | MeanDepth | 0.48 | 1.54E-22 | 1.11E-03 |
| F.Cal.ant.-Sc.Cal. | Width | 0.59 | 1.35E-19 | 4.81E-03 |
| F.Cal.ant.-Sc.Cal. | SurfaceArea | 0.53 | 1.19E-27 | 2.71E-06 |
| F.Coll. | Length | 0.68 | 1.81E-01 | 2.00E-02 |
| F.Coll. | MeanDepth | 0.83 | 1.33E-16 | 1.09E-03 |
| F.Coll. | Width | 0.80 | 7.18E-12 | 3.01E-03 |
| F.Coll. | SurfaceArea | 0.88 | 3.26E-05 | 6.97E-06 |
| F.I.P. | Length | 1.00 | 3.04E-07 | - |
| F.I.P. | MeanDepth | 0.87 | 6.61E-12 | 4.52E-02 |
| F.I.P. | Width | 0.98 | 8.90E-33 | 6.88E-02 |
| F.I.P. | SurfaceArea | 0.87 | 1.76E-09 | 1.95E-05 |
| F.I.P.Po.C.inf. | Length | 0.76 | 9.19E-03 | 3.46E-02 |
| F.I.P.Po.C.inf. | MeanDepth | 0.95 | 5.21E-07 | 9.16E-02 |

| | | | | |
|---|---|---|---|---|
| F.I.P.Po.C.inf. | Width | 0.94 | 2.28E-16 | 3.69E-03 |
| F.I.P.Po.C.inf. | SurfaceArea | 0.75 | 8.19E-04 | 1.45E-01 |
| F.I.P.r.int.1 | Length | 0.09 | 4.64E-01 | 1.11E-02 |
| F.I.P.r.int.1 | MeanDepth | 0.39 | 1.83E-01 | - |
| F.I.P.r.int.1 | Width | 0.60 | 7.71E-02 | 9.96E-04 |
| F.I.P.r.int.1 | SurfaceArea | 0.16 | 6.10E-01 | 6.23E-02 |
| F.I.P.r.int.2 | Length | 0.07 | 9.51E-01 | 1.67E-01 |
| F.I.P.r.int.2 | MeanDepth | 0.01 | 9.78E-01 | 4.18E-01 |
| F.I.P.r.int.2 | Width | 0.23 | 9.36E-01 | 9.62E-02 |
| F.I.P.r.int.2 | SurfaceArea | 0.47 | 9.16E-01 | 8.33E-01 |
| F.P.O. | Length | 0.40 | 2.65E-01 | 2.39E-02 |
| F.P.O. | MeanDepth | 0.95 | 7.52E-19 | 3.10E-02 |
| F.P.O. | Width | 0.89 | 2.32E-23 | 5.31E-04 |
| F.P.O. | SurfaceArea | 0.89 | 1.89E-16 | 1.57E-02 |
| Globals | Length | 0.82 | 2.61E-04 | 3.74E-03 |
| Globals | MeanDepth | 0.97 | 7.88E-07 | 1.33E-02 |
| Globals | Width | 0.96 | 4.06E-23 | 1.80E-02 |
| Globals | SurfaceArea | 0.89 | 3.74E-13 | 1.32E-03 |
| INSULA | Length | -0.41 | 9.74E-01 | 1.67E-01 |
| INSULA | MeanDepth | 0.89 | 2.87E-01 | 3.02E-03 |
| INSULA | Width | 0.77 | 2.03E-01 | 1.73E-03 |
| INSULA | SurfaceArea | 0.79 | 1.45E-04 | 1.02E-01 |
| OCCIPITAL | Length | 0.87 | 6.94E-13 | 1.93E-03 |
| OCCIPITAL | MeanDepth | 0.57 | 3.07E-02 | 6.54E-03 |
| OCCIPITAL | Width | 0.87 | 6.14E-22 | 6.59E-06 |
| OCCIPITAL | SurfaceArea | 0.83 | 1.00E-14 | 9.39E-03 |
| S.C. | Length | 1.00 | 1.65E-02 | - |
| S.C. | MeanDepth | 0.92 | 1.96E-13 | 2.50E-02 |
| S.C. | Width | 0.99 | 6.16E-48 | 2.48E-03 |
| S.C. | SurfaceArea | 1.00 | 2.00E-18 | - |
| S.C.LPC. | Length | -0.67 | 1.59E-01 | - |
| S.C.LPC. | MeanDepth | 0.01 | 8.33E-01 | 1.67E-01 |
| S.C.LPC. | Width | -0.33 | 9.99E-01 | 2.07E-02 |
| S.C.LPC. | SurfaceArea | -0.36 | 9.13E-01 | 1.67E-01 |
| S.C.sylvian. | Length | 0.88 | 4.60E-02 | 2.10E-03 |
| S.C.sylvian. | MeanDepth | 0.69 | 2.02E-01 | 1.59E-04 |
| S.C.sylvian. | Width | 0.86 | 2.97E-03 | 4.34E-03 |
| S.C.sylvian. | SurfaceArea | 1.00 | 2.06E-03 | - |
| S.Call. | Length | 0.85 | 2.49E-17 | 9.19E-04 |
| S.Call. | MeanDepth | 0.67 | 7.94E-08 | 2.22E-04 |
| S.Call. | Width | 0.82 | 8.25E-05 | 7.26E-03 |
| S.Call. | SurfaceArea | 0.81 | 6.80E-23 | 3.79E-03 |
| S.Cu. | Length | 0.08 | 1.53E-01 | 2.14E-06 |
| S.Cu. | MeanDepth | 0.81 | 5.55E-04 | 3.02E-05 |
| S.Cu. | Width | 1.00 | 1.30E-06 | - |
| S.Cu. | SurfaceArea | 0.89 | 7.72E-02 | 1.20E-01 |
| S.F.inf. | Length | 0.33 | 1.76E-02 | - |
| S.F.inf. | MeanDepth | 0.90 | 1.09E-13 | 2.25E-03 |
| S.F.inf. | Width | 0.85 | 8.98E-14 | 4.24E-03 |
| S.F.inf. | SurfaceArea | 0.85 | 3.75E-07 | 1.99E-02 |
| S.F.inf.ant. | Length | 1.00 | 1.49E-02 | - |
| S.F.inf.ant. | MeanDepth | 0.45 | 7.60E-03 | 4.26E-02 |
| S.F.inf.ant. | Width | 0.69 | 4.23E-03 | 7.49E-02 |
| S.F.inf.ant. | SurfaceArea | 0.90 | 3.19E-03 | 9.97E-02 |
| S.F.int. | Length | 0.90 | 6.28E-07 | 2.08E-01 |
| S.F.int. | MeanDepth | 0.72 | 1.79E-06 | 4.58E-02 |
| S.F.int. | Width | 0.91 | 7.71E-12 | 4.40E-02 |

| | | | | |
|---|---|---|---|---|
| S.F.int. | SurfaceArea | 0.84 | 1.07E-06 | 9.58E-03 |
| S.F.inter. | Length | 0.90 | 5.18E-03 | 5.64E-03 |
| S.F.inter. | MeanDepth | 0.87 | 1.73E-04 | 1.83E-03 |
| S.F.inter. | Width | 0.84 | 1.69E-19 | 3.39E-05 |
| S.F.inter. | SurfaceArea | 1.00 | 9.85E-04 | - |
| S.F.marginal. | Length | 0.56 | 7.68E-03 | - |
| S.F.marginal. | MeanDepth | 0.69 | 9.56E-01 | 4.60E-01 |
| S.F.marginal. | Width | 1.00 | 2.91E-02 | - |
| S.F.marginal. | SurfaceArea | 0.63 | 1.64E-01 | 2.23E-02 |
| S.F.median. | Length | 0.96 | 5.43E-06 | 1.06E-02 |
| S.F.median. | MeanDepth | 0.66 | 1.60E-02 | 9.28E-04 |
| S.F.median. | Width | 0.92 | 1.74E-13 | 5.00E-06 |
| S.F.median. | SurfaceArea | 0.88 | 4.54E-05 | 2.62E-02 |
| S.F.orbitaire. | Length | 0.67 | 6.82E-01 | 3.42E-01 |
| S.F.orbitaire. | MeanDepth | 0.70 | 5.53E-01 | 1.09E-01 |
| S.F.orbitaire. | Width | 0.72 | 7.42E-02 | 1.40E-04 |
| S.F.orbitaire. | SurfaceArea | 0.68 | 2.82E-01 | 7.84E-02 |
| S.F.polaire.tr. | Length | 0.98 | 6.80E-10 | 1.49E-02 |
| S.F.polaire.tr. | MeanDepth | 0.84 | 9.61E-04 | 7.71E-05 |
| S.F.polaire.tr. | Width | 0.82 | 9.68E-09 | 6.82E-03 |
| S.F.polaire.tr. | SurfaceArea | 0.98 | 2.22E-08 | 1.32E-01 |
| S.F.sup. | Length | 0.17 | 3.98E-01 | 1.67E-01 |
| S.F.sup. | MeanDepth | 0.93 | 2.72E-15 | 2.18E-02 |
| S.F.sup. | Width | 0.90 | 7.50E-18 | 2.15E-04 |
| S.F.sup. | SurfaceArea | 0.82 | 4.98E-03 | 8.51E-08 |
| S.Li.ant. | Length | 0.71 | 6.80E-02 | 2.55E-07 |
| S.Li.ant. | MeanDepth | 0.57 | 5.28E-02 | 3.21E-04 |
| S.Li.ant. | Width | 0.73 | 8.24E-03 | 1.08E-01 |
| S.Li.ant. | SurfaceArea | 0.51 | 2.06E-01 | 1.32E-03 |
| S.Li.post. | Length | 0.63 | 4.34E-01 | 4.88E-01 |
| S.Li.post. | MeanDepth | 0.85 | 1.82E-03 | 3.75E-05 |
| S.Li.post. | Width | 0.96 | 1.58E-02 | 4.84E-03 |
| S.Li.post. | SurfaceArea | 0.52 | 7.06E-01 | 1.94E-01 |
| S.O.p. | Length | 0.51 | 7.92E-01 | 9.28E-03 |
| S.O.p. | MeanDepth | 0.81 | 5.03E-02 | 3.77E-03 |
| S.O.p. | Width | 1.00 | 2.38E-05 | - |
| S.O.p. | SurfaceArea | 0.70 | 4.79E-02 | 2.43E-02 |
| S.O.T.lat.ant. | Length | 0.93 | 7.51E-05 | 7.15E-02 |
| S.O.T.lat.ant. | MeanDepth | 0.92 | 1.59E-04 | 3.40E-02 |
| S.O.T.lat.ant. | Width | 0.62 | 6.72E-04 | 1.97E-02 |
| S.O.T.lat.ant. | SurfaceArea | 0.88 | 1.56E-05 | 1.32E-01 |
| S.O.T.lat.int. | Length | 0.72 | 3.13E-01 | - |
| S.O.T.lat.int. | MeanDepth | 0.02 | 1.00E+00 | 1.95E-01 |
| S.O.T.lat.int. | Width | 0.46 | 2.98E-01 | - |
| S.O.T.lat.int. | SurfaceArea | 0.63 | 8.20E-01 | 4.54E-01 |
| S.O.T.lat.med. | Length | 0.60 | 7.58E-01 | 1.67E-01 |
| S.O.T.lat.med. | MeanDepth | 0.18 | 1.86E-01 | 8.15E-02 |
| S.O.T.lat.med. | Width | 0.51 | 3.28E-01 | 1.03E-04 |
| S.O.T.lat.med. | SurfaceArea | 0.67 | 3.70E-01 | 9.53E-02 |
| S.O.T.lat.post. | Length | 0.71 | 7.72E-01 | 1.44E-01 |
| S.O.T.lat.post. | MeanDepth | 0.88 | 1.06E-03 | 3.40E-02 |
| S.O.T.lat.post. | Width | 0.68 | 1.33E-02 | 1.44E-02 |
| S.O.T.lat.post. | SurfaceArea | 0.75 | 2.10E-01 | 4.01E-01 |
| S.Olf. | Length | 0.82 | 1.27E-05 | 7.77E-02 |
| S.Olf. | MeanDepth | 0.88 | 1.12E-14 | 4.61E-02 |
| S.Olf. | Width | 0.76 | 5.88E-05 | 5.15E-02 |
| S.Olf. | SurfaceArea | 0.92 | 1.00E-11 | 5.57E-05 |

| | | | | |
|---|---|---|---|---|
| S.Or. | Length | 0.98 | 1.97E-05 | 8.15E-03 |
| S.Or. | MeanDepth | 0.85 | 3.77E-12 | 5.00E-03 |
| S.Or. | Width | 1.00 | 2.18E-13 | - |
| S.Or. | SurfaceArea | 0.94 | 3.66E-10 | 1.67E-02 |
| S.p.C. | Length | 0.67 | 3.95E-02 | 1.64E-02 |
| S.p.C. | MeanDepth | 0.24 | 4.45E-01 | 1.16E-02 |
| S.p.C. | Width | 0.41 | 5.40E-01 | 4.52E-03 |
| S.p.C. | SurfaceArea | 0.32 | 3.22E-01 | 1.07E-02 |
| S.Pa.int. | Length | 1.00 | 1.26E-04 | - |
| S.Pa.int. | MeanDepth | 0.99 | 1.12E-06 | 1.43E-02 |
| S.Pa.int. | Width | 0.95 | 4.00E-12 | 5.85E-02 |
| S.Pa.int. | SurfaceArea | 1.00 | 1.79E-06 | - |
| S.Pa.sup. | Length | 1.00 | 5.62E-02 | - |
| S.Pa.sup. | MeanDepth | 0.37 | 2.01E-02 | - |
| S.Pa.sup. | Width | 1.00 | 1.10E-04 | - |
| S.Pa.sup. | SurfaceArea | 0.70 | 1.73E-01 | 1.93E-02 |
| S.Pa.t. | Length | 1.00 | 3.82E-01 | - |
| S.Pa.t. | MeanDepth | 0.01 | 4.61E-01 | 1.67E-01 |
| S.Pa.t. | Width | 0.49 | 5.96E-01 | 5.04E-02 |
| S.Pa.t. | SurfaceArea | -0.02 | 8.16E-01 | 1.72E-02 |
| S.Pe.C.inf. | Length | 0.94 | 7.44E-02 | 1.43E-02 |
| S.Pe.C.inf. | MeanDepth | 0.87 | 1.10E-02 | 6.55E-02 |
| S.Pe.C.inf. | Width | 1.00 | 3.16E-11 | - |
| S.Pe.C.inf. | SurfaceArea | 0.83 | 4.49E-02 | 2.43E-03 |
| S.Pe.C.inter. | Length | 0.71 | 5.61E-01 | 2.77E-04 |
| S.Pe.C.inter. | MeanDepth | 0.99 | 2.82E-08 | 1.90E-02 |
| S.Pe.C.inter. | Width | 0.99 | 3.42E-13 | 1.24E-02 |
| S.Pe.C.inter. | SurfaceArea | 0.64 | 1.02E-01 | - |
| S.Pe.C.marginal. | Length | -0.49 | 7.08E-01 | 1.67E-01 |
| S.Pe.C.marginal. | MeanDepth | 0.96 | 1.08E-01 | 1.67E-01 |
| S.Pe.C.marginal. | Width | 0.94 | 4.74E-05 | 2.62E-03 |
| S.Pe.C.marginal. | SurfaceArea | -0.38 | 2.23E-01 | - |
| S.Pe.C.median. | Length | 0.90 | 3.45E-01 | 1.21E-02 |
| S.Pe.C.median. | MeanDepth | 0.76 | 5.09E-01 | 2.02E-02 |
| S.Pe.C.median. | Width | 1.00 | 4.51E-05 | - |
| S.Pe.C.median. | SurfaceArea | 0.89 | 1.82E-01 | 6.23E-03 |
| S.Pe.C.sup. | Length | 1.00 | 4.17E-03 | - |
| S.Pe.C.sup. | MeanDepth | 0.96 | 1.18E-02 | 1.08E-02 |
| S.Pe.C.sup. | Width | 1.00 | 4.45E-07 | - |
| S.Pe.C.sup. | SurfaceArea | 0.86 | 3.65E-03 | 1.47E-03 |
| S.Po.C.sup. | Length | 0.96 | 2.14E-01 | 1.67E-01 |
| S.Po.C.sup. | MeanDepth | 0.84 | 1.09E-04 | 1.81E-04 |
| S.Po.C.sup. | Width | 1.00 | 8.77E-21 | - |
| S.Po.C.sup. | SurfaceArea | 0.65 | 7.22E-02 | 9.25E-08 |
| S.R.inf. | Length | 1.00 | 1.52E-03 | - |
| S.R.inf. | MeanDepth | 1.00 | 1.45E-02 | - |
| S.R.inf. | Width | 0.79 | 1.14E-01 | 8.68E-04 |
| S.R.inf. | SurfaceArea | 1.00 | 2.88E-04 | - |
| S.Rh. | Length | 0.66 | 3.02E-02 | 6.20E-02 |
| S.Rh. | MeanDepth | 0.76 | 6.27E-03 | 6.49E-02 |
| S.Rh. | Width | 0.84 | 2.22E-04 | 1.41E-07 |
| S.Rh. | SurfaceArea | 0.44 | 1.90E-01 | 2.81E-03 |
| S.s.P. | Length | 0.76 | 8.70E-04 | 4.81E-03 |
| S.s.P. | MeanDepth | 1.00 | 4.31E-17 | - |
| S.s.P. | Width | 0.93 | 6.95E-11 | 2.01E-04 |
| S.s.P. | SurfaceArea | 0.93 | 6.32E-08 | 4.24E-02 |
| S.T.i.ant. | Length | 0.95 | 2.79E-04 | 1.10E-02 |

| | | | | |
|---|---|---|---|---|
| S.T.i.ant. | MeanDepth | 0.94 | 2.45E-06 | 6.85E-02 |
| S.T.i.ant. | Width | 0.96 | 2.79E-19 | 5.69E-02 |
| S.T.i.ant. | SurfaceArea | 0.97 | 1.34E-09 | 9.53E-02 |
| S.T.i.post. | Length | 0.20 | 5.87E-01 | - |
| S.T.i.post. | MeanDepth | 0.80 | 1.34E-05 | 1.97E-02 |
| S.T.i.post. | Width | 0.88 | 1.04E-12 | 2.13E-02 |
| S.T.i.post. | SurfaceArea | 0.82 | 3.14E-02 | 9.78E-02 |
| S.T.pol. | Length | 0.72 | 3.27E-04 | 2.93E-02 |
| S.T.pol. | MeanDepth | 0.67 | 2.48E-01 | 2.65E-01 |
| S.T.pol. | Width | 0.83 | 1.01E-08 | 1.22E-03 |
| S.T.pol. | SurfaceArea | 0.70 | 3.32E-04 | 5.86E-03 |
| S.T.s. | Length | 0.60 | 4.76E-01 | 4.70E-03 |
| S.T.s. | MeanDepth | 0.81 | 1.33E-09 | 1.57E-04 |
| S.T.s. | Width | 0.83 | 4.21E-12 | 1.87E-03 |
| S.T.s. | SurfaceArea | 0.73 | 3.99E-03 | 1.62E-02 |
| S.T.s.ter.asc.ant. | Length | 0.64 | 4.70E-01 | 1.67E-01 |
| S.T.s.ter.asc.ant. | MeanDepth | 0.53 | 6.56E-01 | 1.45E-01 |
| S.T.s.ter.asc.ant. | Width | 0.96 | 2.71E-04 | 8.00E-03 |
| S.T.s.ter.asc.ant. | SurfaceArea | 0.32 | 9.16E-01 | 8.33E-01 |
| S.T.s.ter.asc.post. | Length | -0.09 | 7.87E-01 | - |
| S.T.s.ter.asc.post. | MeanDepth | 0.83 | 2.94E-03 | 3.14E-02 |
| S.T.s.ter.asc.post. | Width | 0.72 | 6.89E-04 | 7.02E-03 |
| S.T.s.ter.asc.post. | SurfaceArea | 0.59 | 3.86E-01 | 5.41E-02 |
